# Large-Scale Chemical-Genetic Interaction Profiling Identifies a Novel Small-Molecule Inhibitor of *Mycobacterium tuberculosis* Polyketide Synthase 13

**DOI:** 10.64898/2026.02.06.704361

**Authors:** James E. Gomez, Matthew Y. Solomon, Diana K. Hunt, Emily J. Geddes, Austin N. Bond, Cong Liu, Rebecca J. Ulrich, Pooja V. Chaudhary, Sulyman Barkho, Deborah T. Hung

## Abstract

PROSPECT (PRimary screening Of Strains to Prioritize Expanded Chemistry and Targets) is an antimicrobial discovery platform based on chemical-genetic interaction (CGI) profiling of compounds against a pool of *Mycobacterium tuberculosis* (*Mtb)* hypomorphs, each depleted of an essential gene. From prior screening data, we have now identified a novel N-oxolan-3-yl pyrazole carboxamide inhibitor (BRD1554) that had increased, selective activity against strains depleted of polyketide synthase 13 (Pks13), an essential enzyme in mycolic acid synthesis, and Rv2581c, an uncharacterized protein similar to glyoxylase II enzymes. Perturbagen CLass (PCL) analysis, a reference-based approach to mechanism of action (MOA) assignment from PROSPECT, predicted Pks13, a polyketide synthase with five catalytic domains responsible for the terminal condensation step in mycolic acid biosynthesis, was the likely target, potentially implicating the thioesterase domain. We synthesized a more active analogue while assigning the absolute stereochemistry of the active diastereomer, resulting in 1554-06-3R,4S with an MIC_90_ of 3.0 µM against *Mtb* H37Rv. Exposure to 1554-06 led to the upregulation of the *pks13* operon along with the *iniBAC* operon and other genes linked to mycolic acid synthesis. Isolation of mutants resistant to 1554-06 revealed single nucleotide polymorphisms in the thioesterase domain of Pks13. Finally, we biochemically confirmed that 1554-06 inhibits the activity of recombinant Pks13 thioesterase domain, with computational docking of 1554-06 steroisomers consistent with the stereospecific activity seen in whole cell assays. We found unique chemical genetic interactions between inhibitors of the different Pks13 domains and different detoxifying enzymes of *Mtb*, thus revealing novel gene-gene interactions. These results highlight how PROSPECT can not only immediately reveal, with domain-level resolution, the MOA of new whole-cell active chemical inhibitors of *Mtb*, allowing the integration of biological insight into compound triage and accelerated early development, but can also illuminate genetic interactions linked to those mechanisms that could inform predictions of synergy for antitubercular drug development.

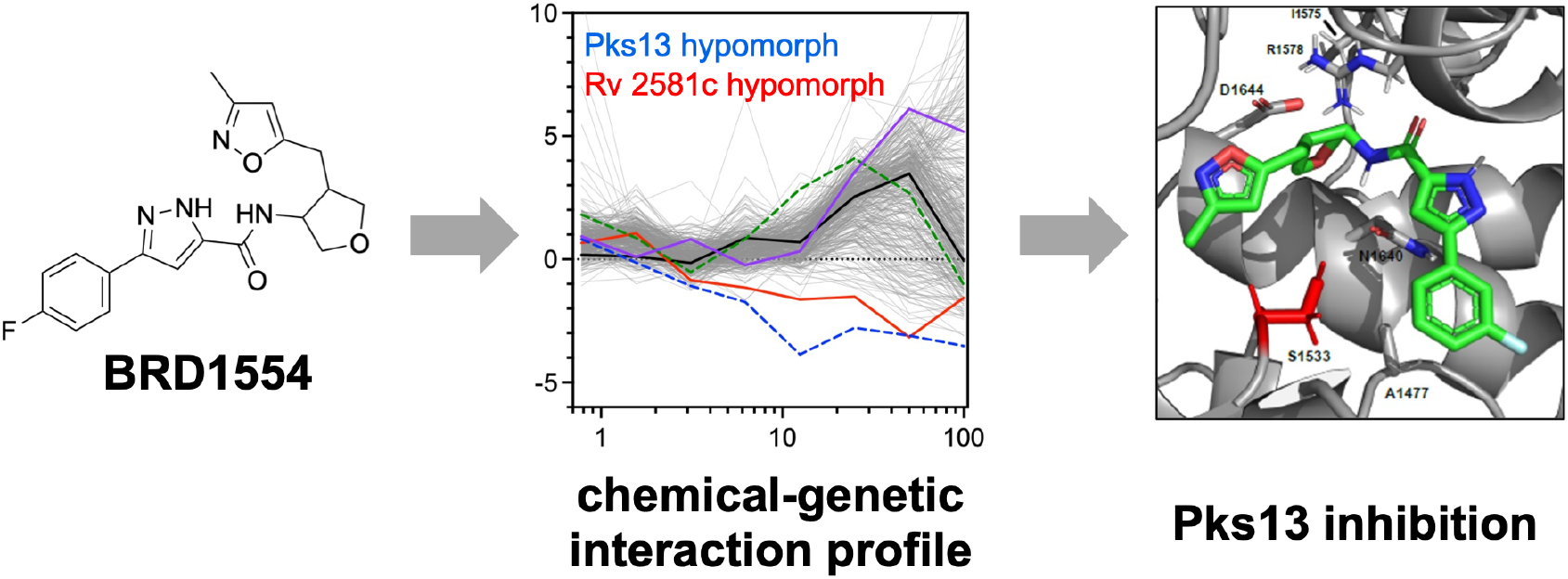

---

*Mycobacterium tuberculosis* (*Mtb*) continues to kill over a million people globally every year^1^, with a concerning rise in drug resistance. Nearly 5% of new infections now involve multidrug-resistant organisms^1^. To address these issues, efforts have been invested in discovering and developing new drugs targeting novel vulnerabilities of *Mtb*, including bedaquiline^2^, delamanid^3^, and pretomanid^4^. New regimens incorporating these drugs have enabled tremendous advances in both treating drug-resistant tuberculosis and reducing the length of time required to achieve lasting cures^5^. Given that tuberculosis must be treated by combinations of drugs to avoid the emergence of resistance, new agents working in combination are needed to achieve rapid, reproducible clearance.

Recently, we developed PROSPECT (PRimary screening Of Strains to Prioritize Expanded Chemistry and Targets), a novel systems chemical biology strategy that integrates whole-cell small molecule discovery with immediate mechanism of action prediction^6^. PROSPECT measures the activity of a small molecule against a pool of hypomorphic (barcoded) bacterial strains, each engineered to be depleted of a different essential protein^7^. Collectively, the interactions between a molecule and the entire pool of hypomorphic mutants define a chemical-genetic interaction (CGI) profile. When applied to chemical libraries, active small molecule discovery can be coupled to mechanism of action (MOA) information based on the behavior of individual strains in response to the molecule, with hypersensitized strains being most informative. The pooled PROSPECT screen thus has the potential to identify inhibitors that target a wide range of protein targets in a single screen, potentially leveraging the hundreds of essential proteins within a bacterial species such as *Mtb*. Thus, the resulting dataset can be mined to discover active molecules with different MOAs. Further, because depletion of the direct target or related pathway proteins can make mutants hypersensitive to certain molecules, PROSPECT can lead to the discovery of compounds that lack initial potent wild-type activity, but with optimization through medicinal chemistry can be converted to potent antibacterial molecules. Taken together, PROSPECT can alter the selection and prioritization of molecules emerging from an antibacterial whole-cell screen through its early integration of MOA information or its identification of wild-type inactive or weakly active molecules with intriguing MOA.

Cell wall biosynthesis inhibitors can be readily identified by PROSPECT^8^. The enzymes that synthesize cell wall components are a common target of antimicrobial drugs for a variety of reasons, including the frequent bactericidal consequence of a weakened cell wall and the accessibility of these enzymes at the outer edges of the cell. The lack of homologs of these enzymes in mammalian systems provides high selectivity for bacteria. In particular, *Mtb* has a broad range of enzymes involved in the synthesis of its complex cell wall^9,10^, which is composed not only of peptidoglycan, but numerous covalently and non-covalently attached glycopolymers and lipids unique to *Mtb* and other closely related members of the Mycobacteriaceae. Perhaps the best studied of the *Mtb* cell wall components are mycolic acids, which are large branched-chain lipids that are a major component of the exterior portion of the cell wall. These molecules are unique to the Mycobacteriaceae, and their synthesis is a target of several drugs in clinical use, including isoniazid, ethionamide^11^, nitroimidazoles^12^, as well as a number of drugs in preclinical development, thus reinforcing the value of these targets.

Here we mined the extensive PROSPECT dataset that we have previously generated to identify a novel small molecule targeting the essential cell wall enzyme Pks13. Pks13 is a large, multidomain type I polyketide synthase that catalyzes the terminal and obligate condensation step in mycolic acid biosynthesis in *Mtb*^13,14^. Structurally, Pks13 is organized into five core catalytic domains: an acyltransferase domain, two acyl carrier protein domains (ACP), a ketosynthase domain, and a thioesterase domain. These domains function together to couple two distinct lipid precursors to produce mycolic acid. The Pks13 acyltransferase domain loads a β-carboxylated C_26_fatty acyl substrate onto the internal ACP, while FadD32 loads a C_56_meromycolate chain onto the N-terminal ACP. The ketosynthase domain then catalyzes a Claisen condensation of these lipid precursors to generate the full-length mycolic acid scaffold, a critical structural component of the mycobacterial cell envelope, tethered to the ACP. Finally, the newly formed mycolic acid is released from the ACP by the Pks13 thioesterase domain by transferring it onto trehalose^15^, forming trehalose monomycolate (TMM), a key cell wall intermediate that is subsequently exported by MmpL3^16^ and further processed into trehalose dimycolate (TDM) or incorporated into mycolyl-arabinogalactan^17^.

From a therapeutic perspective, Pks13 represents a high-value target within the mycolic acid biosynthetic network given its terminal role in mycomembrane biosynthesis. Multiple small-molecule inhibitors have now been described that target distinct functional domains of the enzyme, including compounds acting on the N-terminal ACP domain^18,19^, including a covalent ACP inhibitor, as well as inhibitors of the thioesterase domain^20-25^, which is responsible for product release. Here, we describe the discovery of screening hit BRD1554, a novel Pks13 thioesterase inhibitor, using Perturbagen CLass (PCL) analysis of PROSPECT data, a reference-based method of MOA assignment, along with the further characterization of BRD1554 and expansion via early medicinal chemistry, demonstrating stereoselective engagement with this target. We also reveal unique chemical genetic interactions between inhibitors of the different domains and different detoxifying enzymes of *Mtb*, thereby illustrating the power of PROSPECT to reveal novel gene-gene interactions in the cell.

## RESULTS AND DISCUSSION

### Identification of BRD1554 (1) from PROSPECT screening data

We had previously screened a 50,000 compound library from Charles River Labs using PROSPECT^6^, a multiplexed screening modality that uses induced proteolysis of essential *Mtb* proteins to reduce their intracellular levels, making the corresponding barcoded hypomorphic strains vulnerable to inhibitors that target these proteins, their pathways, or related vulnerabilities. Mining data from this previous screen, we identified an N-(oxolan-3-yl)-1H-pyrazole-5-carboxamide, BRD1554 (**1**) (Figure 1A), as a molecule that moderately suppressed the growth of the majority of strains when assayed at 100 µM but had no clear IC50 against wild-type H37Rv in PROSPECT (Figure 1B). However, we observed that two strains were highly and specifically sensitized: the Pks13 hypomorph and the Rv2581c hypomorph, which has diminished levels of a putative uncharacterized glyoxylase found by Bosch et al. to be among the most vulnerable targets in *Mtb*^26^. Using PCL analysis, a reference based strategy for MOA assignment that predicts MOA from hit compounds by comparing their CGI profiles to those derived from known inhibitors^8^, we found BRD1554 (**1**) to be a high confidence match to a reference cluster formed by several Pks13 inhibitors including several benzofurans (TAM1, TAM3, TAM9)^20^ along with a thiophene (TP4)^19^ (Figure 1C).

**Figure 1.**
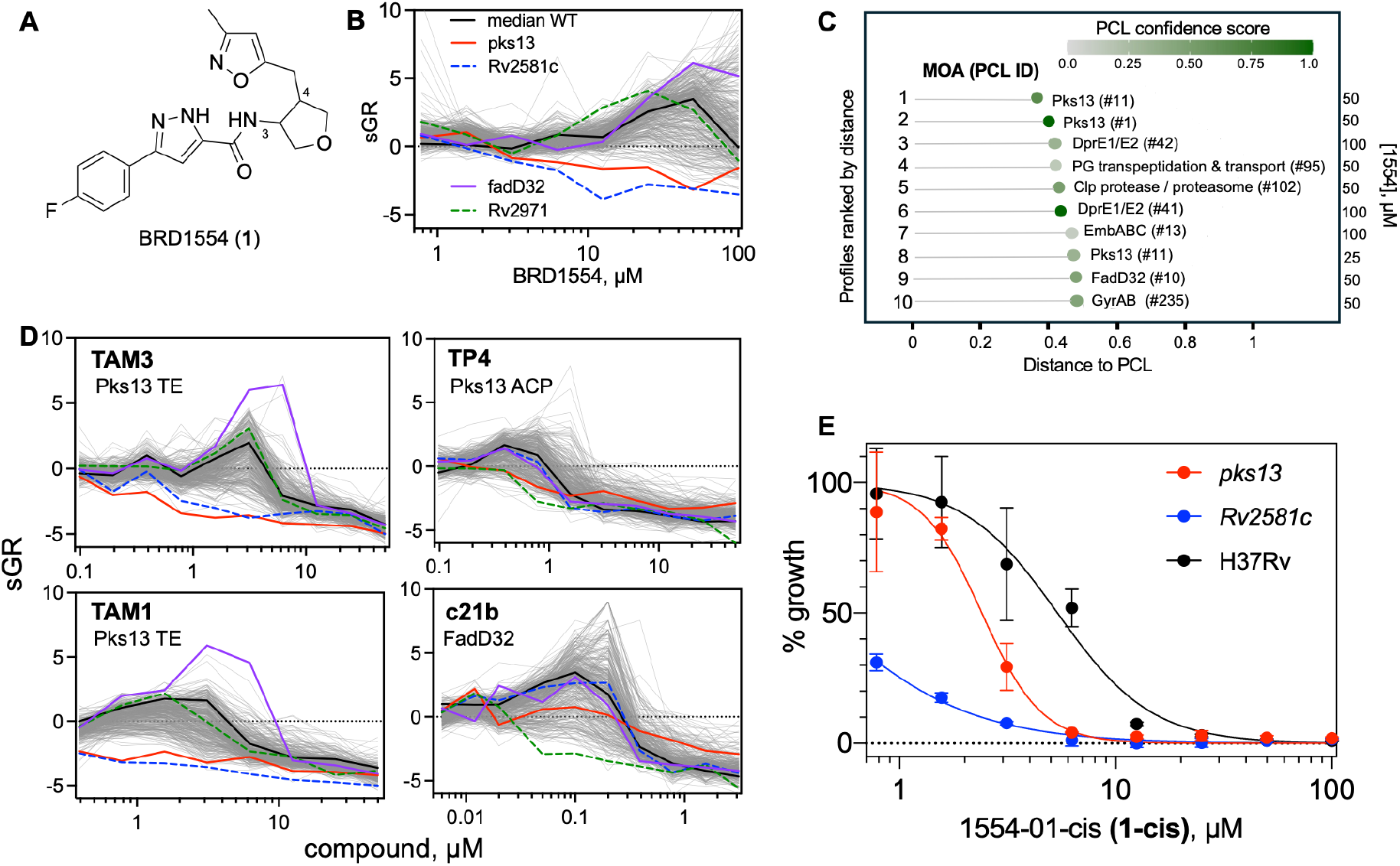
Discovery and confirmation of a novel scaffold targeting Pks13. (A) Structure of the original hit molecule BRD1554 (**1**). (B) Dose-response PROSPECT screening data showing the behavior of 361 strains (grey lines) in response to BRD1554. The y-axis measures the normalized growth rate of each strain (0 = no inhibition, negative values = strong inhibition) over the 14-day assay. The median growth of 7 unique barcoded WT H37Rv clones is highlighted by a black line, and strains depleted of Pks13 and Rv2581c are highlighted in red and dashed blue, respectively. Strains depleted of FadD32 and Rv2971 are shown in purple and dashed green, respectively. The datapoints are the average of 2 replicate screening wells. (C) Perturbagen CLass (PCL) analysis of dose-response CGI profiling data from BRD1554, showing top 10 closest matches in rank order based on distance (closer = greater median Pearson correlation). The compound dose providing the PCL match is given on the right. Darkest green = highest confidence predictions, based on how well that distance successfully predicted the MOA in training^8^. (D) PROSPECT CGI profiles of inhibitors of Pks13 thioesterase (TE; TAM1 and TAM3), ACP (TP4), and FadD32 (c21b). Replicates and strains are as in 2B. (E) Dose-response curves for 1554-01-cis **(1-cis)** in *Mtb* H37Rv parent and hypomorphs of Pks13 and Rv2581c, each grown alone in an OD_600_-based assay. Data are average and standard deviation of 2 replicates.

PCLs are clusters of correlated dose-specific CGI profiles from compounds sharing a common MOA^8^. PCLs can contain multiple doses (often adjacent) of the same compound if the effects on the CGI profile are highly correlated; likewise, the same compound can fall into multiple PCLs based on dose-dependent activity (*e*.*g*., sub-MIC doses elicit different responses than doses near the MIC). In this case, the Pks13 PCL #11 is formed from four dose-specific CGI profiles from TAM1 and TAM3, both benzofuran Pks13 thioesterase inhibitors, with CGI profiles featuring Pks13 and Rv2581c, a putative glyoxylase II^27^, as the two most sensitized strains based on median rank across the 4 treatments (Figure S1A). In contrast, Pks13 PCL #1 features 7 dose-specific CGI profiles from three thioesterase inhibitors (TAM1, TAM3, and TAM9)^20^ and the ACP inhibitor TP4^19^ (Figure S1B). PCL #1 thus contains both thioesterase and ACP inhibitors, indicating that their respective CGI profiles share common features despite mechanistic differences in their mode of Pks13 inhibition. Meanwhile, PCL#11 includes only thioesterase but not ACP inhibitors, suggesting potential unmasking of differences in their chemical genetic interactions with other hypomorphs as a function of specific domain inhibition. The sensitization of Rv2581c is specific to thioesterase inhibition (Figure 1D), while ACP inhibition shows sensitization of Rv2971, an alpha-keto reductase induced by thio-specific oxidative stress^28^ and cAMP^29^. Of note, Rv2971 sensitization is also a prominent feature of FadD32 inhibitors (*i*.*e*, c21b)^30^, which like TP4, block the transfer of meromycolic acids to the ACP of Pks13. Thus, PCL analysis of PROSPECT data can discern different activities of multidomain enzymes.

### Activity Confirmation and Assignment of Absolute Stereochemistry

The BRD1554 screening hit **(1)** was a mixture of 4 stereoisomers involving the stereocenters at positions 3 and 4 of the tetrahydrofuran ring. We resynthesized the hit as a mixture of all four stereoisomers (1554-01 (**1-syn**), 5:2 cis:trans), and the activity of the mixture was confirmed against wild-type *Mtb* H37Rv with an MIC_90_of 24.7 µM (Table S1). Isolation of the cis isomers of 1554-01 (1554-01-cis (**1-cis**)) led to a twofold improvement in potency (13.3 µM). Recapitulating PROSPECT results, the Pks13 and Rv2581c hypomorphs were hypersensitized to **1-cis** relative to wild-type H37Rv, with the Pks13 and Rv2581c hypomorphs showing a 2.8-fold (4.8 µM) and 5.5-fold (2.4 µM) reduction in MIC_90_relative to wild-type, respectively (Figure 1E). Of note, **1-cis** had no measurable toxicity against HepG2 cells up to 120 µM (Table S2).

Seeking compounds with improved potency, we synthesized analogues with alternate substitutions on the phenyl ring at the 5-position of the pyrazole. We found that shifting the fluoro substituent from the 4-position of the phenyl ring in the resynthesized hit (1554-01 (**1-syn**)) to the 3-position (1554-06 (**5**)) improved the MIC_90_of the mixture from 24.7 µM to 3.2 µM. Separation of the diastereomers of **5** into cis and trans mixtures indicated higher activity for the cis mixture (MIC_90_= 3.0 µM for cis, 68.2 µM for trans), while isolation of all four individual stereoisomers indicated further stereospecific activity (Figure 2), with one highly active cis stereoisomer (1554-06-cis2 (**5b**), MIC_90_3.0 µM), one less active trans stereoisomer (1554-06-trans1 (**5c**), MIC_90_53.7 µM), and two stereoisomers (1554-06-cis1 (**5a**) and 1554-06-trans2 (**5d**)) that were completely inactive (MIC_90_>100 µM). We determined the absolute stereochemistry of the isomers using X-ray crystallography (Figure S2A and B), and assigned the absolute stereochemistry of the most active stereoisomer (**5b**) as the (3R,4S) isomer, thus elucidating the structure of **5b** as 3-(3-fluorophenyl)-*N*-((3*R*,4*S*)-4-((3-methylisoxazol-5-yl)methyl)tetrahydrofuran-3-yl)-1*H*-pyrazole-5-carboxamide. Subsequent work focused on **5b** (3R,4S) for mechanistic studies.

**Figure 2.**
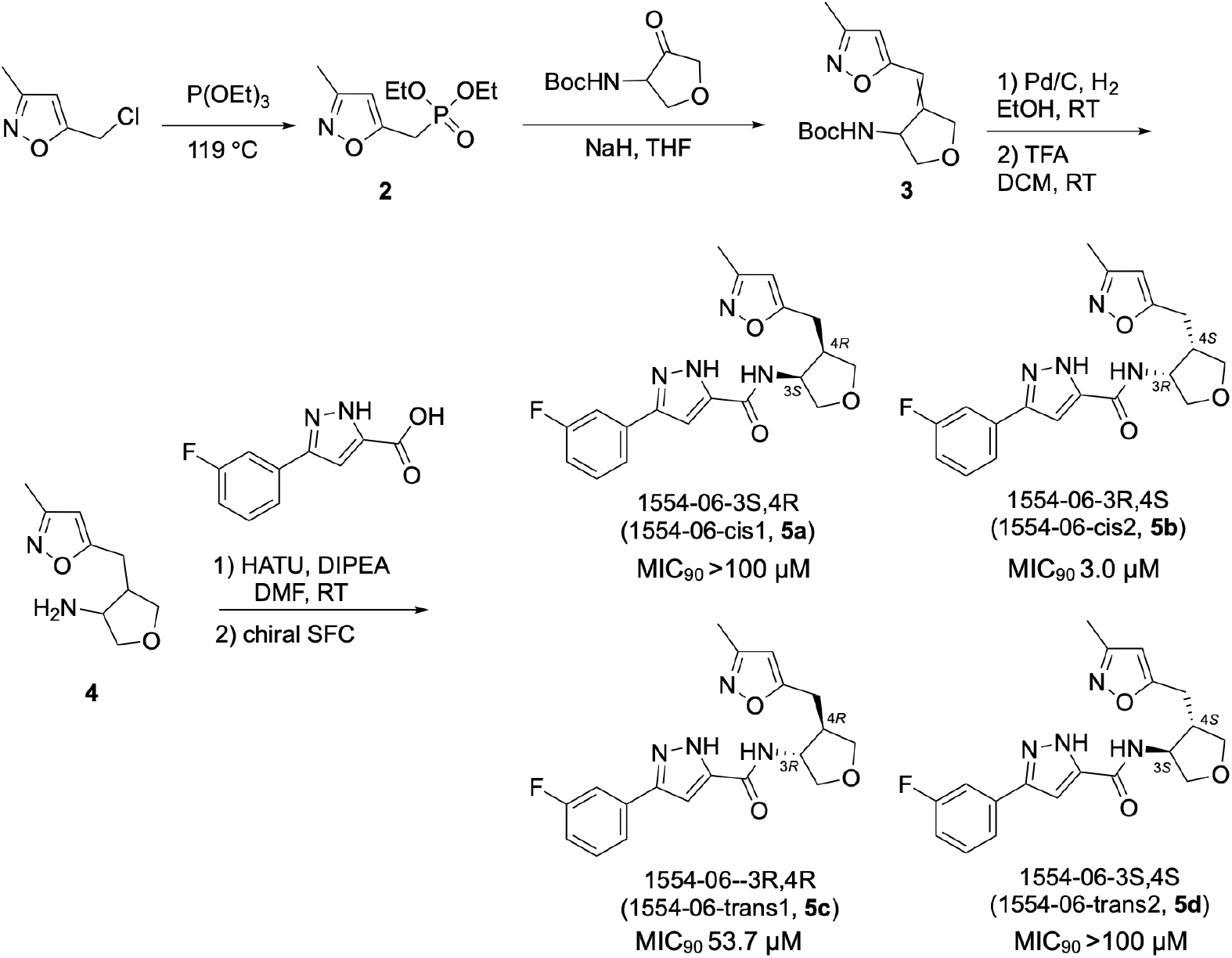
Synthesis and characterization of 1554-06 diastereomers. Synthetic scheme, structures, absolute stereochemical assignment, and H37Rv MIC90s of the 4 purified stereoisomers of 1554-006 are shown. The absolute stereochemistry of cis1 and trans2 stereoisomers was determined by x-ray crystallography, and the stereochemistry of cis2 and trans1 were inferred from those results.

### Transcriptomic Profiling of 1554

Given the PCL prediction, we sought supporting evidence that **5b** indeed inhibits Pks13. We first measured the transcriptional response of *Mtb* to **5b** (Figure 3A). Following 6 hours of exposure to 4X the MIC_90_, we saw strong upregulation of the *iniBAC* operon and genes involved in mycolic acid biosynthesis, including *fas, kasA* and *kasB*. The operon including *fadD32, pks13*, and *accD4*, encoding the final 3 enzymatic activities involved in the condensation of C_54-56_and C_26_fatty acids into mycolic acids, was also significantly upregulated. We saw no upregulation of *Rv2581c*, despite the sensitization of the corresponding hypomorph. Thus, the transcriptional response was consistent with inhibition of mycolic acid biosynthesis via Pks13. Isocitrate lyase (*icl1*), required for fatty acid degradation, was also highly upregulated, consistent with the cell’s need to recycle the long fatty acyl chains that would accumulate upon Pks13 inhibition.

**Figure 3.**
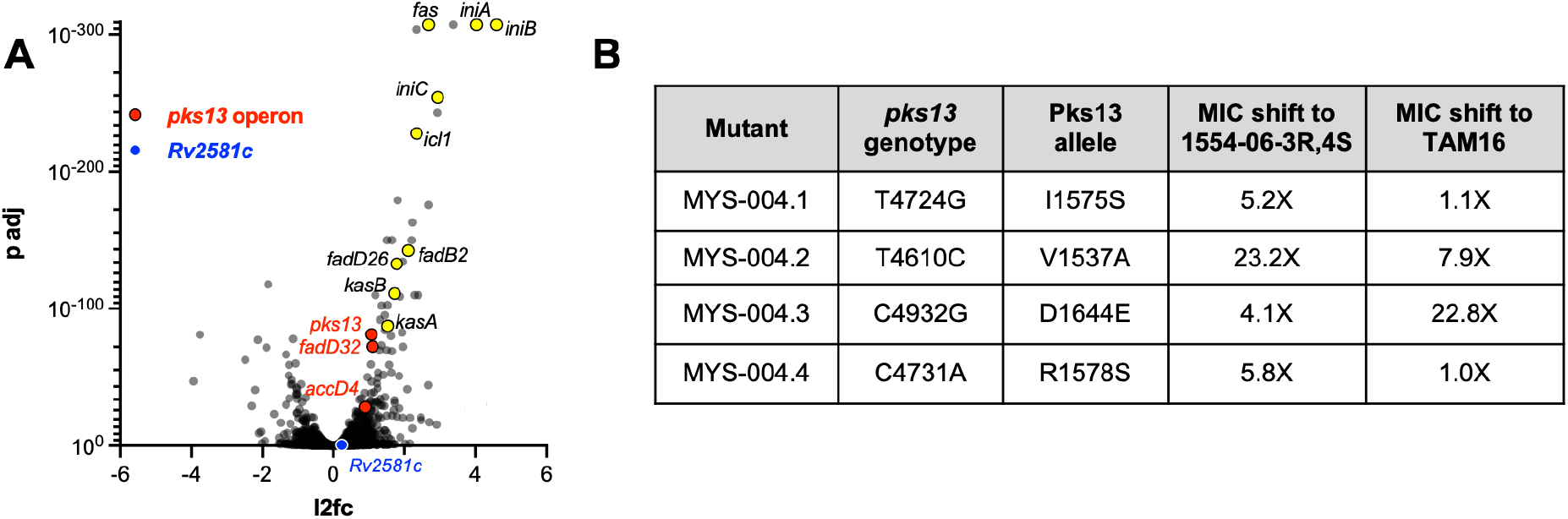
Genetic support for Pks13 as the target of the 1554 series. (A) Transcriptional response of H37Rv to 1554-06 (**5**). Volcano plot of DEseq2 output from analysis of 3 replicate cultures treated with 4X MIC of **5** compared to DMSO vehicle alone. The 3 genes comprising the operon containing *pks13* are highlighted in red, and Rv2581c is highlighted in blue. X-axis is fold change (log2), and y-axis is the adjusted p-value. (B) Genotype of resistant mutants and MIC_90_shifts against **5b** and the advanced benzofuran TAM16.

### Isolation of 1554-Resistant Mutants

We generated resistant mutants to **5** and performed targeted PCR and Sanger sequencing of *pks13* on four mutants derived from two independent parental clones, identifying four distinct single nucleotide polymorphisms (SNPs) in the *pks13* gene (*Rv3800c*), each resulting in a single amino acid substitution in the thioesterase domain of Pks13: V1537A, D1644E, I1575S, and R1578S (Figure 3B). To confirm there were no additional mutations in *Rv2581c* or elsewhere in the genome, we performed whole genome sequencing of three of these mutants, and no other SNPs were identified. All mutations conferred >4X increases in the MIC90 to **5b**, implying that the thioesterase domain of Pks13 was the likely target of this series.

The V1537A missense mutation has previously been implicated in resistance to TAM16, as have other substitutions at D1644 (D1644G/Y)^20,31^. Indeed, we confirmed that the D1644E also conferred high level resistance to TAM16. The I1575 and R1578 mutations do not confer cross-resistance to TAM16.

### Biochemical Confirmation of Pks13 Thioesterase Inhibition

To confirm the predicted target activity for the Pks13 thioesterase domain of *Mtb*, we expressed and purified the domain as previously described^20,31^ (Figure S3). Thioesterase enzymatic activity was monitored using the fluorogenic fatty-acid ester 4-methylumbelliferyl heptanoate (4-MUH) substrate, which is hydrolyzed by the thioesterase domain to release fluorescent 4-methylumbelliferone, enabling continuous plate-reader monitoring of acyl chain cleavage. The four purified 1554-06 isomers demonstrated stereospecific, dose-dependent inhibition of Pks13 thioesterase activity (Figure 4A) that correlated with the H37Rv whole cell activity. **5b** (3R,4S), the most potent whole cell active stereoisomer, was also the most potent in the biochemical assay, with an IC_50_of 0.79 μM. Meanwhile, the trans isomer that showed mild activity in the H37Rv whole cell assay, **5c** (3R,4R), showed a 6-fold reduced activity against the target relative to the most active isomer (Figure 4B). The completely whole cell inactive stereoisomers, **5a** (3S,4R) and **5d** (3S,4S) were inert in the biochemical assay.

**Figure 4.**
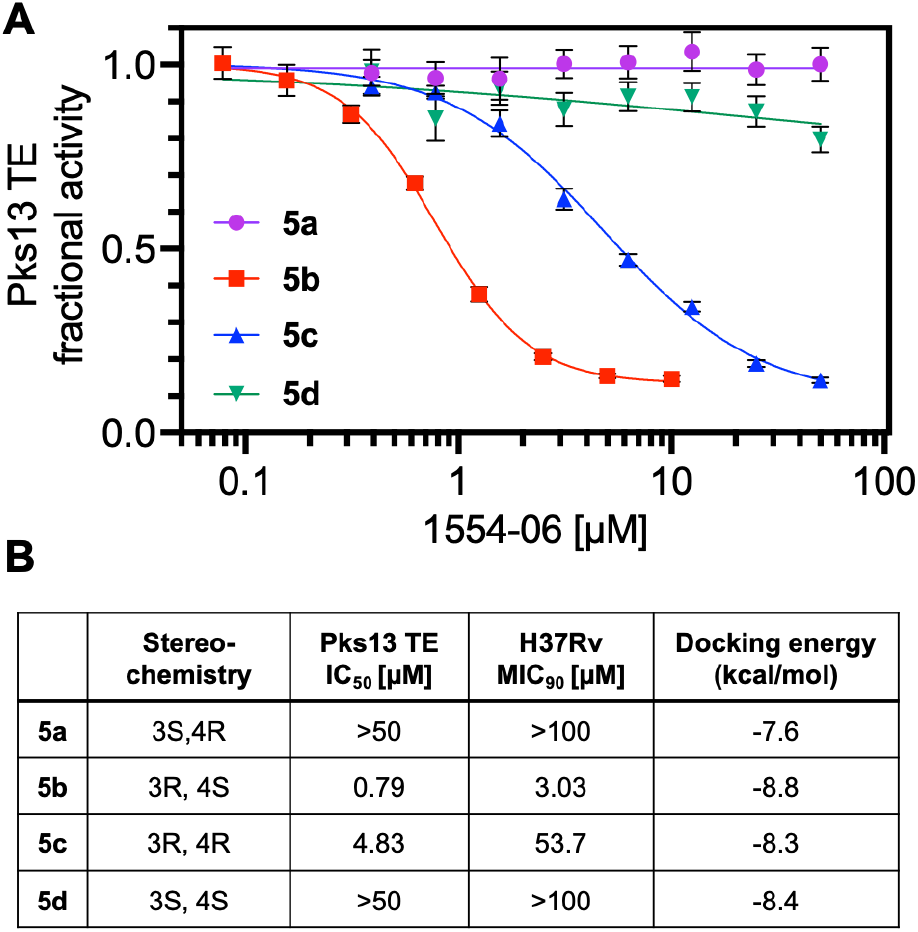
Inhibition of Pks13 thioesterase activity. (A) Biochemical inhibition of recombinant Pks13 thioesterase activity by four purified isomers of 1554-06; 3S,4R (**5a**), 3R,4S (**5b**), 3R,4R (**5c**) and 3S,4S **(5d)**. Initial velocity measurements were conducted in triplicate and normalized to DMSO and TAM16 controls. Error bars = SD. (B) Enzymatic inhibition, whole cell activity, and computational docking values. The highest concentrations used were 50 µM and 100 µM in the enzymatic and whole cell assays, respectively. Docking scores were computed for each isomer of 1554-06 in MOE for the top-ranked ligand poses (see methods).

### Computational Docking of 1554-06 Stereoisomers with the Thioesterase Domain of *Mtb* Pks13

Modest exploration of the structure-activity relationships of the 1554 series via analog synthesis revealed tolerance for alternate phenyl substituents, including the complete removal of the phenyl ring (Table S1). In contrast, replacement of the isoxazole or the pyrazole carboxamide with other heterocycles rendered the analogs inactive, indicating the importance of preserving the scaffold core and side chain (Table S2).

To better understand these medicinal chemistry trends and resistance conferring mutations in the context of Pks13 thioesterase target engagement, we used ligand-receptor co-folding by AlphaFold3 (AF3)^32^, molecular docking, and published X-ray co-crystal structures of the thioesterase domain in the Protein Data Bank (PDB) to evaluate a series of BRD1554 analogs and stereoisomers as ligands in the active-site pocket of Pks13 thioesterase.

The thioesterase domain consists of a conserved α/β-hydrolase core containing the catalytic machinery, capped by a flexible lid region whose conformational dynamics control substrate access and release from the active site^20^. Docking studies suggest that **5b** (3R,4S) largely occupies the same channel previously characterized for Pks13 thioesterase catalytic inhibitor X20404,^22^ which emerged from DNA-encoded library screening (Figure 5A, B).

**Figure 5.**
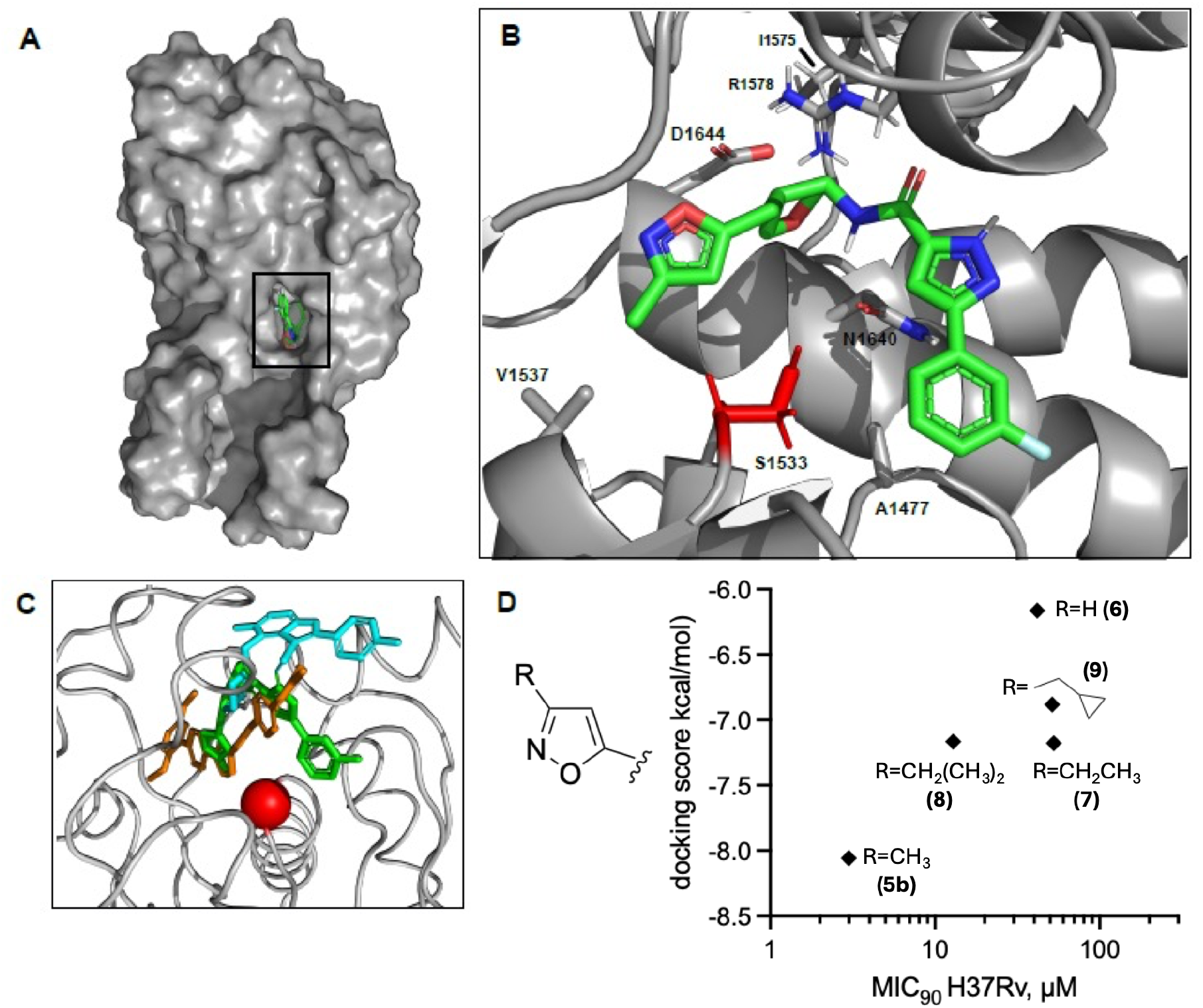
Cofolding and docking analysis of 1554 series. A) The thioesterase domain of *Mtb* Pks13 (UniProt ID: I6X8D2, residues 1451–1727) was cofolded with **5b** (3R,4S). Surface representation of the thioesterase domain (gray) showing the binding grove of the ligand (green). B) Close-up views of inhibitor interactions show **5b** (3R,4S) within the catalytic site; residues implicated in binding and the catalytic serine (red) are shown while other protein segments are hidden for clarity. C) comparison of the docked orientation of **5b** (3R,4S) with the crystal structure poses of TAM16 (cyan) and X20404 (orange) and positions with respect to the catalytic S1533 (red). D) correlation of docking scores (y-axis) of isoxazole substitution analogs with whole cell MIC_90_(x-axis).

Given the different potencies of 1554-06’s four stereoisomers in whole cell MIC and biochemical assays, we calculated the binding energy ranks for the four isomers using molecular docking simulations. The top-ranked docking poses for 1554-06’s 4 stereoisomers varied in their docking energy scores (Figure 4B). Notably, **5b** (3R,4S) was the most favorable ligand, scoring a binding energy of –8.8 kcal.mol^-1^ while its mirror image isomer **5a** (3S,4R) was the least favored, with a calculated docking energy of -7.6 kcal.mol^-1^. The results reflect experimental observations for the two cis isomers (3R,4S; 3S,4R) being the most and least active, respectively, in enzymatic (IC_50_) and whole cell (MIC_90_) assays (Figure 4B). Conversely, the two trans isomers **5c** (3R,4R) and **5d** (3S,4S) scored similarly in docking (-8.3 and -8.4 kcal.mol^-1^) reflecting their reduced activities compared to the active cis isomer.

The co-folding and docking analysis revealed that **5b** adopts a distinctive binding pose within the Pks13 thioesterase active site, positioning the ligand proximally to the catalytic serine S1533 (2.7 Å) while engaging key residues V1537 and D1644 within 4 Å (Figure 5A, B). Notably, **5b** penetrates deeper into the hydrophobic groove than TAM16, enabling the isoxazole moiety to form enhanced hydrophobic interactions with V1537’s isopropyl sidechain while preserving stabilizing contacts with A1477 and N1640. This binding geometry seems unique to **5b** and results in a different pose than that of TAM16 and X20404, Concurrently, it also allows for simultaneous occupancy of both sides of the catalytic serine, suggesting a mechanism for improved active-site engagement and inhibitory potential (Figure 5B, C).

With respect to resistance conferring mutations, the D1644E resistance mutation arises likely due to steric clashes between the larger and longer glutamate sidechain and the tetrahydrofuran moiety of **5b**. Curiously, a V1537A allele conferred the highest MIC_90_shift among all isolated mutations (>23x). The docking pose of **5b** suggests a potential hydrophobic interaction forms between methyl isoxazole and the V1537 sidechain. The V1537A mutation introduces a smaller amino acid sidechain with reduced hydrophobic surface area (Figure S4). Similarly, removal of the isoxazole methyl of **5b** also resulted in loss of activity, suggesting this interaction point is critical for productive inhibition (Figure 5D). Additional analogs substituting the isoxazole methyl of **5b** for larger hydrocarbons were synthesized; however larger groups resulted in both higher docking energy and reduced whole cell activity experimentally (Figure 4D).

Meanwhile, R1578S and I1575S resistance-conferring substitutions are unique to **5b**, resulting in >4x shift in MIC_90_(Figure. 2B), and do not confer resistance to TAM16. These substitutions reside in the α-helix 4 (1572-1588) that constitutes part of the flexible lid domain previously shown to adopt labile states depending on the bound inhibitor^22^. While unlikely to make direct contact with **5b** based on the most favorable docking pose and distance (>10Å), a charge elimination (R1578S) at this position would attenuate existing polar contacts (*e*.*g*., with D1607, D1644) within the lid subdomain and may disfavor inhibitor accessibility and/or stability within the active site (Figure 5B).

### 1554-06-3R,4S is Bactericidal and Active Intracellularly

Bactericidal activity is a valuable property for drugs used to treat tuberculosis, contributing to early bacterial clearance and reduction in transmission of the disease. We tested **5b** for its ability to kill replicating *Mtb*, which is a known property of previously described Pks13 inhibitors (Figure 6A). As with previous inhibitors, treatment with **5b** led to a rapid drop in viability, with a >5 log reduction in colony forming units (cfu) over the first 3 days of treatment, similar to the effect seen with isoniazid (INH). Unlike INH, however, no evidence of culture regrowth due to the rapid emergence of resistance was seen at 11 days.

**Figure 6.**
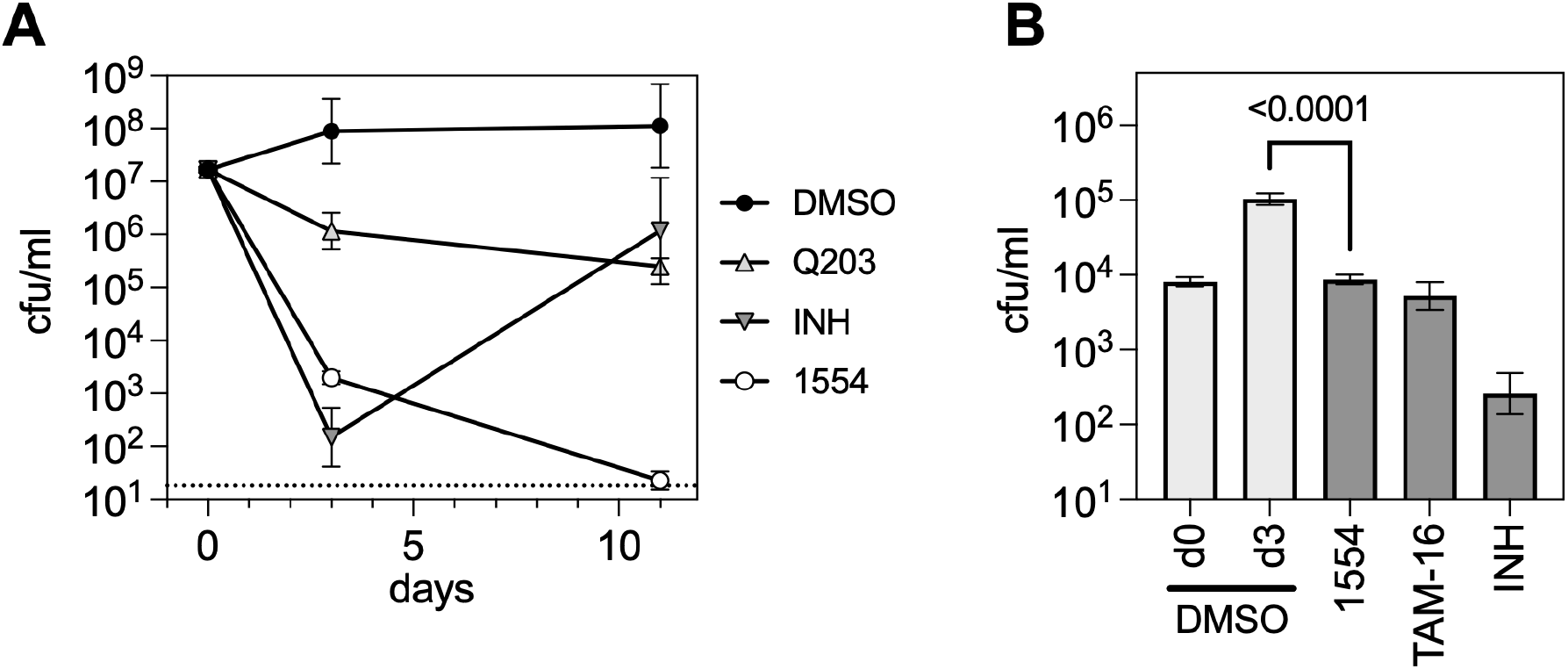
1554-06-3R,4S (5b) is bactericidal and active inside macrophages. (A) Bacterial survival after exposure to **5b** and control antitubercular agents as measured by colony forming units (cfu). Q203 and INH were chosen as bacteriostatic and bactericidal controls, respectively. All compounds were administered at 4X MIC. n=3, error bars: SD. Limit of detection (dashed line) is 18 cfu/ml. (B) Bacterial growth in J774 macrophages, measured as cfu, after 3 days in the presence of **5b** and control compounds. n=3, error bars = SD.

Another valuable property of antitubercular compounds is activity against bacilli residing in macrophages. **5b** was active to a comparable level as the advanced benzofuran thioesterase inhibitor TAM16, fully controlling bacterial replication within J774 macrophages over the course of three days (Figure 6B). In contrast to INH however, no significant reduction in cfu was observed in this time frame with either **5b** or TAM16. Of note, TAM16 has been previously reported^20^ to accumulate well within THP-1 macrophages and show activity in these cells and was efficacious in BALB/c mice.

## CONCLUSIONS

We have previously described PROSPECT, a chemical-genetic interaction profiling strategy for identifying novel molecules with antitubercular activity that also provides immediate insight into mechanism of action. Using PROSPECT, we have not only discovered and characterized inhibitors of novel targets such as the essential efflux pump EfpA^6^, but have also identified new scaffolds for validated targets, including QcrB^8^, the cytochrome b subunit of Complex III in the *Mtb* electron transport chain. Importantly, in these cases, the initial screening hit had weak to no wild-type activity, but with limited chemistry, a relatively potent, wild-type active molecule could be synthesized, thus demonstrating the potential to discover promising scaffolds that would otherwise elude conventional, wild-type screening discovery efforts. Here we demonstrate another example of a potent *Mtb* inhibitor of a high value target discovered through mining of PROSPECT screening data.

Based on PCL analysis, a reference-based method for initial MOA assignment, we identified a N-oxolanyl pyrazole carboxamide series (BRD1554) as a likely Pks13 inhibitor. Further, PROSPECT specifically implicated the thioesterase domain of the multidomain Pks13 enzyme based on unique genetic interactions that distinguished thioesterase and ACP domain inhibitors. Rv2581c, a putative glyoxylase, is hypersensitized to thioesterase inhibitors, while Rv2971, an alpha-keto reductase that also features prominently in FadD32 inhibitor CGI profiles, is hypersensitized to inhibition of the ACP domain of Pks13. FadD32 and Pks13 ACP inhibitors mechanistically mirror each other, with FadD32 required to transfer meromycolic acids to the ACP of Pks13. These results highlight the potential of PROSPECT to predict MOA with high resolution, leveraging the biology embedded in hypomorph sensitization and gene-gene interactions within the cell.

Pks13 is a highly vulnerable target in an essential pathway – mycolic acid biosynthesis – that is limited to a small number of bacteria, including pathogenic mycobacteria and corynebacteria. Thus, despite the relatively weak activity of the screening hit (MIC of 24.7 µM), knowledge that the likely target was potentially of high value for therapeutic development encouraged us to pursue this molecule further. Subsequent genetic and biochemical studies confirmed Pks13 as the protein target, underscoring the value of PROSPECT for rapidly triaging whole cell active compounds by incorporating insight into biological mechanism.

We elucidated the absolute stereochemistry of the Pks13 thioesterase inhibitor, defining the most active stereoisomer as 3-(3-fluorophenyl)-*N*-((3*R*,4*S*)-4-((3-methylisoxazol-5-yl)methyl) tetrahydrofuran-3-yl)-1*H*-pyrazole-5-carboxamide (1554-06-3R,4S, **5b**) and demonstrated its inhibition of the thioesterase domain of Pks13. Computational docking illuminated the basis for the stereospecificity of **5b** (3R,4S), with the docked pose of **5b** bridging contacts between the catalytic site and the previously reported inhibitors TAM16 and X20404, allowing occupancy on both sides of the catalytic residue. This stereospecific binding modality of **5b** (3R,4S) could guide future chemical optimization of this molecule to achieve greater potency.

In addition to identifying novel small molecule candidates as chemical probes or for early drug discovery efforts, PROSPECT has the potential to uniquely elucidate *Mtb* biology by revealing novel gene-gene interactions. By characterizing the nearly 500 hypomorphs targeting essential *Mtb* genes in response to over 400 reference compounds and many thousands of unknown compounds, each serving as a unique perturbation, PROSPECT can illuminate not only hundreds of thousands of chemical-genetic interactions, but also previously unrecognized gene-gene interactions based on correlation of strain behaviors in PROSPECT. In other words, non-obvious hypomorphs that are sensitized to inhibitors of specific targets can reveal previously unrecognized interactions between the hypomorphic gene and the small molecule target; similarly, strains that share sensitivities to common chemical entities could also be functionally related. These shared sensitivities can thus provide insight into the biological roles of genes of unknown function.

As an example, for a number of Pks13 thioesterase inhibitors, in addition to showing strikingly enhanced activity in *Mtb* strains depleted of the target, Pks13, we also found enhanced activity in a strain depleted of an uncharacterized enzyme, Rv2581c, suggesting a previously undescribed link between Pks13 and this enzyme. Rv2581c encodes a putative glyoxylase of the GLX II family, which suggests a possible role in the detoxification of methylglyoxal, a toxic byproduct produced when triose phosphates accumulate^33^. Inhibition of Pks13 thioesterase activity would lead to enzymatic stalling, with the newly condensed mycolic acid tethered to the second ACP domain of Pks13 via a phoshopantethyl thioether^15^. Based on the high degree of similarity of the Rv2581c protein to characterized glyoxylase II enzymes, which detoxify lactoyl-thiols, it seems unlikely that this protein directly releases the stalled mycolic acid but may instead address some downstream consequence of this process. It is currently unresolved whether trapped mycolic acids could be released from TE-inhibited Pks13, and if they are, the fate of these released mycolic acids is unclear, as there is no defined pathway in *Mtb* for the catabolism of mycolic acids. Nonetheless, our transcriptomic data shows strong upregulation of *icl* in response to Pks13 thioesterase inhibition, suggesting increased beta-oxidation, possibly of accumulating fatty acid precursors. This in turn could lead to increased gluconeogenic flux, which would produce triose phosphates, potentially leading to methylglyoxal accumulation and thus a role for the putative glyoxylase Rv2581c. In contrast, Rv2581c appears to be dispensable when Pks13 is inhibited prior to this condensation, given the lack of hypersensitization to Pks13 ACP or FadD32 inhibitors, which prevent meromycolate from being loaded onto the Pks13 ACP by the FadD32 enzyme. Instead, the hypomorph of Rv2971, a putative NADPH-dependent aldo-keto reductase, also suggested to play a role in methylglyoxal detoxification^34^, is sensitized to blockade of ACP loading. Presumably, the difference in accumulated mycolic acid precursors (Pks13-bound mycolic acid versus uncondensed C_26_fatty acids and meromycolates) drives the differential requirements for these detoxifying enzymes. A detailed functional characterization of these enzymes in Mycobacteria will illuminate the precise mechanisms linking them to Pks13 inhibition.

These gene-gene interactions highlight opportunities to identify beneficial synergies to help advance the development of novel *Mtb* therapeutic regimens. Both Rv2581c and Rv2971 are essential in *Mtb*^26,35^, even in the absence of Pks13 inhibition, suggesting that they likely play critical roles in detoxifying products that arise during unperturbed *Mtb* growth. Thus, on their own, they are potential drug targets. But by illuminating their increased importance in the context of domain-specific Pks13 inhibition, PROSPECT reveals new strategies for identifying small molecules inhibitors that can synergize with Pks13 inhibitors in development.

## MATERIALS AND METHODS

### *M. tuberculosis* Strains and Media

*Mycobacterium tuberculosis* H37Rv and hypomorphic derivatives were cultured in Middlebrook 7H9 medium supplemented with OADC, 0.05% Tween 80, and 10 mM sodium acetate. Optical density at 600 nm (OD_600_) was measured using a 1-cm pathlength cuvette. Growth on solid medium used Middlebrook 7H10 supplemented with OADC and 0.2% glycerol. Hypomorphs were grown in medium supplemented with hygromycin (50 µg/ml) and streptomycin (20 µg/ml), and anhydrotetracycline was added to a concentration of 200 ng/ml to repress the hypomorphic phenotype. For MIC determinations, an exponentially growing culture was diluted to an OD_600_of 0.0025, and 100 µl of this was transferred into 96 well plates (Corning) and incubated from 7 days (H37Rv), 10 days (Rv3581c), or 21 days (Pks13), with duration of incubation adjusted to account for the slow growth of hypomorphs, ensuring a comparable number of cell divisions throughout the assay. For MICs of hypomorphs, the mid-log cultures were first washed twice in culture medium to induce proteolysis prior to transfer into assay plates. For MIC assays, small molecules were first diluted in DMSO and then added to assay plates in 1 µl volumes, yielding a final DMSO concentration of 1.0 %. To calculate the MIC, data were first normalized using DMSO and rifampin (1 µM) controls, and a 4-parameter nonlinear regression model was fit to the resulting data, with 90% inhibition used as the response threshold.

### PROSPECT screening and analysis

Chemical-genetic interaction profiling was performed using PROSPECT as previously described^6^. To enable reference-based mechanism of action (MOA) inference, CGI profiles were compared to a curated reference collection (Figshare data deposition https://doi.org/10.6084/m9.figshare.28373561) spanning 438 compounds and 71 MOAs and analyzed using Perturbagen CLass (PCL) analysis as described previously^8^ and using code in MATLAB and R previously made available on Code Ocean (https://doi.org/10.24433/CO.3013890.v1) and GitHub (https://github.com/broadinstitute/Mtb_PROSPECT_PCL_analysis).

### Chemical Methods, Synthesis and Characterization of 1554-06 analogues

All chemical reagents were purchased from commercial sources and used without further purification unless noted in the Supporting Information. Flash chromatography was performed using silica gel (250-400 mesh). NMR spectra were recorded on Bruker 400 spectrometer at 400 MHz for 1H NMR. Additional experimental details may be found in the Supporting Information.

### Diethyl ((3-methylisoxazol-5-yl)methyl)phosphonate (2)

A solution of 5-(chloromethyl)-3-methylisoxazole (282 mg, 2.14 mmol) and P(OEt)_3_(1.84 mL, 10.72 mmol) was stirred at 119°C overnight under N_2_atmosphere. The reaction mixture was concentrated in vacuo. The residue was purified by silica gel chromatography (eluted with DCM/MeOH = 20/1) to give diethyl ((3-methylisoxazol-5-yl)methyl)phosphonate (**2**, 406 mg, 81% yield) as a colorless oil. LC-MS (ESI, *m/z*): [M+H]^+^ = 234.1.

### tert-Butyl (4-((3-methylisoxazol-5-yl)methylene)tetrahydrofuran-3-yl)carbamate (3)

To a solution of diethyl ((3-methylisoxazol-5-yl)methyl)phosphonate (159 mg, 0.68 mmol) in THF (2 mL) was added NaH (60%, 27 mg, 0.68 mmol) at 0 °C. The resulting mixture was stirred at room temperature for 0.5 h. Then, tert-butyl (4-oxotetrahydrofuran-3-yl)carbamate (114 mg, 0.57 mmol) was added. The resulting mixture was stirred at room temperature overnight. The reaction mixture was diluted with water (30 mL) and extracted with ethyl acetate (30 mLx3). The combined organic layers were washed with brine, dried over anhydrous Na_2_SO_4_and concentrated in vacuo. The residue was purified by silica gel column chromatography (eluted with PE/ethyl acetate = 4/1) to give tert-butyl (4-((3-methylisoxazol-5-yl)methylene)tetrahydrofuran-3-yl)carbamate (**3**, 50 mg, 31% yield) as a colorless oil. LC-MS (ESI, *m/z*): [M+H-C4H8]^+^ = 225.1.

### tert-Butyl (4-((3-methylisoxazol-5-yl)methyl)tetrahydrofuran-3-yl)carbamate (4)

To a mixture of tert-butyl (4-((3-methylisoxazol-5-yl)methylene)tetrahydrofuran-3-yl)carbamate (**3**, 817 mg, 2.91 mmol) in EtOH (40 mL) was added Pd/C (10% on carbon, 408 mg), and the mixture placed under H_2_atmosphere. The mixture was stirred at room temperature for 4.0 h. The reaction mixture was filtered on Celite and rinsed with MeOH. The solvent was concentrated in vacuo. The residue was purified by prep-HPLC (eluted with CH_3_CN/H_2_O = 5/95∼ 95/5 with 0.1% TFA) to give tert-butyl (4-((3-methylisoxazol-5-yl)methyl)tetrahydrofuran-3-yl)carbamate (**4**, 192 mg, 23% yield, mixture of 4 diastereomers) as a colorless oil. LC-MS (ESI, *m/z*): [M+H-C4H8]^+^ = 227.1.

### 3-(3-Fluorophenyl)-N-(4-((3-methylisoxazol-5-yl)methyl)tetrahydrofuran-3-yl)-1H-pyrazole-5-carboxamide (1554-06, 5)

A mixture of 4-((3-methylisoxazol-5-yl)methyl)tetrahydrofuran-3-amine (**4**, 50 mg, 0.12 mmol), 3-(3-fluorophenyl)-1H-pyrazole-5-carboxylic acid (36 mg, 0.15 mmol), HATU (56 mg, 0.15 mmol), DIPEA (79 mg, 0.61 mmol) and dry DMF (2.0 mL) was stirred room temperature for 1.0 h. The reaction mixture was purified by reverse phase silica gel column chromatography (eluted with CH_3_CN/H_2_O = 5/95∼95/5 with 0.1% TFA) to give 3-(3-fluorophenyl)-N-(4-((3-methylisoxazol-5-yl)methyl)tetrahydrofuran-3-yl)-1H-pyrazole-5-carboxamide (**1554-06, 5**, 44 mg, TFA salt, 60% yield, cis/trans=5/3) as a white solid. LC-MS: MS(ESI, m/z): [M+H]^+^=371.1 ^1^H NMR (400 MHz, Methanol-d4) δ 7.56 (dd, J = 7.7, 4.0 Hz, 2H), 7.52 – 7.51 (m, 1H), 7.49 (dd, J = 5.4, 2.8 Hz, 2H), 7.47 – 7.43 (m, 1H), 7.16 – 7.13 (m, 1H), 7.13 – 7.07 (m, 2H), 6.13 (s, 1H), 6.04 (s, 1H), 4.84 – 4.81 (m, 1H), 4.41 (q, J = 5.9 Hz, 1H), 4.14 – 4.07 (m, 2H), 4.02 (dd, J = 8.7, 7.4 Hz, 1H), 3.82 (dd, J = 9.3, 3.8 Hz, 1H), 3.72 – 3.66 (m, 2H), 3.62 – 3.58 (m, 1H), 3.06 (dd, J = 15.3, 6.8 Hz, 1H), 3.00 (dd, J = 14.9, 5.9 Hz, 1H), 2.90 (dt, J = 13.1, 7.5 Hz, 2H), 2.79 (dd, J = 14.9, 8.8 Hz, 1H), 2.73 – 2.66 (m, 1H), 2.19 (s, 3H), 2.18 (s, 2H).

### Chiral separation of 3-(3-Fluorophenyl)-N-(4-((3-methylisoxazol-5-yl)methyl)tetrahydrofuran-3-yl)-1H-pyrazole-5-carboxamide (1554-06, 5)

3-(3-fluorophenyl)-N-(4-((3-methylisoxazol-5-yl)methyl)tetrahydrofuran-3-yl)-1H-pyrazole-5-carboxamide (**1554-06, 5**, 130 mg, 0.35 mmol) was separated by chiral preparation on an Waters SFC-150mgm with a Daicel AD (25*250mm,10um) column at 30 °C using a CO2/EtOH[0.5%NH3(7M in MeOH)]=45/55 mobile phase, a flow rate of 100 ml/min with a back pressure of 100 bar, detection at 214 nm and a cycle time of 13 minutes. The sample solution was dissolved at 130mg/6.2 ml MeOH, injection volume was 2.5 ml.

### 1554-06-cis1 (5a)

3-(3-fluorophenyl)-N-((3S,4R)-4-((3-methylisoxazol-5-yl)methyl)tetrahydrofuran-3-yl)-1H-pyrazole-5-carboxamide (39.7 mg). LC-MS (ESI, *m/z*): [M + H]^+^ = 371.1. ^1^H NMR (400 MHz, Methanol-d4) δ 7.55 (s, 1H), 7.48 (s, 2H), 7.11 (s, 2H), 6.04 (s, 1H), 4.84 – 4.80 (m, 1H), 4.09 (dd, J = 9.4, 6.1 Hz, 1H), 4.02 (dd, J = 8.7, 7.4 Hz, 1H), 3.82 (dd, J = 9.4, 3.8 Hz, 1H), 3.69 (t, J = 8.4 Hz, 1H), 3.00 (dd, J = 14.9, 5.9 Hz, 1H), 2.95 – 2.84 (m, 1H), 2.79 (dd, J = 14.8, 8.8 Hz, 1H), 2.19 (s, 3H).

### 1554-06-cis2 (5b)

3-(3-fluorophenyl)-N-((3R,4S)-4-((3-methylisoxazol-5-yl)methyl)tetrahydrofuran-3-yl)-1H-pyrazole-5-carboxamide (33.2 mg). LC-MS (ESI, *m/z*): [M + H]+ = 371.1. ^1^H NMR (400 MHz, Methanol-d4) δ 7.56 (d, J = 7.8 Hz, 1H), 7.53 – 7.42 (m, 2H), 7.16 – 7.07 (m, 2H), 6.04 (s, 1H), 4.85 – 4.80 (m, 1H), 4.09 (dd, J = 9.4, 6.1 Hz, 1H), 4.02 (dd, J = 8.7, 7.4 Hz, 1H), 3.82 (dd, J = 9.4, 3.8 Hz, 1H), 3.69 (t, J = 8.4 Hz, 1H), 3.00 (dd, J = 14.9, 5.9 Hz, 1H), 2.95 – 2.84 (m, 1H), 2.79 (dd, J = 14.9, 8.8 Hz, 1H), 2.19 (s, 3H).

### 1554-06-trans1 (5c)

3-(3-fluorophenyl)-N-((3R,4R)-4-((3-methylisoxazol-5-yl)methyl)tetrahydrofuran-3-yl)-1H-pyrazole-5-carboxamide (7.8 mg). LC-MS (ESI, *m/z*): [M + H]+ = 371.1.^1^H NMR (400 MHz, Methanol-d4) δ 7.55 (dt, J = 7.8, 1.2 Hz, 1H), 7.52 – 7.41 (m, 2H), 7.15 – 7.06 (m, 2H), 6.13 (s, 1H), 4.41 (dt, J = 6.8, 5.5 Hz, 1H), 4.11 (dt, J = 9.1, 7.2 Hz, 2H), 3.70 (dd, J = 9.1, 5.5 Hz, 1H), 3.60 (dd, J = 9.0, 6.6 Hz, 1H), 3.06 (dd, J = 15.4, 6.7 Hz, 1H), 2.91 (dd, J = 15.4, 8.2 Hz, 1H), 2.70 (dqd, J = 8.2, 6.8, 5.5 Hz, 1H), 2.18 (s, 3H).

### 1554-06-trans2 (5d)

3-(3-fluorophenyl)-N-((3S,4S)-4-((3-methylisoxazol-5-yl)methyl)tetrahydrofuran-3-yl)-1H-pyrazole-5-carboxamide (9.3 mg). LC-MS (ESI, *m/z*): [M + H]+ = 371.1. ^1^H NMR (400 MHz, Methanol-d4) δ 7.55 (dt, J = 7.7, 1.3 Hz, 1H), 7.53 – 7.41 (m, 2H), 7.15 – 7.06 (m, 2H), 6.13 (s, 1H), 4.41 (dt, J = 6.9, 5.6 Hz, 1H), 4.11 (dt, J = 9.2, 7.2 Hz, 2H), 3.70 (dd, J = 9.1, 5.5 Hz, 1H), 3.60 (dd, J = 9.0, 6.6 Hz, 1H), 3.06 (dd, J = 15.4, 6.8 Hz, 1H), 2.91 (dd, J = 15.4, 8.2 Hz, 1H), 2.76 – 2.63 (m, 1H), 2.18 (s, 3H).

### Resistant Mutant Selection

Two independent single-colony isolates of H37Rv (SC11 and SC18) were grown in parallel and used for isolation of mutants resistant to 1554-06 **(5)**, which ensured that any mutations common to these two parental cultures arose independently. Selection was done at 2 cell concentrations (2×10^8^ and 1×10^7^) and 2 compound concentrations (4X and 8X the measured MIC_90_). Colonies were picked into media containing 1X MIC_90_and expanded. Genomic DNA was isolated using a Cetrimide-based protocol^36^. Libraries were constructed using the NexTerra kit (Illumina) and sequenced on a NextSeq X instrument. Reads were aligned to NC_000962.3 and polymorphisms were identified using Pilon^37^.

### Transcriptomic Analysis

H37Rv was grown at 37°C to mid log phase (OD_600_of 0.57) and back-diluted to an OD_600_of 0.4 prior to exposure to compound at 4X MIC_90_in 1.2 ml volume in a 24 well plate in triplicate. Cultures were collected by centrifugation at 8000 xg for 2 minutes and resuspended in 500 µL Trizol reagent (ThermoFisher Scientific). For RNA extraction, cell pellets in Trizol reagent were transferred to 2mL FastPrep tubes (MP Biomedicals) containing 0.1 mm Zirconia/Silica beads (BioSpec Products) and bead beaten for 90 seconds at 10 m/sec speed using the FastPrep-24 5G (MP Biomedicals). After addition of 200 µl chloroform, each sample tube was mixed thoroughly by inversion, incubated for 3 minutes at room temperature, and spun down 15 minutes at 4°C. The aqueous phase was mixed with an equal volume of 40% ethanol, transferred to a Direct-zol spin plate (Zymo Research), and RNA was extracted according to the Direct-zol protocol (Zymo Research).

Illumina cDNA libraries were generated using a modified version of the RNAtag-seq protocol^38,39^. Briefly, up to 100-250 ng of total RNA was fragmented, depleted of genomic DNA, dephosphorylated, and ligated to DNA adapters carrying 5’-AN_8_-3’ barcodes of known sequence with a 5’ phosphate and a 3’ blocking group. Barcoded RNAs were pooled and depleted of rRNA using the riboPOOLS rRNA depletion kit (siTOOLS). Pools of barcoded RNAs were converted to Illumina cDNA libraries in 2 main steps: (i) reverse transcription of the RNA using a primer designed to the constant region of the barcoded adaptor with addition of an adapter to the 3’ end of the cDNA by template switching using SMARTScribe (Clontech) as described^40^; (ii) PCR amplification using primers whose 5’ ends target the constant regions of the 3’ adaptors and whose 3’ ends contain the full Illumina P5 or P7 sequences. cDNA libraries were sequenced on an Illumina NovaSeq X platform, generating 150 bp paired-end reads.

Sequencing reads from each sample in a pool were demultiplexed based on their associated barcode sequence using custom scripts (https://github.com/broadinstitute/split_merge_pl). Up to 1 mismatch in the barcode was allowed provided it did not make assignment of the read to a different barcode possible. Barcode sequences were removed from the first read as were terminal Gs from the second read that may have been added by SMARTScribe during template switching. Reads were aligned to NC_000962.3 using BWA^41^ and read counts were assigned to genes and other genomic features using custom scripts (https://github.com/broadinstitute/BactRNASeqCount). Differential expression analysis was conducted with DESeq2^42^.

### Cloning, expression, and purification of the thioesterase domain of *Mycobacterium tuberculosis* Pks13

The thioesterase domain encoding region of *Mtb pks13* was amplified by PCR using the full length *pks13* gene (Uniprot: I6X8D2) as template DNA, with a N-terminal His(6x) tag and thrombin and TEV cleavage sites incorporated (HHHHHHGSLVPRGSASENLYFQGGSGS). The amplified DNA was cloned into the pET-21(+) bacterial expression vector by Gibson assembly, the plasmid was transformed into *E. coli* BL21(DE3) strain (NEB), and a primary culture was started from a single colony obtained after transformation. The primary culture was diluted into a secondary culture of LB medium containing carbenicillin (100 µg/mL) and was grown at 37°C to an OD_600_of ∼0.6. Expression of the thioesterase construct was induced with 0.5 mM IPTG and cells were harvested after 16 hr of growth at 20°C.

The harvested cells were resuspended in the lysis buffer (50 mM Tris-HCl pH 8.0, 0.5 M NaCl, 10% (v/v) glycerol, 1 mM TCEP, protease inhibitor, and DNase) and lysed by passing through a chilled microfluidizer 3 times (Avestin Emulsiflex-C3, 10-15k PSI). The resulting cell extract was clarified by centrifugation (15,000 x *g*) for 1 hr at 4°C and filtered (PES, 0.22 µm). The cleared supernatant was loaded onto a Ni NTA-affinity column, and the His-tagged thioesterase domain constructs were eluted with a linear gradient of 10-250 mM imidazole in 20 mM Tris-HCl, pH 8.0 and 0.5 M NaCl. The peak fractions were pooled and were concentrated for loading onto a Superdex-200 gel filtration column (GE Healthcare). The thioesterase domain eluted under a single peak as a monomer and was > 95% pure as observed by SDS-PAGE and intact mass spectrometry analysis. The purified protein was concentrated to 20-25 mg/ml, flash-frozen and stored at ™80°C.

### Measurement of Thioesterase Enzymatic Activity

Activity of the Pks13 thioesterase domain was evaluated using 4-methylumbelliferyl heptanoate (4-MUH; Sigma) as a fluorogenic substrate in a 96-well plate format^20^. For initial velocity measurements, Pks13-TE (1 μM) in 0.1 M Tris-HCl, pH7 buffer was incubated with compounds from the 1554 series in a 100 µL reaction volume. Compounds were prepared at 50x desired final concentration, serially diluted 2-fold for a total of eight concentrations, and 2 μL of each dilution was added to protein. Protein samples treated with 2% (v/v) DMSO were used as controls to determine the baseline activity of the Pks13-TE domain. The plates with protein and compound were incubated at 37°C for 30 min, then the reaction was initiated with 20 µM 4-MUH. The fluorescence of the hydrolyzed product 4-methylumbelliferone was read every 5 minutes for 90 minutes at 37°C (excitation at 355 nm and emission at 460 nm) using a SpectraMax Paradigm Plate Reader (Molecular Devices). Background hydrolysis of the 4-MUH substrate in buffer as a control and subtracted from all data points. Enzymatic activity upon treatment with the 1554 series was normalized between 100% (enzyme only) and 0% (full inhibition by 5 μM of the control compound TAM16). Data points were plotted as an average of three independent experiments. The IC_50_values were calculated using a 4-parameter nonlinear regression model in GraphPad Prism.

### Pks13:1554-06 Ligand-Complex Docking

The thioesterase (TE) domain of Pks13 (UniProt ID: I6X8D2, residues 1451–1727) was co-folded with 1554-006-cis-3R,4S to identify the ligand-binding pocket and to prepare the protein conformation for subsequent docking. AlphaFold 3 (PiperOrigin-RevId: 695389383) was employed to generate five cofolding predictions using default parameters and a random seed of 314. The top-ranked model, based on the ranking_score, was selected for further analysis.

The cofolded ligand was removed from this model, and the remaining protein structure was used for molecular docking studies. Docking simulations were conducted using MOE (version 2022.02) with four stereoisomers of 1554-006-3R,4S, -3S,4R, -3S,4S, and -3R,4R—as well as five analogs of 1554-006-cis-3R,4S.

Protein preparation was performed using MOE’s QuickPrep module, which assigned appropriate protonation states, optimized sidechain rotamers, and minimized steric clashes. Ligands were then docked to the prepared receptor using the Triangle Matcher placement method with a rigid receptor setting. The top-ranked docking pose for each ligand was recorded along with its corresponding binding energy.

#### Determination of bactericidal activity

*Mtb* was grown to mid log phase and diluted to an OD_600_of 0.01 prior to exposure to 4x the measured MIC of each of the compounds in 100 µl volumes in 96 well plates. Plates were incubated for 3 and 11 days, at which time 3 individual wells per treatment were sampled, serially diluted, and plated onto 7H10 agar plates for colony enumeration.

#### M. tuberculosis infection assay in J774 macrophages

J774 murine macrophages were thawed, expanded, and maintained in DMEM supplemented with 10% heat-inactivated FBS at 37°C. Cells were seeded in flat, clear-bottom 96-well plates at 3,000 cells/well (50 µL; 60,000 cells/mL) and incubated overnight. *Mtb* H37Rv(pUV3583cGFP) was cultured in 7H9 medium supplemented with OADC, sodium acetate, Tween-80, and hygromycin. On the day of infection, bacteria were washed, de-clumped by low-speed centrifugation, diluted in DMEM/FBS to 6×10^4^ CFU/mL, and added to macrophages at MOI 1 (50 µL/well). Plates were centrifuged briefly to synchronize infection and incubated for 4 h. Following infection, wells were washed with PBS, and either lysed immediately (T0) with 0.5% Triton X-100 for CFU enumeration or replenished with DMEM/FBS containing vehicle (0.5% DMSO) or test compounds. For CFU determination, lysates were serially diluted in PBST and plated on 7H10 agar. Infected cultures were incubated for 72 h, after which additional lysis and serial dilutions were performed for CFU plating. Plates were incubated at 37 °C until colony formation for enumeration.

#### HepG2 cytotoxicity assay

HepG2 cells were maintained in DMEM supplemented with 10% fetal bovine serum at 37 °C and 5% CO_2_. Cells were thawed from liquid nitrogen, expanded in T75 flasks, and passaged by trypsinization at near confluency. For assays, cells were harvested, counted by trypan blue exclusion, and diluted to 1×10^5^ cells/mL, then seeded into white, clear-bottom 96-well plates at 1×10^4^ cells/well (100 µL) and incubated overnight. Compounds were prepared as serial dilutions in culture medium and added to cells (1.2 µL per well) to achieve a top final concentration of 120 µM, with DMSO-treated wells serving as controls. Plates were incubated for 48 hours, after which cell viability was assessed using CellTiter-Glo diluted 1:1 in PBS. Luminescence was measured following a 2-minute shake and 10-minute incubation at room temperature using a SpectraMax M5 plate reader. Cytotoxicity was determined by comparing luminescence from compound-treated wells to DMSO controls.

## ASSOCIATED CONTENT

Supporting information is available in .pdf format and contains the following:

1. Figures describing the PCL MOA predictions, crystallographic methods and data, QC of recombinant Pks13, and details of molecular docking
2. Tables of MIC and HepG2 activity of 1554 series analogs
3. Experimental Procedures including Schemes, Synthesis, and Characterization of 1554 series analogs
4. ^1^H NMR spectra and chiral HPLC spectra for 1554 series

## NOTES

The Authors declare no competing financial interests

## ACKNOWLEDGEMENTS

RNA-Seq libraries were constructed and sequenced at the Broad Institute of MIT and Harvard by the Microbial ‘Omics Core and Genomics Platform, respectively. The Microbial ‘Omics Core also provided guidance on experimental design and conducted preliminary analysis for all RNA-Seq data.

We thank 3 Point Bio and Viva Biotech for compound synthesis and analytical characterization.

This work was funded by the Bill and Melinda Gates Foundation.

## ABREVIATIONS

PROSPECT: Primary screening Of Strains to Prioritize Expanded Chemistry and Targets
*Mtb*: *Mycobacterium tuberculosis*
CGI: chemical-genetic interaction
PCL: Perturbagen Class
MIC_90_: Minimum Inhibitory Concentration inhibiting 90% of detectable growth
cfu: colony forming unit
DMSO: dimethyl sulfoxide
ACP: acyl carrier protein
KS: ketosynthase
AT: acyltransferase
TE: thioesterase
Boc: tert-butylcarbonate
TFA: trifluoroacetic acid
EtOH: ethanol
DMF: dimethylformamide
HATU: Hexafluorophosphate Azabenzotriazole Tetramethyl Uronium (O-(7-azabenzotriazol-1-yl)-N,N,N’,N’-tetramethyluronium hexafluorophosphate)
RT: room temperature
DIPEA: N,N-diisopropylethylamine

