## Supporting Information for "Large-Scale Chemical-Genetic Interaction Profiling Identifies a Novel Small-Molecule Inhibitor of *Mycobacterium tuberculosis* Polyketide Synthase 13"

**Table of Contents**

Supplemental Figures S1-S4……………….………………..……………………………………….....2

Crystallographic methods and data…………………………….……..……………...............................6

MIC and HepG2 activity of 1554 series analogs (Tables S1-S3)…………………………………............8

Experimental Procedures

Methods and Characterization………..…………………………………………………….……...12

Abbreviations…………………………………………………………………….....................12

Schemes, Synthesis, and Characterization of 1554 series analogs………..…………………….……13

[1]H NMR spectra and chiral HPLC spectra for 1554 series……………………………........................68

#### Supplemental Figures

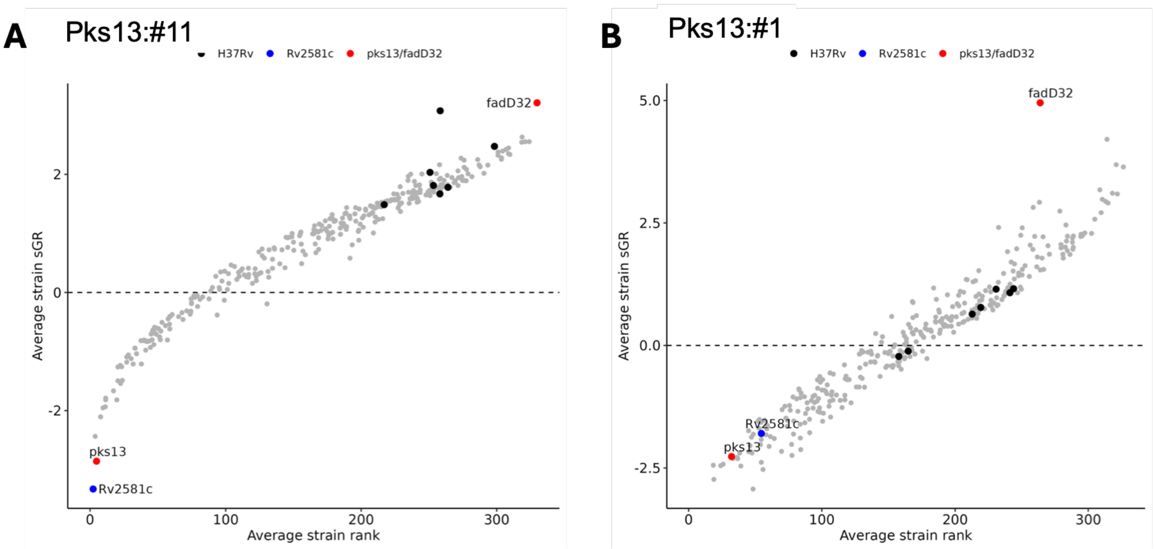


#### Figure S1.

Plots showing the relative sensitization of strains across treatments comprising the two Pks13 PCLs used to assign Pks13 as the likely target of BRD1554 (**1**). Pks13:#11 is comprised of 4 dose-specific CGI profiles each from the reference compounds TAM1 and TAM3. The Pks13:#1 PCL is comprised of 7 dose-specific CGI profiles from 4 compounds (TAM1, TAM3, TAM9, TP4). X axis is average rank of each strain across all dose-specific CGI profiles, Y axis is the average standardized growth rate, a z-scored measure of effect; negative values indicate sensitization.

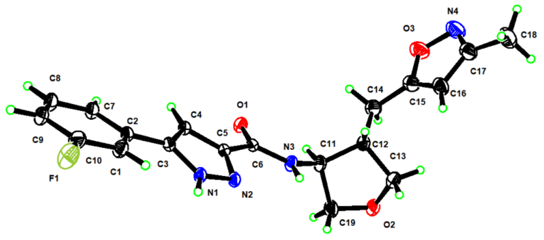


**Figure S2A.** Molecular three-dimensional structure thermoellipsoid diagram of 1554-06-trans2 (**2e**)

<u>Crystal data and structure refinement</u>

| | |
|---|---|
| Empirical formula | $C_{19}H_{19}FN_4O_3$ |
| Formula weight | 370.38 |
| Temperature | 170(2) K |
| Wavelength | 1.54178 Å |
| Crystal system | Orthorhombic |
| Space group | $P2_12_12_1$ |
| Unit cell dimensions | a = 8.7740 (9) Å |
| | b = 11.6981 (12) Å |
| | c = 17.1994 (17) Å |
| Volume | 1765.3 (3) $Å^3$ |
| Z | 4 |
| Density (calculated) | 1.394 $Mg/m^3$ |
| Absorption coefficient | 0.87 $mm^{-1}$ |
| F(000) | 776 |
| Crystal size | 0.15 x 0.08 x 0.05 mm |
| Theta range | 4.6 to 74.7° |
| Index ranges | -10<=h<=10, -13<=k<=14, -21<=l<=21 |
| Reflections collected | 25705 |
| Independent reflections | 3575 [R(int) = 0.072] |
| Absorption correction | multiscan |
| Ratio Max. to min. trans. | 0.7985 |
| Refinement | Full-matrix least-squares on $F^2$ |
| Data/restaints/parameters | 3575/ 0 / 259 |
| Goodness-of-fit on F2 | 1.09 |
| Final R indices | $R[F^2>2\sigma(F^2)] = 0.036$, $wR(F^2) = 0.094$ |
| Absolute structure parameter | 0.07 (8) |
| Largest diff. peak and hole | 0.22 and -0.18 e $Å^{-3}$ |

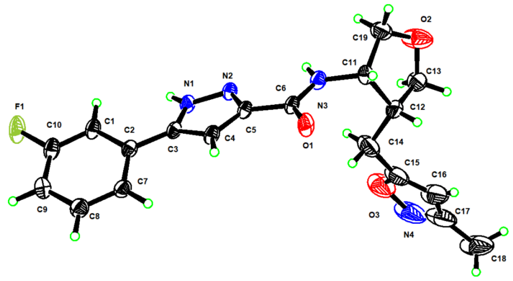


**Figure S2B.** Molecular three-dimensional structure thermoellipsoid diagram of 1554-06-cis1 (**2b**)

<u>Crystal data and structure refinement</u>

| | |
|---|---|
| Empirical formula | $C_{19}H_{19}FN_4O_3$ |
| Formula weight | 370.38 |
| Temperature | 200(2) K |
| Wavelength | 1.54178 Å |
| Crystal system | Tetragonal |
| Space group | $P4_12_12$ |
| Unit cell dimensions | a = 7.1055 (2) Å |
| | b = 7.1055 (2) Å |
| | c = 71.442 (3) Å |
| Volume | 3607.0 (3) $Å^3$ |
| Z | 8 |
| Density (calculated) | 1.364 $Mg/m^3$ |
| Absorption coefficient | 0.85 $mm^{-1}$ |
| F(000) | 1552 |
| Crystal size | 0.18 × 0.09 × 0.06 |
| Theta range | 5.0 to 74.4° |
| Index ranges | -7<=h<=8, -8<=k<=7, -88<=l<=89 |
| Reflections collected | 30261 |
| Independent reflections | 3684 [R(int) = 0.074] |
| Absorption correction | multiscan |
| Refinement | Full-matrix least-squares on $F^2$ |
| Data/restaints/parameters | 3684/ 0 / 245 |
| Goodness-of-fit on F2 | 1.07 |
| Final R indices | $R[F^2>2\sigma(F^2)] = 0.072$, $wR(F^2) = 0.195$ |
| Final R indices [all data] | R1 = 0.073, wR2 = 0.195 |
| Absolute structure parameter | 0.03 (8) |
| Largest diff. peak and hole | 0.58 and -0.51 e $Å^{-3}$ |

#### Crystallographic Methods and Data

**Experimental protocol:** Intensity data were collected on a Bruker APEX-II CCD diffractometer with a CMOS surface detector. A Cu-K source (= 1.54178 Å). Data was collected at 200K. The collection, cell refinement, and integration of intensity data was carried out with SAINT software. A semi-empirical absorption correction was performed with SADABS. The structure was phased with intrinsic methods using SHELXT and refined by the least-squares method.

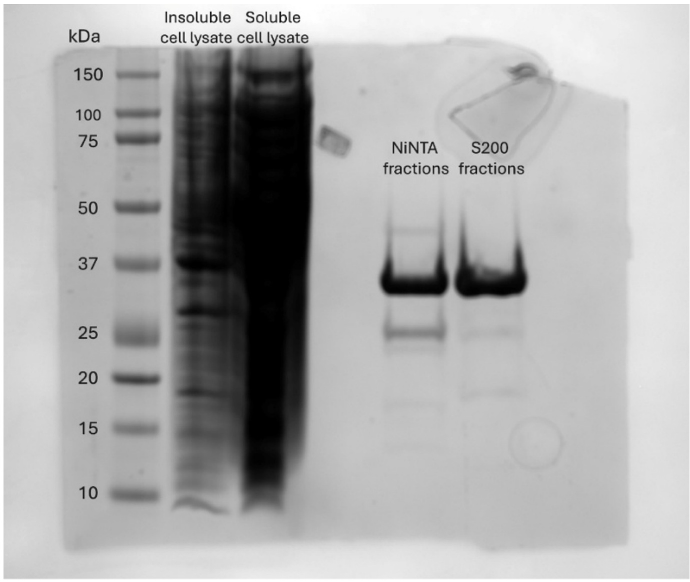


**Figure S3A.** SDS-PAGE gel demonstrating the presence of a pure protein at ~34 kDa, consistent with the expected mass of the *M. tuberculosis* pks13 thioesterase domain construct. The soluble cell lysate was purified using a Ni NTA affinity column, followed by the S200 size exclusion column. Pure fractions from the S200 were pooled for further analyses.

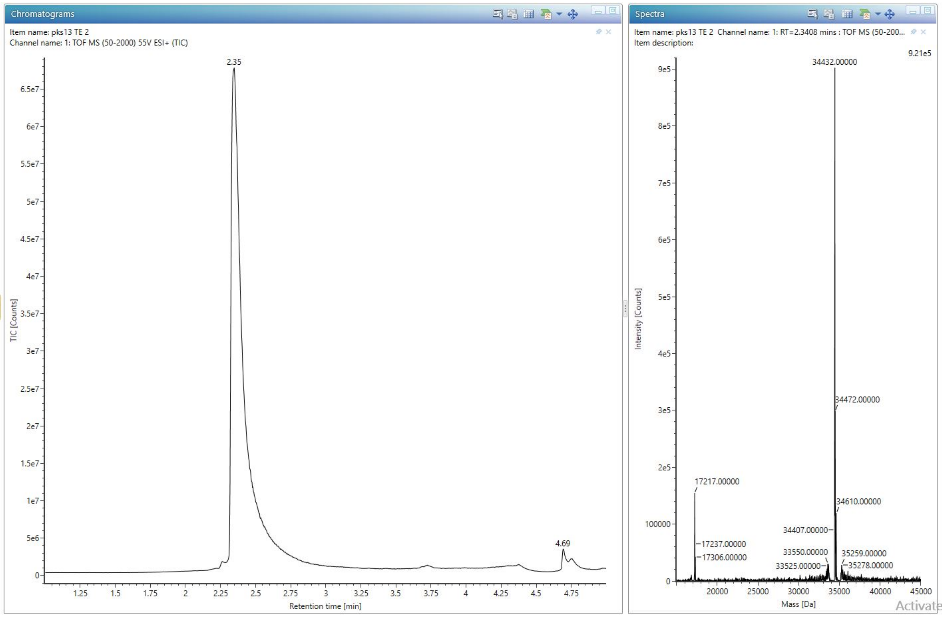


**Figure S3B.** Intact mass trace (Bioaccord) of the purified, recombinant, His-tagged *M. tuberculosis* pks13 thioesterase domain. The observed mass of 34432 Da corresponds with the expected mass of the domain (34432.46 Da), with the loss of the starting methionine residue.

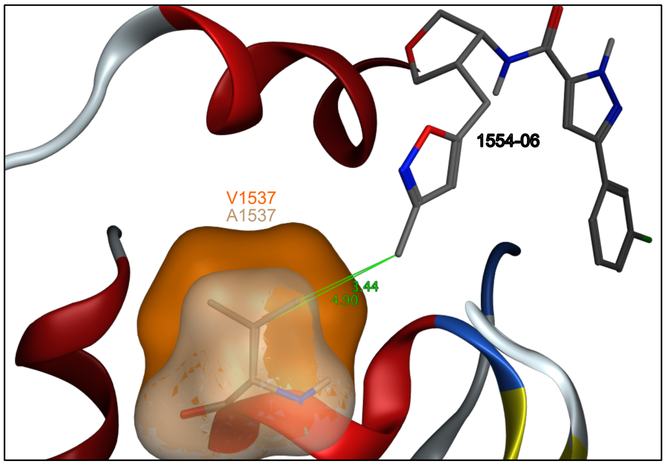


**Figure S4**

**Alanine substitution at site 1537 in Pks13 TE**. Potential size of hydrophobic patches provided by the valine (orange) and alanine (grey) sidechains at this position and proximity to 1554-06. attenuating hydrophobic contacts between the ligand and receptor.

**Table S1.** Activity data for selected analogs, pyrazole core.

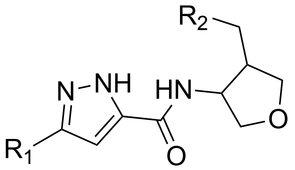


| Compound ID, (paper ID) | R1 | R2 | WT H37Rv MIC 90 (μM) |
|---|---|---|---|
| 1554-01 (**1-syn**)<br>1554-01-cis (**1-cis**) | 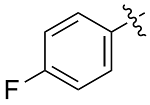 | 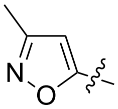 | 24.7<br>cis 13.3 |
| 1554-03 | 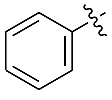 | 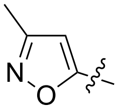 | 16.4 |
| 1554-04 | 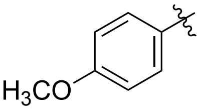 | 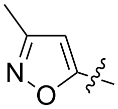 | 10.0 |
| 1554-05 | 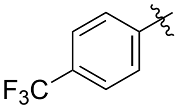 | 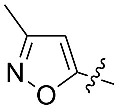 | 30.2 |
| 1554-06 (**5**)<br>1554-06-cis<br>1554-06-trans<br>1554-06-cis1 (**5a**)<br>1554-06-cis2 (**5b**)<br>1554-06-trans1(**5c**)<br>1554-06-trans2 (**5d**) | 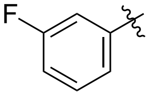 | 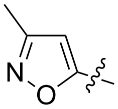 | 3.5<br>cis: 3.0<br>trans: 68.2<br>cis1: >100<br>cis2: 3.0<br>trans1: 53.7<br>trans2: >100 |
| 1554-13 | 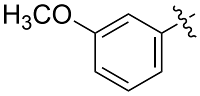 | 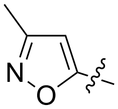 | 10.0 |
| 1554-14 | 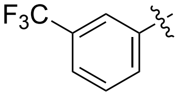 | 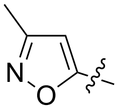 | 11.9 |
| 1554-15 | 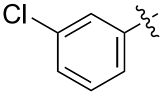 | 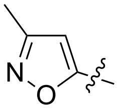 | 6.2 |
| 1554-07 | - | 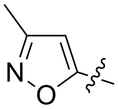 | 5.8 |
| 1554-08 | 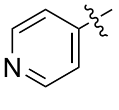 | 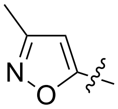 | 49.9 |

| | | | |
|---|---|---|---|
| 1554-17 | 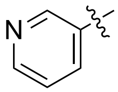 | 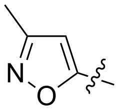 | >100 |
| 1554-18 | 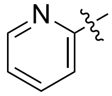 | 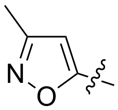 | >100 |
| 1554-11 | 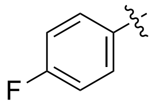 | 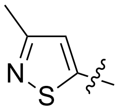 | >100 |
| 1554-23<br>1554-23-cis1<br>1554-23-cis2 (**7**)<br>1554-23-trans1<br>1554-23-trans2 | 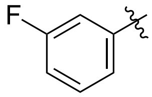 | 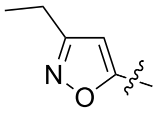 | 65.0<br>cis1: >100<br>cis2: 52.6<br>trans1: >100<br>trans2: >100 |
| 1554-31-cis (**6**)<br>1554-31-trans | 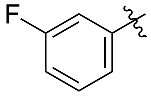 | 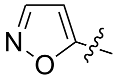 | cis: 41.7<br>trans: >100 |
| 1554-24<br>1554-24-cis1<br>1554-24-cis2<br>1554-24-trans1<br>1554-24-trans2 | 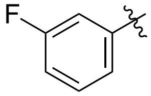 | 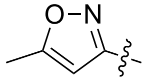 | 7.0<br>cis1: >100<br>cis2: 4.3<br>trans1: 51.6<br>trans2: >100 |
| 1554-26 | 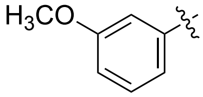 | 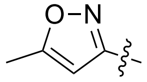 | 26.4 |
| 1554-25 | 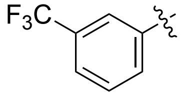 | 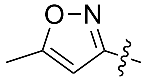 | 65.2 |
| 1554-34-cis<br>1554-34-trans | 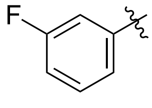 | 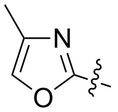 | cis: >100<br>trans: >100 |
| 1554-35-cis1 (**8**)<br>1554-35-cis2<br>1554-35-trans1<br>1554-35-trans2 | 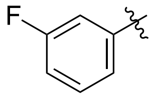 | 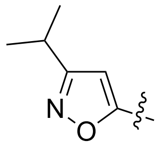 | cis1: 12.9<br>cis2: >100<br>trans1: >100<br>trans2: >100 |
| 1554-36-cis1<br>1554-36-cis2 (**9**)<br>1554-36-trans1 | 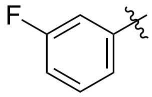 | 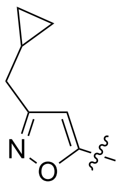 | cis1: >100<br>cis2: 51.4<br>trans1: 64.0 |

**Table S2.** HepG2 data for selected analogs

| **Compound ID (paper ID)** | **HepG2 IC50 (µM)** |
|---|---|
| 1554-01-cis (**1-cis**) | >120 |
| 1554-06-cis2 (1554-06-3R,4S) (5b) | >120 |
| 1554-06-trans1 (1554-06-3R,4R) (5c) | >120 |
| 1554-07 | >120 |
| 1554-13 | >120 |
| 1554-14 | >120 |
| 1554-15 | >120 |
| 1554-24-cis2 | >120 |

**Table S3.** SAR of selected analogs, alternate cores.

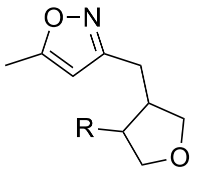


| Compound ID | R | WT H37Rv MIC 90 (µM) |
|---|---|---|
| 1554-02 | 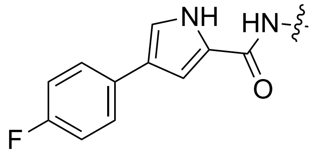 | >100 |
| 1554-10 | 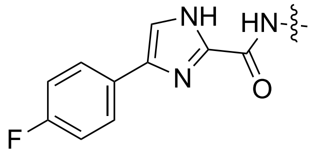 | >100 |
| 1554-12 | 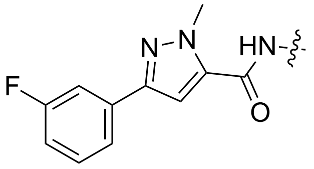 | >100 |
| 1554-19 | 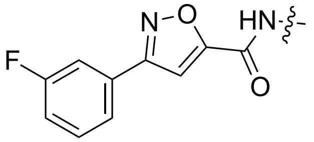 | >100 |

| | | |
|---|---|---|
| 1554-28 | 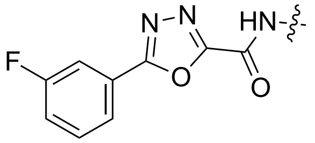 | >100 |
| 1554-09 | 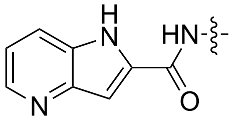 | >100 |
| 1554-21 | 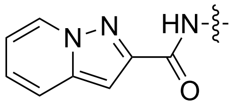 | >100 |
| 1554-22 | 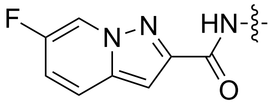 | >100 |
| 1554-27 | 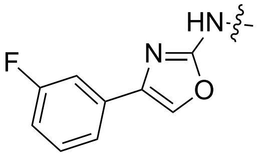 | >100 |
| 1554-29 | 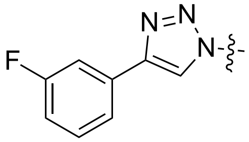 | >100 |

#### Experimental Procedures

#### Methods and Characterization

All chemical reagents were purchased from commercial sources and used without further purification. Flash chromatography was performed using silica gel (250-400 mesh). NMR spectra (including 2-D) were recorded on Bruker 400 spectrometer at 400 MHz for 1H NMR. Chemical shifts (δ scale) are reported in parts per million (ppm) relative to the central peak of the solvent. Coupling constants (J) are given in hertz (Hz). Spectra were obtained in the following solvents (reference peaks included for 1H): $CDCl_3$ (7.26 ppm), $CD_3OD$ (3.31 ppm) and DMSO-d6 (2.50 ppm). NMR experiments were performed at room temperature. Chemical shift values for all 1H NMR spectra are reported in parts per million (ppm). 1H NMR multiplicities are reported as: s = singlet, d = doublet, t = triplet, q = quartet, br = broad, m = multiplet. LCMS spectra were obtained on an Agilent 1260 series high performance liquid chromatography (HPLC) system operating in reverse-phase mode coupled to an Agilent 6125B mass spectrometer using an ESI source. Welch Boltimate EXT-C18 column (50 mm*4.6 mm*2.7 um), 40°C column temperature, Water with 0.05% TFA (mobile phase A), acetonitrile (mobile phase B), 1.8.0 mL/min flow rate, 10%B for 0.2 min, increase to 95%B within 0.9 min, 95%B for 0.9 min, back to 10%B within 1.3 min and UV absorbance detection at 190 to 400 nm.

#### Abbreviations

AgOTf: silver trifluoromethanesulfonate
$Boc_2O$: di-tert-butylcarbonate
Boc: tert-butylcarbonate
tBuOK: potassium tert-butoxide
tBuOH: tert-butanol
nBuLi: n-buthyllithium
DCM: dichoromethane
DIPEA: N,N-diisopropylethylamine
DMF: dimethylformamide
DMP: Dess-Martin periodinane
EtOH: ethanol
HATU: Hexafluorophosphate Azabenzotriazole Tetramethyl Uronium (O-(7-azabenzotriazol-1-yl)-N,N,N',N'-tetramethyluronium hexafluorophosphate)
LiHMDS: lithium bis(trimethylsilyl)amide
MeOH: methanol
NCS: N-chlorosuccinimide
$NH_4OAc$: ammonium acetate
$NEt_3$: triethylamine
PCC: pyridinium chlorochromate
$Pd(dppf)Cl_2$: [1,1'-Bis(diphenylphosphino)ferrocene]palladium(II) dichloride
PE: petroleum ether
RT: room temperature
TFA: trifluoroacetic acid
TsCl: para-toluenesulfonyl chloride
THF: tetrahydrofuran
pTsOH: para-toluenesulfonic acid
TosMIC: toluenesulfonylmethyl isocyanide

#### Schemes, Synthesis, and Characterization of 1554 series analogs

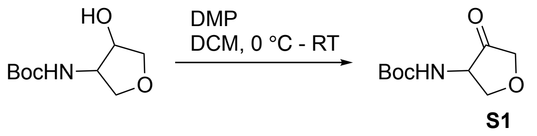


##### Procedure for preparation of tert-butyl (4-oxotetrahydrofuran-3-yl)carbamate (S1)

To a solution of tert-butyl (4-hydroxytetrahydrofuran-3-yl)carbamate (2.50 g, 12.30 mmol) in DCM (50 mL) was added DMP (6.26 g, 14.76 mmol) at 0 ºC. The resulting mixture was stirred at 25ºC for 4.0 h. The reaction mixture was concentrated in vacuo. The residue was filtered and rinsed with ethyl acetate. The mixture was diluted with water and extracted with ethyl acetate (40 mLx3). The combined organic layers were washed with brine, dried over anhydrous $Na_2SO_4$ and concentrated in vacuo. The residue was purified by silica gel column chromatography (eluted with PE/ethyl acetate = 5/1) to give tert-butyl (4-oxotetrahydrofuran-3-yl)carbamate (**S1**, 2.10 g, 85% yield) as a white solid.

LC-MS (ESI, *m/z*): $[M+H-C4H8]^+$ = 146.1.

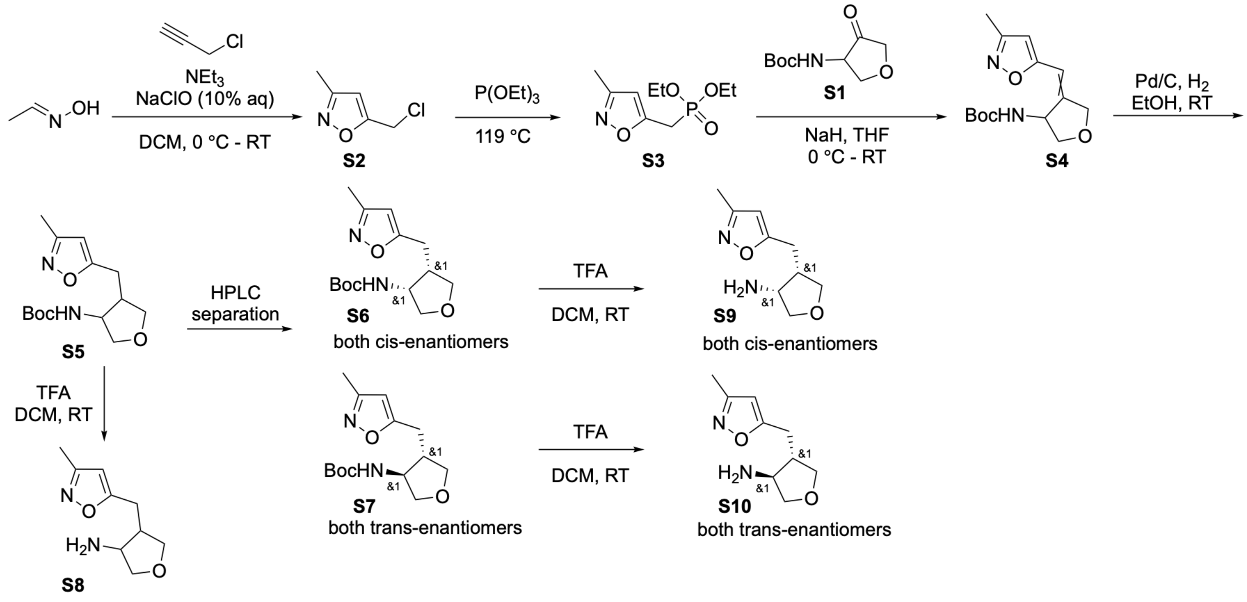


##### Procedure for preparation of 5-(chloromethyl)-3-methylisoxazole (S2)

To a mixture of (*E*)-acetaldehyde oxime (1.0 g, 16.93 mmol), 3-chloroprop-1-yne (1.27 g, 17.10 mmol), and $Et_3N$ (0.94 mL, 6.77 mmol) in DCM (8 mL) was added NaClO (aq.) (10%, 39 mL) at 0 ºC. The resulting mixture was stirred at 25 ºC overnight. The mixture was quenched with water and extracted with DCM (30 mLx3). The combined organic layers were washed with brine, dried over anhydrous $Na_2SO_4$ and concentrated in vacuo. The residue was purified by silica gel column chromatography (eluted with PE/ethyl acetate = 5/1) to give 5-(chloromethyl)-3-methylisoxazole (**S2**, 282 mg, 13% yield) as a colorless oil.

LC-MS (ESI, *m/z*): $[M+H]^+$ = 132.0.

**Procedure for preparation of diethyl ((3-methylisoxazol-5-yl)methyl)phosphonate (S3)**

A solution of 5-(chloromethyl)-3-methylisoxazole (**S2**, 282 mg, 2.14 mmol) and $P(OEt)_3$ (1.84 mL, 10.72 mmol) was stirred at 119 °C overnight under $N_2$ atmosphere. The reaction mixture was concentrated in vacuo. The residue was purified by silica gel chromatography (eluted with DCM/MeOH = 20/1) to give diethyl ((3-methylisoxazol-5-yl)methyl)phosphonate (**S3**, 406 mg, 81% yield) as a colorless oil.

LC-MS (ESI, *m/z*): $[M+H]^+$ = 234.1.

**Procedure for preparation of tert-butyl (4-((3-methylisoxazol-5-yl)methylene)tetrahydrofuran-3-yl)carbamate (S4)**

To a solution of diethyl ((3-methylisoxazol-5-yl)methyl)phosphonate (**S3**, 159 mg, 0.68 mmol) in THF (2 mL) was added NaH (60%, 27 mg, 0.68 mmol) at 0 °C. The resulting mixture was stirred at room temperature for 0.5 h. Then, tert-butyl (4-oxotetrahydrofuran-3-yl)carbamate (114 mg, 0.57 mmol) was added. The resulting mixture was stirred at room temperature overnight. The reaction mixture was diluted with water (30 mL) and extracted with ethyl acetate (30 mLx3). The combined organic layers were washed with brine, dried over anhydrous $Na_2SO_4$ and concentrated in vacuo. The residue was purified by silica gel column chromatography (eluted with PE/ethyl acetate = 4/1) to give tert-butyl (4-((3-methylisoxazol-5-yl)methylene)tetrahydrofuran-3-yl)carbamate (**S4**, 50 mg, 31% yield) as a colorless oil.

LC-MS (ESI, *m/z*): $[M+H-C4H8]^+$ = 225.1.

**Procedure for preparation of tert-butyl (4-((3-methylisoxazol-5-yl)methyl)tetrahydrofuran-3-yl)carbamate (S5) and separation of cis (3R,4S; 3S,4R) (S6) and trans (3R,4R; 3S,4S) (S7) isomeric mixtures**

**For diastereomeric mixture and cis isomers:** To a mixture of tert-butyl (4-((3-methylisoxazol-5-yl)methylene)tetrahydrofuran-3-yl)carbamate (**S4**, 817 mg, 2.91 mmol) in EtOH (40 mL) was added Pd/C (10% on carbon, 408 mg), and the mixture placed under $H_2$ atmosphere. The mixture was stirred at room temperature for 4.0 h. The reaction mixture was filtered on Celite and rinsed with MeOH. The solvent was concentrated in vacuo. The residue was purified by prep-HPLC (eluted with $CH_3CN/H_2O$ = 5/95~ 95/5 with 0.1% TFA) to give tert-butyl (4-((3-methylisoxazol-5-yl)methyl)tetrahydrofuran-3-yl)carbamate

(**S5**, 192 mg, 23% yield, mixture of 4 diastereomers) as a colorless oil and a mixture of tert-butyl ((3R,4S)-4-((3-methylisoxazol-5-yl)methyl)tetrahydrofuran-3-yl)carbamate and ((3S,4R)-4-((3-methylisoxazol-5-yl)methyl)tetrahydrofuran-3-yl)carbamate (**S6**, 116 mg, 14% yield, cis isomers) as a colorless oil.

LC-MS (ESI, *m/z*): $[M+H-C4H8]^+$ = 227.1.

**For cis and trans isomers:**

To a mixture of tert-butyl (4-((3-methylisoxazol-5-yl)methylene)tetrahydrofuran-3-yl)carbamate (**S4**, 350 mg, 1.25 mmol) in EtOH (18 mL) was added Pd/C (10% on carbon, 175 mg) and the mixture placed under $H_2$ atmosphere. The mixture was stirred at room temperature for 4.0 h. The reaction mixture was filtered on Celite and rinsed with MeOH. The solvent was concentrated in vacuo. The residue was purified by prep-HPLC (eluted with $CH_3CN/H_2O$ = 5/95~ 95/5 with 0.1% TFA) to give a mixture of tert-butyl 3R,4S)-4-((3-methylisoxazol-5-yl)methyl)tetrahydrofuran-3-yl)carbamate and tert-butyl ((3S,4R)-4-((3-methylisoxazol-5-yl)methyl)tetrahydrofuran-3-yl)carbamate (**S6**, 36 mg, 10% yield, cis isomers) as a colorless oil and a mixture of tert-butyl ((3R,4R)-4-((3-methylisoxazol-5-yl)methyl)tetrahydrofuran-3-yl)carbamate and tert-butyl ((3S,4S)-4-((3-methylisoxazol-5-yl)methyl)tetrahydrofuran-3-yl)carbamate (**S7**, 29 mg, 8% yield, trans isomers) as a colorless oil.

LC-MS (ESI, *m/z*): $[M+H-C4H8]^+$ = 227.1.

**Procedure for preparation of 4-((3-methylisoxazol-5-yl)methyl)tetrahydrofuran-3-amine (S8)**

A mixture of tert-butyl (4-((3-methylisoxazol-5-yl)methyl)tetrahydrofuran-3-yl)carbamate (**S5**, 34 mg, 0.12 mmol) in DCM (2 mL) and TFA (1 mL) was stirred at room temperature for 1.0 h. The reaction mixture was concentrated in vacuo to give 63 mg of 4-((3-methylisoxazol-5-yl)methyl)tetrahydrofuran-3-amine (**S8**) TFA salt crude as a colorless oil.

LC-MS (ESI, *m/z*): $[M+H]^+$ = 183.1.

**Procedure for preparation of a mixture of (3R,4S)-4-((3-methylisoxazol-5-yl)methyl)tetrahydrofuran-3-amine and (3S,4R)-4-((3-methylisoxazol-5-yl)methyl)tetrahydrofuran-3-amine (cis isomers) (S9)**

A mixture of tert-butyl ((3R,4S)-4-((3-methylisoxazol-5-yl)methyl)tetrahydrofuran-3-yl)carbamate and tert-butyl ((3S,4R)-4-((3-methylisoxazol-5-yl)methyl)tetrahydrofuran-3-yl)carbamate (**S6**, 116 mg, 0.41 mmol) in DCM (2 mL) and TFA (2 mL) was stirred at room temperature for 1.0 h. The reaction mixture was concentrated in vacuo to give 171 mg of (3R,4S)-4-((3-methylisoxazol-5-yl)methyl)tetrahydrofuran-

3-amine and 3S,4R)-4-((3-methylisoxazol-5-yl)methyl)tetrahydrofuran-3-amine (**S9**) TFA salt as a colorless oil.

LC-MS (ESI, *m/z*): $[M+H]^+$ = 183.1.

**Procedure for preparation of a mixture of (3R,4R)-4-((3-methylisoxazol-5-yl)methyl)tetrahydrofuran-3-amine and (3S,4S)-4-((3-methylisoxazol-5-yl)methyl)tetrahydrofuran-3-amine (trans isomers) (S10)**

A mixture of tert-butyl ((3S,4S)-4-((3-methylisoxazol-5-yl)methyl)tetrahydrofuran-3-yl)carbamate and ((3R,4R)-4-((3-methylisoxazol-5-yl)methyl)tetrahydrofuran-3-yl)carbamate (**S7**, 36 mg, 0.13 mmol) in DCM (3 mL) and TFA (1 mL) was stirred at room temperature for 1.0 h. The reaction mixture was concentrated in vacuo to give 53 mg of (3S,4S)-4-((3-methylisoxazol-5-yl)methyl)tetrahydrofuran-3-amine and (3R,4R)-4-((3-methylisoxazol-5-yl)methyl)tetrahydrofuran-3-amine (**S10**) TFA salt as a colorless oil.

LC-MS (ESI, *m/z*): $[M+H]^+$ = 183.1.

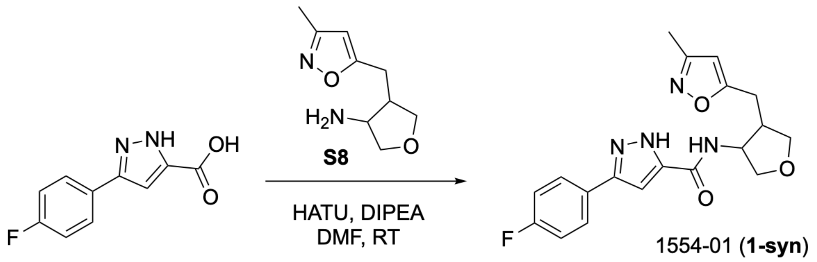


**Procedure for preparation of 3-(4-fluorophenyl)-N-(4-((3-methylisoxazol-5-yl)methyl)tetrahydrofuran-3-yl)-1H-pyrazole-5-carboxamide (1554-01, 1-syn)**

A mixture of 4-((3-methylisoxazol-5-yl)methyl)tetrahydrofuran-3-amine (**S8**, 63 mg, 0.21 mmol), 3-(4-fluorophenyl)-1H-pyrazole-5-carboxylic acid (66 mg, 0.32 mmol), DIPEA (137 mg, 1.06 mmol), HATU (121 mg, 0.32 mmol) and dry DMF (2.0 mL) was stirred at room temperature overnight. The reaction mixture was concentrated in vacuo. The residue was purified by reverse phase silica gel column chromatography (eluted with $CH_3CN/H_2O$ = 5/95 ~ 95/5 with 0.1% TFA) to give 3-(4-fluorophenyl)-N-(4-((3-methylisoxazol-5-yl)methyl)tetrahydrofuran-3-yl)-1H-pyrazole-5-carboxamide (**1554-01**, 18 mg, 31% yield for two steps) TFA salt as a white solid.

LC-MS (ESI, *m/z*): $[M+H]^+$ = 371.2.

$^{1}$H NMR (400 MHz, CD3OD, cis/trans = 5/2) δ 7.77 – 7.73 (m, 2.81H), 7.23 – 7.15 (m, 2.83H), 7.06 (s, 0.92H), 7.02 (s, 0.38H), 6.13 (s, 0.36H), 6.04 (s, 1.00H), 4.84 – 4.81 (m, 1.06H), 4.41 (q, *J* = 6.0 Hz, 0.37H), 4.14 – 4.06 (m, 1.80H), 4.04 – 4.00 (m, 1.05H), 3.82 (dd, *J* = 9.2, 3.6 Hz, 1.06H), 3.73 – 3.65 (m, 1.44H), 3.60 (dd, *J* = 8.8, 6.4 Hz, 0.39H), 3.09 – 3.03 (m, 0.36H), 3.02 – 2.97 (m, 1.08H), 2.94 – 2.84 (m, 1.60H), 2.82 – 2.76 (m, 1.05H), 2.72 – 2.67 (m, 0.43H), 2.19 (s, 2.94H), 2.18 (s, 1.21H).

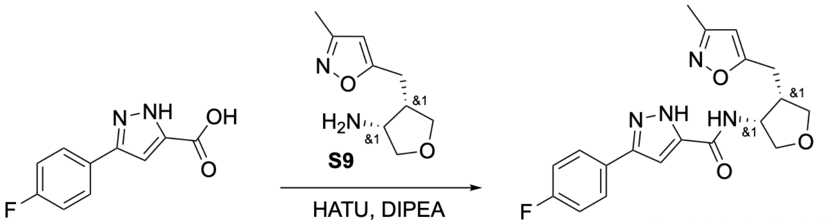

1554-01-cis (**1-cis**)

**Procedure for preparation of 3-(4-fluorophenyl)-N-((3R,4S)-4-((3-methylisoxazol-5-yl)methyl)tetrahydrofuran-3-yl)-1H-pyrazole-5-carboxamide and 3-(4-fluorophenyl)-N-((3S,4R)-4-((3-methylisoxazol-5-yl)methyl)tetrahydrofuran-3-yl)-1H-pyrazole-5-carboxamide (cis isomers) (1554-01-cis, 1-cis)**

A mixture of **S9** (70 mg, 0.17 mmol), 3-(4-fluorophenyl)-1H-pyrazole-5-carboxylic acid (42 mg, 0.20 mmol), DIPEA (110 mg, 0.85 mmol), HATU (78 mg, 0.20 mmol) and dry DMF (3.0 mL) was stirred at room temperature overnight. The reaction mixture was concentrated in vacuo. The residue was purified by reverse phase silica gel column chromatography (eluted with $CH_3CN/H_2O$ = 5/95 ~ 95/5 with 0.1% TFA) to give 3-(4-fluorophenyl)-N-((3R,4S)-4-((3-methylisoxazol-5-yl)methyl)tetrahydrofuran-3-yl)-1H-pyrazole-5-carboxamide (**1554-01-cis**, 34 mg, 41% yield) TFA salt as a white solid.

LC-MS (ESI, *m/z*): $[M+H]^+$ = 371.2.

$^{1}$H NMR (400 MHz, CD3OD, cis) δ 7.78 – 7.73 (m, 2H), 7.23 – 7.17 (m, 2H), 7.06 (s, 1H), 6.04 (s, 1H), 4.84 – 4.81 (m, 1H), 4.09 (dd, *J* = 9.6, 6.0 Hz, 1H), 4.02 (dd, *J* = 8.8, 7.6 Hz, 1H), 3.82 (dd, *J* = 9.6, 4.0 Hz, 1H), 3.68 (t, *J* = 8.4 Hz, 1H), 3.00 (dd, *J* = 14.8, 6.0 Hz, 1H), 2.94 – 2.84 (m, 1H), 2.79 (dd, *J* = 14.8, 8.8 Hz, 1H), 2.19 (s, 3H).

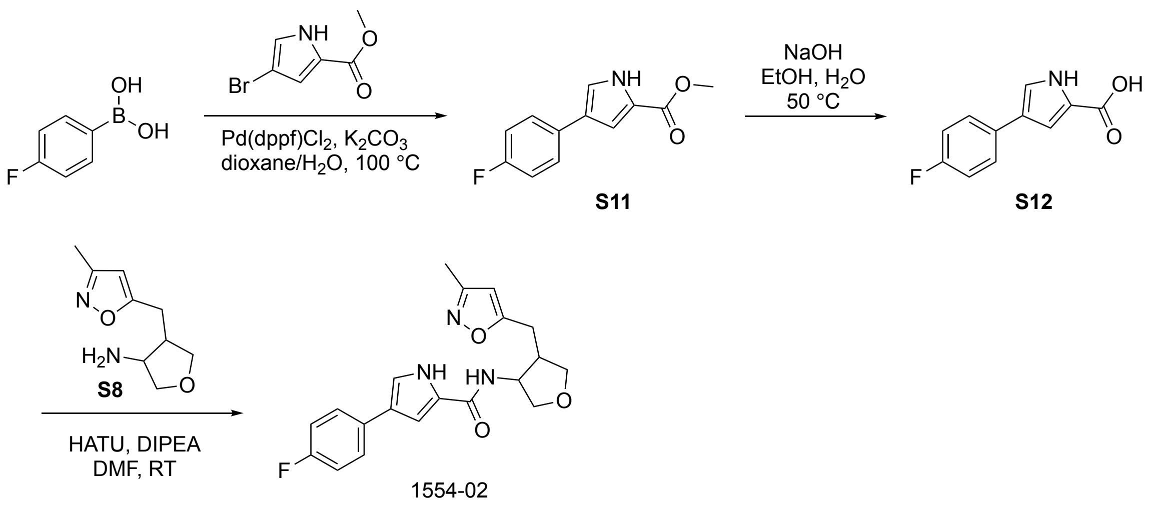


**Procedure for preparation of methyl 4-(4-fluorophenyl)-1H-pyrrole-2-carboxylate (S11)**

To a mixture of methyl 4-bromo-1H-pyrrole-2-carboxylate (200 mg, 0.98 mmol), (4-fluorophenyl)boronic acid (178 mg, 1.27 mmol) and Pd(dppf)Cl$_2$ (48 mg, 0.06 mmol) was added $K_2CO_3$ (271 mg, 1.96 mmol) in Dioxane/$H_2O$ (4/1 mL) at room temperature under $N_2$ atmosphere. The resulting mixture was stirred at 100 °C overnight. The reaction mixture was diluted with $H_2O$ (50 mL) and extracted with ethyl acetate (30 mLx3). The combined organic layers were washed with brine, dried over anhydrous $Na_2SO_4$ and concentrated in vacuo. The residue was purified by silica gel column chromatography (eluted with PE/ethyl acetate = 2/1) to give methyl 4-(4-fluorophenyl)-1H-pyrrole-2-carboxylate (**S11**, 109 mg, 51% yield) as a white solid.

LC-MS: MS(ESI, m/z): $[M+H]^+$=220.1.

**Procedure for preparation of 4-(4-fluorophenyl)-1H-pyrrole-2-carboxylic acid (S12)**

A mixture of **S11** (109 mg, 0.50 mmol), NaOH (199 mg, 4.97 mmol) and EtOH/$H_2O$ (2 /2 mL) was stirred at 50 °C for 1.0 h. The reaction mixture was acidified with HCl (aqueous, 2 N) to pH 4. The mixture was purified by reverse phase silica gel column chromatography (eluted with $CH_3CN/H_2O$ = 5/95~95/5 with 0.1% HCl) to give 4-(4-fluorophenyl)-1H-pyrrole-2-carboxylic acid (**S12**, 110 mg, 91% yield) as a yellow oil.

LC-MS: MS(ESI, m/z): $[M+H]^+$=206.1

**Procedure for preparation of 4-(4-fluorophenyl)-N-(4-((3-methylisoxazol-5-yl)methyl)tetrahydrofuran-3-yl)-1H-pyrrole-2-carboxamide (1554-02)**

A mixture of 4-((3-methylisoxazol-5-yl)methyl)tetrahydrofuran-3-amine (**S8**, 40 mg, 0.10 mmol), 4-(4-fluorophenyl)-1H-pyrrole-2-carboxylic acid (**S12**, 28 mg, 0.12 mmol), HATU (44 mg, 0.12 mmol),

DIPEA (63 mg, 0.49 mmol) and dry DMF (2.0 mL) was stirred room temperature for 1.0 h. The reaction mixture was purified by reverse phase silica gel column chromatography (eluted with $CH_3CN/H_2O$ = 5/95~95/5 with 0.1% TFA) to give 4-(4-fluorophenyl)-N-(4-((3-methylisoxazol-5-yl)methyl)tetrahydrofuran-3-yl)-1H-pyrrole-2-carboxamide (**1554-02**, 36 mg, 39% yield, cis/trans=5/3) as a white solid.

LC-MS: MS(ESI, m/z): $[M+H]^+$=370.1

1H NMR (400 MHz, Methanol-d4) δ 7.54 (ddd, J = 8.8, 4.4, 1.6 Hz, 3H), 7.25 (d, J = 1.7 Hz, 1H), 7.23 (d, J = 1.7 Hz, 1H), 7.20 (d, J = 1.8 Hz, 1H), 7.09 (d, J = 1.7 Hz, 1H), 7.07 (d, J = 2.3 Hz, 1H), 7.05 (d, J = 2.3 Hz, 2H), 7.03 (d, J = 2.3 Hz, 1H), 6.10 (s, 1H), 6.03 (s, 1H), 4.81 (dt, J = 6.6, 3.3 Hz, 1H), 4.37 (q, J = 6.1 Hz, 1H), 4.10 (ddd, J = 9.2, 6.8, 5.1 Hz, 2H), 4.00 (dd, J = 8.7, 7.3 Hz, 1H), 3.79 (dd, J = 9.3, 4.3 Hz, 1H), 3.71 – 3.64 (m, 2H), 3.59 (dd, J = 8.9, 6.8 Hz, 1H), 3.04 (dd, J = 15.4, 6.8 Hz, 1H), 2.98 (dd, J = 15.0, 5.6 Hz, 1H), 2.93 – 2.83 (m, 2H), 2.75 (dd, J = 15.0, 9.1 Hz, 1H), 2.70 – 2.63 (m, 1H), 2.18 (s, 3H), 2.16 (s, 2H).

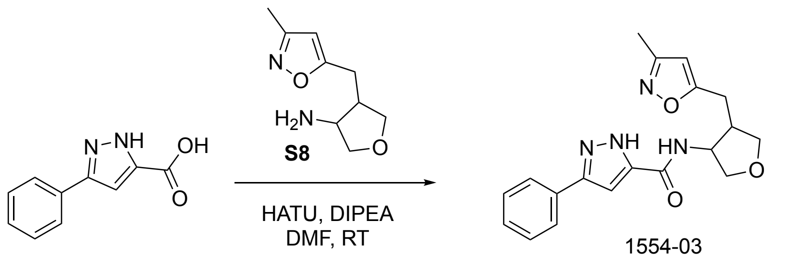


**Procedure for preparation of N-(4-((3-methylisoxazol-5-yl)methyl)tetrahydrofuran-3-yl)-3-phenyl-1H-pyrazole-5-carboxamide (1554-03)**

A mixture of 4-((3-methylisoxazol-5-yl)methyl)tetrahydrofuran-3-amine (**S8**, 50 mg, 0.12 mmol), 3-phenyl-1H-pyrazole-5-carboxylic acid (28 mg, 0.15 mmol), DIPEA (79 mg, 0.61 mmol), HATU (56 mg, 0.15 mmol) and dry DMF (2.0 mL) was stirred at room temperature for 3.0 h. The reaction mixture was purified by reverse phase silica gel column chromatography (eluted with $CH_3CN/H_2O$ = 5/95 ~ 95/5 with 0.1% TFA) to give N-(4-((3-methylisoxazol-5-yl)methyl)tetrahydrofuran-3-yl)-3-phenyl-1H-pyrazole-5-carboxamide (**1554-03**, 31 mg, 55% yield) TFA salt as a white solid.

LC-MS (ESI, *m/z*): $[M+H]^+$ = 353.2.

$^1$H NMR (400 MHz, CD3OD, cis/trans = 20/7) δ 7.74 – 7.71 (m, 2.80H), 7.48 – 7.43 (m, 2.82H), 7.40 – 7.37 (m, 1.02H), 7.37 – 7.35 (m, 0.35H), 7.09 (s, 1.00H), 7.05 (s, 0.35H), 6.13 (s, 0.35H), 6.04 (s, 1.00H), 4.85 – 4.81 (m, 1.00H), 4.42 (p, *J* = 6.4 Hz, 0.36H), 4.14 – 4.08 (m, 1.82H), 4.02 (dd, *J* = 8.4, 7.6 Hz,

1.07H), 3.82 (dd, $J$ = 9.2, 3.6 Hz, 1.02H), 3.72 – 3.67 (m, 1.44H), 3.60 (dd, $J$ = 8.8, 6.4 Hz, 0.36H), 3.09 – 3.04 (m, 0.35H), 3.00 (dd, $J$ = 14.8, 9.6 Hz, 1.06H), 2.94 – 2.85 (m, 1.52H), 2.79 (dd, $J$ = 14.8, 8.8 Hz, 1.07H), 2.70 (dd, $J$ = 13.6, 6.8 Hz, 0.38H), 2.19 (s, 3.00H), 2.19 (s, 1.05H).

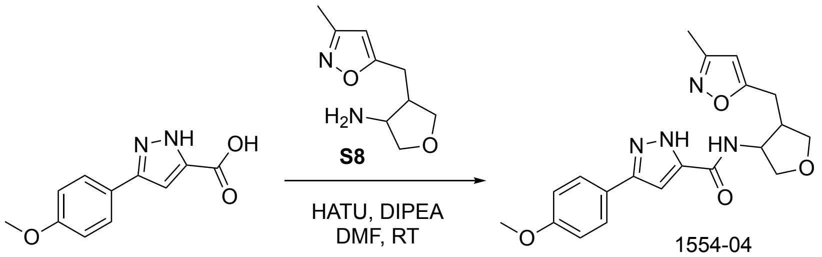


**Procedure for preparation of 3-(4-methoxyphenyl)-N-(4-((3-methylisoxazol-5-yl)methyl)tetrahydrofuran-3-yl)-1H-pyrazole-5-carboxamide (1554-04)**

To a mixture of 3-(4-methoxyphenyl)-1H-pyrazole-5-carboxylic acid (**S8**, 50 mg, 0.23 mmol, 1.0 eq.), 4-((3-methylisoxazol-5-yl)methyl)tetrahydrofuran-3-amine (100 mg, TFA salt, 0.23 mmol, 1.0 eq.) in DMF (2 mL) was added DIPEA (0.5 mL) and HATU (92 mg, 0.24 mmol, 1.05 eq), the resulting mixture was stirred at room temperature for 2 h. The reaction mixture was diluted with water (20 mL), extracted with ethyl acetate (20 mL×3). The combined organic phases were dried over anhydrous $Na_2SO_4$ and concentrated under vacuum. The residue was purified by silica gel column chromatography (eluted with $CH_3CN/H_2O$ = 5/95 ~ 95/5) to give 3-(4-methoxyphenyl)-N-(4-((3-methylisoxazol-5-yl)methyl)tetrahydrofuran-3-yl)-1H-pyrazole-5-carboxamide (**1554-04**, 29 mg, 33% yield) as a white solid.

LC-MS (ESI, *m/z*): $[M+H]^+$ = 383.2.

$^1$H NMR (400 MHz, Methanol-d4)(cis:trans=10:7) δ7.76 – 7.56 (m, 3.4H), 7.05 – 6.96 (m, 4H), 6.92 (d, $J$ = 12.0 Hz, 1H), 6.13 (s, 1H), 6.04 (s, 1H), 4.82 (dd, J = 6.4, 3.7 Hz, 1H), 4.41 (q, J = 5.9 Hz, 1H), 4.09 (ddd, J = 9.3, 6.3, 2.5 Hz, 3H), 4.02 (dd, J = 8.7, 7.4 Hz, 1H), 3.83 (d, J = 1.0 Hz, 5H), 3.83 – 3.78 (m, 1H), 3.74 – 3.65 (m, 2H), 3.60 (dd, J = 8.9, 6.6 Hz, 1H), 3.09 – 2.84 (m, 4H), 2.84 – 2.64 (m, 2H), 2.19 (d, J = 1.5 Hz, 5H).

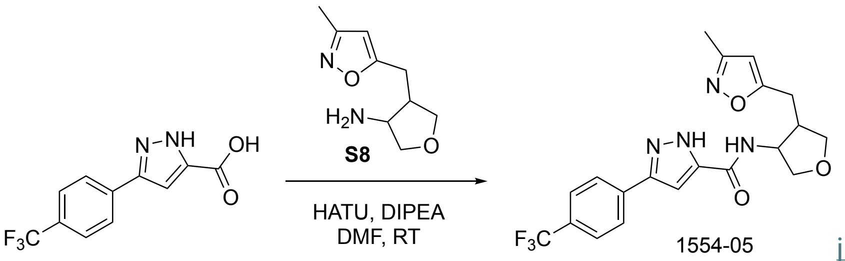


javascript:

**Procedure for the preparation of N-(4-((3-methylisoxazol-5-yl)methyl)tetrahydrofuran-3-yl)-3-(4-(trifluoromethyl)phenyl)-1H-pyrazole-5-carboxamide (1554-05)**

A mixture of 3-(4-methoxyphenyl)-1H-pyrazole-5-carboxylic acid (**S8**, 50 mg, 0.12 mmol), 3-(4-(trifluoromethyl)phenyl)-1*H*-pyrazole-5-carboxylic acid (37 mg, 0.15 mmol), HATU (56 mg, 0.15 mmol), DIPEA (79 mg, 061 mmol) and dry DMF (2.0 mL) was stirred room temperature for 1.0 h. The reaction mixture was purified by reverse phase silica gel column chromatography (eluted with $CH_3CN/H_2O$ = 5/95~95/5) to give N-(4-((3-methylisoxazol-5-yl)methyl)tetrahydrofuran-3-yl)-3-(4-(trifluoromethyl)phenyl)-1H-pyrazole-5-carboxamide (**1554-05**, 26 mg, 51% yield, cis/trans=4/3) as a white solid.

LC-MS: MS(ESI, m/z): $[M+H]^+$=421.1.

$^1$H NMR (400 MHz, DMSO-$d_6$) δ 13.87 (s, 2H), 8.72 – 8.53 (m, 1H), 8.46 – 8.23 (m, 1H), 8.02 (s, 2H), 8.00 (s, 1H), 7.85 (s, 1H), 7.82 (s, 2H), 7.80 (s, 1H), 7.47 (d, *J* = 46.7 Hz, 1H), 7.26 (d, *J* = 12.3 Hz, 1H), 6.20 (s, 1H), 6.13 (s, 1H), 4.80 – 4.70 (m, 1H), 4.30 – 4.24 (m, 1H), 4.05 – 3.98 (m, 2H), 3.91 (d, *J* = 5.7 Hz, 1H), 3.72 (dd, *J* = 8.4, 3.3 Hz, 1H), 3.62 – 3.56 (m, 2H), 3.47 (d, *J* = 7.9 Hz, 1H), 3.01 (dd, *J* = 13.6, 4.3 Hz, 1H), 2.86 (dd, *J* = 14.3, 10.2 Hz, 2H), 2.78 – 2.71 (m, 2H), 2.69 (d, *J* = 10.7 Hz, 1H), 2.15 (s, 2H), 2.14 (s, 3H).

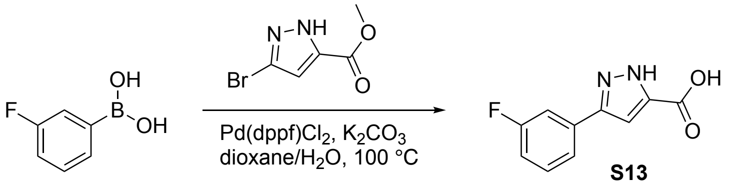


javascript:

**Procedure for the preparation of 3-(3-fluorophenyl)-1H-pyrazole-5-carboxylic acid (S13)**

To a mixture of methyl 3-bromo-1H-pyrazole-5-carboxylate (100 mg, 0.49 mmol), (3-fluorophenyl)boronic acid (89 mg, 0.63 mmol) and Pd(dppf)$Cl_2$ (24 mg, 0.03 mmol) was added $Na_2CO_3$ (103 mg, 0.98 mmol) in Dioxane/$H_2O$ (4/1 mL) at room temperature under $N_2$ atmosphere. The resulting mixture was stirred at 100 ºC overnight. The reaction mixture was purified by reverse phase silica gel column chromatography (eluted with $CH_3CN/H_2O$ = 5/95~95/5 with 0.1% HCl) to give 3-(3-fluorophenyl)-1H-pyrazole-5-carboxylic acid (**S13**, 36 mg, HCl salt, 15% yield) as a yellow oil.

LC-MS: MS(ESI, m/z): $[M+H]^+$=207.1

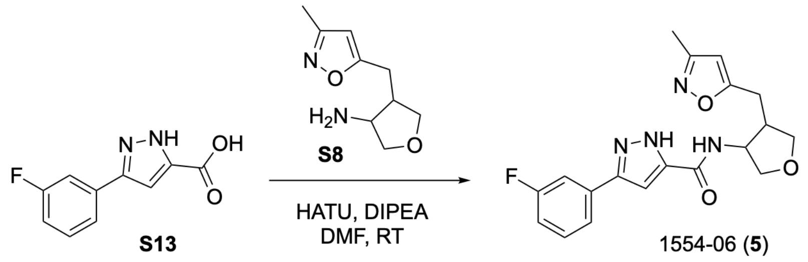


**Procedure for the preparation of 3-(3-fluorophenyl)-N-(4-((3-methylisoxazol-5-yl)methyl)tetrahydrofuran-3-yl)-1H-pyrazole-5-carboxamide (1554-06, 5)**

A mixture of 4-((3-methylisoxazol-5-yl)methyl)tetrahydrofuran-3-amine (**S8**, 50 mg, 0.12 mmol), 3-(3-fluorophenyl)-1H-pyrazole-5-carboxylic acid (**S13**, 36 mg, 0.15 mmol), HATU (56 mg, 0.15 mmol), DIPEA (79 mg, 0.61 mmol) and dry DMF (2.0 mL) was stirred room temperature for 1.0 h. The reaction mixture was purified by reverse phase silica gel column chromatography (eluted with $CH_3CN/H_2O$ = 5/95~95/5 with 0.1% TFA) to give 3-(3-fluorophenyl)-N-(4-((3-methylisoxazol-5-yl)methyl)tetrahydrofuran-3-yl)-1H-pyrazole-5-carboxamide (**1554-06**, 44 mg, TFA salt, 60% yield, cis/trans=5/3) as a white solid.

LC-MS: MS(ESI, m/z): $[M+H]^+$=371.1

1H NMR (400 MHz, Methanol-d4) δ 7.56 (dd, J = 7.7, 4.0 Hz, 2H), 7.52 – 7.51 (m, 1H), 7.49 (dd, J = 5.4, 2.8 Hz, 2H), 7.47 – 7.43 (m, 1H), 7.16 – 7.13 (m, 1H), 7.13 – 7.07 (m, 2H), 6.13 (s, 1H), 6.04 (s, 1H), 4.84 – 4.81 (m, 1H), 4.41 (q, J = 5.9 Hz, 1H), 4.14 – 4.07 (m, 2H), 4.02 (dd, J = 8.7, 7.4 Hz, 1H), 3.82 (dd, J = 9.3, 3.8 Hz, 1H), 3.72 – 3.66 (m, 2H), 3.62 – 3.58 (m, 1H), 3.06 (dd, J = 15.3, 6.8 Hz, 1H), 3.00 (dd, J = 14.9, 5.9 Hz, 1H), 2.90 (dt, J = 13.1, 7.5 Hz, 2H), 2.79 (dd, J = 14.9, 8.8 Hz, 1H), 2.73 – 2.66 (m, 1H), 2.19 (s, 3H), 2.18 (s, 2H).

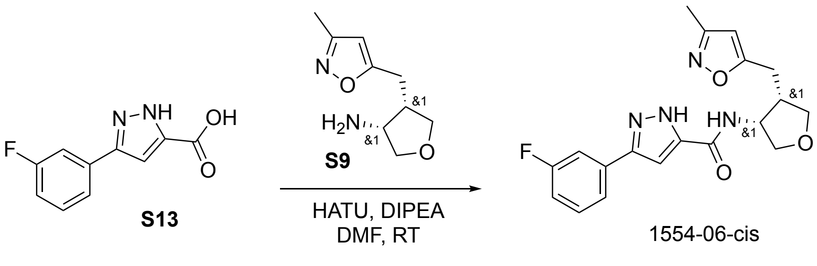


**Procedure for preparation of 3-(3-fluorophenyl)-N-((3S,4R)-4-((3-methylisoxazol-5-yl)methyl)tetrahydrofuran-3-yl)-1H-pyrazole-5-carboxamide and 3-(3-fluorophenyl)-N-((3R,4S)-4-((3-methylisoxazol-5-yl)methyl)tetrahydrofuran-3-yl)-1H-pyrazole-5-carboxamide (cis isomers) (1554-06-cis)**

A mixture of **S9** (53 mg, 0.13 mmol), 3-(3-fluorophenyl)-1H-pyrazole-5-carboxylic acid (**S13**, 32 mg, 0.16 mmol), DIPEA (83 mg, 0.65 mmol), HATU (59 mg, 0.16 mmol) and dry DMF (3.0 mL) was stirred at room temperature for 1.0 h. The reaction mixture was purified by reverse phase silica gel column chromatography (eluted with $CH_3CN/H_2O$ = 5/95 ~ 95/5 with 0.1% TFA) to give 3-(3-fluorophenyl)-N-((3S,4R)-4-((3-methylisoxazol-5-yl)methyl)tetrahydrofuran-3-yl)-1H-pyrazole-5-carboxamide and 3-(3-fluorophenyl)-N-((3R,4S)-4-((3-methylisoxazol-5-yl)methyl)tetrahydrofuran-3-yl)-1H-pyrazole-5-carboxamide (**1554-06-cis**, 40 mg, 63% yield for two steps, mixture of cis isomers) TFA salt as a white solid.

LC-MS (ESI, *m/z*): $[M+H]^+$ = 371.1.

$^1$H NMR (400 MHz, CD3OD, cis) δ 7.59 – 7.54 (m, 1H), 7.52 – 7.44 (m, 2H), 7.15 – 7.08 (m, 2H), 6.04 (s, 1H), 4.84 – 4.81 (m, 1H), 4.10 (dd, *J* = 9.2, 6.0 Hz, 1H), 4.02 (dd, *J* = 8.8, 7.6 Hz, 1H), 3.82 (dd, *J* = 9.6, 4.0 Hz, 1H), 3.69 (t, *J* = 8.4 Hz, 1H), 3.00 (dd, *J* = 14.8, 6.0 Hz, 1H), 2.94 – 2.85 (m, 1H), 2.79 (dd, *J* = 14.8, 8.4 Hz, 1H), 2.19 (s, 3H).

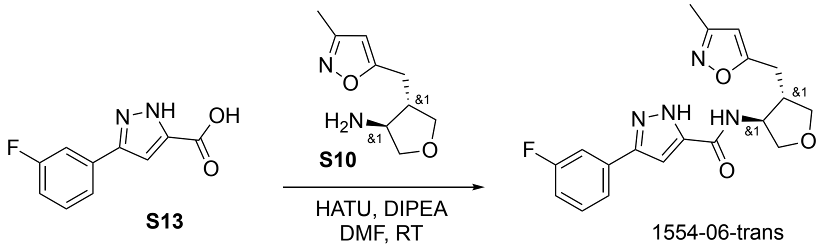


**Procedure for preparation of 3-(3-fluorophenyl)-N-((3R,4R)-4-((3-methylisoxazol-5-yl)methyl)tetrahydrofuran-3-yl)-1H-pyrazole-5-carboxamide and 3-(3-fluorophenyl)-N-((3S,4S)-4-((3-methylisoxazol-5-yl)methyl)tetrahydrofuran-3-yl)-1H-pyrazole-5-carboxamide (trans isomers) (1554-06-trans)**

A mixture of **S10** (41 mg, 0.10 mmol), 3-(3-fluorophenyl)-1H-pyrazole-5-carboxylic acid (**S13**, 25 mg, 0.12 mmol), DIPEA (65 mg, 0.50 mmol), HATU (46 mg, 0.12 mmol) and dry DMF (3.0 mL) was stirred at room temperature for 1.0 h. The reaction mixture was purified by reverse phase silica gel column chromatography (eluted with $CH_3CN/H_2O$ = 5/95 ~ 95/5 with 0.1% TFA) to give 3-(3-fluorophenyl)-N-((3R,4R)-4-((3-methylisoxazol-5-yl)methyl)tetrahydrofuran-3-yl)-1H-pyrazole-5-carboxamide and 3-(3-fluorophenyl)-N-((3S,4S)-4-((3-methylisoxazol-5-yl)methyl)tetrahydrofuran-3-yl)-1H-pyrazole-5-carboxamide (**1554-06-trans**, 23 mg, 48% yield for two steps, mixture of trans isomers) TFA salt as a white solid.

LC-MS (ESI, *m/z*): $[M+H]^+$ = 371.1.

$^{1}$H NMR (400 MHz, $CD_3OD$, trans) δ 7.57 – 7.54 (m, 1H), 7.51 – 7.44 (m, 2H), 7.14 – 7.08 (m, 2H), 6.13 (s, 1H), 4.41 (dd, *J* = 12.4, 5.6 Hz, 1H), 4.14 – 4.08 (m, 2H), 3.70 (dd, *J* = 9.2, 5.6 Hz, 1H), 3.60 (dd, *J* = 9.2, 6.8 Hz, 1H), 3.06 (dd, *J* = 15.2, 6.8 Hz, 1H), 2.91 (dd, *J* = 15.6, 8.4 Hz, 1H), 2.74 – 2.65 (m, 1H), 2.18 (s, 3H).

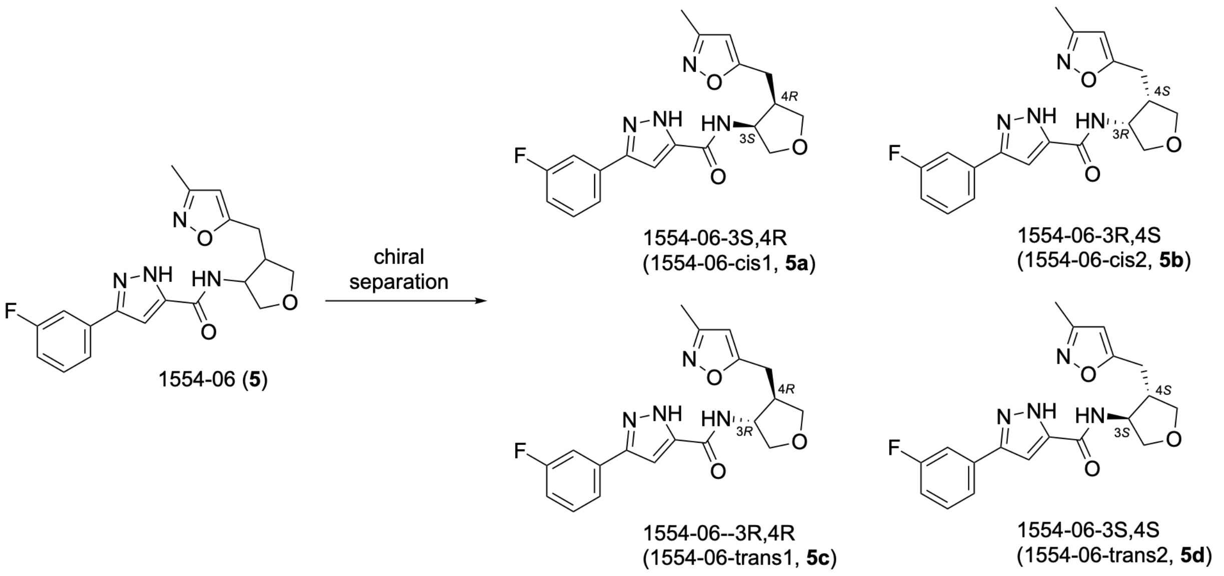


**Procedure for chiral separation of 3-(3-fluorophenyl)-N-(4-((3-methylisoxazol-5-yl)methyl)tetrahydrofuran-3-yl)-1H-pyrazole-5-carboxamide (1554-06)**

3-(3-fluorophenyl)-N-(4-((3-methylisoxazol-5-yl)methyl)tetrahydrofuran-3-yl)-1H-pyrazole-5-carboxamide (**1554-06**, 130 mg, 0.35 mmol) was separated by chiral preparation on an Waters SFC-150mgm with a Daicel AD (25*250mm,10um) column at 30 °C using a CO2/EtOH[0.5%NH3(7M in MeOH)]=45/55 mobile phase, a flow rate of 100 ml/min with a back pressure of 100 bar, detection at 214 nm and a cycle time of 13 minutes. The sample solution was dissolved at 130mg/6.2 ml MeOH, injection volume was 2.5 ml.

**1554-06-cis1 (5a)**: 3-(3-fluorophenyl)-N-((3S,4R)-4-((3-methylisoxazol-5-yl)methyl)tetrahydrofuran-3-yl)-1H-pyrazole-5-carboxamide (39.7 mg). Absolute configuration was confirmed by X-ray single crystal diffraction.

LC-MS (ESI, *m/z*): $[M + H]^+ = 371.1$.

$^{1}$H NMR (400 MHz, Methanol-d4) δ 7.55 (s, 1H), 7.48 (s, 2H), 7.11 (s, 2H), 6.04 (s, 1H), 4.84 – 4.80 (m, 1H), 4.09 (dd, J = 9.4, 6.1 Hz, 1H), 4.02 (dd, J = 8.7, 7.4 Hz, 1H), 3.82 (dd, J = 9.4, 3.8 Hz, 1H), 3.69 (t, J

= 8.4 Hz, 1H), 3.00 (dd, J = 14.9, 5.9 Hz, 1H), 2.95 – 2.84 (m, 1H), 2.79 (dd, J = 14.8, 8.8 Hz, 1H), 2.19 (s, 3H).

**1554-06-cis2 (5b)**: 3-(3-fluorophenyl)-N-((3R,4S)-4-((3-methylisoxazol-5-yl)methyl)tetrahydrofuran-3-yl)-1H-pyrazole-5-carboxamide (33.2 mg).

LC-MS (ESI, *m/z*): [M + H]+ = 371.1.

$^1$H NMR (400 MHz, Methanol-d4) δ 7.56 (d, J = 7.8 Hz, 1H), 7.53 – 7.42 (m, 2H), 7.16 – 7.07 (m, 2H), 6.04 (s, 1H), 4.85 – 4.80 (m, 1H), 4.09 (dd, J = 9.4, 6.1 Hz, 1H), 4.02 (dd, J = 8.7, 7.4 Hz, 1H), 3.82 (dd, J = 9.4, 3.8 Hz, 1H), 3.69 (t, J = 8.4 Hz, 1H), 3.00 (dd, J = 14.9, 5.9 Hz, 1H), 2.95 – 2.84 (m, 1H), 2.79 (dd, J = 14.9, 8.8 Hz, 1H), 2.19 (s, 3H).

**1554-06-trans1 (5c)**: 3-(3-fluorophenyl)-N-((3R,4R)-4-((3-methylisoxazol-5-yl)methyl)tetrahydrofuran-3-yl)-1H-pyrazole-5-carboxamide (7.8 mg).

LC-MS (ESI, *m/z*): [M + H]+ = 371.1.

$^1$H NMR (400 MHz, Methanol-d4) δ 7.55 (dt, J = 7.8, 1.2 Hz, 1H), 7.52 – 7.41 (m, 2H), 7.15 – 7.06 (m, 2H), 6.13 (s, 1H), 4.41 (dt, J = 6.8, 5.5 Hz, 1H), 4.11 (dt, J = 9.1, 7.2 Hz, 2H), 3.70 (dd, J = 9.1, 5.5 Hz, 1H), 3.60 (dd, J = 9.0, 6.6 Hz, 1H), 3.06 (dd, J = 15.4, 6.7 Hz, 1H), 2.91 (dd, J = 15.4, 8.2 Hz, 1H), 2.70 (dqd, J = 8.2, 6.8, 5.5 Hz, 1H), 2.18 (s, 3H).

**1554-06-trans2 (5d)**: 3-(3-fluorophenyl)-N-((3S,4S)-4-((3-methylisoxazol-5-yl)methyl)tetrahydrofuran-3-yl)-1H-pyrazole-5-carboxamide (9.3 mg). Absolute configuration was confirmed by X-ray single crystal diffraction.

LC-MS (ESI, *m/z*): [M + H]+ = 371.1.

$^1$H NMR (400 MHz, Methanol-d4) δ 7.55 (dt, J = 7.7, 1.3 Hz, 1H), 7.53 – 7.41 (m, 2H), 7.15 – 7.06 (m, 2H), 6.13 (s, 1H), 4.41 (dt, J = 6.9, 5.6 Hz, 1H), 4.11 (dt, J = 9.2, 7.2 Hz, 2H), 3.70 (dd, J = 9.1, 5.5 Hz, 1H), 3.60 (dd, J = 9.0, 6.6 Hz, 1H), 3.06 (dd, J = 15.4, 6.8 Hz, 1H), 2.91 (dd, J = 15.4, 8.2 Hz, 1H), 2.76 – 2.63 (m, 1H), 2.18 (s, 3H).

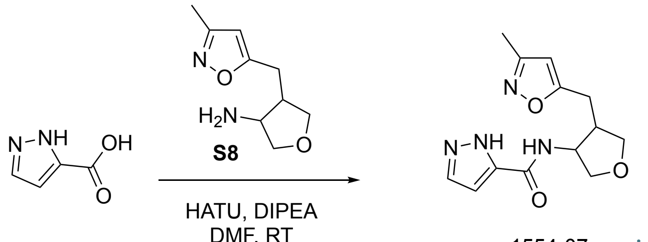


1554-07 javascript:

**Procedure for preparation of N-(4-((3-methylisoxazol-5-yl)methyl)tetrahydrofuran-3-yl)-1H-pyrazole-5-carboxamide (1554-07)**

A mixture of 4-((3-methylisoxazol-5-yl)methyl)tetrahydrofuran-3-amine (**S8**, 140 mg, 0.34 mmol), 1H-pyrazole-5-carboxylic acid (46 mg, 0.41 mmol), HATU (156 mg, 0.41 mmol), DIPEA (221 mg, 1.71 mmol) and dry DMF (3.0 mL) was stirred at room temperature for 1.0 h. The reaction mixture was purified by reverse phase silica gel column chromatography (eluted with $CH_3CN/H_2O$ = 5/95~95/5) to give N-(4-((3-methylisoxazol-5-yl)methyl)tetrahydrofuran-3-yl)-1H-pyrazole-5-carboxamide (**1554-07**, 28 mg, 30% yield, cis/trans=10/7) as a white solid.

LC-MS: MS(ESI, m/z): $[M+H]^+$=277.1

$^1$H NMR (400 MHz, DMSO-$d_6$) δ 13.08 (s, 1H), 8.34 (d, $J$ = 7.4 Hz, 1H), 8.20 (d, $J$ = 8.7 Hz, 1H), 7.76 (dd, $J$ = 3.6, 2.2 Hz, 2H), 6.71 (d, $J$ = 2.3 Hz, 1H), 6.67 (d, $J$ = 2.3 Hz, 1H), 6.17 (s, 1H), 6.11 (s, 1H), 4.71 (p, $J$ = 6.4 Hz, 1H), 4.23 (q, $J$ = 6.4 Hz, 1H), 3.97 (ddd, $J$ = 8.7, 6.9, 3.7 Hz, 2H), 3.87 (dd, $J$ = 8.4, 6.6 Hz, 1H), 3.69 (dd, $J$ = 9.0, 5.0 Hz, 1H), 3.58 – 3.53 (m, 2H), 3.48 – 3.45 (m, 1H), 2.99 – 2.93 (m, 1H), 2.84 – 2.77 (m, 2H), 2.73 – 2.67 (m, 2H), 2.64 (dd, $J$ = 8.4, 6.1 Hz, 1H), 2.15 (s, 2H), 2.14 (s, 3H).

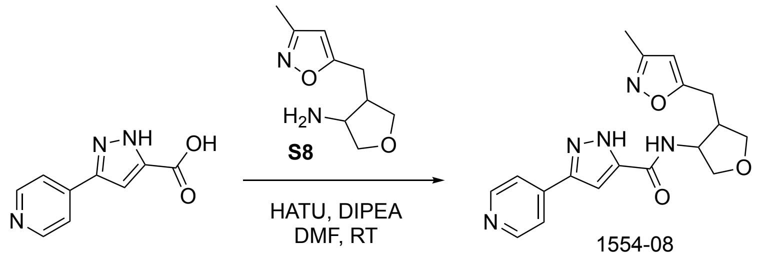


1554-08

**Procedure for preparation of N-(4-((3-methylisoxazol-5-yl)methyl)tetrahydrofuran-3-yl)-3-(pyridin-4-yl)-1H-pyrazole-5-carboxamide (1554-08)**

To a mixture of 3-(4-methoxyphenyl)-1H-pyrazole-5-carboxylic acid (50 mg, 0.26 mmol, 1.0 eq.), 4-((3-methylisoxazol-5-yl)methyl)tetrahydrofuran-3-amine (**S8**, 100 mg, TFA salt, 0.26 mmol, 1.0 eq.) in DMF (2 mL) was added DIPEA (0.5 mL) and HATU (105 mg, 0.28 mmol, 1.05 eq). The resulting mixture was stirred at room temperature for 2 h. The reaction mixture was diluted with water (20 mL), extracted with ethyl acetate (20 mL×3). The combined organic phases were dried over anhydrous $Na_2SO_4$ and concentrated under vacuum. The residue was purified by silica gel column chromatography (eluted with $CH_3CN/H_2O$ = 5/95 ~ 95/5) to give N-(4-((3-methylisoxazol-5-yl)methyl)tetrahydrofuran-3-yl)-3-(pyridin-4-yl)-1H-pyrazole-5-carboxamide (**1554-08**, 20 mg, 21% yield) as a white solid.

LC-MS (ESI, *m/z*): $[M+H]^+$ = 354.2.

$^1$H NMR (400 MHz, Methanol-d4)(cis:trans=4:1) δ 8.79 (d, *J* = 6.4 Hz, 2.5H), 8.31 (s, 2.5H), 7.58 (s, 1H), 7.50 (s, 0.25H), 6.13 (s, 0.25H), 6.05 (s, 1H), 4.42 (q, *J* = 6.0 Hz, 0.25H), 4.17 – 4.07 (m, 1.5H), 4.03 (dd, *J* = 8.8, 7.6 Hz, 1H), 3.82 (dd, *J* = 9.4, 3.8 Hz, 1H), 3.73 – 3.66 (m, 1.25H), 3.60 (dd, *J* = 9.4, 3.8 Hz, 1H), 3.09 – 2.84 (m, 2.5H), 2.84 – 2.66 (m, 1.25H), 2.19 (s, 3H), 2.17 (s, 0.75H).

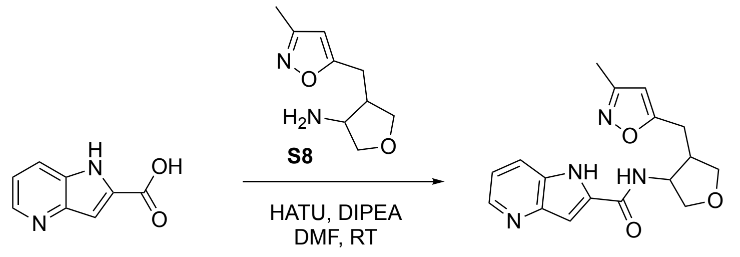


1554-09

**Procedure for preparation of N-(4-((3-methylisoxazol-5-yl)methyl)tetrahydrofuran-3-yl)-1H-pyrrolo[3,2-b]pyridine-2-carboxamide (1554-09)**

To a mixture of 1H-pyrrolo[3,2-b]pyridine-2-carboxylic acid (50 mg, 0.25 mmol, 1.0 eq.), 4-((3-methylisoxazol-5-yl)methyl)tetrahydrofuran-3-amine (**S8**, 100 mg, TFA salt, 0.25 mmol, 1.0 eq.) in DMF (2 mL) was added DIPEA (0.5 mL) and HATU (98 mg, 0.26 mmol, 1.05 eq), the resulting mixture was stirred at room temperature for 2 h. The reaction mixture was diluted with water (20 mL), extracted with ethyl acetate (20 mL×3). The combined organic phases were dried over anhydrous $Na_2SO_4$ and concentrated under vacuum. The residue was purified by silica gel column chromatography (eluted with $CH_3CN/H_2O$ = 5/95 ~ 95/5) to give N-(4-((3-methylisoxazol-5-yl)methyl)tetrahydrofuran-3-yl)-1H-pyrrolo[3,2-b]pyridine-2-carboxamide (**1554-09**, 21.5 mg, 27% yield) as a white solid.

LC-MS (ESI, *m/z*): $[M+H]^+$ = 327.2.

$^1$H NMR (400 MHz, Methanol-d4)(cis:trans=5:3) δ 8.40 – 8.38 (m, 2H), 7.98 – 7.90 (m, 2H), 7.34 (d, *J* = 0.8 Hz, 1H), 7.32 – 7.22 (m, 2.2H), 6.12 (s, 0.6H), 6.03 (s, 1H), 4.91 – 4.89 (m, 1H), 4.47 – 4.43 (m, 0.6H), 4.15 – 4.10(m, 2H), 4.03 (dd, *J* = 8.8, 7.2 Hz, 1H), 3.85 (dd, *J* = 9.4, 4.2 Hz, 1H), 3.76 – 3.67 (m, 2H), 3.61 (dd, *J* = 9.0, 6.7 Hz, 0.6H), 3.11 – 2.88 (m, 3H), 2.82 – 2.69 (m, 1.8H), 2.15 (s, 4.8H).

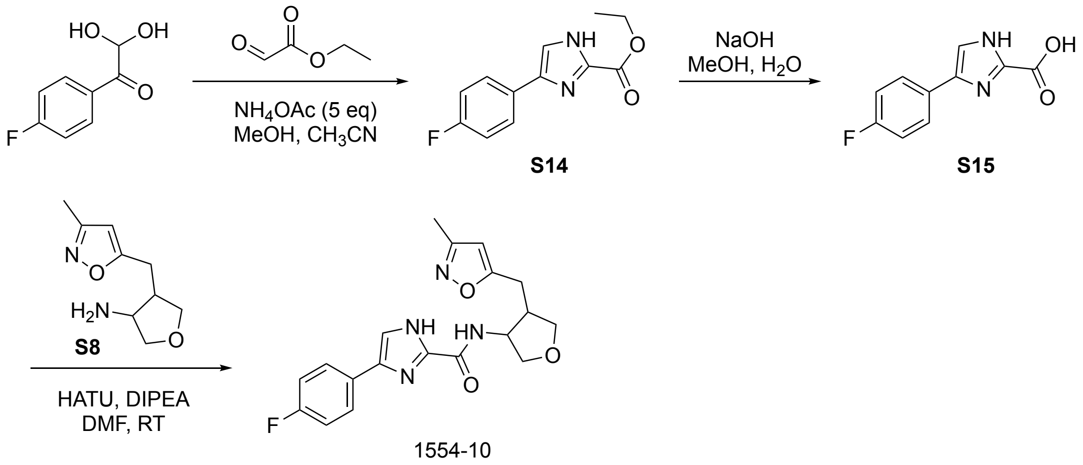

#### Procedure for preparation of ethyl 4-(4-fluorophenyl)-1H-imidazole-2-carboxylate (S14)

To a mixture of 1-(4-fluorophenyl)-2,2-dihydroxyethanone (500 mg, 2.94 mmol, 1.0 eq.) and ethyl 2-oxoacetate (900 mg, 8.82 mmol, 3.0 eq.) in MeOH (20 mL) was added $NH_4OAc$ (1.13 g, 14.70 mmol, 5.0 eq). The resulting mixture was stirred at room temperature for 16 h. The reaction mixture was concentrated under vacuum. The residue was purified by flash silica gel column chromatography (eluted with PE/ethyl acetate = 2/1) to give ethyl 4-(4-fluorophenyl)-1H-imidazole-2-carboxylate (**S14**, 330 mg, 48% yield) as a yellow solid.

LC-MS (ESI, *m/z*): $[M+H]^+$ = 235.1

#### Procedure for preparation of 4-(4-fluorophenyl)-1H-imidazole-2-carboxylic acid (S15)

To a mixture of ethyl 4-(4-fluorophenyl)-1H-imidazole-2-carboxylate (**S14**, 100 mg, 0.45 mmol, 1.0 eq.) in THF/MeOH/$H_2O$ (2 mL/0.5 mL/0.5 mL) was added NaOH (91 mg, 2.27 mmol, 5.0 eq). The resulting mixture was stirred at 50°C for 5 h. The reaction mixture was acidified to pH 3 with HCl (aqueous, 4M), then concentrated under vacuum. The residue was purified by reverse phase silica gel column chromatography (eluted with MeCN/$H_2O$ = 5/95~95/5) to give 4-(4-fluorophenyl)-1H-imidazole-2-carboxylic acid (**S15**, 70 mg, 75% yield) as a white solid.

LC-MS (ESI, *m/z*): $[M+H]^+$ = 207.1

#### Procedure for preparation of 4-(4-fluorophenyl)-N-(4-((3-methylisoxazol-5-yl)methyl)tetrahydrofuran-3-yl)-1H-imidazole-2-carboxamide (1554-10)

To a mixture of 4-(4-fluorophenyl)-1H-imidazole-2-carboxylic acid (**S15**, 40 mg, 0.19 mmol, 1.0 eq.), 4-((3-methylisoxazol-5-yl)methyl)tetrahydrofuran-3-amine (**S8**, 80 mg, TFA salt, 0.19 mmol, 1.0 eq.) in DMF (2 mL) was added DIPEA (0.5 mL) and HATU (77 mg, 0.20 mmol, 1.05 eq). The resulting mixture was stirred at room temperature for 2 h. The reaction mixture was diluted with water (20 mL), extracted with ethyl acetate (20 mL×3). The combined organic phases were dried over anhydrous $Na_2SO_4$ and concentrated under vacuum. The residue was purified by silica gel column chromatography (eluted with $CH_3CN/H_2O$ = 5/95 ~ 95/5) to give 4-(4-fluorophenyl)-N-(4-((3-methylisoxazol-5-yl)methyl)tetrahydrofuran-3-yl)-1H-imidazole-2-carboxamide (**1554-10**, 21 mg, 29% yield) as a white solid.

LC-MS (ESI, *m/z*): $[M+H]^+$ = 371.2.

$^1$H NMR (400 MHz, Methanol-d4)(cis:trans=4:3) δ 7.88 – 7.66 (m, 3.6H), 7.61 – 7.56 (m, 1.5H), 7.24 – 7.05 (m, 3.6H), 6.14 (s, 0.75H), 6.04 (s, 1H), 4.81 – 4.76 (m, 1H), 4.39 (q, *J* = 5.8 Hz, 1H), 4.16 – 3.98 (m, 3.75H), 3.84 (dd, *J* = 9.4, 3.4 Hz, 1H), 3.76 – 3.67 (m, 1.8H), 3.61 (dd, *J* = 9.0, 6.4 Hz, 0.8H), 3.16 – 2.97 (m, 2.4H), 2.97 – 2.77 (m, 3.4H), 2.77 – 2.63 (m, 1H), 2.19 (s, 5.5H).

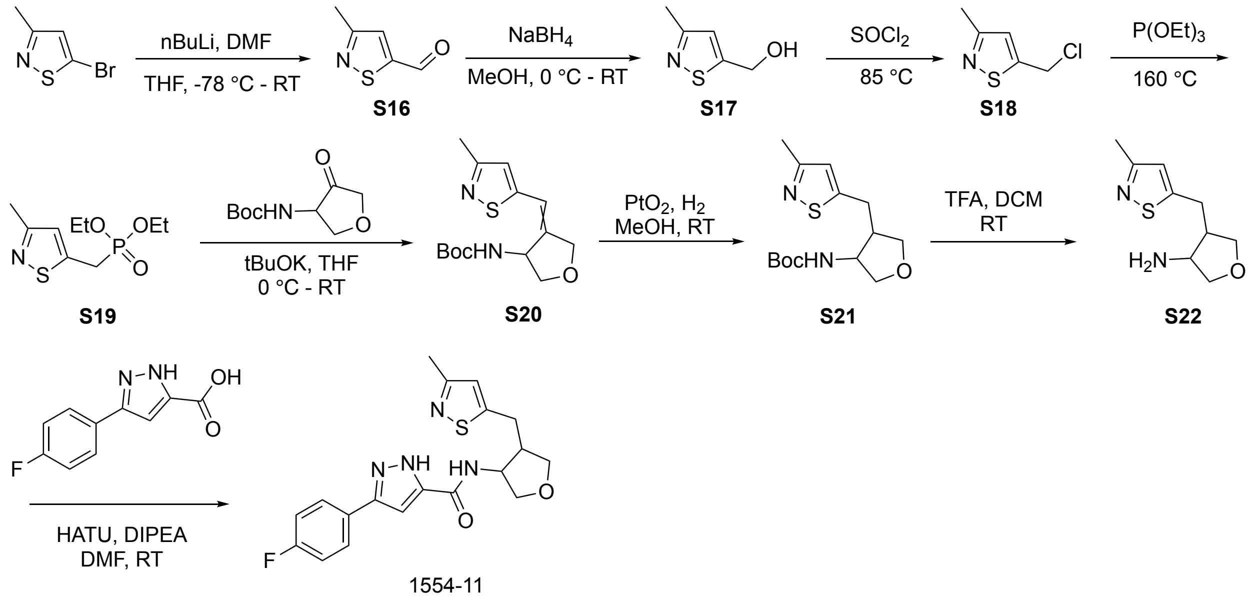


**Procedure for preparation of 3-methylisothiazole-5-carbaldehyde (S16)**

To a solution of 5-bromo-3-methylisothiazole (500 mg, 2.81 mmol) in dry THF (10 mL) was added n-BuLi (1.35 mL, 3.37 mmol, 2.5 M in hexane) dropwise at -78 °C under Ar atmosphere. After 30 minutes, the mixture was cooled to -78 °C and DMF (0.65 mL, 8.42 mmol) was added dropwise under Ar atmosphere. The resulting mixture was stirred at room temperature for 3.0 h. The reaction mixture was quenched by 2 N aqueous HCl (3 mL) at 0 °C. The mixture was stirred at room temperature for 30 minutes and then extracted with ethyl acetate (40 mLx3). The combined organic layers were washed with brine, dried over anhydrous $Na_2SO_4$ and concentrated in vacuo. The residue was purified by silica gel column chromatography (eluted with PE/ethyl acetate = 9/1) to give 3-methylisothiazole-5-carbaldehyde (**S16**, 462 mg crude) as a colorless oil.

$^1$H NMR (400 MHz, $CDCl_3$) δ 10.08 (s, 1H), 7.54 (s, 1H), 2.59 (s, 3H).

**Procedure for preparation of (3-methylisothiazol-5-yl)methanol (S17)**

To a solution of crude 3-methylisothiazole-5-carbaldehyde (**S16**, 462 mg, 3.63 mmol) in MeOH (5 mL) was added $NaBH_4$ (206 mg, 5.45 mmol) at 0 °C. The resulting mixture was stirred at room temperature for 1.0 h. The reaction mixture was concentrated in vacuo. The residue was purified by silica gel column chromatography (eluted with DCM/MeOH = 15/1) to give (3-methylisothiazol-5-yl)methanol (**S17**, 221 mg, 61% yield for two steps) as a yellow oil.

LC-MS (ESI, *m/z*): $[M+H]^+$ = 130.1.

**Procedure for preparation of 5-(chloromethyl)-3-methylisothiazole (S18)**

A solution of (3-methylisothiazol-5-yl)methanol (**S17**, 221 mg, 1.71 mmol) and $SOCl_2$ (2.5 mL, 34.22 mmol) was stirred at 85 °C for 3.0 h. The reaction mixture was concentrated in vacuo. The residue was basified with saturated aqueous $NaHCO_3$ until pH 8 was achieved, and the mixture was extracted with DCM (30 mLx3). The combined organic layers were washed with brine, dried over anhydrous $Na_2SO_4$, concentrated in vacuo. The residue was purified by silica gel column chromatography (eluted with PE/ethyl acetate = 9/1) to give 5-(chloromethyl)-3-methylisothiazole (**S18**, 204 mg, 81% yield) as a brown oil.

LC-MS (ESI, *m/z*): $[M+H]^+$ = 148.0.

**Procedure for preparation of diethyl ((3-methylisothiazol-5-yl)methyl)phosphonate (S19)**

A solution of 5-(chloromethyl)-3-methylisothiazole (**S18**, 204 mg, 1.38 mmol) and $P(OEt)_3$ (0.95 mL, 5.53 mmol) was stirred at 160 °C for 8.0 h under Ar atmosphere. The reaction mixture was concentrated in vacuo. The residue was purified by silica gel chromatography (eluted with DCM/MeOH = 20/1) to give diethyl ((3-methylisothiazol-5-yl)methyl)phosphonate (**S19**, 316 mg, 92% yield) as a colorless oil.

LC-MS (ESI, *m/z*): $[M+H]^+$ = 250.0.

**Procedure for preparation of tert-butyl (4-((3-methylisothiazol-5-yl)methylene)tetrahydrofuran-3-yl)carbamate (S20)**

To a solution of diethyl ((3-methylisothiazol-5-yl)methyl)phosphonate (**S19**, 266 mg, 1.07 mmol) in THF(3 mL) was added $^t$BuOK (1.0 M in THF, 2.14 mL, 2.14 mmol) at 0 °C under Ar atmosphere. The resulting mixture was stirred at room temperature for 2.0 h. Then, tert-butyl (4-oxotetrahydrofuran-3-yl)carbamate (215 mg, 1.07 mmol) in THF (3 mL) was added at 0 ºC. The resulting mixture was stirred at room temperature for 2.5 h. The reaction mixture was diluted with water (30 mL) and extracted with ethyl acetate (30 mLx3). The combined organic layers were washed with brine, dried over anhydrous $Na_2SO_4$ and concentrated in vacuo. The residue was purified by silica gel column chromatography (eluted with PE/ethyl acetate = 4/1) to give tert-butyl (4-((3-methylisothiazol-5-yl)methylene)tetrahydrofuran-3-yl)carbamate (**S20**, 243 mg, 74% yield) as a colorless oil.

LC-MS (ESI, *m/z*): $[M+H]^+$ = 297.1.

**Procedure for preparation of tert-butyl (4-((3-methylisothiazol-5-yl)methyl)tetrahydrofuran-3-yl)carbamate (S21)**

A mixture of tert-butyl (4-((3-methylisothiazol-5-yl)methylene)tetrahydrofuran-3-yl)carbamate (**S20**, 90 mg, 0.30 mmol) in MeOH (8 mL) and $PtO_2$ (90 mg) was stirred at room temperature overnight under $H_2$ atmosphere. The reaction mixture was filtered on Celite and rinsed with MeOH. The mixture was concentrated in vacuo. The residue was purified by silica gel column chromatography (eluted with PE/ethyl acetate = 1/5) to give tert-butyl (4-((3-methylisothiazol-5-yl)methyl)tetrahydrofuran-3-yl)carbamate (**S21**, 7 mg, 8% yield) as a colorless oil.

LC-MS (ESI, *m/z*): $[M+H]^+$ = 299.1.

**Procedure for preparation of 4-((3-methylisothiazol-5-yl)methyl)tetrahydrofuran-3-amine (S22)**

A mixture of tert-butyl (4-((3-methylisothiazol-5-yl)methyl)tetrahydrofuran-3-yl)carbamate (**S21**, 7.0 mg, 0.02 mmol) in DCM (2 mL) and TFA (0.5 mL) was stirred at room temperature for 1.0 h. The reaction mixture was concentrated in vacuo to give 4-((3-methylisothiazol-5-yl)methyl)tetrahydrofuran-3-amine (**S22**, 15 mg) TFA salt crude as a colorless oil.

LC-MS (ESI, *m/z*): $[M+H]^+$ = 199.1.

**Procedure for preparation of 3-(4-fluorophenyl)-N-(4-((3-methylisothiazol-5-yl)methyl)tetrahydrofuran-3-yl)-1H-pyrazole-5-carboxamide (1554-11)**

A mixture of 4-((3-methylisothiazol-5-yl)methyl)tetrahydrofuran-3-amine 2,2,2-trifluoroacetate (1 eq), 3-(4-fluorophenyl)-1H-pyrazole-5-carboxylic acid (1.4 eq), DIPEA (5 eq), HATU (1.1 eq) and dry DMF (0.1 M) was stirred at room temperature for 2.0 h. The reaction mixture was concentrated in vacuo. The residue was purified by reverse phase silica gel column chromatography (eluted with $CH_3CN/H_2O$ =

5/95~95/5 with 0.1% TFA) to give 3-(4-fluorophenyl)-N-(4-((3-methylisothiazol-5-yl)methyl)tetrahydrofuran-3-yl)-1H-pyrazole-5-carboxamide (**1554-11**) TFA salt as a white solid.

LC-MS (ESI, *m/z*): $[M+H]^+$ = 387.0.

$^1$H NMR (400 MHz, $CD_3OD$) δ 7.78 – 7.73 (m, 3.00H), 7.23 – 7.17 (m, 3.03H), 7.13 – 6.97 (m, 2.01H), 6.93 (s, 1.00H), 4.84 – 4.80 (m, 1.00H), 4.41 (dd, *J* = 12.0, 6.0 Hz, 0.50H), 4.13 – 4.07 (m, 2.02H), 4.00 (t, *J* = 8.4 Hz, 1.03H), 3.82 (dd, *J* = 9.2, 3.6 Hz, 1.03H), 3.73 – 3.66 (m, 1.53H), 3.60 (dd, *J* = 8.8, 6.4 Hz, 0.51H), 3.24 (dd, *J* = 14.8, 6.8 Hz, 0.54H), 3.17 (dd, *J* = 15.2, 6.4 Hz, 1.04H), 3.09 (dd, *J* = 14.8, 8.0 Hz, 0.53H), 2.98 (dd, *J* = 15.2, 8.8 Hz, 1.05H), 2.91 – 2.81 (m, 1.07H), 2.71 – 2.64 (m, 0.50H), 2.37 (s, 4.50H).

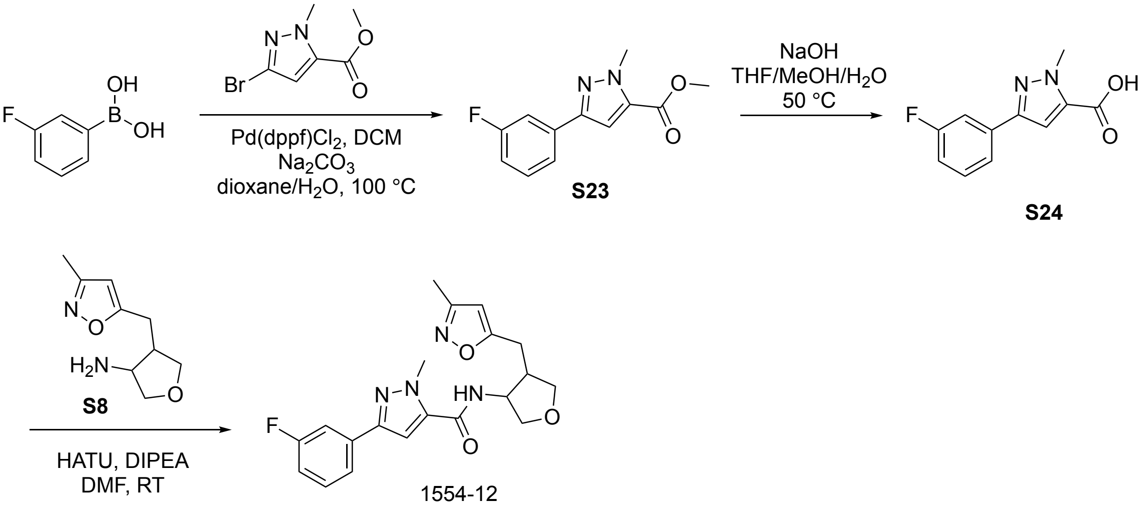


**Procedure for preparation of methyl 3-(3-fluorophenyl)-1-methyl-1H-pyrazole-5-carboxylate (S23)**

A mixture of methyl 3-bromo-1-methyl-1H-pyrazole-5-carboxylate (125 mg, 0.57 mmol) in dioxane/$H_2O$ (3/0.8 mL), (3-fluorophenyl)boronic acid (120 mg, 0.86 mmol), Pd(dppf)$Cl_2$.DCM (23 mg, 0.03 mmol) and $Na_2CO_3$ (121 mg, 1.14 mmol) was stirred at 100 °C overnight under $N_2$ atmosphere. The reaction mixture was adjusted to pH 4.0 with HCl (aqueous, 2 N). The reaction mixture was concentrated in vacuo. The residue was purified by reverse phase silica gel column chromatography (eluted with $CH_3CN/H_2O$ = 5/95~95/5 with 0.1% HCl) to give methyl 3-(3-fluorophenyl)-1-methyl-1H-pyrazole-5-carboxylate (**S23**, 100 mg) as the crude HCl salt as a white solid.

LC-MS (ESI, *m/z*): $[M+H]^+$ = 235.0.

**Procedure for preparation of 3-(3-fluorophenyl)-1-methyl-1H-pyrazole-5-carboxylic acid (S24)**

A mixture of methyl 3-(3-fluorophenyl)-1-methyl-1H-pyrazole-5-carboxylate (**S23**, 100 mg, 0.43 mmol) in MeOH/THF/$H_2O$ (2 mL/2 mL/2 mL) and NaOH (171 mg, 4.27 mmol) was stirred at 50 °C for 1.0 h.

The reaction mixture was adjusted to pH 4.0 with HCl (aqueous, 2 N). The reaction mixture was concentrated in vacuo. The residue was purified by reverse phase silica gel column chromatography (eluted with $CH_3CN/H_2O$ = 5/95~95/5 with 0.1% HCl) to give 3-(3-fluorophenyl)-1-methyl-1H-pyrazole-5-carboxylic acid (**S24**, 93 mg, 64% yield for two steps) HCl salt as a white solid.

LC-MS (ESI, *m/z*): $[M+H]^+$ = 207.0.

**Procedure for preparation of 3-(3-fluorophenyl)-1-methyl-N-(4-((3-methylisoxazol-5-yl)methyl)tetrahydrofuran-3-yl)-1H-pyrazole-5-carboxamide (1554-12)**

A mixture of 4-((3-methylisoxazol-5-yl)methyl)tetrahydrofuran-3-amine (**S8**, 70 mg, 0.17 mmol), 3-(3-fluorophenyl)-1-methyl-1H-pyrazole-5-carboxylic acid (**S24**, 45 mg, 0.20 mmol), DIPEA (110 mg, 0.85 mmol), HATU (78 mg, 0.20 mmol) and dry DMF (2.0 mL) was stirred at room temperature for 1.0 h. The reaction mixture was purified by reverse phase silica gel column chromatography (eluted with $CH_3CN/H_2O$ = 5/95 ~ 95/5 with 0.1% TFA) to give 3-(3-fluorophenyl)-1-methyl-N-(4-((3-methylisoxazol-5-yl)methyl)tetrahydrofuran-3-yl)-1H-pyrazole-5-carboxamide (**1554-12**, 26 mg, 31% yield) TFA salt as a white solid.

LC-MS (ESI, *m/z*): $[M+H]^+$ = 385.1.

$^1$H NMR (400 MHz, $CD_3OD$, cis/trans = 2/1) δ 7.62 – 7.58 (m, 1.50H), 7.55 – 7.49 (m, 1.51H), 7.45 – 7.39 (m, 1.51H), 7.20 (s, 1.00H), 7.10 (s, 0.50H), 7.08 – 7.03 (m, 1.50H), 6.12 (s, 0.50H), 6.06 (s, 1.00H), 4.83 – 4.80 (m, 1.00H), 4.38 (dd, *J* = 12.8, 6.0 Hz, 0.51H), 4.13 (s, 1.50H), 4.12 (s, 3.01H), 4.12 – 4.11 (m, 0.51H), 4.11 – 4.07 (m, 1.50H), 4.02 (dd, *J* = 8.8, 7.6 Hz, 1.00H), 3.80 (dd, *J* = 9.6, 4.0 Hz, 1.00H), 3.70 – 3.65 (m, 1.51H), 3.59 (dd, *J* = 9.2, 6.8 Hz, 0.50H), 3.07 – 3.02 (m, 0.51H), 3.01 – 2.96 (m, 1.00H), 2.95 – 2.86 (m, 1.52H), 2.82 – 2.76 (m, 1.00H), 2.72 – 2.67 (m, 0.51H), 2.19 (s, 3H), 2.18 (s, 1.50H).

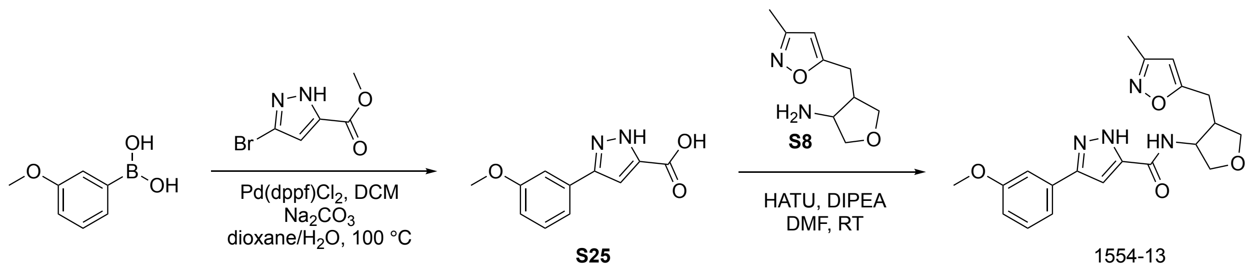


**Procedure for preparation of 3-(3-methoxyphenyl)-1H-pyrazole-5-carboxylic acid (S25)**

To a mixture of methyl 3-bromo-1H-pyrazole-5-carboxylate (200 mg, 0.98 mmol), (3-methoxyphenyl)boronic acid (193 mg, 1.27 mmol) and Pd(dppf)$Cl_2$ (48 mg, 0.06 mmol) was added $Na_2CO_3$ (207 mg, 1.95 mmol) in Dioxane/$H_2O$ (4/1 mL) at room temperature under $N_2$ atmosphere. The resulting mixture was stirred 100°C overnight. The reaction mixture was purified by reverse phase silica gel column chromatography (eluted with $CH_3CN/H_2O$ = 5/95~95/5 with 0.1% HCl) to give 3-(3-methoxyphenyl)-1H-pyrazole-5-carboxylic acid (**S25**, 40 mg, HCl salt, 15% yield) as a white solid.

LC-MS: MS(ESI, m/z): $[M+H]^+$=219.1

**Procedure for preparation of 3-(3-methoxyphenyl)-N-(4-((3-methylisoxazol-5-yl)methyl)tetrahydrofuran-3-yl)-1H-pyrazole-5-carboxamide (1554-13)**

A mixture of 4-((3-methylisoxazol-5-yl)methyl)tetrahydrofuran-3-amine (**S8**, 43 mg, 0.10 mmol), 3-(3-methoxyphenyl)-1H-pyrazole-5-carboxylic acid (**S25**, 25 mg, 0.12 mmol), DIPEA (68 mg, 0.52 mmol), HATU (48 mg, 0.13 mmol) and dry DMF (3.0 mL) was stirred at room temperature for 1.0 h. The reaction mixture was purified by reverse phase silica gel column chromatography (eluted with $CH_3CN/H_2O$ = 5/95 ~ 95/5 with 0.1% TFA) to give 3-(3-methoxyphenyl)-N-(4-((3-methylisoxazol-5-yl)methyl)tetrahydrofuran-3-yl)-1H-pyrazole-5-carboxamide (**1554-13**, 24 mg, 48% yield) TFA salt as a white solid.

LC-MS (ESI, *m/z*): $[M+H]^+$ = 383.1.

$^1$H NMR (400 MHz, $CD_3OD$, cis/trans = 5/2) δ 7.39 – 7.34 (m, 1.40H), 7.31 – 7.28 (m, 2.82H), 7.08 (s, 1.00H), 7.04 (s, 0.40H), 6.96 – 6.93 (m, 1.40H), 6.13 (s, 0.40H), 6.04 (s, 1.00H), 4.84 – 4.82 (m, 1.01H), 4.41 (dd, *J* = 12.4, 6.0 Hz, 0.41H), 4.14 – 4.07 (m, 1.80H), 4.02 (dd, *J* = 8.8, 7.6 Hz, 1.01H), 3.86 (s, 3.01H), 3.85 (s, 1.22H), 3.82 (dd, *J* = 9.2, 4.0 Hz, 1.00H), 3.72 – 3.66 (m, 1.41H), 3.60 (dd, *J* = 8.8, 6.4 Hz, 0.41H), 3.09 – 3.03 (m, 0.41H), 3.00 (dd, *J* = 14.8, 5.6 Hz, 1.03H), 2.91 (dd, *J* = 15.2, 8.0 Hz, 1.41H), 2.79 (dd, *J* = 14.8, 8.8 Hz, 1.01H), 2.72 – 2.67 (m, 0.41H), 2.19 (s, 3.02H), 2.19 (s, 1.20H).

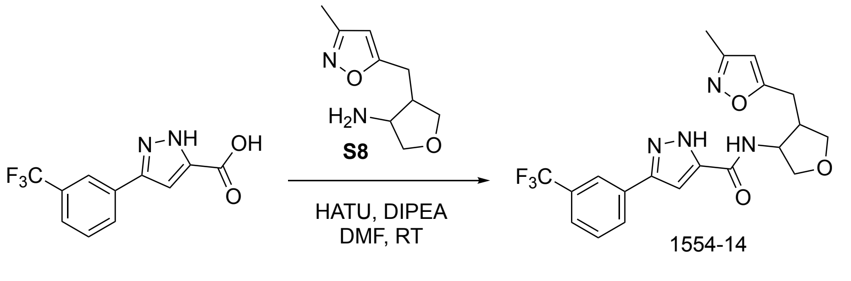


**Procedure for preparation of N-(4-((3-methylisoxazol-5-yl)methyl)tetrahydrofuran-3-yl)-3-(3-(trifluoromethyl)phenyl)-1H-pyrazole-5-carboxamide (1554-14)**

To a mixture of 3-(3-(trifluoromethyl)phenyl)-1H-pyrazole-5-carboxylic acid (50 mg, 0.20 mmol, 1.0 eq.), 4-((3-methylisoxazol-5-yl)methyl)tetrahydrofuran-3-amine (**S8**, 80 mg, TFA salt, 0.20 mmol, 1.0 eq.) in DMF (2 mL) was added DIPEA (0.5 mL) and HATU (84 mg, 0.22 mmol, 1.1 eq), the resulting mixture was stirred at room temperature for 2 h. The reaction mixture was diluted with water (20 mL), extracted with ethyl acetate (20 mL×3). The combined organic phases were dried over anhydrous $Na_2SO_4$ and concentrated under vacuum. The residue was purified by silica gel column chromatography (eluted with $CH_3CN/H_2O$ with 0.1% TFA = 5/95 ~ 95/5) to give N-(4-((3-methylisoxazol-5-yl)methyl)tetrahydrofuran-3-yl)-3-(3-(trifluoromethyl)phenyl)-1H-pyrazole-5-carboxamide (**1554-14**, 35.3 mg, 34% yield) as a white solid.

LC-MS (ESI, *m/z*): $[M+H]^+$ = 421.1.

$^1$H NMR (400 MHz, DMSO-d6) (cis:trans=5:3) δ 13.87 (s, 1H), 8.55 (s, 0.6H), 8.43 (s, 1H), 8.12 – 8.09 (m, 3.2H), 7.73 – 7.68 (m, 3.2H), 7.46 (s, 1H), 7.37 (s, 0.6H), 6.20 (s, 0.6H), 6.13 (s, 1H), 4.78 – 4.72 (m, 1H), 4.33 – 4.22 (m, 0.7H), 4.05 – 3.95 (m, 2.4H), 3.91 (dd, *J* = 8.4, 6.9 Hz, 1.3H), 3.72 (dd, *J* = 9.1, 4.3 Hz, 1H), 3.63 – 3.57 (m, 2.4H), 3.48 (dd, *J* =9.2, 2.4 Hz, 1H), 3.00 (dd, *J* = 15.6, 6.1 Hz, 0.8H), 2.91 – 2.80 (m, 1.7H), 2.80 – 2.68 (m, 2.1H), 2.66 – 2.61 (q, *J* = 7.0 Hz, 0.7H), 2.16 (s, 1.8H), 2.14 (s, 3H).

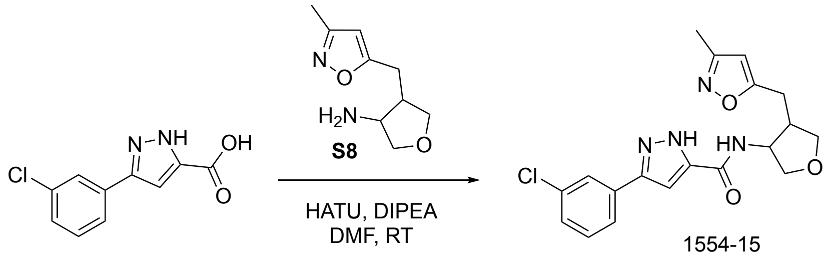


1554-15

**Procedure for preparation of 3-(3-chlorophenyl)-N-(4-((3-methylisoxazol-5-yl)methyl)tetrahydrofuran-3-yl)-1H-pyrazole-5-carboxamide (1554-15)**

To a mixture of 3-(3-chlorophenyl)-1H-pyrazole-5-carboxylic acid (40 mg, 0.18 mmol, 1.0 eq.), 4-((3-methylisoxazol-5-yl)methyl)tetrahydrofuran-3-amine (**S8**, 80 mg, TFA salt, 0.18 mmol, 1.0 eq.) in DMF (2 mL) was added DIPEA (0.5 mL) and HATU (77 mg, 0.20 mmol, 1.1 eq). The resulting mixture was stirred at room temperature for 2 h. The reaction mixture was diluted with water (20 mL), and extracted with ethyl acetate (20 mL×3). The combined organic layers were dried over anhydrous $Na_2SO_4$ and concentrated under vacuum. The residue was purified by reverse phase silica gel column chromatography (eluted with $CH_3CN/H_2O$ = 5/95 ~ 95/5) to give 3-(3-chlorophenyl)-N-(4-((3-methylisoxazol-5-yl)methyl)tetrahydrofuran-3-yl)-1H-pyrazole-5-carboxamide (**1554-15**, 8 mg, 11% yield) as a white solid.

LC-MS (ESI, *m/z*): $[M+H]^+$ = 387.0.

$^1$H NMR (400 MHz, Methanol-d4)(cis:trans=9:4) δ 13.75 (d, *J* = 17.5 Hz, 1H), 8.65 – 8.22 (m, 1H), 7.95 – 7.71 (m, 3H), 7.55 – 7.16 (m, 4.6H), 6.19 (d, *J* = 9.2 Hz, 1H), 6.13 (s, 1H), 4.77 – 4.71 (m, 1H), 4.29 – 4.24 (m, 0.4H), 4.03 – 3.98 (m, 1.8H), 3.94 – 3.90 (m, 1H), 3.77 – 3.67 (m, 1H), 3.60 – 3.56 (m, 1.5H), 3.50 – 3.46 (m, 0.4H), 3.04 - 2.95 (m, 0.7H), 2.89 – 2.66 (m, 4H), 2.15 (s, 1.3H), 2.14 (s, 3H).

javascript:

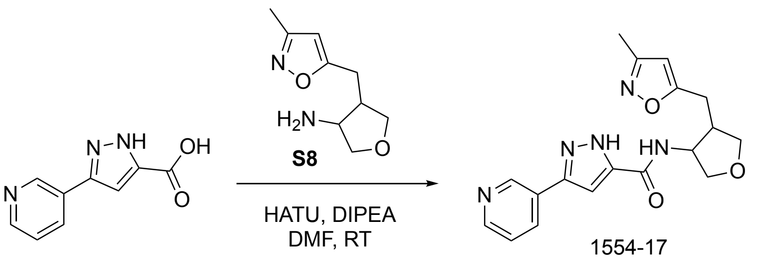


**Procedure for preparation N-(4-((3-methylisoxazol-5-yl)methyl)tetrahydrofuran-3-yl)-3-(pyridin-3-yl)-1H-pyrazole-5-carboxamide (1554-17)**

A mixture of 4-((3-methylisoxazol-5-yl)methyl)tetrahydrofuran-3-amine (**S8**, 80 mg, 0.19 mmol), 3-(pyridin-3-yl)-1H-pyrazole-5-carboxylic acid (44 mg, 0.23 mmol), HATU (104 mg, 0.27 mmol), DIPEA (126 mg, 0.97 mmol) and dry DMF (2.0 mL) was stirred at room temperature for 1.0 h. The reaction mixture was purified by reverse phase silica gel column chromatography (eluted with $CH_3CN/H_2O$ = 5/95~95/5 with 0.1% TFA) to give N-(4-((3-methylisoxazol-5-yl)methyl)tetrahydrofuran-3-yl)-3-(pyridin-3-yl)-1H-pyrazole-5-carboxamide (**1554-17**, 27 mg, TFA salt, 29% yield, cis/trans=10/7) as a white solid.

LC-MS (ESI, m/z): $[M+H]^+$=354.1

HNMR 1554-017: $^1$H NMR (400 MHz, Methanol-*d*$_4$) δ 9.32 – 9.13 (m, 2H), 8.82 (s, 1H), 8.80 (s, 1H), 8.79 – 8.76 (m, 1H), 8.06 – 8.00 (m, 2H), 7.43 (s, 1H), 7.35 (s, 1H), 6.13 (s, 1H), 6.05 (s, 1H), 4.86 – 4.83 (m, 1H), 4.41 (q, *J* = 5.8 Hz, 1H), 4.15 – 4.12 (m, 1H), 4.12 – 4.10 (m, 1H), 4.10 – 4.07 (m, 1H), 4.03 (dd, *J* = 8.7, 7.4 Hz, 1H), 3.83 (dd, *J* = 9.4, 3.8 Hz, 1H), 3.74 – 3.67 (m, 2H), 3.62 – 3.58 (m, 1H), 3.06 (dd, *J* = 15.4, 6.9 Hz, 1H), 3.02 – 2.97 (m, 1H), 2.95 (d, *J* = 8.1 Hz, 1H), 2.91 (d, *J* = 6.1 Hz, 1H), 2.82 – 2.76 (m, 1H), 2.71 (ddd, *J* = 8.0, 6.8, 5.4 Hz, 1H), 2.19 (s, 2H), 2.18 (s, 3H).

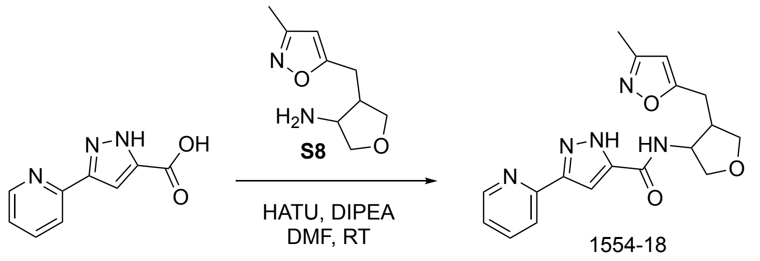

**Procedure for preparation of N-(4-((3-methylisoxazol-5-yl)methyl)tetrahydrofuran-3-yl)-3-(pyridin-2-yl)-1H-pyrazole-5-carboxamide (1554-18)**

A mixture of 4-((3-methylisoxazol-5-yl)methyl)tetrahydrofuran-3-amine (**S8**, 40 mg, 0.10 mmol, Source V4505-108), 3-(pyridin-2-yl)-1H-pyrazole-5-carboxylic acid (22 mg, 0.12 mmol), HATU (52 mg, 0.14 mmol), DIPEA (63 mg, 0.49 mmol) and dry DMF (2.0 mL) was stirred at room temperature for 1.0 h. The reaction mixture was purified by reverse phase silica gel column chromatography (eluted with $CH_3CN/H_2O$ = 5/95~95/5) to give N-(4-((3-methylisoxazol-5-yl)methyl)tetrahydrofuran-3-yl)-3-(pyridin-3-yl)-1H-pyrazole-5-carboxamide (**1554-18**, 22 mg, 62% yield, cis/trans=1/1) as a white solid.

LC-MS: MS(ESI, m/z): $[M+H]^+$=354.1

HNMR 1554-018: $^1$H NMR (400 MHz, Methanol-$d_4$) δ 8.51 (s, 2H), 7.95 (t, *J* = 9.3 Hz, 2H), 7.84 – 7.80 (m, 2H), 7.28 (s, 2H), 7.25 (s, 2H), 6.15 (s, 1H), 6.04 (s, 1H), 4.82 (d, *J* = 3.5 Hz, 1H), 4.42 – 4.38 (m, 1H), 4.14 – 4.12 (m, 1H), 4.10 (d, *J* = 3.9 Hz, 1H), 4.08 (d, *J* = 6.1 Hz, 1H), 4.02 (t, *J* = 8.0 Hz, 1H), 3.82 (dd, *J* = 9.3, 3.4 Hz, 1H), 3.72 (d, *J* = 5.2 Hz, 1H), 3.71 – 3.68 (m, 1H), 3.61 (dd, *J* = 9.0, 6.3 Hz, 1H), 3.12 – 3.06 (m, 1H), 3.06 – 3.01 (m, 1H), 2.93 – 2.89 (m, 1H), 2.86 (d, *J* = 6.7 Hz, 1H), 2.81 (s, 1H), 2.68 (q, *J* = 6.8 Hz, 1H), 2.20 (s, 3H), 2.19 (s, 3H).

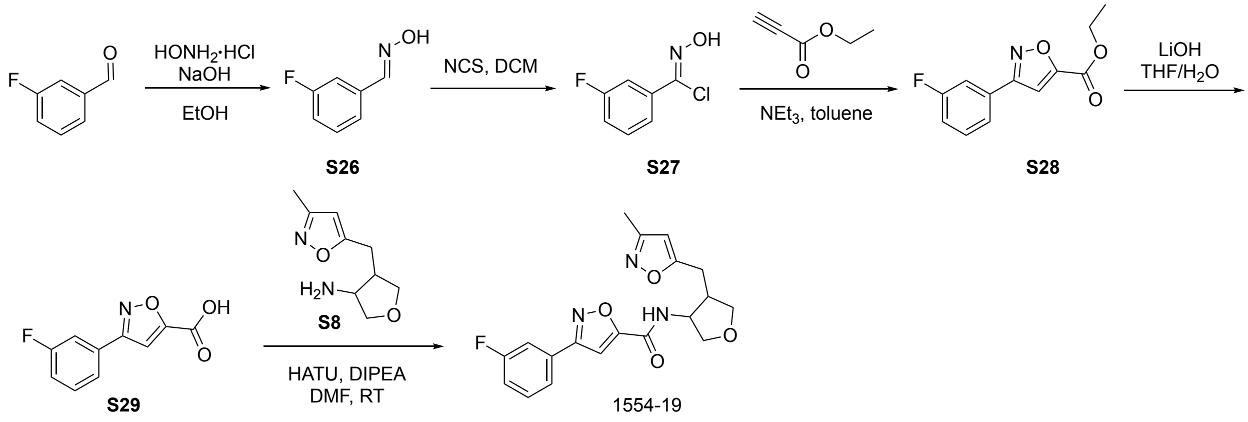


**Procedure for preparation of (E)-3-fluorobenzaldehyde oxime (S26)**

To a solution of 3-fluorobenzaldehyde (5.0 g, 40.3 mmol, 1.0 eq.) and hydroxylamine hydrochloride (3.4 g, 48.3 mmol, 1.2eq.) in EtOH (4 mL)/$H_2O$(13 mL) was added a solution of NaOH (4.03 g, 100.8 mmol, 2.5 eq.) in water (20 mL) drop-wise at 0 °C. The resulting mixture was stirred at room temperature for 30 min. After a white solid precipitated, the mixture was diluted with water and acidified with HCl (aqueous, 4M). The white precipitate was filtered, the filter cake was washed with water, dried in vacuo to give (E)-3-fluorobenzaldehyde oxime (**S26**, 5.1 g, 92% yield) as a white solid.

LC-MS (ESI, *m/z*): $[M+H]^+$ = 140.1.

**Procedure for preparation of (Z)-3-fluoro-N-hydroxybenzimidoyl chloride (S27)**

To a solution of (E)-3-fluorobenzaldehyde oxime (**S26**, 1.0 g, 7.2 mmol, 1.0 eq.) in DMF (20 mL) was added NCS (1.0 g, 7.6 mmol, 1.05 eq.). The resulting mixture was stirred at 35 °C for 3 h. The reaction mixture was concentrated in vacuo. The residue was dissolved in ethyl acetate (100 mL), washed with water (50 mL) and brine (50 mL), dried over anhydrous $Na_2SO_4$ and concentrated under vacuum, followed by purification with flash silica gel column chromatography (eluted with PE/ethyl acetate =3/1) to give (Z)-3-fluoro-N-hydroxybenzimidoyl chloride (**S27**, 1.1 g, 88% yield) as a yellow oil.

LC-MS (ESI, *m/z*): $[M+H]^+$ = 174.1.

**Procedure for preparation of ethyl 3-(3-fluorophenyl)isoxazole-5-carboxylate (S28)**

To a solution of (Z)-3-fluoro-N-hydroxybenzimidoyl chloride (**S27**, 1.1 g, 6.34 mmol, 1.0 eq.) and ethyl propiolate (622 mg, 6.34 mmol, 1.0 eq.) in toluene (20 mL) was added $Et_3N$ (1.28 g, 12.7 mmol, 2.0 eq.). The resulting mixture was stirred at 65 °C for 3 h. The reaction mixture was concentrated in vacuo. The residue was purified via flash column chromatography on silica gel (eluted with PE/ethyl acetate =3/1) to give ethyl 3-(3-fluorophenyl)isoxazole-5-carboxylate (**S28**, 440 mg, 30% yield) as a white solid.

LC-MS (ESI, m/z): $[M+H]^+$ = 236.1.

**Procedure for preparation of 3-(3-fluorophenyl)isoxazole-5-carboxylic acid (S29)**

To a solution of ethyl 3-(3-fluorophenyl)isoxazole-5-carboxylate (**S28**, 200 mg, 0.85 mmol, 1.0 eq.) in THF (5 mL)/$H_2O$ (1 mL) was added LiOH (61 mg, 2.55 mmol, 3.0 eq.). The resulting mixture was stirred at room temperature for 2 h. The reaction mixture was acidified with HCl (aqueous, 4M) to pH 3, then extracted with ethyl acetate (10 mL×3). The combined organic phases were dried over anhydrous $Na_2SO_4$ and concentrated under vacuum. The residue was purified by slurrying from PE, afford 3-(3-fluorophenyl)isoxazole-5-carboxylic acid (**S29**, 130 mg, 74% yield) as a white solid.

LC-MS (ESI, *m/z*): $[M+H]^+$ = 208.1.

**Procedure for preparation of 3-(3-fluorophenyl)-N-(4-((3-methylisoxazol-5-yl)methyl)tetrahydrofuran-3-yl)isoxazole-5-carboxamide (1554-19)**

To a mixture of 3-(3-fluorophenyl)isoxazole-5-carboxylic acid (**S29**, 40 mg, 0.19 mmol, 1.0 eq.) and 4-((3-methylisoxazol-5-yl)methyl)tetrahydrofuran-3-amine (**S8**, 40 mg, TFA salt, 0.19 mmol, 1.0 eq.) in DMF (2 mL) was added DIPEA (0.5 mL) and HATU (77 mg, 0.20 mmol, 1.05 eq). The resulting mixture was stirred at room temperature for 2 h. The reaction mixture was diluted with water (20 mL) and extracted with ethyl acetate (20 mL×3). The combined organic phases were dried over anhydrous $Na_2SO_4$ and concentrated under vacuum. The residue was purified by silica gel column chromatography (eluted with $CH_3CN/H_2O$ = 5/95 ~ 95/5) to give 3-(3-fluorophenyl)-N-(4-((3-methylisoxazol-5-yl)methyl)tetrahydrofuran-3-yl)isoxazole-5-carboxamide (**1554-19**, 14 mg, 20% yield) as a white solid.

LC-MS (ESI, *m/z*): $[M+H]^+$ = 372.0.

$^1$H NMR (400 MHz, Methanol-d4)(cis:trans=7:1) δ 7.74 - 7.70 (m, 1.14H), 7.68 - 7.63 (m, 1.14H), 7.57 - 7.52 (m, 1H), 7.44 (s, 1H), 7.40 (s, 0.14H), 7.29 - 7.23 (m, 1.14H), 6.13 (s, 0.14H), 6.05 (s, 1H), 4.88 – 4.86 (m, 1H),4.44 – 4.40 (m, 0.14H), 4.16 – 4.05 (m, 1.3H), 4.02 (dd, *J* = 8.7, 7.3 Hz, 1H), 3.83 (dd, *J* = 9.4, 4.0 Hz, 1H), 3.69 (t, J = 8.2 Hz, 1.14H), 3.60 (dd, *J* = 8.8, 6.4 Hz, 0.14H) 3.08 - 2.87 (m, 2.4H), 2.83 - 2.70 (m, 1.28H), 2.19 (s, 3.4H).

**Procedure for preparation of N-(4-((3-methylisoxazol-5-yl)methyl)tetrahydrofuran-3-yl)pyrazolo[1,5-a]pyridine-2-carboxamide (1554-21)**

To a mixture of pyrazolo[1,5-a]pyridine-2-carboxylic acid (30 mg, 0.19 mmol, 1.0 eq.) and 4-((3-methylisoxazol-5-yl)methyl)tetrahydrofuran-3-amine (**S8**, 80 mg, TFA salt, 0.19 mmol, 1.0 eq.) in DMF (2 mL) was added DIPEA (0.5 mL) and HATU (77 mg, 0.20 mmol, 1.05 eq). The resulting mixture was stirred at room temperature for 2.0 h. The reaction mixture was diluted with water (20 mL) and extracted with ethyl acetate (20 mL×3). The combined organic phases were dried over anhydrous $Na_2SO_4$ and concentrated under vacuum. The residue was purified by silica gel column chromatography (eluted with $CH_3CN/H_2O$ = 5/95 ~ 95/5) to give N-(4-((3-methylisoxazol-5-yl)methyl)tetrahydrofuran-3-yl)pyrazolo[1,5-a]pyridine-2-carboxamide (**1554-21**, 32 mg, 53% yield) as a white solid.

LC-MS (ESI, *m/z*): $[M+H]^+$ = 327.1.

$^1$H NMR (400 MHz, Methanol-d4)(cis:trans=3:2) δ 8.58 – 8.55 (m, 1.7H), 7.71 – 7.67 (m, 1.7H), 7.28 – 7.22 (m, 1.7H), 7.05 – 6.94 (m, 3.4H), 6.13 (s, 0.7H), 6.03 (s, 1H), 4.91 – 4.88 (m, 1H), 4.48 – 4.44 (m, 0.7H), 4.17 – 4.07 (m, 2.5H), 4.03 (dd, *J* = 8.7, 7.4 Hz, 1H), 3.85 (dd, *J* = 9.3, 3.8 Hz, 1H), 3.77 – 3.66 (m, 1.8H), 3.61 (dd, *J* = 9.0, 6.6 Hz, 0.7H), 3.10 – 2.98 (m, 1.8H), 2.96 – 2.85 (m, 1.8H), 2.85 – 2.68 (m, 1.8H), 2.18 (s, 3H), 2.17 (s, 2.1H).

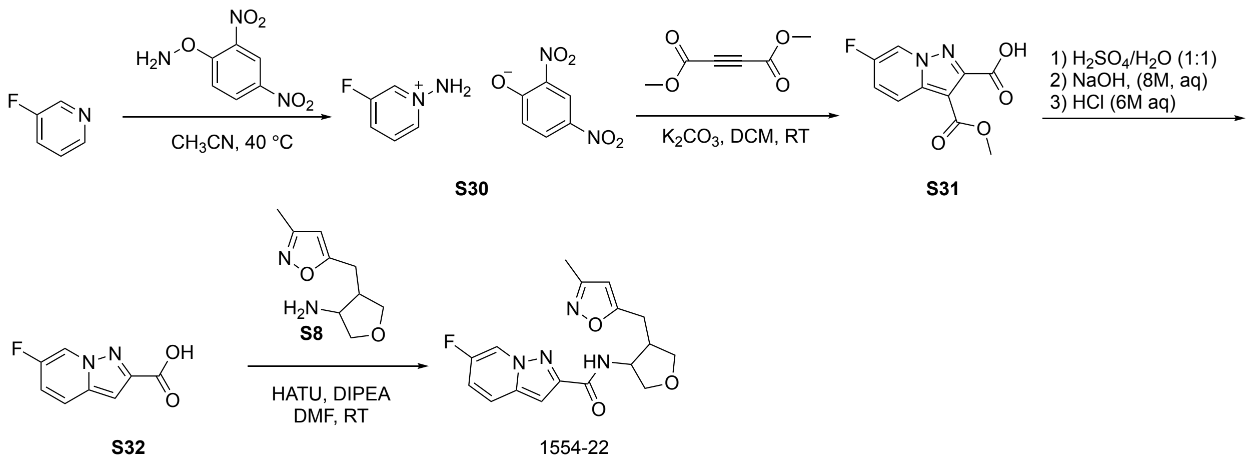


**Procedure for preparation of 1-amino-3-fluoropyridin-1-ium 2,4-dinitrophenolate (S30)**

A mixture of 3-fluoropyridine (1 g, 10.30 mmol, 1.0 eq.) and o-(2,4-dinitrophenyl)hydroxylamine (2.05 g, 10.30 mmol, 1.1 eq.) in $CH_3CN$ (35 mL) was stirred at 40 °C overnight. The reaction mixture was concentrated to give 1-amino-3-fluoropyridin-1-ium 2,4-dinitrophenolate (**S30**, 3.2g, crude) as a red oil.

LC-MS (ESI, *m/z*): $[M+H]^+$ = 113.0

**Procedure for preparation of dimethyl 6-fluoropyrazolo[1,5-a]pyridine-2,3-dicarboxylate (S31)**

To a mixture of 1-amino-3-fluoropyridin-1-ium 2,4-dinitrophenolate (**S30**, crude, 3.2g, 10.30 mmol, 1.0 eq.) and dimethyl but-2-ynedioate (2.2 g, 15.45 mmol, 1.5 eq.) in DMF (50 mL) was added $K_2CO_3$ (2.84 g, 20.60 mmol, 2.0 eq.). The resulting mixture was stirred at room temperature for 6.0 h. The mixture was poured into $H_2O$ (100 mL) and extracted with ethyl acetate (50 mL×3). The combined organic layers were washed with brine, dried over anhydrous $Na_2SO_4$, and concentrated in vacuo. The residue was purified by silica gel column chromatography (eluted with PE ethyl acetate = 5/1~1/1) to afford dimethyl 6-fluoropyrazolo[1,5-a]pyridine-2,3-dicarboxylate (**S31**, 140 mg, 5% yield ) as a light-yellow oil.

LC-MS: m/z 253.1 (M+H)+.

**Procedure for preparation of 6-fluoropyrazolo[1,5-a]pyridine-2-carboxylic acid (S32)**

A solution of dimethyl 6-fluoropyrazolo[1,5-a]pyridine-2,3-dicarboxylate (**S31**, 140 mg, 056 mmol, 1.0 eq.) in conc. $H_2SO_4$ (3 mL)/$H_2O$ (3 mL) was stirred at 100 °C for 3.0 h. After cooling in ice bath, pH 8 was attained via addition of an aqueous solution of NaOH (8 M). This mixture was then acidified to pH 3 with aqueous 6M HCl. The mixture was then extracted with DCM (5 mL×3). The combined organic layers were dried over anhydrous $Na_2SO_4$, concentrated in vacuum, followed by purification by slurrying from PE, give 6-fluoropyrazolo[1,5-a]pyridine-2-carboxylic acid (**S32**, 83 mg, 83% yield) as a white solid.

LC-MS: m/z 181.1 (M+H)+.

**Procedure for preparation of 6-fluoro-N-(4-((3-methylisoxazol-5-yl)methyl)tetrahydrofuran-3-yl)pyrazolo[1,5-a]pyridine-2-carboxamide (1554-22)**

To a mixture of 6-fluoropyrazolo[1,5-a]pyridine-2-carboxylic acid (**S32**, 40 mg, 0.22 mmol, 1.0 eq.) and 4-((3-methylisoxazol-5-yl)methyl)tetrahydrofuran-3-amine (**S8**, 50 mg, TFA salt, 0.22 mmol, 1.0 eq.) in DMF (2 mL) was added DIPEA (0.5 mL) and HATU (92 mg, 0.24 mmol, 1.1 eq). The resulting mixture was stirred at room temperature for 2.0 h. The reaction mixture was diluted with water (20 mL), extracted with ethyl acetate (20 mL×3). The combined organic phases were dried over anhydrous $Na_2SO_4$ and concentrated under vacuum. The residue was purified by silica gel column chromatography (eluted with $CH_3CN/H_2O$ = 5/95 ~ 95/5) to give 6-fluoro-N-(4-((3-methylisoxazol-5-yl)methyl)tetrahydrofuran-3-yl)pyrazolo[1,5-a]pyridine-2-carboxamide (**1554-22**, 15.8 mg, 21% yield) as a white solid.

LC-MS (ESI, *m/z*): $[M+H]^+$ = 345.0.

$^1$H NMR (400 MHz, Methanol-d4)(cis:trans=5:4) δ 8.69 (dtd, J = 11.6, 2.5, 1.3 Hz, 2H), 7.75 (dt, J = 10.4, 5.3 Hz, 2H), 7.28 (dddd, J = 10.1, 8.1, 4.5, 2.1 Hz, 2H), 7.08 (d, J = 10.5 Hz, 2H), 6.13 (s, 1H), 6.04 (s, 1H), 4.90 (d, J = 6.4 Hz, 1H), 4.46 (dt, J = 7.0, 5.6 Hz, 1H), 4.17 – 4.07 (m, 3H), 4.03 (dd, J = 8.7, 7.4 Hz, 1H), 3.85 (dd, J = 9.3, 3.8 Hz, 1H), 3.77 – 3.65 (m, 2H), 3.61 (dd, J = 9.0, 6.5 Hz, 1H), 3.12 – 2.84 (m, 4H), 2.84 – 2.66 (m, 2H), 2.18 (d, J = 3.4 Hz, 6H).

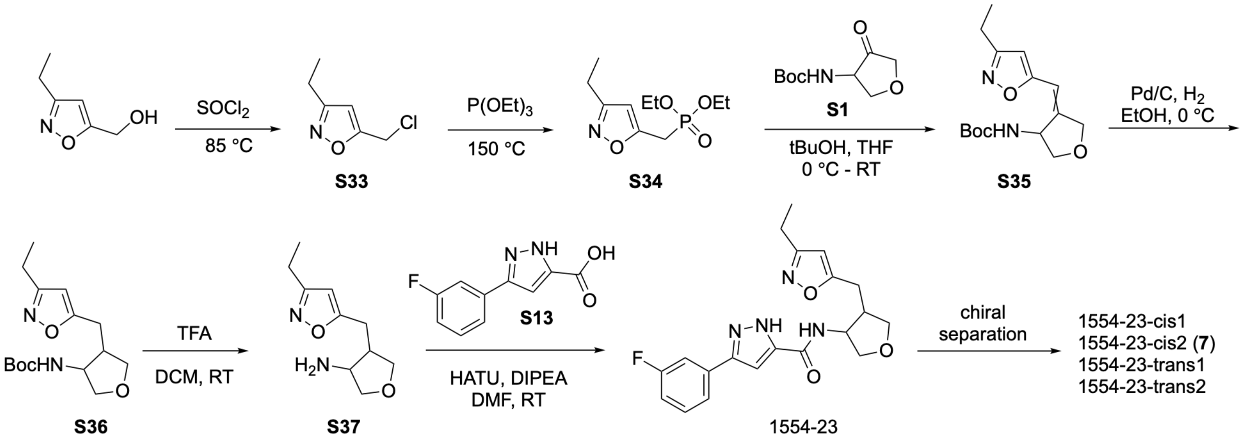


**Procedure for preparation of 5-(chloromethyl)-3-ethylisoxazole (S33)**

A solution of (3-ethylisoxazol-5-yl)methanol (400 mg, 3.15 mmol) and $SOCl_2$ (2.29 mL, 31.46 mmol) was stirred at 85 ºC for 1.0 h. The reaction mixture was concentrated in vacuo. The residue was basified to pH 8 with saturated aqueous $NaHCO_3$ and extracted with DCM (30 mLx3). The combined organic layers were washed with brine, dried over anhydrous $Na_2SO_4$, concentrated in vacuo. The residue was purified by silica gel column chromatography (eluted with PE/ethyl acetate = 9/1) to give 5-(chloromethyl)-3-ethylisoxazole (**S33**, 320 mg, 70% yield) as a brown oil.

LC-MS (ESI, *m/z*): $[M+H]^+$ = 146.0.

**Procedure for preparation of diethyl ((3-ethylisoxazol-5-yl)methyl)phosphonate (S34)**

A solution of **S33** (393 mg, 2.70 mmol) and $P(OEt)_3$ (1.85 mL, 10.80 mmol) was stirred at 150 °C for 7.0 h under $N_2$ atmosphere. The reaction mixture was concentrated in vacuo. The residue was purified by silica gel chromatography (eluted with DCM/MeOH = 20/1) to give diethyl ((3-ethylisoxazol-5-yl)methyl)phosphonate (**S34**, 640 mg, 96% yield) as a colorless oil.

LC-MS (ESI, *m/z*): $[M+H]^+$ = 248.1.

**Procedure for preparation of tert-butyl (4-((3-ethylisoxazol-5-yl)methylene)tetrahydrofuran-3-yl)carbamate (S35)**

To a solution of diethyl ((3-ethylisoxazol-5-yl)methyl)phosphonate (**S34**, 640 mg, 2.59 mmol) in THF (4 mL) was added $^t$BuOK (1.0 M in THF, 5.18 mL, 5.18 mmol) at 0 °C under $N_2$ atmosphere. The resulting mixture was stirred at room temperature for 1.0 h. Then, tert-butyl (4-oxotetrahydrofuran-3-yl)carbamate (**S1**, 521 mg, 2.59 mmol) in THF (6 mL) was added at 0 °C. The resulting mixture was stirred at room temperature for 3.0 h. The reaction mixture was diluted with water (30 mL) and extracted with ethyl acetate (30 mLx3). The combined organic layers were washed with brine, dried over anhydrous $Na_2SO_4$ and concentrated in vacuo. The residue was purified by silica gel column chromatography (eluted with PE/ethyl acetate = 4/1) to give tert-butyl (4-((3-ethylisoxazol-5-yl)methylene)tetrahydrofuran-3-yl)carbamate (**S35**, 471 mg, 62% yield) as a yellow oil.

LC-MS (ESI, *m/z*): $[M+H-C_4H_8]^+$ = 239.1.

**Procedure for preparation of tert-butyl (4-((3-ethylisoxazol-5-yl)methyl)tetrahydrofuran-3-yl)carbamate (S36)**

To a mixture of tert-butyl (4-((3-ethylisoxazol-5-yl)methylene)tetrahydrofuran-3-yl)carbamate (**S35**, 223 mg, 0.76 mmol) in EtOH (15 mL) was added Pd/C (111 mg), and the reaction was placed under $H_2$ atmosphere and stirred at room temperature. The reaction mixture was filtered on Celite and rinsed with MeOH. The solvent was concentrated in vacuo. The residue was purified by silica gel column chromatography (eluted with PE/ethyl acetate = 4/1) to give tert-butyl (4-((3-ethylisoxazol-5-yl)methyl)tetrahydrofuran-3-yl)carbamate (**S36**, 58 mg, 26% yield) as a colorless oil.

LC-MS (ESI, *m/z*): $[M+H-C_4H_8]^+$ = 241.1.

**Procedure for preparation of 4-((3-ethylisoxazol-5-yl)methyl)tetrahydrofuran-3-amine (S37)**

A mixture of tert-butyl (4-((3-ethylisoxazol-5-yl)methyl)tetrahydrofuran-3-yl)carbamate (**S36**, 58 mg, 0.20 mmol) in DCM (2 mL) and TFA (0.5 mL) was stirred at room temperature for 1.0 h. The reaction mixture was concentrated in vacuo to give 4-((3-ethylisoxazol-5-yl)methyl)tetrahydrofuran-3-amine (**S37**, 86 mg), crude TFA salt as a colorless oil.

LC-MS (ESI, *m/z*): $[M+H]^+$ = 197.1.

**Procedure for preparation of N-(4-((3-ethylisoxazol-5-yl)methyl)tetrahydrofuran-3-yl)-3-(3-fluorophenyl)-1H-pyrazole-5-carboxamide (1554-23, 7)**

A mixture of 4-((3-ethylisoxazol-5-yl)methyl)tetrahydrofuran-3-amine (**S37**, 59 mg, 0.14 mmol), 3-(3-fluorophenyl)-1H-pyrazole-5-carboxylic acid (**S13**, 34 mg, 0.17 mmol), DIPEA (90 mg, 0.70 mmol), HATU (63 mg, 0.17 mmol) and dry DMF (2.5 mL) was stirred at room temperature for 1.0 h. The reaction mixture was purified by reverse phase silica gel column chromatography (eluted with $CH_3CN/H_2O$ = 5/95 ~ 95/5 with 0.1% TFA) to give N-(4-((3-ethylisoxazol-5-yl)methyl)tetrahydrofuran-3-yl)-3-(3-fluorophenyl)-1H-pyrazole-5-carboxamide (**1554-23**, 29 mg, 43% yield for two steps) TFA salt as a white solid.

LC-MS (ESI, *m/z*): $[M+H]^+$ = 385.1.

$^1$H NMR (400 MHz, $CD_3OD$, cis/trans = 5/4) δ 7.57 – 7.54 (m, 1.80H), 7.52 – 7.44 (m, 3.61H), 7.15 – 7.06 (m, 3.60H), 6.17 (s, 0.80H), 6.08 (s, 1.00H), 4.83 – 4.78 (m, 1.00H), 4.42 (dd, *J* = 12.4, 6.0 Hz, 0.81H), 4.15 – 4.08 (m, 2.65H), 4.03 (dd, *J* = 8.4, 7.6 Hz, 1.01H), 3.82 (dd, *J* = 9.2, 4.0 Hz, 1.01H), 3.72 – 3.67 (m, 1.82H), 3.60 (dd, *J* = 8.8, 6.8 Hz, 0.81H), 3.09 – 2.98 (m, 1.80H), 2.96 – 2.87 (m, 1.87H), 2.80 (dd, *J* = 14.8, 8.8 Hz, 1.03H), 2.71 (dd, *J* = 13.6, 7.2 Hz, 0.82H), 2.62 – 2.55 (m, 3.61H), 1.18 (q, *J* = 7.6 Hz, 5.40H).

**Procedure for chiral separation of N-(4-((3-ethylisoxazol-5-yl)methyl)tetrahydrofuran-3-yl)-3-(3-fluorophenyl)-1H-pyrazole-5-carboxamide (1554-23)**

A sample of 186 mg of N-(4-((3-ethylisoxazol-5-yl)methyl)tetrahydrofuran-3-yl)-3-(3-fluorophenyl)-1H-pyrazole-5-carboxamide (1554-23) was purified by chiral separation (see 1554-06 for conditions, eluted with $CO_2$/EtOH[1%$NH_3$(7M in MeOH)]=80/20) to give N-(4-((3-ethylisoxazol-5-yl)methyl)tetrahydrofuran-3-yl)-3-(3-fluorophenyl)-1H-pyrazole-5-carboxamide (28 mg, 15% yield, (3R,4S *or* 3S,4R, **1554-23-cis1**) as a white solid, N-(4-((3-ethylisoxazol-5-yl)methyl)tetrahydrofuran-3-

yl)-3-(3-fluorophenyl)-1H-pyrazole-5-carboxamide (37 mg, 19% yield, 3R,4S *or* 3S,4R, **1554-23-cis2, 10**) as a white solid, N-(4-((3-ethylisoxazol-5-yl)methyl)tetrahydrofuran-3-yl)-3-(3-fluorophenyl)-1H-pyrazole-5-carboxamide (18 mg, 9% yield, 3S,4S *or* 3R,4R, **1554-23-trans1**) as a white solid, N-((3R,4R)-4-((3-ethylisoxazol-5-yl)methyl)tetrahydrofuran-3-yl)-3-(3-fluorophenyl)-1H-pyrazole-5-carboxamide (11 mg, 6% yield, 3S,4S *or* 3R,4R, **1554-23-trans2**) as a white solid.

LC-MS: MS (ESI, *m/z*): $[M+H]^+$=385.1

**1554-23-cis1**: $^1$H NMR (400 MHz, Methanol-d4) δ 7.56 (d, J = 7.7 Hz, 1H), 7.48 (t, J = 9.8 Hz, 2H), 7.11 (t, J = 8.3 Hz, 2H), 6.08 (s, 1H), 4.85 – 4.82 (m, 1H), 4.10 (dd, J = 9.4, 6.1 Hz, 1H), 4.03 (dd, J = 8.7, 7.4 Hz, 1H), 3.82 (dd, J = 9.4, 3.9 Hz, 1H), 3.69 (t, J = 8.3 Hz, 1H), 3.04 – 2.98 (m, 1H), 2.94 – 2.85 (m, 1H), 2.80 (dd, J = 14.9, 8.7 Hz, 1H), 2.59 (q, J = 7.6 Hz, 2H), 1.19 (t, J = 7.6 Hz, 3H).

**1554-023-cis2 (4)**: $^1$H NMR (400 MHz, Methanol-d4) δ 7.56 (d, J = 7.8 Hz, 1H), 7.52 – 7.44 (m, 2H), 7.17 – 7.06 (m, 2H), 6.08 (s, 1H), 4.84 – 4.82 (m, 1H), 4.10 (dd, J = 9.4, 6.1 Hz, 1H), 4.03 (dd, J = 8.7, 7.4 Hz, 1H), 3.82 (dd, J = 9.3, 3.9 Hz, 1H), 3.69 (t, J = 8.3 Hz, 1H), 3.01 (dd, J = 15.0, 6.0 Hz, 1H), 2.94 – 2.86 (m, 1H), 2.80 (dd, J = 14.9, 8.7 Hz, 1H), 2.59 (q, J = 7.6 Hz, 2H), 1.19 (t, J = 7.6 Hz, 3H).

**1554-023-trans1**: $^1$H NMR (400 MHz, Methanol-d4) δ 7.59 – 7.53 (m, 1H), 7.52 – 7.43 (m, 2H), 7.12 (dd, J = 8.7, 2.6 Hz, 1H), 7.09 (s, 1H), 6.17 (s, 1H), 4.42 (q, J = 6.1 Hz, 1H), 4.15 – 4.07 (m, 2H), 3.69 (dd, J = 9.1, 5.6 Hz, 1H), 3.60 (dd, J = 9.0, 6.7 Hz, 1H), 3.06 (dd, J = 15.3, 7.0 Hz, 1H), 2.93 (dd, J = 15.3, 7.9 Hz, 1H), 2.71 (q, J = 6.9 Hz, 1H), 2.58 (q, J = 7.6 Hz, 2H), 1.17 (t, J = 7.6 Hz, 3H).

**1554-023-trans2**: $^1$H NMR (401 MHz, Methanol-d4) δ 7.59 – 7.52 (m, 1H), 7.48 (d, J = 9.1 Hz, 2H), 7.08 (d, J = 11.0 Hz, 2H), 6.17 (s, 1H), 4.42 (q, J = 6.1 Hz, 1H), 4.11 (ddd, J = 10.2, 9.1, 7.1 Hz, 2H), 3.69 (dd, J = 9.1, 5.7 Hz, 1H), 3.60 (dd, J = 8.9, 6.7 Hz, 1H), 3.06 (dd, J = 15.4, 7.0 Hz, 1H), 2.93 (dd, J = 15.4, 8.0 Hz, 1H), 2.74 – 2.68 (m, 1H), 2.58 (q, J = 7.6 Hz, 2H), 1.17 (t, J = 7.6 Hz, 3H).

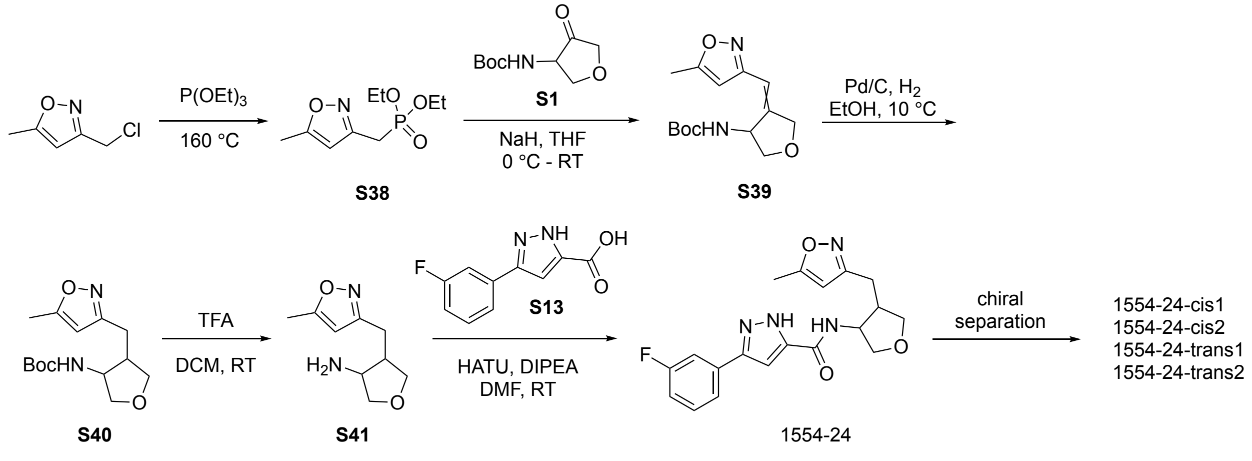

**Procedure for preparation of diethyl ((5-methylisoxazol-3-yl)methyl)phosphonate (S38)**

A mixture of 3-(chloromethyl)-5-methylisoxazole (1 g, 7.60 mmol) in $P(OEt)_3$ (10 mL) was stirred at 160 ºC overnight. The reaction mixture was concentrated in vacuo. The residue was purified by silica gel column chromatography (eluted with DCM/MeOH = 10/1) to give diethyl ((5-methylisoxazol-3-yl)methyl)phosphonate (**S38**, 711 mg, 40% yield) as a yellow oil.

LC-MS: MS (ESI, *m/z*): $[M+H]^+$=234.1.

**Procedure for preparation of tert-butyl (4-((5-methylisoxazol-3-yl)methylene)tetrahydrofuran-3-yl)carbamate (S39)**

To a solution of diethyl ((5-methylisoxazol-3-yl)methyl)phosphonate (**S38**, 711 mg, 3.05 mmol) in THF (10 mL) was added NaH (155 mg, 3.88 mmol) at 0 ºC under Ar atmosphere. The resulting mixture was stirred at room temperature for 1.0 h. Then, tert-butyl (4-oxotetrahydrofuran-3-yl)carbamate (**S1**, 558 mg, 2.77 mmol) was added at 0 ºC. The resulting mixture was stirred at room temperature for 3.0 h. The reaction mixture was diluted with water (30 mL) and extracted with ethyl acetate (30 mLx3). The combined organic layers were washed with brine, dried over anhydrous $Na_2SO_4$ and concentrated in vacuo. The residue was purified by silica gel column chromatography (eluted with DCM/MeOH = 8/1) to give tert-butyl (4-((5-methylisoxazol-3-yl)methylene)tetrahydrofuran-3-yl)carbamate (**S39**, 264 mg, 30% yield) as a colorless oil.

LC-MS: MS (ESI, *m/z*): $[M+H]^+$=281.1.

**Procedure for preparation of tert-butyl (4-((5-methylisoxazol-3-yl)methyl)tetrahydrofuran-3-yl)carbamate (S40)**

A mixture of tert-butyl (4-((5-methylisoxazol-3-yl)methylene)tetrahydrofuran-3-yl)carbamate (**S39**, 264 mg, 0.94 mmol), Pd/C(10%wt, 132 mg), and EtOH (20 mL) was stirred at 10 ºC for 4 h. The reaction mixture was concentrated in vacuo. The residue was purified by reverse phase silica gel column chromatography (eluted with $CH_3CN/H_2O$ = 5/95~95/5 with 0.1% TFA) to give tert-butyl (4-((5-methylisoxazol-3-yl)methyl)tetrahydrofuran-3-yl)carbamate (**S40**, 125 mg TFA salt, 26% yield) as a yellow oil.

LC-MS: MS (ESI, *m/z*): $[M+H]^+$=283.1.

**Procedure for preparation of 4-((5-methylisoxazol-3-yl)methyl)tetrahydrofuran-3-amine (S41)**

A mixture of tert-butyl (4-((5-methylisoxazol-3-yl)methyl)tetrahydrofuran-3-yl)carbamate (**S40**, 125 mg, 0.44 mmol) and DCM/TFA (3/1 mL) was stirred at room temperature for 1 h. The reaction mixture was concentrated in vacuo to give 4-((5-methylisoxazol-3-yl)methyl)tetrahydrofuran-3-amine (**S41**, 147 mg TFA salt, crude) as a yellow oil.

LC-MS: MS (ESI, *m/z*): $[M+H]^+$=183.1.

**Procedure for preparation of 3-(3-fluorophenyl)-N-(4-((5-methylisoxazol-3-yl)methyl)tetrahydrofuran-3-yl)-1H-pyrazole-5-carboxamide (1554-24)**

A mixture of 4-((5-methylisoxazol-3-yl)methyl)tetrahydrofuran-3-amine TFA salt (**S41**, 40 mg, 0.10 mmol), 3-(3-fluorophenyl)-1H-pyrazole-5-carboxylic acid (**S13**, 24 mg, 0.12 mmol), DIPEA (63 mg, 0.49 mmol), HATU (44 mg, 0.12 mmol) and dry DMF (2.0 mL) was stirred at room temperature for 1.0 h. The reaction mixture was purified by reverse phase silica gel column chromatography (eluted with $CH_3CN/H_2O$ = 5/95 ~ 95/5 with 0.1% TFA) to give 3-(3-fluorophenyl)-N-(4-((5-methylisoxazol-3-yl)methyl)tetrahydrofuran-3-yl)-1H-pyrazole-5-carboxamide (**1554-24**, 25 mg, 52% yield) TFA salt as a white solid.

LC-MS (ESI, *m/z*): $[M+H]^+$ = 371.1.

$^1$H NMR (400 MHz, $CD_3OD$, cis/trans or trans/cis = 10/7) δ 7.59 – 7.54 (m, 1.70H), 7.52 – 7.44 (m, 3.41H), 7.17 – 7.06 (m, 3.40H), 6.09 (s, 0.70H), 6.04 (s, 1.00H), 4.83 – 4.81 (m, 1.00H), 4.39 (dd, *J* = 12.4, 5.6 Hz, 0.71H), 4.13 – 4.07 (m, 2.46H), 4.01 – 3.96 (m, 1.00H), 3.82 (dd, *J* = 9.6, 4.0 Hz, 1.00H), 3.73 – 3.67 (m, 1.00H), 3.66 – 3.63 (m, 0.70H), 3.58 (dd, *J* = 8.8, 6.4 Hz, 0.73H), 2.97 – 2.91 (m, 1.00H), 2.88 – 2.83 (m, 1.71H), 2.77 (dd, *J* = 14.8, 8.8 Hz, 0.78H), 2.71 – 2.63 (m, 1.77H), 2.35 (s, 3.00H), 2.34 (s, 2.10H).

**Procedure for chiral separation of 3-(3-fluorophenyl)-N-(4-((5-methylisoxazol-3-yl)methyl)tetrahydrofuran-3-yl)-1H-pyrazole-5-carboxamide (1554-24)**

A sample of 3-(3-fluorophenyl)-N-(4-((5-methylisoxazol-3-yl)methyl)tetrahydrofuran-3-yl)-1H-pyrazole-5-carboxamide (**1554-24**, 123 mg TFA salt, cis/trans=5/3) was purified by chiral separation (see 1554-06 for conditions, eluted with $CO_2$/MeOH[0.2%$NH_3$(7M in MeOH)]=85/15) to give 3-(3-fluorophenyl)-N-(4-((5-methylisoxazol-3-yl)methyl)tetrahydrofuran-3-yl)-1H-pyrazole-5-carboxamide (27 mg, 28% yield, 3R,4S *or* 3S,4R, **1554-24-cis1**) as a white solid, 3-(3-fluorophenyl)-N-(-4-((5-methylisoxazol-3-yl)methyl)tetrahydrofuran-3-yl)-1H-pyrazole-5-carboxamide (18 mg, 19% yield, 3S,4S *or* 3R,4R, **1554-24-trans1**) as a white solid, 3-(3-fluorophenyl)-N-(4-((5-methylisoxazol-3-yl)methyl)tetrahydrofuran-3-yl)-1H-pyrazole-5-carboxamide (16 mg, 17% yield, 3S,4S *or* 3R,4R, **1554-24-trans2**) as a white solid, 3-

(3-fluorophenyl)-N-(4-((5-methylisoxazol-3-yl)methyl)tetrahydrofuran-3-yl)-1H-pyrazole-5-carboxamide (: 9 mg, 30% yield, 3R,4S *or* 3S,4R, **1554-24-cis2**) as a white solid.

LC-MS: MS (ESI, *m/z*): $[M+H]^+$=371.1

**1554-24-cis1**: $^1$H MR (400 MHz, Methanol-d$_4$) δ 7.56 (dt, J = 7.7, 1.3 Hz, 1H), 7.53 – 7.48 (m, 1H), 7.48 – 7.43 (m, 1H), 7.14 (d, J = 6.8 Hz, 1H), 7.13 – 7.03 (m, 1H), 6.04 (d, J = 1.1 Hz, 1H), 4.85 – 4.80 (m, 1H), 4.09 (dd, J = 9.3, 6.1 Hz, 1H), 3.99 (dd, J = 8.7, 7.3 Hz, 1H), 3.82 (dd, J = 9.3, 3.7 Hz, 1H), 3.66 (t, J = 8.3 Hz, 1H), 2.92 – 2.82 (m, 2H), 2.71 – 2.63 (m, 1H), 2.34 (d, J = 0.9 Hz, 3H).

**1554-24-trans1**: $^1$H NMR (400 MHz, Methanol-d$_4$) δ 7.56 (d, J = 7.8 Hz, 1H), 7.53 – 7.48 (m, 1H), 7.45 (dd, J = 8.0, 5.9 Hz, 1H), 7.11 (dt, J = 8.8, 3.1 Hz, 2H), 6.09 (d, J = 1.1 Hz, 1H), 4.39 (dt, J = 6.8, 5.4 Hz, 1H), 4.13 – 4.10 (m, 1H), 4.10 – 4.07 (m, 1H), 3.70 (dd, J = 9.1, 5.4 Hz, 1H), 3.58 (dd, J = 9.0, 6.5 Hz, 1H), 2.94 (dd, J = 14.7, 6.3 Hz, 1H), 2.77 (dd, J = 14.6, 8.6 Hz, 1H), 2.71 – 2.64 (m, 1H), 2.34 (d, J = 0.9 Hz, 3H).

**1554-24-trans2**: $^1$H NMR (400 MHz, Methanol-d$_4$) δ 7.55 (dt, J = 7.8, 1.2 Hz, 1H), 7.49 (dt, J = 7.4, 1.9 Hz, 1H), 7.47 – 7.41 (m, 1H), 7.10 (d, J = 3.0 Hz, 2H), 6.08 (s, 1H), 4.39 (dt, J = 6.8, 5.5 Hz, 1H), 4.10 (ddd, J = 8.9, 7.0, 4.3 Hz, 2H), 3.70 (dd, J = 9.1, 5.4 Hz, 1H), 3.58 (dd, J = 9.0, 6.5 Hz, 1H), 2.94 (dd, J = 14.6, 6.3 Hz, 1H), 2.77 (dd, J = 14.6, 8.6 Hz, 1H), 2.67 (dt, J = 8.4, 6.2 Hz, 1H), 2.33 (d, J = 0.9 Hz, 3H).

**1554-24-cis2**: $^1$H NMR (400 MHz, Methanol-d$_4$) δ 7.57 (dt, J = 7.8, 1.2 Hz, 1H), 7.53 – 7.49 (m, 1H), 7.48 – 7.44 (m, 1H), 7.14 (d, J = 3.7 Hz, 1H), 7.13 – 7.08 (m, 1H), 6.04 (d, J = 1.0 Hz, 1H), 4.84 – 4.81 (m, 1H), 4.09 (dd, J = 9.4, 6.0 Hz, 1H), 3.99 (dd, J = 8.7, 7.3 Hz, 1H), 3.82 (dd, J = 9.3, 3.7 Hz, 1H), 3.66 (t, J = 8.3 Hz, 1H), 2.91 – 2.87 (m, 1H), 2.86 – 2.82 (m, 1H), 2.70 – 2.62 (m, 1H), 2.35 (d, J = 0.9 Hz, 3H).

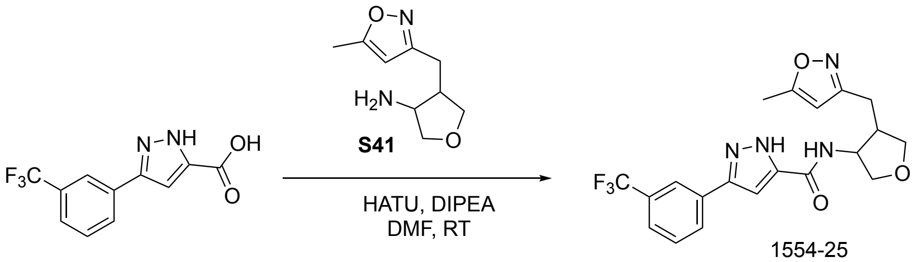


javascript:

**Procedure for preparation of N-(4-((5-methylisoxazol-3-yl)methyl)tetrahydrofuran-3-yl)-3-(3-(trifluoromethyl)phenyl)-1H-pyrazole-5-carboxamide (1554-25)**

A mixture of 4-((5-methylisoxazol-3-yl)methyl)tetrahydrofuran-3-amine (**S41**, 18 mg, 0.04 mmol), 3-(3-(trifluoromethyl)phenyl)-1H-pyrazole-5-carboxylic acid (13 mg, 0.05 mmol), HATU (23 mg, 0.06 mmol),

DIPEA (28 mg, 0.22 mmol) and dry DMF (2.0 mL) was stirred at room temperature for 1.0 h. The reaction mixture was purified by reverse phase silica gel column chromatography (eluted with $CH_3CN/H_2O$ = 5/95~95/5 with 0.1% TFA) to give N-(4-((5-methylisoxazol-3-yl)methyl)tetrahydrofuran-3-yl)-3-(3-(trifluoromethyl)phenyl)-1H-pyrazole-5-carboxamide (**1554-25**, 12 mg, TFA salt, 53% yield, cis/trans=10/7) as a white solid.

LC-MS 1554-025: MS(ESI, m/z): $[M+H]^+$=421.1

1H NMR (401 MHz, Methanol-d4) δ 8.07 (d, J = 5.6 Hz, 2H), 8.01 (s, 2H), 7.67 (d, J = 2.4 Hz, 2H), 7.66 (d, J = 2.6 Hz, 2H), 7.24 (s, 1H), 7.18 (s, 1H), 6.09 (s, 1H), 6.05 (s, 1H), 4.83 – 4.81 (m, 1H), 4.42 – 4.36 (m, 1H), 4.14 – 4.11 (m, 1H), 4.11 (d, J = 2.9 Hz, 1H), 4.08 (s, 1H), 3.99 (dd, J = 8.7, 7.3 Hz, 1H), 3.82 (dd, J = 9.3, 3.7 Hz, 1H), 3.71 (dd, J = 9.1, 5.3 Hz, 1H), 3.66 (t, J = 8.3 Hz, 1H), 3.59 (dd, J = 9.0, 6.4 Hz, 1H), 2.94 (dd, J = 14.7, 6.3 Hz, 1H), 2.88 (d, J = 5.3 Hz, 1H), 2.87 – 2.82 (m, 1H), 2.78 (dd, J = 14.7, 8.6 Hz, 1H), 2.69 (d, J = 11.0 Hz, 1H), 2.65 (d, J = 11.1 Hz, 1H), 2.35 (s, 3H), 2.34 (s, 2H).

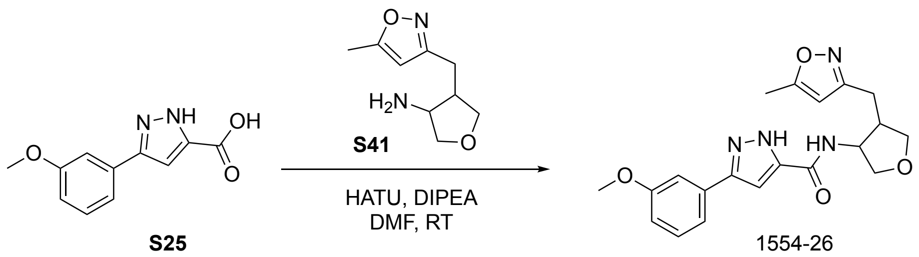


**Procedure for preparation of 3-(3-methoxyphenyl)-N-(4-((5-methylisoxazol-3-yl)methyl)tetrahydrofuran-3-yl)-1H-pyrazole-5-carboxamide (1554-26)**

A mixture of 4-((5-methylisoxazol-3-yl)methyl)tetrahydrofuran-3-amine (**S41**, 30 mg, 0.07 mmol), 3-(3-methoxyphenyl)-1H-pyrazole-5-carboxylic acid (**S25**, 19 mg, 0.09 mmol), HATU (39 mg, 0.10 mmol), DIPEA (47 mg, 0.37 mmol) and dry DMF (2.0 mL) was stirred at room temperature for 1.0 h. The reaction mixture was purified by reverse phase silica gel column chromatography (eluted with $CH_3CN/H_2O$ = 5/95~95/5 with 0.1% TFA) to give 3-(3-methoxyphenyl)-N-(4-((5-methylisoxazol-3-yl)methyl)tetrahydrofuran-3-yl)-1H-pyrazole-5-carboxamide (**1554-26**, 18 mg, TFA salt, 53% yield, cis/trans=10/9) as a white solid.

LC-MS 1554-026: MS(ESI, m/z): $[M+H]^+$=383.1

$^1$H NMR (400 MHz, Methanol-$d_4$) δ 7.39 – 7.36 (m, 1H), 7.36 – 7.33 (m, 1H), 7.31 – 7.30 (m, 1H), 7.29 (t, $J$ = 2.3 Hz, 2H), 7.28 (d, $J$ = 1.2 Hz, 1H), 7.09 (s, 1H), 7.05 (s, 1H), 6.95 (d, $J$ = 1.1 Hz, 1H), 6.93 (d, $J$ = 1.2 Hz, 1H), 6.09 (d, $J$ = 0.9 Hz, 1H), 6.04 (d, $J$ = 1.0 Hz, 1H), 4.84 – 4.81 (m, 1H), 4.41 – 4.36 (m,

1H), 4.13 – 4.11 (m, 1H), 4.11 – 4.09 (m, 1H), 4.08 (dd, $J$ = 5.7, 1.3 Hz, 1H), 3.99 (dd, $J$ = 8.7, 7.3 Hz, 1H), 3.86 (s, 3H), 3.85 (s, 3H), 3.84 – 3.80 (m, 1H), 3.73 – 3.68 (m, 1H), 3.66 (t, $J$ = 8.3 Hz, 1H), 3.58 (dd, $J$ = 8.9, 6.5 Hz, 1H), 2.94 (dd, $J$ = 14.7, 6.3 Hz, 1H), 2.88 (d, $J$ = 3.4 Hz, 1H), 2.84 (d, $J$ = 5.7 Hz, 1H), 2.80 – 2.74 (m, 1H), 2.69 (d, $J$ = 11.0 Hz, 1H), 2.66 (s, 1H), 2.35 (d, $J$ = 0.9 Hz, 3H), 2.34 (d, $J$ = 0.9 Hz, 3H).

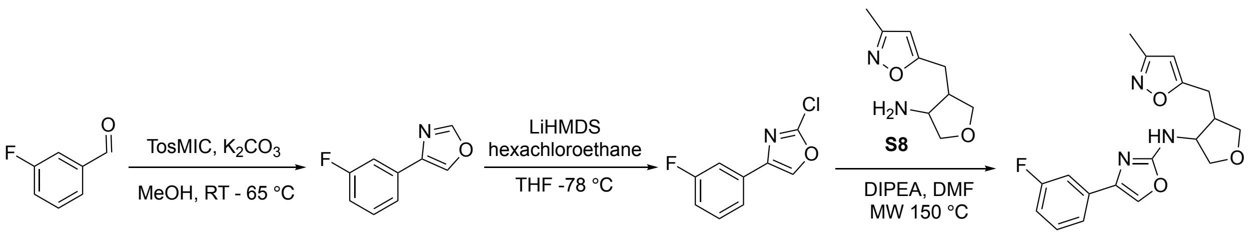


javascript:

**Procedure for preparation of 4-(3-fluorophenyl)oxazole (S42)**

To a solution of 3-fluorobenzaldehyde (300 mg, 2.42 mmol) in MeOH (5 mL) was added $K_2CO_3$ (367 mg, 2.66 mmol) at room temperature under $N_2$ atmosphere. The resulting mixture was stirred at room temperature for 10 min. Then, toluenesulfonylmethyl isocyanide (TosMIC) (519 mg, 2.66 mmol) in MeOH (5 mL) was added at room temperature. The resulting mixture was stirred at 65°C for 4 h. The reaction mixture was diluted with water (30 mL) and extracted with ethyl acetate (30 mLx3). The combined organic layers were washed with brine, dried over anhydrous $Na_2SO_4$ and concentrated in vacuo. The residue was purified by silica gel column chromatography (eluted with PE/ethyl acetate = 3/1) to give 4-(3-fluorophenyl)oxazole (**S42**, 298 mg, 75% yield) as a yellow oil.

LC-MS: MS(ESI, m/z): $[M+H]^+$=164.1.

**Procedure for preparation of 2-chloro-4-(3-fluorophenyl)oxazole (S43)**

To a solution of 4-(3-fluorophenyl)oxazole (**S42**, 298 mg, 1.83 mmol) in THF (10 mL) was added LiHMDS (1 moL/L, 2 mL) at -78°C under $N_2$ atmosphere. The resulting mixture was stirred at room temperature for 30 min. Then, hexachloroethane (476 mg, 2.01mmol) was added at room temperature. The resulting mixture was stirred at room temperature for 3.0 h. The reaction mixture was diluted with water (30 mL) and extracted with ethyl acetate (30 mLx3). The combined organic layers were washed with brine, dried over anhydrous $Na_2SO_4$, and concentrated in vacuo. The residue was purified by silica gel column chromatography (eluted with PE/ethyl acetate = 4/1) to give 2-chloro-4-(3-fluorophenyl)oxazole (**S43**, 270 mg, 74% yield) as a yellow oil.

LC-MS: MS(ESI, m/z): $[M+H]^+$=198.1

**Procedure for preparation of 4-(3-fluorophenyl)-N-(4-((3-methylisoxazol-5-yl)methyl)tetrahydrofuran-3-yl)oxazol-2-amine (1554-27)**

A mixture of 4-((3-methylisoxazol-5-yl)methyl)tetrahydrofuran-3-amine (**S8**, 70 mg, 0.17 mmol), 2-chloro-4-(3-fluorophenyl)oxazole (**S43**, 101 mg, 0.51 mmol), DIPEA (110 mg, 0.85 mmol) and dry DMF (2.0 mL) was stirred and heated at 150 °C via microwave irradiation for 2.0 h under $N_2$ atmosphere. The reaction mixture was purified by reverse phase silica gel column chromatography (eluted with $CH_3CN/H_2O$ = 5/95~95/5) to give 4-(3-fluorophenyl)-N-(4-((3-methylisoxazol-5-yl)methyl)tetrahydrofuran-3-yl)oxazol-2-amine (**1554-27**, 31 mg, 52% yield, cis/trans=5/2) as a white solid.

LC-MS: MS(ESI, m/z): $[M+H]^+$=344.1

$^1$H NMR (400 MHz, Methanol-$d_4$) δ 7.40 – 7.35 (m, 1H), 7.33 (t, $J$ = 1.4 Hz, 1H), 7.24 – 7.21 (m, 1H), 7.17 (s, 1H), 6.94 (dddd, $J$ = 9.1, 7.8, 2.6, 1.3 Hz, 1H), 5.99 (s, 1H), 4.44 (td, $J$ = 6.1, 3.7 Hz, 1H), 4.08 (d, $J$ = 5.9 Hz, 1H), 4.03 (dd, $J$ = 8.7, 7.4 Hz, 1H), 3.76 (dd, $J$ = 9.3, 3.7 Hz, 1H), 3.68 – 3.63 (m, 1H), 3.09 – 2.99 (m, 1H), 2.94 – 2.90 (m, 1H), 2.81 (dd, $J$ = 14.7, 7.8 Hz, 1H), 2.19 (s, 1H), 2.12 (s, 3H).

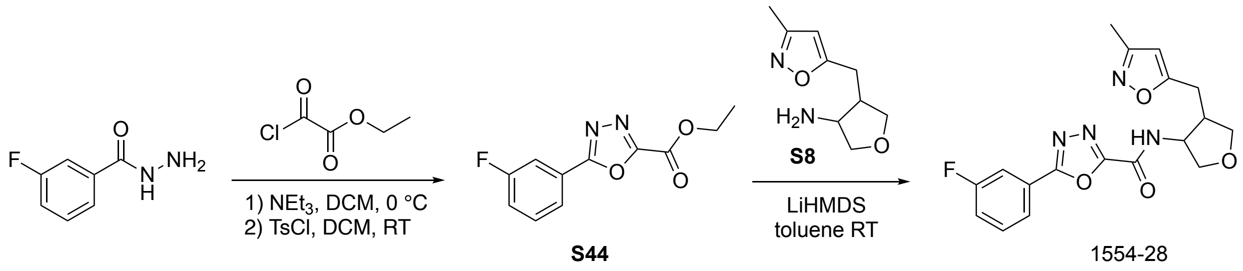


**Procedure for preparation of ethyl 5-(3-fluorophenyl)-1,3,4-oxadiazole-2-carboxylate (S44)**

To a mixture of 3-fluorobenzohydrazide (350 mg, 2.27 mmol) and $Et_3N$ (0.63 mL, 4.54 mmol) in DCM (8 mL) was added ethyl 2-chloro-2-oxoacetate (0.28 mL, 2.50 mmol) dropwise at 0 °C. The mixture was stirred at room temperature for 1.5 h. Then, $Et_3N$ (0.32 mL, 2.27 mmol) and TsCl (433 mg, 2.27 mmol) were added at room temperature. The resulting mixture was stirred at room temperature overnight. The reaction mixture was diluted with $H_2O$ (50 mL) and extracted with DCM (30 mLx3). The combined organic layers were washed with brine, dried over anhydrous $Na_2SO_4$ and concentrated in vacuo. The residue was purified by silica gel column chromatography (eluted with PE/ethyl acetate =9/1) to give ethyl 5-(3-fluorophenyl)-1,3,4-oxadiazole-2-carboxylate (**S44**, 373 mg, 70% yield) as a colorless oil.

LC-MS (ESI, *m/z*): $[M+H]^+$ = 235.0.

**Procedure for preparation of 5-(3-fluorophenyl)-N-(4-((3-methylisoxazol-5-yl)methyl)tetrahydrofuran-3-yl)-1,3,4-oxadiazole-2-carboxamide (1554-28)**

To a solution of ethyl 5-(3-fluorophenyl)-1,3,4-oxadiazole-2-carboxylate (**S44**, 30 mg, 0.13 mmol) and 4-((3-methylisoxazol-5-yl)methyl)tetrahydrofuran-3-amine (**S8**, 63 mg, 0.15 mmol) in toluene (1.5 mL) was added LiHMDS (1.0 M in THF, 0.32 mL, 0.32 mmol) at room temperature under Ar atmosphere. The resulting mixture was stirred at room temperature overnight. The reaction mixture was quenched with water (0.5 mL) and concentrated in vacuo. The residue was purified by reverse phase silica gel column chromatography (eluted with $CH_3CN/H_2O$ = 5/95 ~ 95/5 with 0.1% TFA) to give 5-(3-fluorophenyl)-N-(4-((3-methylisoxazol-5-yl)methyl)tetrahydrofuran-3-yl)-1,3,4-oxadiazole-2-carboxamide (**1554-28**, 7.9 mg, 13% yield) TFA salt as a white solid.

LC-MS (ESI, *m/z*): $[M+H]^+$ = 373.1.

$^1H$ NMR (400 MHz, $CD_3OD$, cis/trans = 1/1) δ 8.00 – 7.96 (m, 2.00H), 7.90 – 7.86 (m, 2.00H), 7.67 – 7.61 (m, 2.09H), 7.46 – 7.38 (m, 2.07H), 6.14 (s, 1.05H), 6.07 (s, 0.95H), 4.91 – 4.87 (m, 1.03H), 4.46 (dd, *J* = 12.0, 5.6 Hz, 1.09H), 4.15 – 4.08 (m, 3.06H), 4.03 (dd, *J* = 8.4, 7.2 Hz, 1.01H), 3.87 (dd, *J* = 9.6, 4.0 Hz, 1.01H), 3.78 – 3.68 (m, 2.08H), 3.61 (dd, *J* = 9.2, 6.8 Hz, 1.03H), 3.07 (dd, *J* = 15.2, 6.8 Hz, 1.08H), 3.04 – 2.98 (m, 1.05H), 2.97 – 2.89 (m, 2.05H), 2.83 (dd, *J* = 14.4, 8.0 Hz, 1.03H), 2.79 – 2.72 (m, 1.05H), 2.20 (s, 3.00H), 2.20 (s, 3.05H).

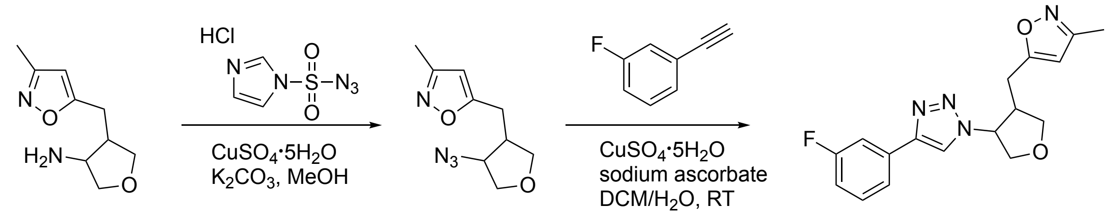


S8 S45 1554-29 javascript:

**Procedure for preparation of 5-((4-azidotetrahydrofuran-3-yl)methyl)-3-methylisoxazole (S45)**

To a mixture of 4-((3-methylisoxazol-5-yl)methyl)tetrahydrofuran-3-amine (**S8**, 30 mg, 0.07 mmol), $CuSO_4(5H_2O)$ (1 mg, 0.0015 mmol) and $K_2CO_3$ (51 mg, 0.37 mmol) in dry MeOH (2 mL) was added 1H-imidazole-1-sulfonyl azide hydrochloride (31 mg, 0.15 mmol) at room temperature under $N_2$ atmosphere. The resulting mixture was stirred at room temperature overnight. The reaction mixture was diluted with water (30 mL) and extracted with ethyl acetate (30 mLx3). The combined organic layers were washed with brine, dried over anhydrous $Na_2SO_4$ and concentrated in vacuo. The residue was purified by silica gel column chromatography (eluted with PE/ethyl acetate = 4/1) to give 5-((4-azidotetrahydrofuran-3-yl)methyl)-3-methylisoxazole (**S45**, 8 mg, 51% yield) as a yellow oil.

LC-MS: MS(ESI, m/z): $[M+H]^+$=209.1.

**Procedure for preparation of 5-((4-(4-(3-fluorophenyl)-1H-1,2,3-triazol-1-yl)tetrahydrofuran-3-yl)methyl)-3-methylisoxazole (1554-29)**

To a mixture of 5-((4-azidotetrahydrofuran-3-yl)methyl)-3-methylisoxazole (**S45**, 8 mg, 0.04 mmol), $CuSO_4(5H_2O)$ (3 mg, 0.01 mmol) and sodium ascorbate (5mg, 0.02 mmol) in $DCM/H_2O$ (1/1 mL) was added 1-ethynyl-3-fluorobenzene (5 mg, 0.04 mmol) at room temperature under $N_2$ atmosphere. The resulting mixture was stirred at room temperature overnight. The reaction mixture was concentrated in vacuo. The residue was purified by reverse phase silica gel column chromatography (eluted with $CH_3CN/H_2O$ = 5/95~95/5 with 0.1% TFA) to give 5-((4-(4-(3-fluorophenyl)-1H-1,2,3-triazol-1-yl)tetrahydrofuran-3-yl)methyl)-3-methylisoxazole (**1554-29**, 10 mg, 51% yield, cis/trans=5/4) as a yellow oil.

LC-MS: MS(ESI, m/z): $[M+H]^+$=329.1

$^1$H NMR (400 MHz, Methanol-$d_4$) δ 8.36 (s, 1H), 8.34 (s, 1H), 7.66 (dt, $J$ = 7.8, 1.3 Hz, 1H), 7.65 – 7.62 (m, 1H), 7.62 – 7.59 (m, 1H), 7.58 – 7.55 (m, 1H), 7.48 – 7.45 (m, 1H), 7.45 – 7.41 (m, 1H), 7.12 – 7.06 (m, 2H), 6.08 (s, 1H), 5.98 (s, 1H), 5.45 (ddd, $J$ = 7.1, 4.7, 2.8 Hz, 1H), 5.14 (dt, $J$ = 6.8, 4.0 Hz, 1H), 4.38 – 4.34 (m, 1H), 4.33 (d, $J$ = 3.0 Hz, 1H), 4.32 (d, $J$ = 1.1 Hz, 1H), 4.28 – 4.23 (m, 1H), 4.19 (t, $J$ = 8.5 Hz, 1H), 4.14 (dd, $J$ = 10.0, 4.1 Hz, 1H), 3.89 (t, $J$ = 9.0 Hz, 1H), 3.70 (dd, $J$ = 9.1, 6.0 Hz, 1H), 3.27 – 3.19 (m, 1H), 3.17 – 3.12 (m, 1H), 3.10 (d, $J$ = 2.6 Hz, 1H), 3.09 – 3.03 (m, 1H), 2.56 (dd, $J$ = 15.8, 7.3 Hz, 1H), 2.43 (dd, $J$ = 15.8, 8.0 Hz, 1H), 2.18 (s, 5H).

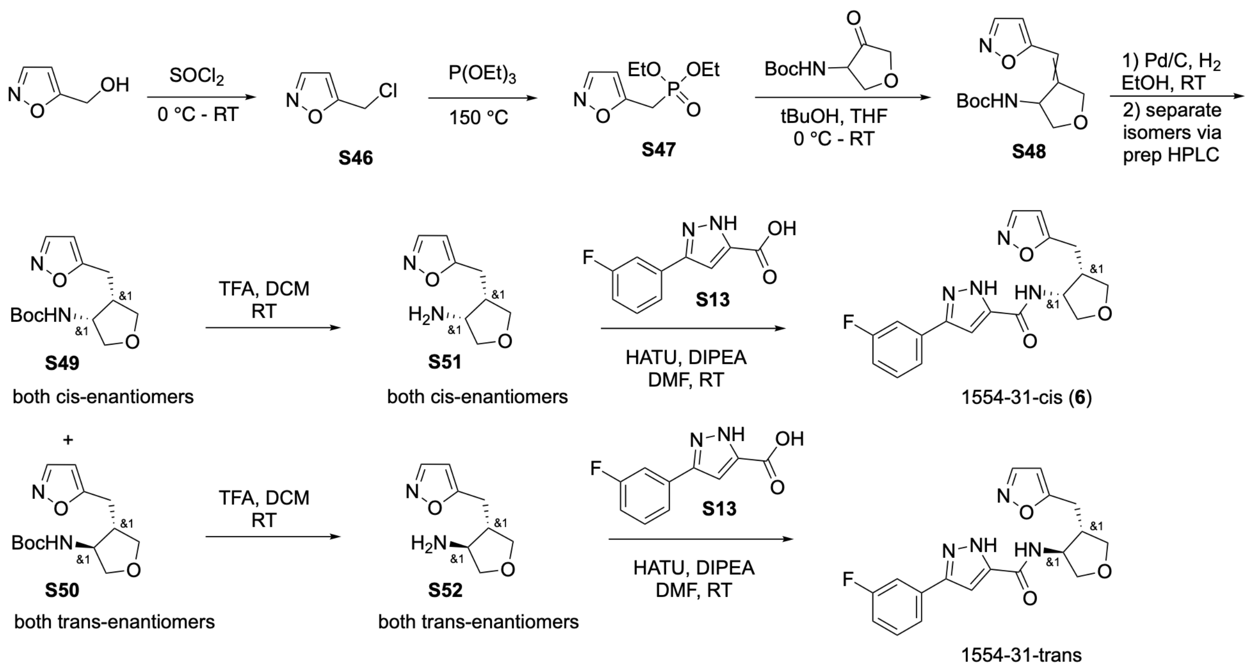


**Procedure for preparation of 5-(chloromethyl)isoxazole (S46)**

Isoxazol-5-ylmethanol (1 g, 10.10 mmol, 1.0 eq.) was added to $SOCl_2$ (10 mL) at 0 ºC. The resulting mixture was stirred at room temperature for 2.0 h. The reaction mixture was concentrated under high vacuum to give 5-(chloromethyl)isoxazole (**S46**, 2.2 g, crude) as a dark oil, which was used in the next step without further purification.

LC-MS (ESI, *m/z*): $[M+H]^+$ = 118.1.

**Procedure for preparation of diethyl (isoxazol-5-ylmethyl)phosphonate (S47)**

A mixture of 5-(chloromethyl)isoxazole (**S46**, 2.2 g, crude, 10.10 mmol, 1.0 eq.) and triethyl phosphite (10 mL) was stirred at 150 ºC for 3.0 h. The mixture was concentrated in high vacuum. The residue was purified by silica gel column chromatography (eluted with DCM/MeOH = 20/1) to afford diethyl (isoxazol-5-ylmethyl)phosphonate (**S47**, 1.5 g, 68% yield for 2 steps) as a colorless oil.

LC-MS (ESI, *m/z*): $[M+H]^+$ = 220.1.

**Procedure for preparation of tert-butyl (4-(isoxazol-5-ylmethylene)tetrahydrofuran-3-yl)carbamate (S48)**

To a solution of diethyl (isoxazol-5-ylmethyl)phosphonate (**S47**, 190 mg, 0.87 mmol, 1.0 eq.) in THF (5 mL) was added NaH (60% w/w, 42 mg, 1.04 mmol, 1.2 eq.) at 0 ºC. The resulting mixture was stirred at 0 ºC for 30 min. tert-Butyl (4-oxotetrahydrofuran-3-yl)carbamate (192 mg, 0.96 mmol, 1.1 eq.) was added and the mixture was stirred at room temperature for 1.0 h. The reaction mixture was diluted with water (10 mL), extracted with ethyl acetate (10 mL×3). The combined organic layers were dried over anhydrous $Na_2SO_4$, concentrated in vacuum, followed by purification by flash silica gel column chromatography (eluted with PE/ethyl acetate =3/1), to give tert-butyl (4-(isoxazol-5-ylmethylene)tetrahydrofuran-3-yl)carbamate (**S48**, 55 mg, 24% yield) as a white solid.

LC-MS: m/z 211.1 $(M+H-C_4H_8)^+$.

**Procedure for preparation of tert-butyl (4-(isoxazol-5-ylmethylene)tetrahydrofuran-3-yl)carbamate and separation of cis (3R,4S; 3S,4R) (S49) and trans (3R,4R; 3S,4S) (S50) isomeric mixtures**

To a mixture of tert-butyl (4-(isoxazol-5-ylmethylene)tetrahydrofuran-3-yl)carbamate (**S48**, 150 mg mg, 0.56 mmol, 1.0 eq) in EtOH (15 mL) was added Pd/C (75 mg). The resulting mixture was stirred at 10 ºC for 3 h under $H_2$ atmosphere. The reaction mixture was filtered through Celite to remove Pd/C. The mixture was concentrated in vacuo, followed by purification by prep-HPLC to give tert-butyl (4-(isoxazol-5-ylmethyl)tetrahydrofuran-3-yl)carbamate (**S49**, 32 mg, 3R,4S; 3S,4R, cis isomers), tert-butyl (4-(isoxazol-5-ylmethyl)tetrahydrofuran-3-yl)carbamate (**S50**, 32 mg, 3R,4R; 3S,4S, trans isomers) as white solids.

LC-MS (ESI, *m/z*): $[M+H-C_4H_8]^+ = 213.1$

**Procedure for preparation of (3R,4S)-4-(isoxazol-5-ylmethyl)tetrahydrofuran-3-amine and (3S,4R)-4-(isoxazol-5-ylmethyl)tetrahydrofuran-3-amine (cis isomers) (S51)**

To a solution of **S49** (32 mg, cis isomers, 0.12 mmol) in DCM (1 mL) was added TFA (0.5 mL). The resulting mixture was stirred at room temperature for 1 h. The reaction mixture was concentrated under high vacuum to give (3R,4S)-4-(isoxazol-5-ylmethyl)tetrahydrofuran-3-amine and (3S,4R)-4-(isoxazol-5-ylmethyl)tetrahydrofuran-3-amine (**S51**, cis isomers, crude, TFA salt) as a black oil.

LC-MS (ESI, *m/z*): $[M+H]^+ = 169.1$

**Procedure for preparation of (3R,4R)-4-(isoxazol-5-ylmethyl)tetrahydrofuran-3-amine and (3S,4S)-4-(isoxazol-5-ylmethyl)tetrahydrofuran-3-amine (trans isomers) (S52)**

To a solution of **S50** (27 mg, trans isomers, 0.10 mmol) in DCM (1 mL) was added TFA (0.5 mL). The resulting mixture was stirred at room temperature for 1 h. The reaction mixture was concentrated in high vacuum to give (3R,4R)-4-(isoxazol-5-ylmethyl)tetrahydrofuran-3-amine and (3S,4S)-4-(isoxazol-5-ylmethyl)tetrahydrofuran-3-amine (**S52**, crude, TFA salt) as a black oil.

LC-MS (ESI, *m/z*): $[M+H]^+$ = 169.1

**Procedure for preparation of 3-(3-fluorophenyl)-N-((3R,4S)-4-(isoxazol-5-ylmethyl)tetrahydrofuran-3-yl)-1H-pyrazole-5-carboxamide and 3-(3-fluorophenyl)-N-((3R,4S)-4-(isoxazol-5-ylmethyl)tetrahydrofuran-3-yl)-1H-pyrazole-5-carboxamide (1554-31-cis, 6)**

To a mixture of **S51** (crude, TFA salt, 0.12 mmol, 1.0 eq.) and 3-(3-fluorophenyl)-1H-pyrazole-5-carboxylic acid (**S13**, 25 mg, 0.12 mmol, 1.0 eq.) in DMF (2 mL) was added DIPEA (0.5 mL) and HATU (49 mg, 0.13 mmol, 1.1 eq). The resulting mixture was stirred at room temperature for 2 h. The reaction mixture was diluted with water (10 mL) and extracted with ethyl acetate (10 mL×3). The combined organic phases were dried over anhydrous $Na_2SO_4$ and concentrated under vacuum. The residue was purified by silica gel column chromatography (eluted with $CH_3CN/H_2O$ with 0.1% TFA = 5/95 ~ 95/5) to give 3-(3-fluorophenyl)-N-((3R,4S)-4-(isoxazol-5-ylmethyl)tetrahydrofuran-3-yl)-1H-pyrazole-5-carboxamide and 3-(3-fluorophenyl)-N-((3R,4S)-4-(isoxazol-5-ylmethyl)tetrahydrofuran-3-yl)-1H-pyrazole-5-carboxamide (**1554-31-cis**, 25 mg, mixture of cis isomers, 59% yield for 2 steps) as a white solid.

LC-MS (ESI, *m/z*): $[M+H]^+$ = 357.1

$^1$H NMR (400 MHz, Methanol-d4) δ8.24 (d, J = 1.8 Hz, 1H), 7.57 (d, J = 7.8 Hz, 1H), 7.54 – 7.42 (m, 2H), 7.16 – 7.07 (m, 2H), 6.19 (dd, J = 1.7, 0.8 Hz, 1H), 4.10 (dd, J = 9.4, 6.0 Hz, 1H), 4.02 (dd, J = 8.7, 7.3 Hz, 1H), 3.83 (dd, J = 9.3, 3.7 Hz, 1H), 3.69 (t, J = 8.3 Hz, 1H), 3.07 (dd, J = 14.3, 5.0 Hz, 1H), 2.99 – 2.81 (m, 2H).

**Procedure for preparation of 3-(3-fluorophenyl)-N-((3R,4R)-4-(isoxazol-5-ylmethyl)tetrahydrofuran-3-yl)-1H-pyrazole-5-carboxamide and 3-(3-fluorophenyl)-N-((3S,4S)-4-(isoxazol-5-ylmethyl)tetrahydrofuran-3-yl)-1H-pyrazole-5-carboxamide (1554-31-trans)**

To a mixture of **S52** (crude, TFA salt, trans isomers, 0.10 mmol, 1.0 eq.) and 3-(3-fluorophenyl)-1H-pyrazole-5-carboxylic acid (**S13**, 21 mg, 0.10 mmol, 1.0 eq.) in DMF (2 mL) was added DIPEA (0.5 mL) and HATU (42 mg, 0.11 mmol, 1.1 eq). The resulting mixture was stirred at room temperature for 2 h. The reaction mixture was diluted with water (10 mL) and extracted with ethyl acetate (10 mL×3). The combined organic phases were dried over anhydrous $Na_2SO_4$ and concentrated under vacuum. The residue was purified by silica gel column chromatography (eluted with $CH_3CN/H_2O$ with 0.1% TFA = 5/95 ~ 95/5) to give 3-(3-fluorophenyl)-N-((3R,4R)-4-(isoxazol-5-ylmethyl)tetrahydrofuran-3-yl)-1H-pyrazole-5-carboxamide and 3-(3-fluorophenyl)-N-((3S,4S)-4-(isoxazol-5-ylmethyl)tetrahydrofuran-3-

yl)-1H-pyrazole-5-carboxamide (**1554-31-trans**, 6 mg, mixture of trans isomers, 17% yield for 2 steps) as a white solid.

LC-MS (ESI, *m/z*): $[M+H]^+$ = 357.0.

$^1$H NMR (400 MHz, Methanol-d4) δ 8.26 (d, J = 1.8 Hz, 1H), 7.56 (dt, J = 7.7, 1.2 Hz, 1H), 7.53 – 7.41 (m, 2H), 7.11 (q, J = 5.9 Hz, 2H), 6.28 (d, J = 1.7 Hz, 1H), 4.42 (q, J = 5.8 Hz, 1H), 4.11 (ddd, J = 9.2, 7.0, 4.1 Hz, 2H), 3.71 (dd, J = 9.2, 5.3 Hz, 1H), 3.61 (dd, J = 9.0, 6.5 Hz, 1H), 3.15 (dd, J = 15.4, 6.5 Hz, 1H), 2.99 (dd, J = 15.4, 8.4 Hz, 1H), 2.79 – 2.66 (m, 1H).

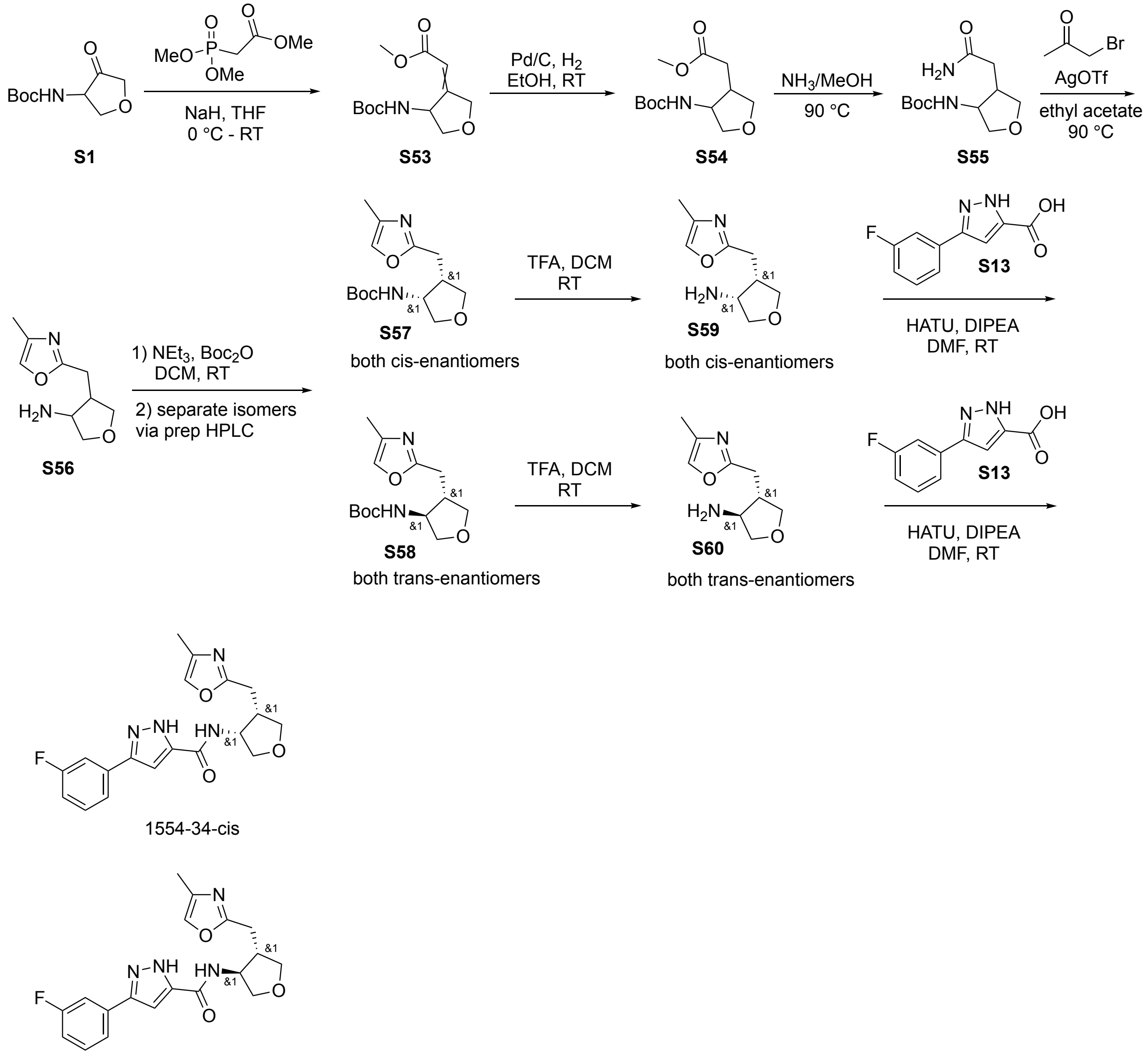


1554-34-trans

**Procedure for preparation of methyl 2-(4-((tert-butoxycarbonyl)amino)dihydrofuran-3(2H)-ylidene)acetate (S53)**

To a solution of methyl 2-(dimethoxyphosphoryl)acetate (498 mg, 2.73 mmol) in THF (10 mL) was added NaH (60%, 119 mg, 2.98 mmol) at 0 ºC. The resulting mixture was stirred at room temperature for 1.0 h. tert-Butyl (4-oxotetrahydrofuran-3-yl)carbamate (**S1**, 500 mg, 2.48 mmol) was added to the mixture at 0 ºC. The resulting mixture was stirred at room temperature overnight. The reaction mixture was diluted with water (30 mL) and extracted with ethyl acetate (30 mLx3). The combined organic layers were washed with brine, dried over anhydrous $Na_2SO_4$ and concentrated in vacuo. The residue was purified by silica gel column chromatography (eluted with PE/ethyl acetate = 4/1) to give methyl 2-(4-((tert-butoxycarbonyl)amino)dihydrofuran-3(2H)-ylidene)acetate (**S53**, 427 mg, 67% yield) as a colorless oil.

LC-MS (ESI, *m/z*): $[M+H-C_4H_8]^+$ = 202.1.

**Procedure for preparation of methyl 2-(4-((tert-butoxycarbonyl)amino)tetrahydrofuran-3-yl)acetate (S54)**

A mixture of methyl 2-(4-((tert-butoxycarbonyl)amino)dihydrofuran-3(2H)-ylidene)acetate (**S53**, 427 mg, 1.66 mmol) in EtOH (10 mL) and Pd/C (107 mg) was stirred at room temperature under $H_2$ atmosphere overnight. The reaction mixture was filtered on Celite and rinsed with MeOH. The mixture was concentrated in vacuo to give methyl 2-(4-((tert-butoxycarbonyl)amino)tetrahydrofuran-3-yl)acetate (**S54**, 425 mg, 99% yield) as a colorless oil.

LC-MS (ESI, *m/z*): $[M+H-C_4H_8]^+$ = 204.1.

**Procedure for preparation of tert-butyl (4-(2-amino-2-oxoethyl)tetrahydrofuran-3-yl)carbamate (S55)**

A mixture of methyl 2-(4-((tert-butoxycarbonyl)amino)tetrahydrofuran-3-yl)acetate (445 mg, 1.72 mmol) and $NH_3$ in MeOH (7 N, 8.0 mL) was stirred at 90 ºC overnight. The reaction mixture was concentrated in vacuo. The residue was purified by silica gel column chromatography (eluted with DCM/MeOH = 15/1) to give tert-butyl (4-(2-amino-2-oxoethyl)tetrahydrofuran-3-yl)carbamate (**S55**, 318 mg, 72% yield for two steps) as a white solid.

LC-MS (ESI, *m/z*): $[M+H]^+$ = 245.1.

**Procedure for preparation of tert-butyl (4-((4-methyloxazol-2-yl)methyl)tetrahydrofuran-3-yl)carbamate (S56)**

A mixture of tert-butyl (4-(2-amino-2-oxoethyl)tetrahydrofuran-3-yl)carbamate (**S55**, 318 mg, 1.30 mmol), 1-bromopropan-2-one (446 mg, 3.25 mmol), AgOTf (669 mg, 2.60 mmol) and ethyl acetate (10 mL) was stirred at 90 °C overnight under $N_2$ atmosphere. The reaction mixture was filtered and rinsed with ethyl acetate. The filtrate was concentrated in vacuo to give 4-((4-methyloxazol-2-yl)methyl)tetrahydrofuran-3-amine (S56, 240 mg) as a crude brown oil.

LC-MS (ESI, *m/z*): $[M+H]^+$ = 183.1.

**Procedure for preparation of tert-butyl (4-((4-methyloxazol-2-yl)methyl)tetrahydrofuran-3-yl)carbamate and separation of cis (3R,4S; 3S,4R) (S57) and trans (3R,4R; 3S,4S) (S58) isomeric mixtures**

A mixture of 4-((4-methyloxazol-2-yl)methyl)tetrahydrofuran-3-amine (**S56**, 240 mg, 1.32 mmol), $Boc_2O$ (575 mg, 2.63 mmol), $Et_3N$ (400 mg, 3.95 mmol) and DCM (5.0 mL) was stirred at room temperature for 2.0 h. The reaction mixture was concentrated in vacuo. The residue was purified by reverse phase silica gel column chromatography (eluted with $CH_3CN/H_2O$ = 5/95 ~ 95/5) to give tert-butyl (4-((4-methyloxazol-2-yl)methyl)tetrahydrofuran-3-yl)carbamate (120 mg, 33% yield for two steps) as a brown oil.

LC-MS (ESI, *m/z*): $[M+H]^+$ = 283.2.

The tert-butyl (4-((4-methyloxazol-2-yl)methyl)tetrahydrofuran-3-yl)carbamate (120 mg, 0.43 mmol) was purified by prep-HPLC (eluted with $CH_3CN/H_2O$ = 5/95 ~ 95/5 with 0.1% TFA) to give tert-butyl ((3R,4S)-4-((4-methyloxazol-2-yl)methyl)tetrahydrofuran-3-yl)carbamate and ((3S,4R)-4-((4-methyloxazol-2-yl)methyl)tetrahydrofuran-3-yl)carbamate (**S57**, 19 mg, 16% yield, cis isomers) as a colorless oil and tert-butyl ((3R,4R)-4-((4-methyloxazol-2-yl)methyl)tetrahydrofuran-3-yl)carbamate and tert-butyl ((3S,4S)-4-((4-methyloxazol-2-yl)methyl)tetrahydrofuran-3-yl)carbamate (**S58**, 62 mg, 52% yield, trans isomers) as a colorless oil.

LC-MS (ESI, *m/z*): $[M+H]^+$ = 283.2.

**Procedure for preparation of (3R,4S)-4-((4-methyloxazol-2-yl)methyl)tetrahydrofuran-3-amine and (3S,4R)-4-((4-methyloxazol-2-yl)methyl)tetrahydrofuran-3-amine (S59, cis isomers)**

A mixture of **S57** (19 mg, cis isomers, 0.07 mmol) in DCM (3 mL) and TFA (1 mL) was stirred at room temperature for 1.0 h. The reaction mixture was concentrated in vacuo to give (3R,4S)-4-((4-methyloxazol-2-yl)methyl)tetrahydrofuran-3-amine and (3S,4R)-4-((4-methyloxazol-2-yl)methyl)tetrahydrofuran-3-amine (**S59**, 29 mg, cis isomers) crude TFA salt as a colorless oil.

LC-MS (ESI, *m/z*): $[M+H]^+$ = 183.2.

**Procedure for preparation of (3R,4R)-4-((4-methyloxazol-2-yl)methyl)tetrahydrofuran-3-amine and (3S,4S)-4-((4-methyloxazol-2-yl)methyl)tetrahydrofuran-3-amine (S60, trans isomers)**

A mixture of **S58** (44 mg, trans isomers, 0.16 mmol) in DCM (3 mL) and TFA (1 mL) was stirred at room temperature for 1.0 h. The reaction mixture was concentrated in vacuo to give (3R,4R)-4-((4-methyloxazol-2-yl)methyl)tetrahydrofuran-3-amine and (3S,4S)-4-((4-methyloxazol-2-yl)methyl)tetrahydrofuran-3-amine (**S60**, 66 mg, trans isomers) crude TFA salt as a colorless oil.

LC-MS (ESI, *m/z*): $[M+H]^+$ = 183.2.

**Procedure for preparation of 3-(3-fluorophenyl)-N-((3R,4S)-4-((4-methyloxazol-2-yl)methyl)tetrahydrofuran-3-yl)-1H-pyrazole-5-carboxamide and 3-(3-fluorophenyl)-N-((3S,4R)-4-((4-methyloxazol-2-yl)methyl)tetrahydrofuran-3-yl)-1H-pyrazole-5-carboxamide (1554-34-cis)**

A mixture of **S59** (39 mg, cis isomers, 0.10 mmol), 3-(3-fluorophenyl)-1H-pyrazole-5-carboxylic acid (**S13**, 24 mg, 0.11 mmol), DIPEA (61 mg, 0.48 mmol), HATU (43 mg, 0.11 mmol) and dry DMF (3.0 mL) was stirred at room temperature for 1.0 h. The reaction mixture was purified by reverse phase silica gel column chromatography (eluted with $CH_3CN/H_2O$ = 5/95 ~ 95/5 with 0.1% TFA) to give 3-(3-fluorophenyl)-N-((3R,4S)-4-((4-methyloxazol-2-yl)methyl)tetrahydrofuran-3-yl)-1H-pyrazole-5-carboxamide and 3-(3-fluorophenyl)-N-((3S,4R)-4-((4-methyloxazol-2-yl)methyl)tetrahydrofuran-3-yl)-1H-pyrazole-5-carboxamide (**1554-34-cis**, 12 mg, cis isomeric mixture, 25% yield) TFA salt as a white solid.

LC-MS (ESI, *m/z*): $[M+H]^+$ = 371.1.

$^1$H NMR (400 MHz, $CD_3OD$, cis) δ 7.56 (d, *J* = 8.0 Hz, 1H), 7.53 – 7.42 (m, 3H), 7.16 – 7.05 (m, 2H), 4.84 – 4.82 (m, 1H), 4.10 (dd, *J* = 9.2, 6.4 Hz, 1H), 4.03 (dd, *J* = 8.8, 7.6 Hz, 1H), 3.81 (dd, *J* = 9.2, 4.0 Hz, 1H), 3.69 (t, *J* = 8.0 Hz, 1H), 3.06 – 2.94 (m, 2H), 2.84 – 2.75 (m, 1H), 2.06 (d, *J* = 1.2 Hz, 3H).

**Procedure for preparation of 3-(3-fluorophenyl)-N-((3R,4R)-4-((4-methyloxazol-2-yl)methyl)tetrahydrofuran-3-yl)-1H-pyrazole-5-carboxamide and 3-(3-fluorophenyl)-N-((3S,4S)-4-((4-methyloxazol-2-yl)methyl)tetrahydrofuran-3-yl)-1H-pyrazole-5-carboxamide (1554-34-trans)**

A mixture of **S60** (66 mg, trans, isomers, 0.16 mmol), 3-(3-fluorophenyl)-1H-pyrazole-5-carboxylic acid (**S13**, 40 mg, 0.19 mmol), DIPEA (104 mg, 0.80 mmol), HATU (73 mg, 0.19 mmol) and dry DMF (3.0 mL) was stirred at room temperature for 1.0 h. The reaction mixture was purified by reverse phase silica gel column chromatography (eluted with $CH_3CN/H_2O$ = 5/95 ~ 95/5 with 0.1% TFA) to give 3-(3-

fluorophenyl)-N-((3R,4R)-4-((4-methyloxazol-2-yl)methyl)tetrahydrofuran-3-yl)-1H-pyrazole-5-carboxamide and 3-(3-fluorophenyl)-N-((3S,4S)-4-((4-methyloxazol-2-yl)methyl)tetrahydrofuran-3-yl)-1H-pyrazole-5-carboxamide (**1554-34-trans**, 16 mg, mixture of trans isomers, 21% yield) TFA salt as a white solid.

LC-MS (ESI, *m/z*): $[M+H]^+$ = 371.1.

$^1$H NMR (400 MHz, $CD_3OD$, trans) δ 7.58 – 7.53 (m, 1H), 7.53 – 7.47 (m, 2H), 7.47 – 7.44 (m, 1H), 7.14 – 7.05 (m, 2H), 4.41 (dd, *J* = 12.4, 5.6 Hz, 1H), 4.16 – 4.10 (m, 2H), 3.71 (dd, *J* = 9.2, 5.2 Hz, 1H), 3.62 (dd, *J* = 8.8, 6.4 Hz, 1H), 3.08 (dd, *J* = 15.2, 6.8 Hz, 1H), 2.95 (dd, *J* = 15.2, 8.0 Hz, 1H), 2.82 – 2.73 (m, 1H), 2.10 (d, *J* = 1.2 Hz, 3H).

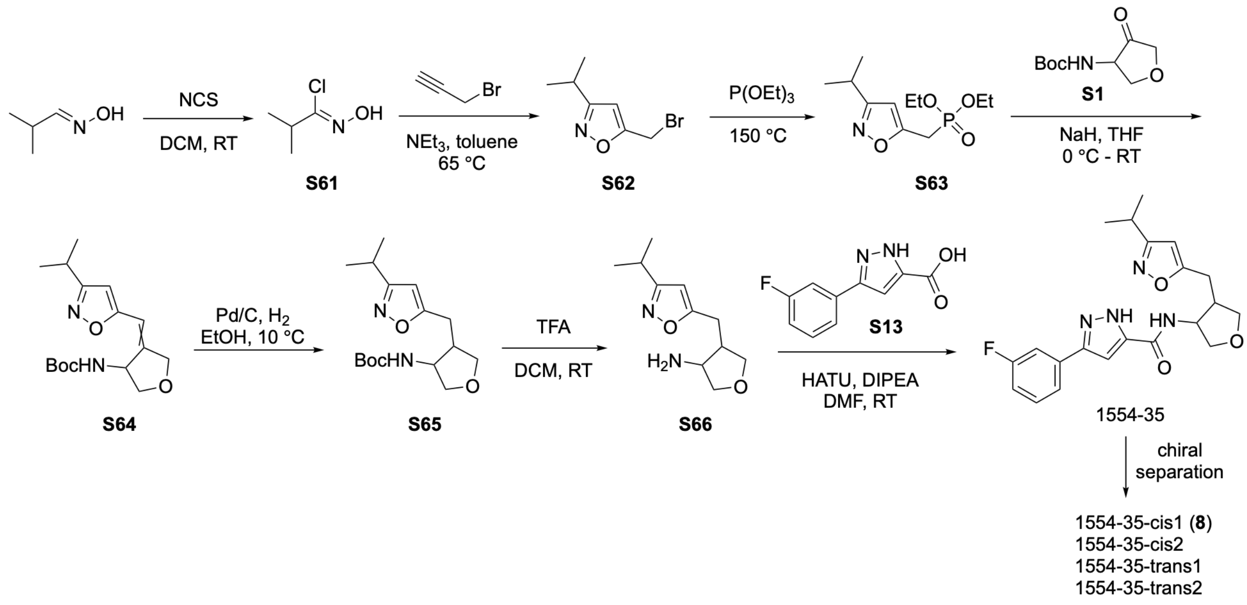


**Procedure for preparation of (Z)-N-hydroxyisobutyrimidoyl chloride (S61)**

To a solution of isobutyraldehyde oxime (600 mg, 6.89 mmol, 1.0 eq.) in DMF (5 mL) was added NCS (1.01 g, 6.89 mmol, 1.0 eq.) at 0 ℃. The resulting mixture was stirred at room temperature for 1.0 h. The reaction mixture was diluted with water (10 mL), and was extracted with ethyl acetate (10 mL×3). The combined organic layers were dried over anhydrous $Na_2SO_4$ and concentrated under vacuum to give (Z)-N-hydroxyisobutyrimidoyl chloride (**S61**, 860 mg, crude) as a light-yellow oil, which was used in the next step without further purification.

LC-MS (ESI, *m/z*): $[M+H]^+$ = 122.1.

**Procedure for preparation of 5-(bromomethyl)-3-isopropylisoxazole (S62)**

To a solution of (Z)-N-hydroxyisobutyrimidoyl chloride (**S61**, 730 mg, 6.01 mmol, 1.0 eq.) in DMF (5 mL) was added propargyl bromide (1.07 g, 9.01 mmol, 1.5 eq.) and $K_2CO_3$ (1.66 g, 12.02 mmol, 3.0 eq.). The resulting mixture was stirred at room temperature for 1 h. The reaction mixture was diluted with water (30 mL) and extracted with ethyl acetate (30 mL×3). The combined organic layers were dried over anhydrous $Na_2SO_4$, concentrated in vacuum, followed by purification by flash column chromatography (eluted with PE), to give 5-(bromomethyl)-3-isopropylisoxazole (**S62**, 830 mg, 70% yield) as a light-yellow oil.

$^1$H NMR (400 MHz, Chloroform-d) δ 6.19 (d, J = 1.7 Hz, 1H), 4.51 (dd, J = 59.8, 0.6 Hz, 2H), 3.06 (heptd, J = 7.0, 4.2 Hz, 1H), 1.29 (dd, J = 7.0, 1.6 Hz, 7H).

**Procedure for preparation of diethyl ((3-isopropylisoxazol-5-yl)methyl)phosphonate (S63)**

A mixture of 5-(bromomethyl)-3-isopropylisoxazole (**S62**, 830 mg, 4.07 mmol, 1.0 equiv) and triethyl phosphite (50 mL) was stirred at 150 °C for 3 h. The mixture was concentrated under vacuum. The residue was purified by silica gel column chromatography (eluted with DCM/MeOH = 20/1) to afford diethyl ((3-isopropylisoxazol-5-yl)methyl)phosphonate (**S63**, 1.06 g, 99% yield) as a color-less oil.

LC-MS (ESI, *m/z*): $[M+H]^+$ = 262.1.

**Procedure for preparation of tert-butyl (4-((3-isopropylisoxazol-5-yl)methylene)tetrahydrofuran-3-yl)carbamate (S64)**

To a solution of diethyl ((3-isopropylisoxazol-5-yl)methyl)phosphonate (**S63**, 1.06 g, 4.06 mmol, 1.0 eq.) in THF (15 mL) at 0 °C was added NaH (60% w/w, 117 mg, 4.87 mmol, 1.2 eq.). The resulting mixture was stirred at 0 °C for 30 min. tert-Butyl (4-oxotetrahydrofuran-3-yl)carbamate (**S1**, 816 mg, 4.06 mmol, 1.0 eq.) was added and the mixture was stirred at room temperature for 1 h. The reaction mixture was diluted with water (20 mL) and extracted with ethyl acetate (20 mL×3). The combined organic layers were dried over anhydrous $Na_2SO_4$, concentrated in vacuum, followed by purification by flash silica gel column chromatography (eluted with PE/ethyl acetate =3/1) to give tert-butyl (4-((3-isopropylisoxazol-5-yl)methylene)tetrahydrofuran-3-yl)carbamate (**S64**, 470 mg, 38% yield) as a white solid.

LC-MS: m/z 253.1 $(M+H-C_4H_8)^+$.

**Procedure for preparation of tert-butyl (4-((3-isopropylisoxazol-5-yl)methyl)tetrahydrofuran-3-yl)carbamate (S65)**

To a mixture of tert-butyl (4-((3-isopropylisoxazol-5-yl)methylene)tetrahydrofuran-3-yl)carbamate (**S64**, 470 mg, 1.52 mmol, 1.0 eq) in EtOH (25 mL) was added Pd/C (10% w/w, 230 mg), the resulting mixture was stirred at 10 °C for 4 h under $H_2$ atmosphere. The reaction mixture was filtered through Celite to remove Pd/C. The mixture was concentrated in vacuo, followed by purification by reverse phase silica gel column chromatography (eluted with $CH_3CN/H_2O$ with 0.1% TFA=5/95~95/5) to give tert-butyl (4-((3-isopropylisoxazol-5-yl)methyl)tetrahydrofuran-3-yl)carbamate (**S65**, 375 mg, 79% yield) as a light-yellow oil.

LC-MS (ESI, *m/z*): $[M+H-C_4H_8]^+$ = 255.1

**Procedure for preparation of 4-((3-isopropylisoxazol-5-yl)methyl)tetrahydrofuran-3-amine (S66)**

To a solution of tert-butyl (4-((3-isopropylisoxazol-5-yl)methyl)tetrahydrofuran-3-yl)carbamate (**S65**, 375 mg, 1.21 mmol, 1.0 eq) in DCM (5 mL) was added TFA (2 mL) and the resulting mixture was stirred at room temperature for 1 h. The reaction mixture was concentrated under vacuum to give 4-((3-isopropylisoxazol-5-yl)methyl)tetrahydrofuran-3-amine (**S66**, crude, TFA salt) as a black oil.

LC-MS (ESI, *m/z*): $[M+H-C_4H_8]^+$ = 211.1

**Procedure for preparation of 3-(3-fluorophenyl)-N-(4-((3-isopropylisoxazol-5-yl)methyl)tetrahydrofuran-3-yl)-1H-pyrazole-5-carboxamide (1554-35)**

To a mixture of 4-((3-isopropylisoxazol-5-yl)methyl)tetrahydrofuran-3-amine (**S66**, crude, TFA salt, 1.21 mmol, 1.0 eq.) and 3-(3-fluorophenyl)-1H-pyrazole-5-carboxylic acid (**S13**, 293 mg, 1.42 mmol, 1.2 eq.) in DMF (3 mL) was added DIPEA (0.5 mL) and HATU (695 mg, 1.83 mmol, 1.5 eq). The resulting mixture was stirred at room temperature for 2.0 h. The reaction mixture was diluted with water (20 mL) and extracted with ethyl acetate (20 mL×3). The combined organic phases were dried over anhydrous $Na_2SO_4$ and concentrated under vacuum. The residue was purified by silica gel column chromatography (eluted with $CH_3CN/H_2O$ with 0.1% TFA = 5/95 ~ 95/5) to give 3-(3-fluorophenyl)-N-(4-((3-isopropylisoxazol-5-yl)methyl)tetrahydrofuran-3-yl)-1H-pyrazole-5-carboxamide (**1554-35**, 150 mg, 31% yield) as a white solid.

LC-MS (ESI, *m/z*): $[M+H]^+$ = 399.2.

***Procedure for chiral separation of* 3-(3-fluorophenyl)-N-(4-((3-isopropylisoxazol-5-yl)methyl)tetrahydrofuran-3-yl)-1H-pyrazole-5-carboxamide (1554-35)**

3-(3-fluorophenyl)-N-(4-((3-isopropylisoxazol-5-yl)methyl)tetrahydrofuran-3-yl)-1H-pyrazole-5-carboxamide (**1554-35**, 150 mg, 0.38 mmol) was separated by chiral preparation (for method, see 1554-06, eluent CO2/EtOH[0.5%NH3(7M in MeOH)]=45/55) to give 3-(3-fluorophenyl)-N-(4-((3-isopropylisoxazol-5-yl)methyl)tetrahydrofuran-3-yl)-1H-pyrazole-5-carboxamide (27.5 mg, 3R,4S *or* 3S,4R, **1554-35-cis1, 8**), 3-(3-fluorophenyl)-N-(4-((3-isopropylisoxazol-5-yl)methyl)tetrahydrofuran-3-

yl)-1H-pyrazole-5-carboxamide (41.4 mg, 3R,4S *or* 3S,4R, **1554-24-cis2**), 3-(3-fluorophenyl)-N-((3S,4S)-4-((3-isopropylisoxazol-5-yl)methyl)tetrahydrofuran-3-yl)-1H-pyrazole-5-carboxamide (8.7 mg, 3R,4R *or* 3S,4S, **1554-35-trans1**), and 3-(3-fluorophenyl)-N-((3R,4R)-4-((3-isopropylisoxazol-5-yl)methyl)tetrahydrofuran-3-yl)-1H-pyrazole-5-carboxamide (9.4 mg, 3R,4R *or* 3S,4S, **1554-35-trans2**).

**1554-35-cis1 (8)**: $^{1}$H NMR (400 MHz, Methanol-d4) δ 7.56 (d, J = 7.8 Hz, 1H), 7.53 – 7.42 (m, 2H), 7.12 (dd, J = 11.8, 5.3 Hz, 2H), 6.10 (s, 1H), 4.85 – 4.81 (m, 1H), 4.06 (ddd, J = 25.3, 9.0, 6.8 Hz, 2H), 3.81 (dd, J = 9.3, 4.0 Hz, 1H), 3.69 (t, J = 8.3 Hz, 1H), 3.02 (dd, J = 15.0, 6.1 Hz, 1H), 2.92 (dq, J = 13.9, 6.9 Hz, 2H), 2.80 (dd, J = 15.0, 8.6 Hz, 1H), 1.20 (d, J = 7.0 Hz, 6H).

LC-MS (ESI, *m/z*): [M + H]+ = 399.2.

**1554-24-cis2**: $^{1}$H NMR (400 MHz, Methanol-d4) δ 7.56 (dt, J = 7.7, 1.3 Hz, 1H), 7.53 – 7.42 (m, 2H), 7.12 (tt, J = 6.6, 3.2 Hz, 2H), 6.10 (s, 1H), 4.85 – 4.81 (m, 1H), 4.06 (ddd, J = 25.3, 9.0, 6.8 Hz, 2H), 3.81 (dd, J = 9.3, 4.0 Hz, 1H), 3.69 (t, J = 8.3 Hz, 1H), 3.02 (ddd, J = 15.0, 6.2, 0.8 Hz, 1H), 2.92 (dq, J = 13.9, 6.9 Hz, 2H), 2.80 (ddd, J = 14.9, 8.6, 0.8 Hz, 1H), 1.20 (d, J = 7.0 Hz, 6H).

LC-MS (ESI, *m/z*): [M + H]+ = 399.2.

**1554-35-trans1**: $^{1}$H NMR (400 MHz, Methanol-d4) δ 7.55 (d, J = 7.6 Hz, 1H), 7.47 (t, J = 8.3 Hz, 2H), 7.09 (d, J = 9.5 Hz, 2H), 6.19 (s, 1H), 4.43 (q, J = 6.2 Hz, 1H), 4.12 (ddd, J = 12.1, 9.0, 7.1 Hz, 2H), 3.69 (dd, J = 9.1, 5.7 Hz, 1H), 3.60 (dd, J = 8.9, 6.8 Hz, 1H), 3.05 (dd, J = 15.4, 7.2 Hz, 1H), 2.99 – 2.84 (m, 2H), 2.71 (td, J = 7.4, 6.0 Hz, 1H), 1.18 (dd, J = 7.0, 2.1 Hz, 6H).

LC-MS (ESI, *m/z*): [M + H]+ = 399.2.

**1554-35-trans2**: $^{1}$H NMR (400 MHz, Methanol-d4) δ 7.57 (d, J = 7.8 Hz, 1H), 7.54 – 7.44 (m, 2H), 7.12 (d, J = 10.8 Hz, 2H), 6.21 (s, 1H), 4.45 (dt, J = 7.1, 5.8 Hz, 1H), 4.14 (ddd, J = 12.3, 9.0, 7.1 Hz, 2H), 3.71 (dd, J = 9.1, 5.7 Hz, 1H), 3.63 (dd, J = 9.0, 6.8 Hz, 1H), 3.07 (dd, J = 15.4, 7.2 Hz, 1H), 2.94 (ddd, J = 14.0, 9.3, 7.4 Hz, 2H), 2.73 (h, J = 7.2 Hz, 1H), 1.21 (dd, J = 7.0, 2.1 Hz, 6H).

LC-MS (ESI, *m/z*): [M + H]+ = 399.2.

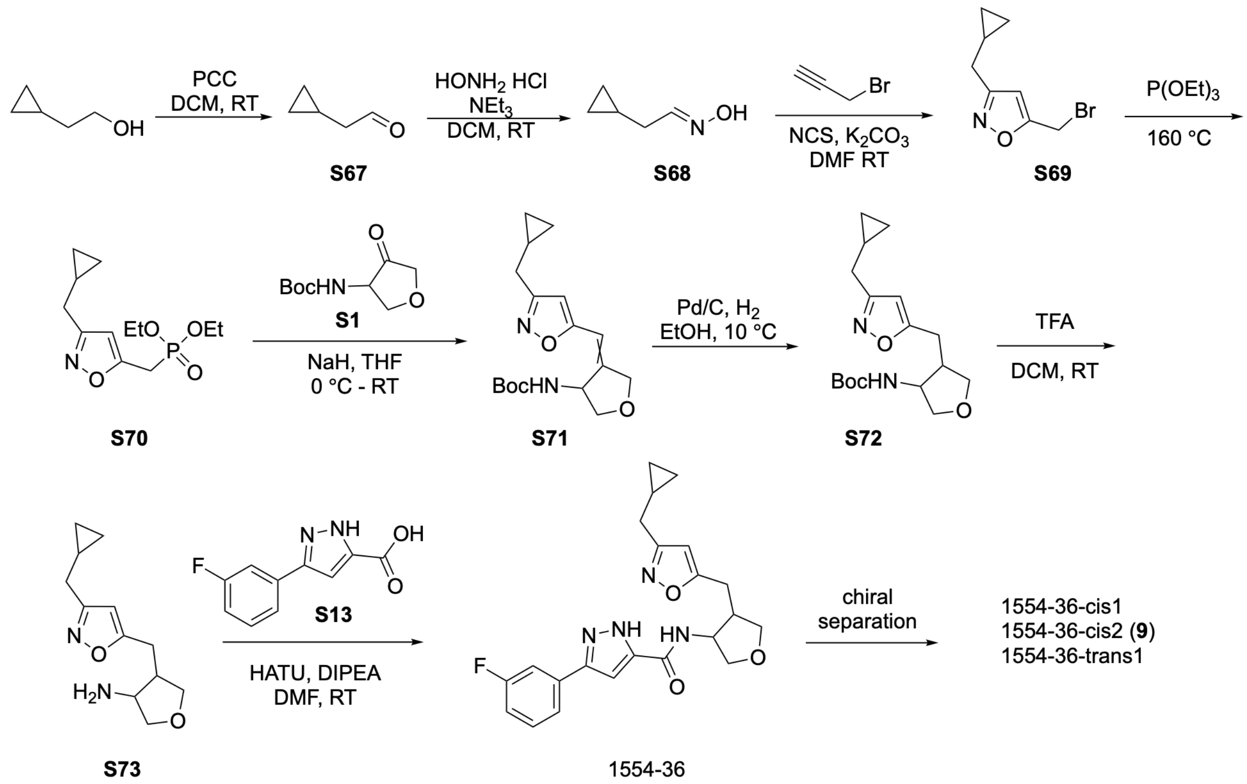


**Procedure for preparation of 2-cyclopropylacetaldehyde (S67)**

To a solution of 2-cyclopropylethan-1-ol (6.0 g, 69.7 mmol, 1.0 eq.) in DCM (100 mL) was added PCC (30.0 g, 139.3 mmol, 2 eq.) at 0 ºC. The resulting mixture was stirred at room temperature for 2 h. The reaction mixture was filtered, the filtrate was concentrated in vacuo, followed by purification by flash silica gel column chromatography (eluted with PE), give 2-cyclopropylacetaldehyde (**S67**, 5.5 g, 94% yield) as a color-less oil.

LC-MS (ESI, *m/z*): $[M+H]^+$ = 85.1.

**Procedure for preparation of 2-cyclopropylacetaldehyde oxime (S68)**

To a solution of 2-cyclopropylacetaldehyde (**S67**, 5.50 g, 65.4 mmol, 1.0 eq.) in DCM (50 mL) was added $NH_2OH{\cdot}HCl$ (5.45 g, 69.5 mmol, 1.2 eq.) and $Et_3N$ (6.62 g, 65.4 mmol, 1.0 eq.), The resulting mixture was stirred at room temperature for 2 h. The reaction mixture was concentrated in vacuo, the residue was diluted with PE (200 mL) and stirred for 30 min. The mixture was filtered through celite. The filtrate was concentrated to give 2-cyclopropylacetaldehyde oxime (**S68**, 4.7 g, 73% yield) as a yellow oil.

LC-MS (ESI, *m/z*): $[M+H]^+$ = 100.1.

**Procedure for preparation of 5-(bromomethyl)-3-(cyclopropylmethyl)isoxazole (S69)**

To a solution of 2-cyclopropylacetaldehyde oxime (**S68**, 4.7 g, 47.4 mmol, 1.0 eq.) in DMF (100 mL) was added NCS (6.96 g, 52.1 mmol, 1.1 eq.) at 0 °C. The resulting mixture was stirred at room temperature for 1 h. Then propargyl bromide (8.46 g, 71.1 mmol, 1.5 eq.) and $K_2CO_3$ (19.6 g, 142.2 mmol, 3.0 eq.) was added in sequence. The resulting mixture was stirred at room temperature for 1 h. The reaction mixture was diluted with water (200 mL), extracted with ethyl acetate (200 mL×3). The combined organic layers were dried over anhydrous $Na_2SO_4$, concentrated in vacuo, followed by purification by flash silica gel column chromatography (eluted with PE), give 5-(bromomethyl)-3-(cyclopropylmethyl)isoxazole (**S69**, 3.3 g, 32% yield) as a light-yellow oil.

$^1$H NMR (400 MHz, Chloroform-d) δ 6.28 (s, 1H), 4.59 (d, J = 0.7 Hz, 2H), 2.57 (d, J = 7.2 Hz, 2H), 1.09 – 0.93 (m, 1H), 0.69 – 0.52 (m, 2H), 0.33 – 0.20 (m, 2H).

**Procedure for preparation of diethyl ((3-(cyclopropylmethyl)isoxazol-5-yl)methyl)phosphonate (S70)**

A mixture of 5-(bromomethyl)-3-(cyclopropylmethyl)isoxazole (**S69**, 1.8 g, 8.33 mmol, 1.0 equiv), triethyl phosphite (20 mL) was stirred at 150 °C for 3 h. The mixture was concentrated under vacuum. The residue was purified by silica gel column chromatography (eluted with DCM/MeOH = 20/1) to afford diethyl ((3-(cyclopropylmethyl)isoxazol-5-yl)methyl)phosphonate (**S70**, 830 mg, 36% yield) as a color-less oil.

LC-MS (ESI, *m/z*): $[M+H]^+$ = 274.1.

**Procedure for preparation of tert-butyl (4-((3-(cyclopropylmethyl)isoxazol-5-yl)methylene)tetrahydrofuran-3-yl)carbamate (S71)**

To a solution of diethyl ((3-(cyclopropylmethyl)isoxazol-5-yl)methyl)phosphonate (**S70**, 830 mg, 3.04 mmol, 1.0 eq.) in THF (15 mL) was added NaH (60% w/w, 146 mg, 3.65 mmol, 1.2 eq.) at 0 °C. The resulting mixture was stirred at 0 °C for 30 min. tert-Butyl (4-oxotetrahydrofuran-3-yl)carbamate (**S1**, 611 mg, 3.04 mmol, 1.0 eq.) was added and the mixture was stirred at room temperature for 1 h. The reaction mixture was diluted with water (20 mL) and extracted with ethyl acetate (20 mL×3). The combined organic layers were dried over anhydrous $Na_2SO_4$, concentrated in vacuo, followed by purification by flash silica gel column chromatography (eluted with PE/ethyl acetate =3/1), give tert-butyl (4-((3-(cyclopropylmethyl)isoxazol-5-yl)methylene)tetrahydrofuran-3-yl)carbamate (**S71**, 410 mg, 42% yield) as a white solid.

LC-MS: m/z 265.1 $(M+H-C_4H_8)^+$.

**Procedure for preparation of tert-butyl (4-((3-(cyclopropylmethyl)isoxazol-5-yl)methyl)tetrahydrofuran-3-yl)carbamate (S72)**

To a mixture of tert-butyl (4-((3-(cyclopropylmethyl)isoxazol-5-yl)methylene)tetrahydrofuran-3-yl)carbamate (**S71**, 410 mg, 1.28 mmol, 1.0 eq) in EtOH (25 mL) was added Pd/C (10% w/w, 200 mg). The resulting mixture was stirred at 10 °C for 4 h under $H_2$ atmosphere. The reaction mixture was filtered over celite to remove Pd/C. The mixture was concentrated in vacuo, followed by purification by reverse phase silica gel column chromatography (eluted with $CH_3CN/H_2O$ with 0.1% TFA=5/95~95/5) to give tert-butyl (4-((3-(cyclopropylmethyl)isoxazol-5-yl)methyl)tetrahydrofuran-3-yl)carbamate (**S72**, 340 mg, 82% yield) as a light-yellow oil.

LC-MS (ESI, *m/z*): $[M+H-C_4H_8]^+$ = 267.1

**Procedure for preparation of 4-((3-(cyclopropylmethyl)isoxazol-5-yl)methyl)tetrahydrofuran-3-amine (S73)**

To a solution of tert-butyl (4-((3-(cyclopropylmethyl)isoxazol-5-yl)methyl)tetrahydrofuran-3-yl)carbamate (340 mg, 1.05 mmol, 1.0 eq) in DCM (5 mL) was added TFA (2 mL). The resulting mixture was stirred at room temperature for 1.0 h. The reaction mixture was concentrated under vacuum to give 4-((3-(cyclopropylmethyl)isoxazol-5-yl)methyl)tetrahydrofuran-3-amine (**S73**, crude, TFA salt) as a black oil.

LC-MS (ESI, *m/z*): $[M+H]^+$ = 223.1

**Procedure for preparation of N-(4-((3-(cyclopropylmethyl)isoxazol-5-yl)methyl)tetrahydrofuran-3-yl)-3-(3-fluorophenyl)-1H-pyrazole-5-carboxamide (1554-36)**

To a mixture of 4-((3-(cyclopropylmethyl)isoxazol-5-yl)methyl)tetrahydrofuran-3-amine (**S73**, crude, TFA salt, 1.05 mmol, 1.0 eq.) and 3-(3-fluorophenyl)-1H-pyrazole-5-carboxylic acid (**S13**, 260 mg, 1.26 mmol, 1.2 eq.) in DMF (3 mL) was added DIPEA (0.5 mL) and HATU (479 mg, 1.26 mmol, 1.2 eq). The resulting mixture was stirred at room temperature for 2 h. The reaction mixture was diluted with water (20 mL), extracted with ethyl acetate (20 mL×3). The combined organic phases were dried over anhydrous $Na_2SO_4$ and concentrated under vacuum. The residue was purified by silica gel column chromatography (eluted with $CH_3CN/H_2O$ with 0.1% TFA = 5/95 ~ 95/5) to give N-(4-((3-(cyclopropylmethyl)isoxazol-5-yl)methyl)tetrahydrofuran-3-yl)-3-(3-fluorophenyl)-1H-pyrazole-5-carboxamide (**1554-36**, 190 mg, 44% yield) as a white solid.

LC-MS (ESI, *m/z*): $[M+H]^+$ = 411.2.

**Procedure for the chiral separation of N-(4-((3-(cyclopropylmethyl)isoxazol-5-yl)methyl)tetrahydrofuran-3-yl)-3-(3-fluorophenyl)-1H-pyrazole-5-carboxamide (1554-36)**

N-(4-((3-(cyclopropylmethyl)isoxazol-5-yl)methyl)tetrahydrofuran-3-yl)-3-(3-fluorophenyl)-1H-pyrazole-5-carboxamide (**1554-36**, 190 mg, 0.46 mmol) was separated by chiral preparation (for method, see 1554-06, eluent $CO_2$/EtOH[0.5%NH3(7M in MeOH)]=45/55) to give N-(4-((3-(cyclopropylmethyl)isoxazol-5-yl)methyl)tetrahydrofuran-3-yl)-3-(3-fluorophenyl)-1H-pyrazole-5-carboxamide (24 mg, 3R,4S *or* 3S,4R, **1554-36-cis1**), N-(4-((3-(cyclopropylmethyl)isoxazol-5-yl)methyl)tetrahydrofuran-3-yl)-3-(3-fluorophenyl)-1H-pyrazole-5-carboxamide (41 mg, 3R,4S *or* 3S,4R, **1554-36-cis2, 9**), N-(4-((3-(cyclopropylmethyl)isoxazol-5-yl)methyl)tetrahydrofuran-3-yl)-3-(3-fluorophenyl)-1H-pyrazole-5-carboxamide (15 mg, 3R,4R *or* 3S,4S, **1554-36-trans1**)

**1554-36-cis1**: $^1$H NMR (400 MHz, Methanol-d4) δ 7.54 (s, 1H), 7.48 (d, J = 8.8 Hz, 2H), 7.11 (d, J = 14.8 Hz, 2H), 6.15 (s, 1H), 4.83 (s, 1H), 4.10 (dd, J = 9.3, 6.2 Hz, 1H), 4.03 (dd, J = 8.7, 7.3 Hz, 1H), 3.82 (dd, J = 9.3, 3.9 Hz, 1H), 3.70 (t, J = 8.3 Hz, 1H), 3.03 (dd, J = 14.9, 6.0 Hz, 1H), 2.90 (dt, J = 14.2, 7.4 Hz, 1H), 2.81 (dd, J = 14.9, 8.7 Hz, 1H), 2.46 (d, J = 7.1 Hz, 2H), 0.96 – 0.87 (m, 1H), 0.54 – 0.43 (m, 2H), 0.17 (q, J = 4.8 Hz, 2H).

LC-MS (ESI, *m/z*): [M + H]+ = 411.2.

**1554-36-cis2 (9)**: $^1$H NMR (400 MHz, Methanol-d4) δ 7.51 (dd, J = 24.1, 8.3 Hz, 3H), 7.29 – 6.97 (m, 2H), 6.15 (s, 1H), 4.84 (d, J = 6.5 Hz, 1H), 4.10 (dd, J = 9.3, 6.1 Hz, 1H), 4.04 (dd, J = 8.7, 7.3 Hz, 1H), 3.82 (dd, J = 9.3, 3.9 Hz, 1H), 3.70 (t, J = 8.3 Hz, 1H), 3.03 (dd, J = 14.9, 6.0 Hz, 1H), 2.99 – 2.87 (m, 1H), 2.81 (dd, J = 14.9, 8.7 Hz, 1H), 2.46 (d, J = 7.1 Hz, 2H), 1.00 – 0.85 (m, 1H), 0.49 (dt, J = 8.2, 3.0 Hz, 2H), 0.17 (q, J = 5.0 Hz, 2H).

LC-MS (ESI, *m/z*): [M + H]+ = 411.2.

**1554-36-trans1**: $^1$H NMR (400 MHz, Methanol-d4) δ 7.32 (t, J = 13.3 Hz, 3H), 6.96 (s, 1H), 6.89 (s, 1H), 6.07 (s, 1H), 4.42 (s, 3H), 4.27 (q, J = 6.1 Hz, 1H), 3.95 (ddd, J = 11.8, 9.1, 7.1 Hz, 2H), 3.53 (dd, J = 9.1, 5.6 Hz, 1H), 3.44 (dd, J = 9.0, 6.8 Hz, 1H), 2.90 (dd, J = 15.5, 7.1 Hz, 1H), 2.78 (dd, J = 15.4, 7.8 Hz, 1H), 2.62 – 2.49 (m, 1H), 2.28 (d, J = 7.0 Hz, 2H), 0.77 – 0.69 (m, 1H), 0.32 (t, J = 6.5 Hz, 2H), -0.07 (s, 2H).

LC-MS (ESI, *m/z*): [M + H]+ = 411.2.

#### $^{1}$H NMR spectra and chiral HPLC for 1554 series

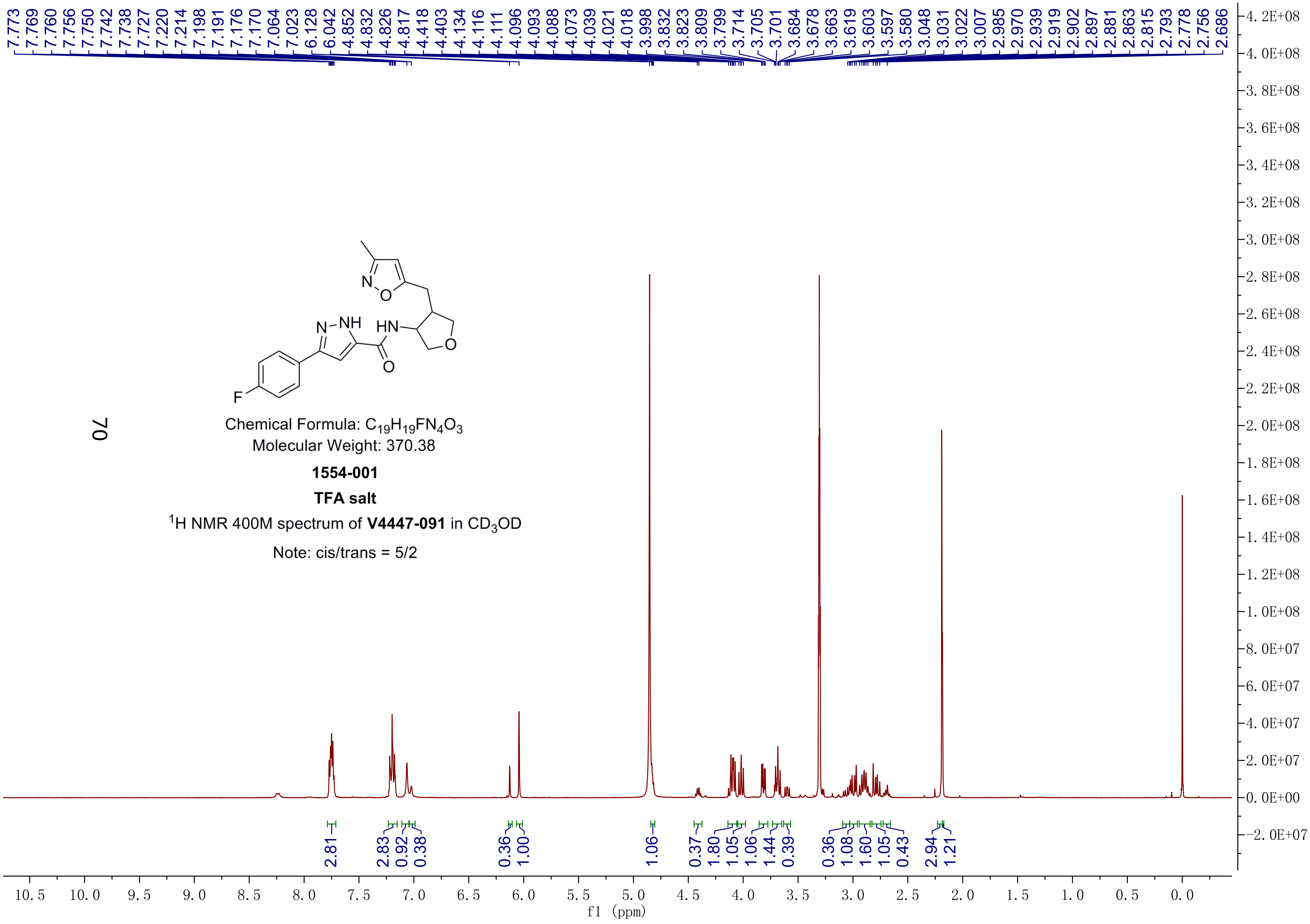


Chemical Formula: $C_{19}H_{19}FN_4O_3$
Molecular Weight: 370.38

**1554-001**

**TFA salt**

$^1$H NMR 400M spectrum of **V4447-091** in $CD_3OD$

Note: cis/trans = 5/2

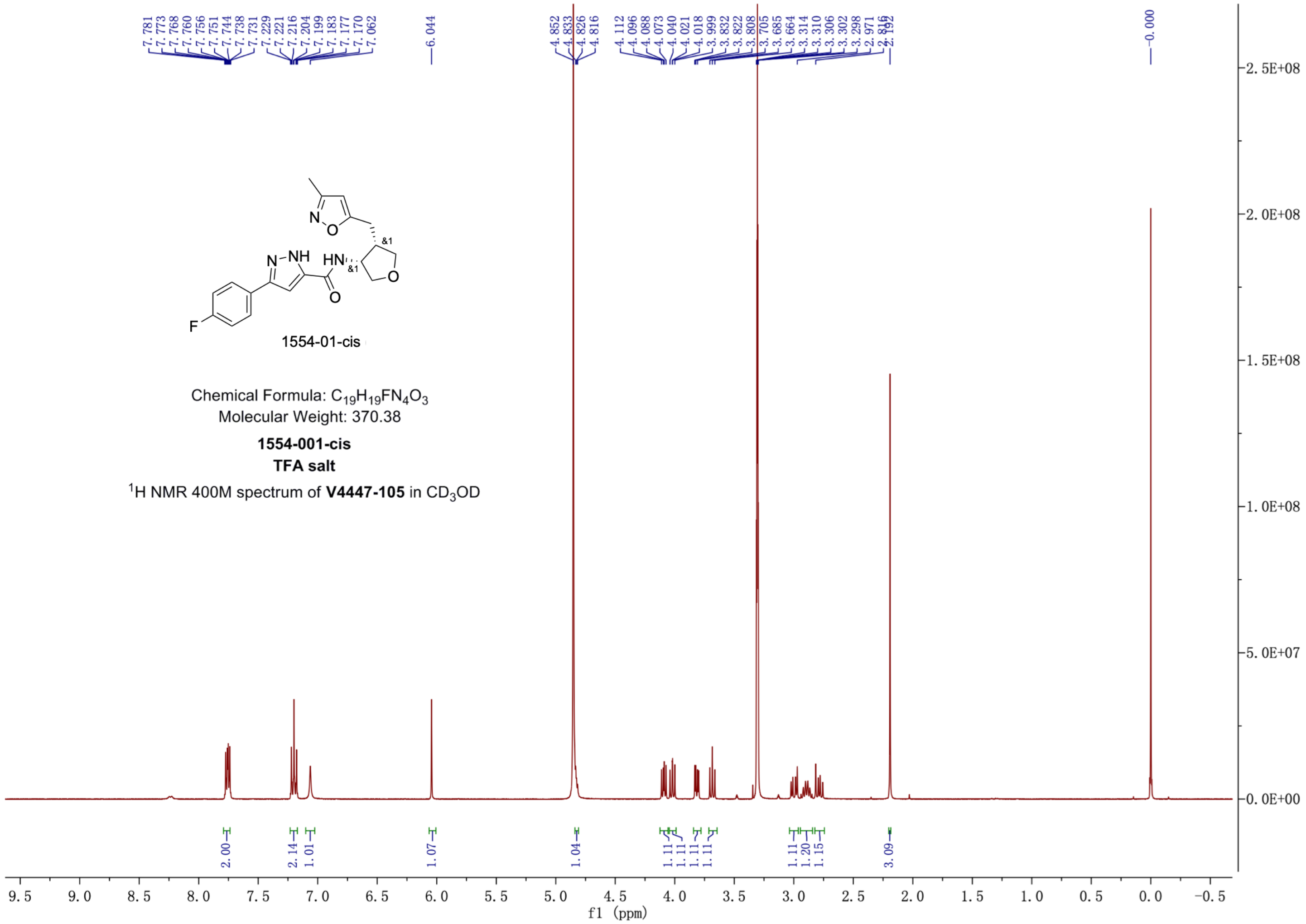


1554-01-cis

Chemical Formula: $C_{19}H_{19}FN_4O_3$
Molecular Weight: 370.38

**1554-001-cis**
**TFA salt**

$^1$H NMR 400M spectrum of **V4447-105** in $CD_3OD$

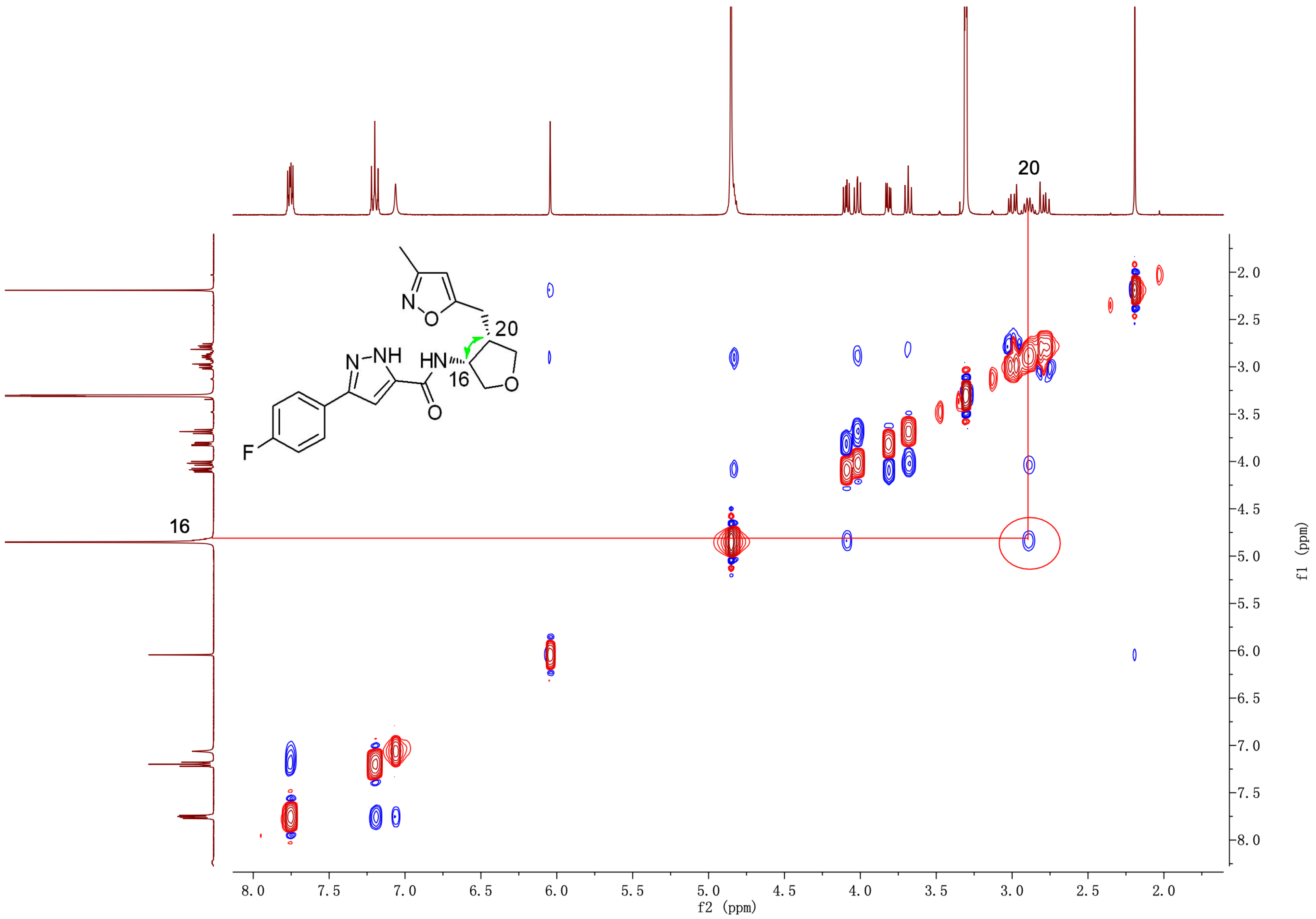

20
16
HN
N–NH
N
O
F
f2 (ppm)
f1 (ppm)

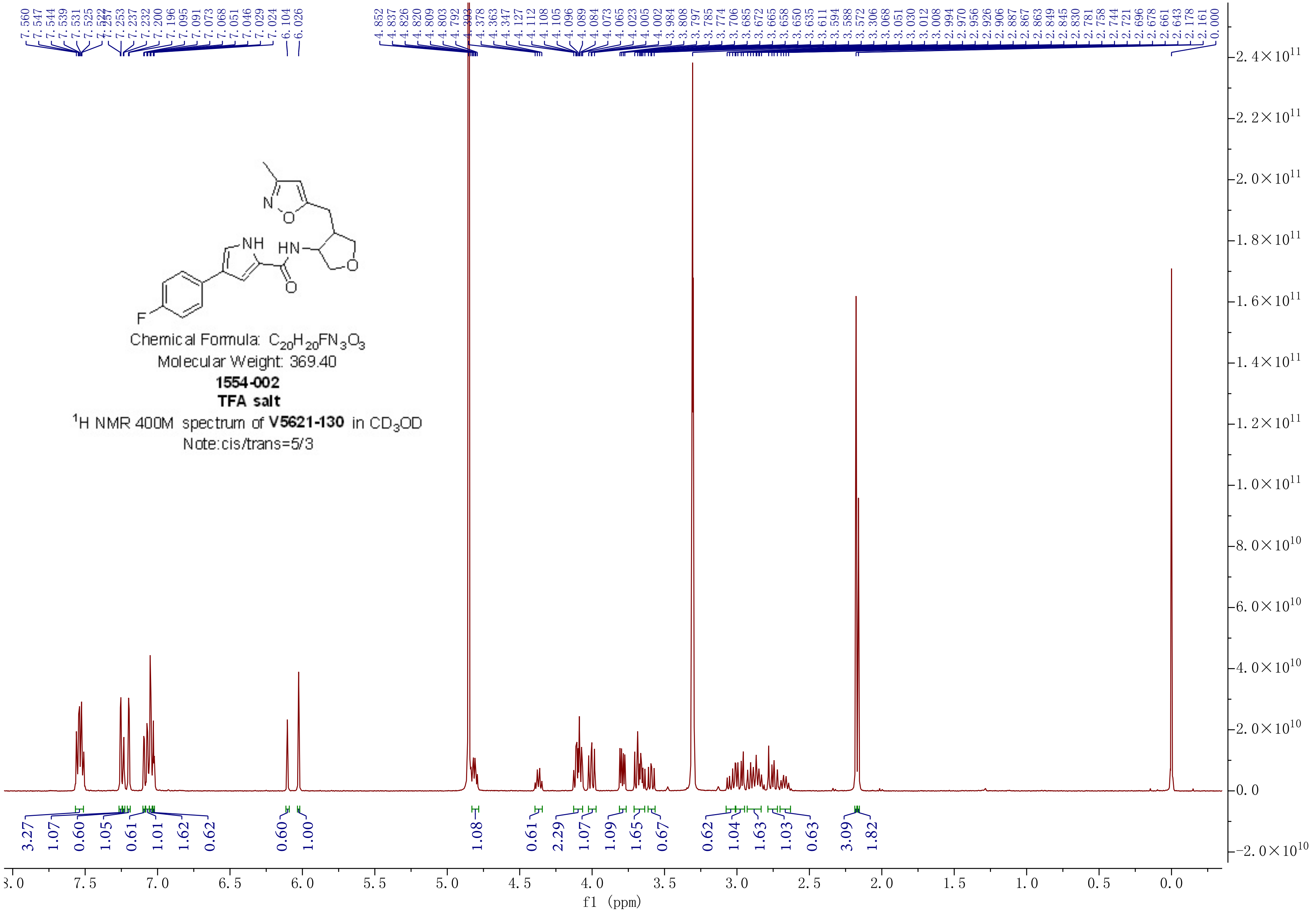

Chemical Formula: $C_{20}H_{20}FN_3O_3$
Molecular Weight: 369.40
**1554-002**
**TFA salt**
$^1$H NMR 400M spectrum of **V5621-130** in $CD_3OD$
Note:cis/trans=5/3
f1 (ppm)

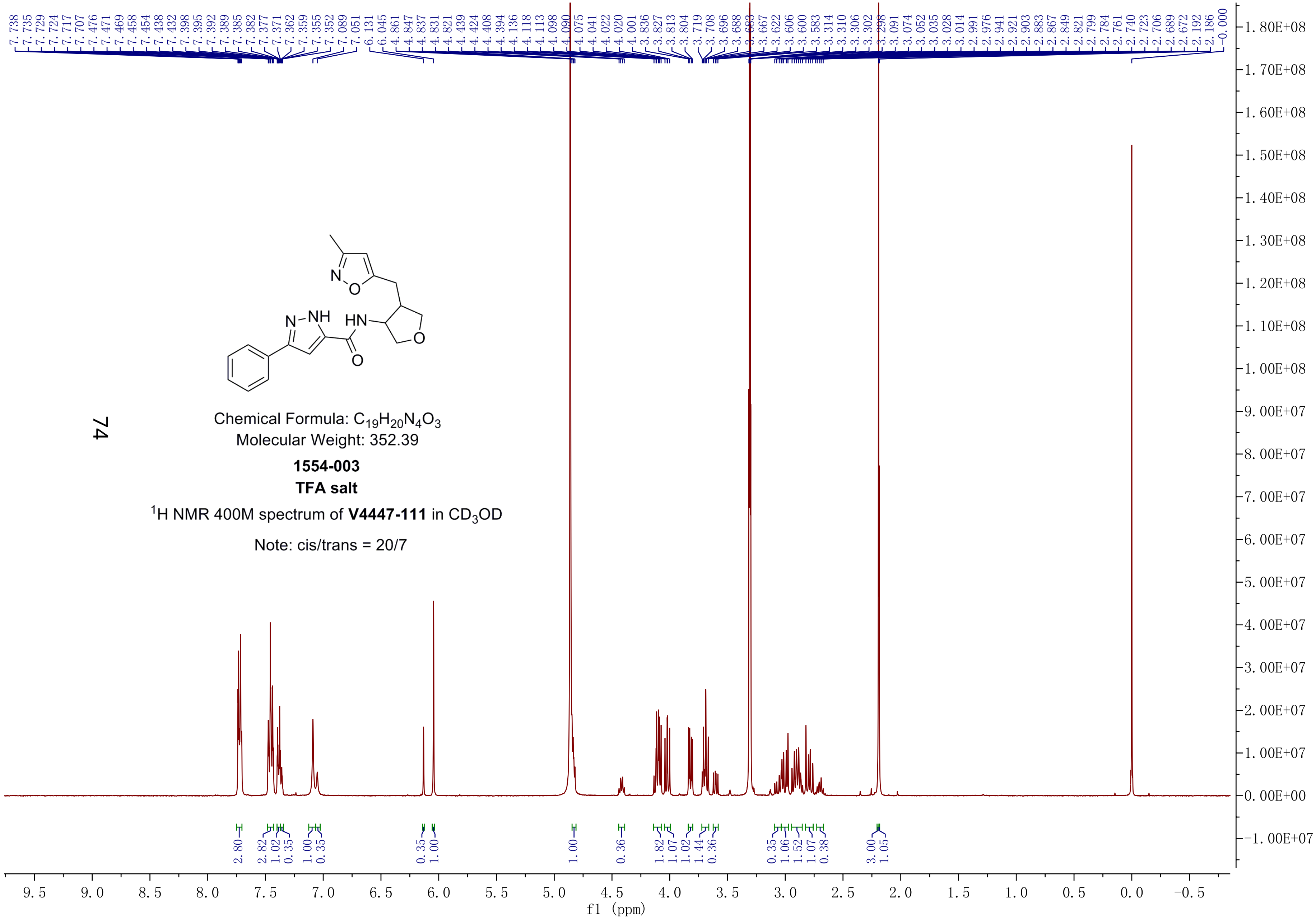


Chemical Formula: $C_{19}H_{20}N_4O_3$
Molecular Weight: 352.39

**1554-003**
**TFA salt**

$^{1}$H NMR 400M spectrum of **V4447-111** in $CD_3OD$

Note: cis/trans = 20/7

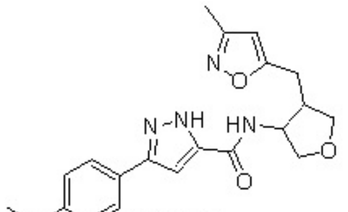

**1554-004**

Chemical Formula: $C_{20}H_{22}N_4O_4$
Molecular Weight: 382.41
$^1$H NMR 400M spectrum of V5618-092 in MeOH-$d_4$
cis:trans= 10:7

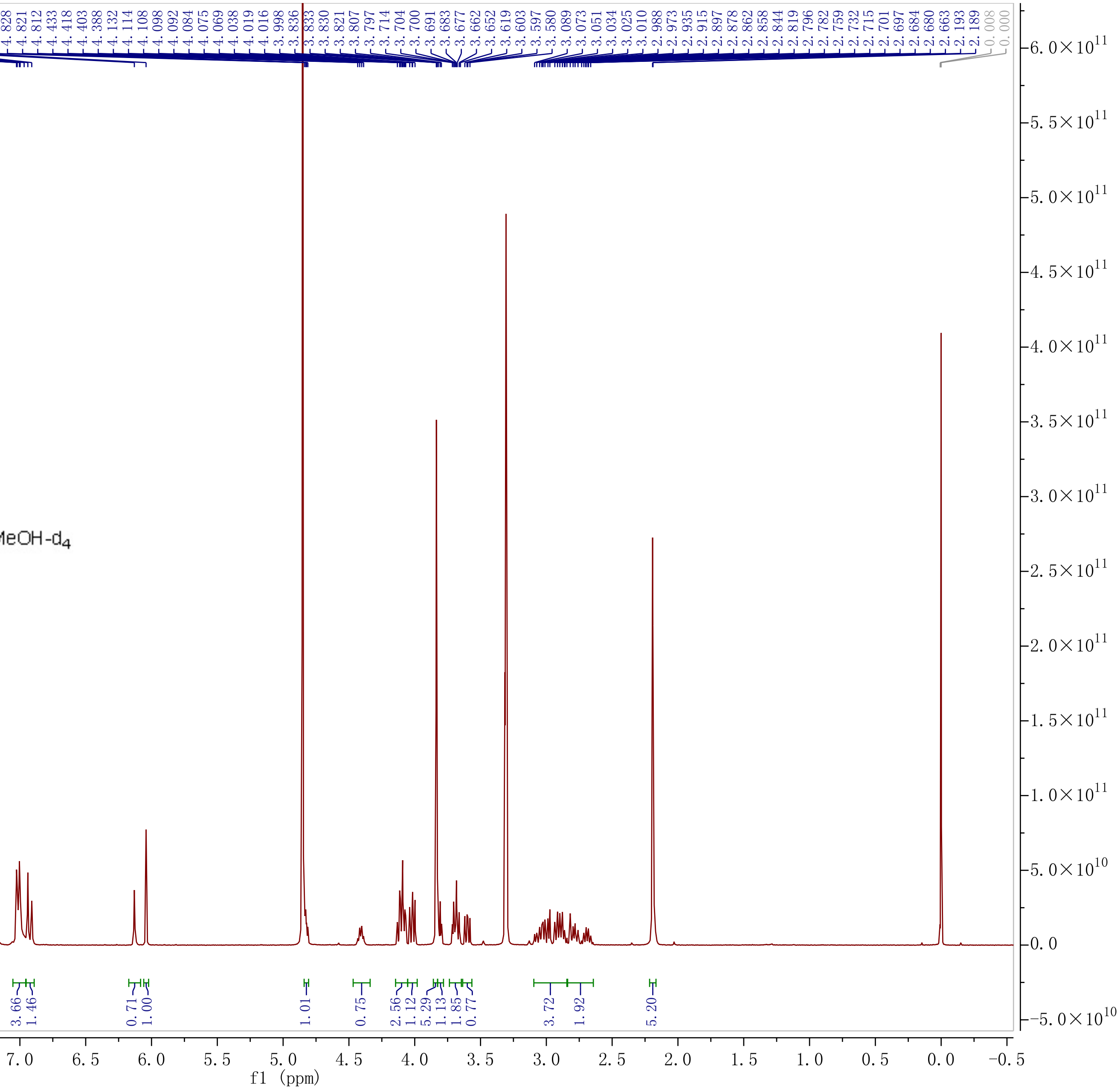

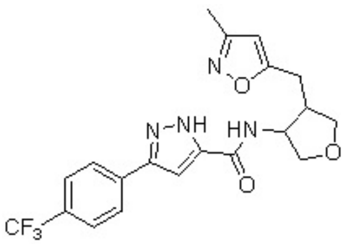

Chemical Formula: $C_{20}H_{19}F_3N_4O_3$
Molecular Weight: 420.39
**1554-005**
$^1$H NMR 400M spectrum of **V5621-135** in DMSO-$d_6$
Note:cis/trans=4/3

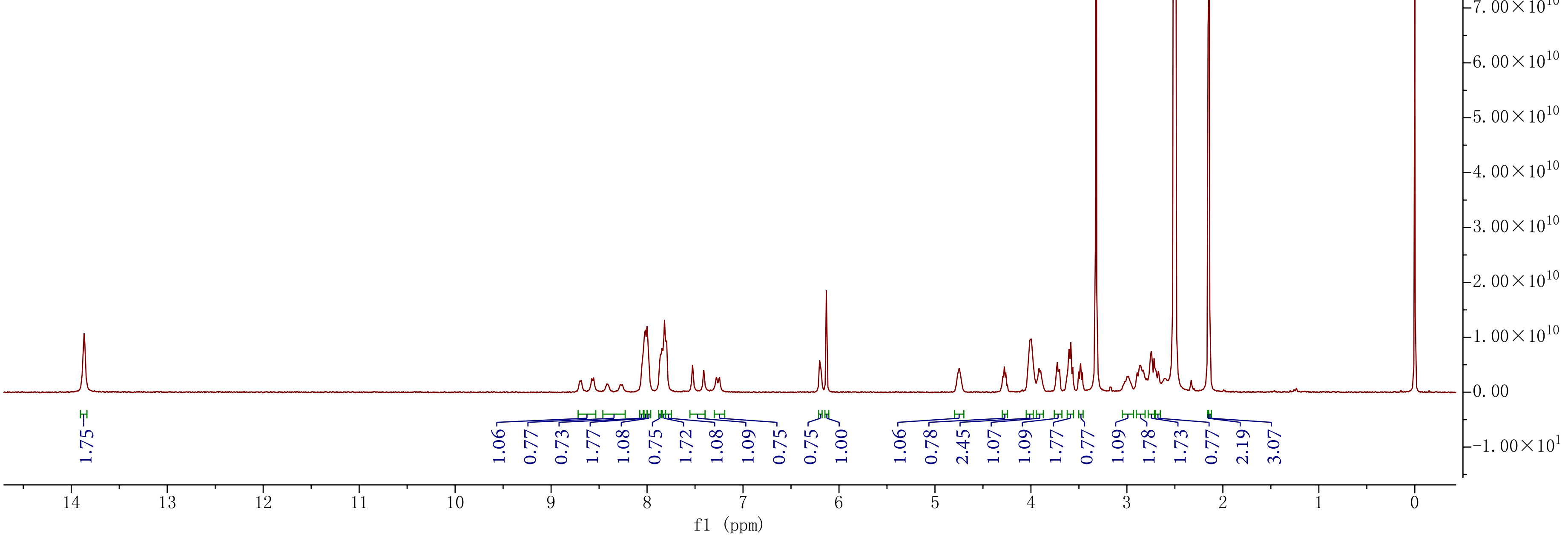

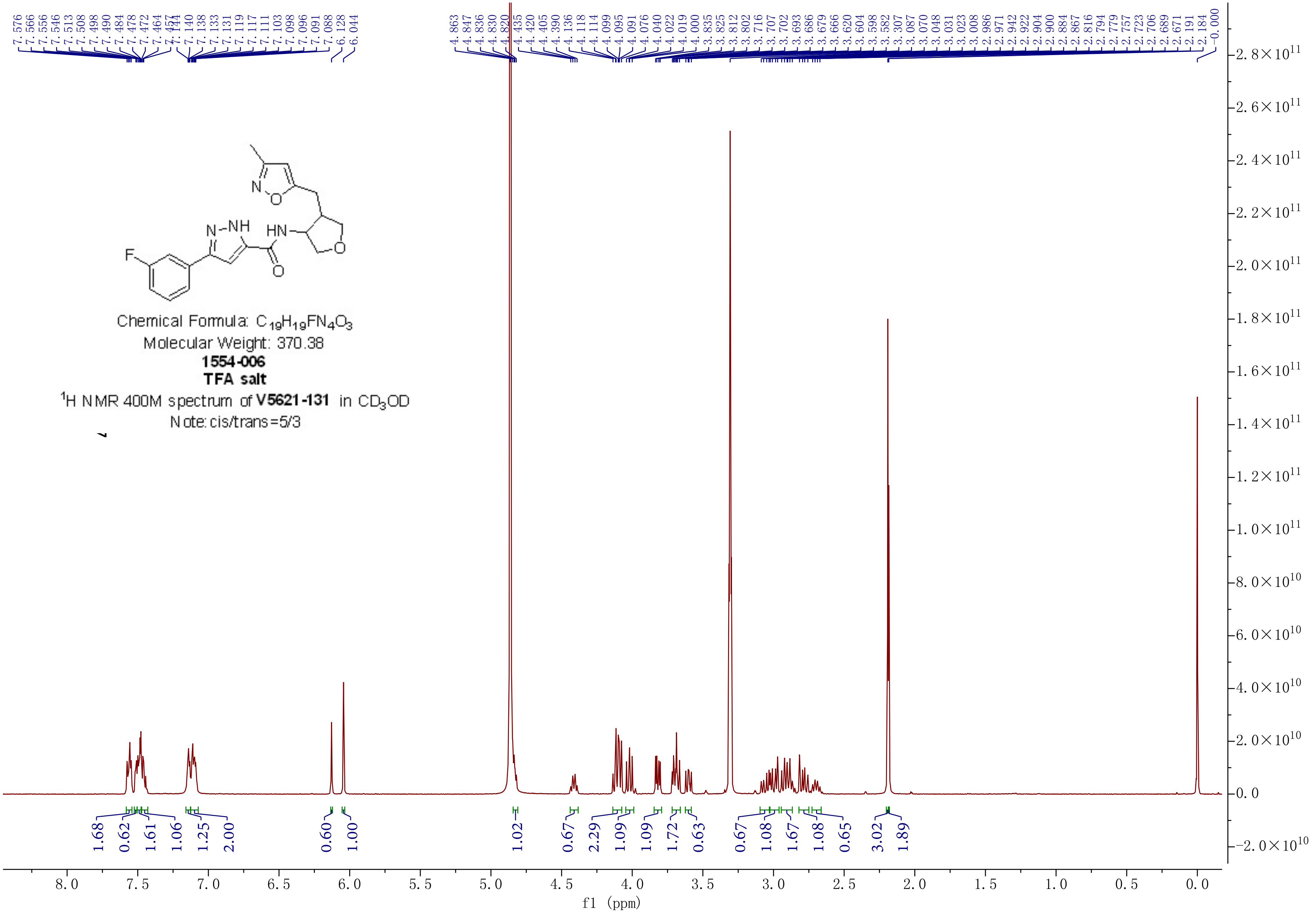

Chemical Formula: $C_{19}H_{19}FN_4O_3$
Molecular Weight: 370.38
**1554-006**
**TFA salt**
$^1$H NMR 400M spectrum of **V5621-131** in $CD_3OD$
Note: cis/trans=5/3

**1554-006-cis**
**TFA salt**

$^{1}$H NMR 400M spectrum of **V5601-105** in $CD_3OD$

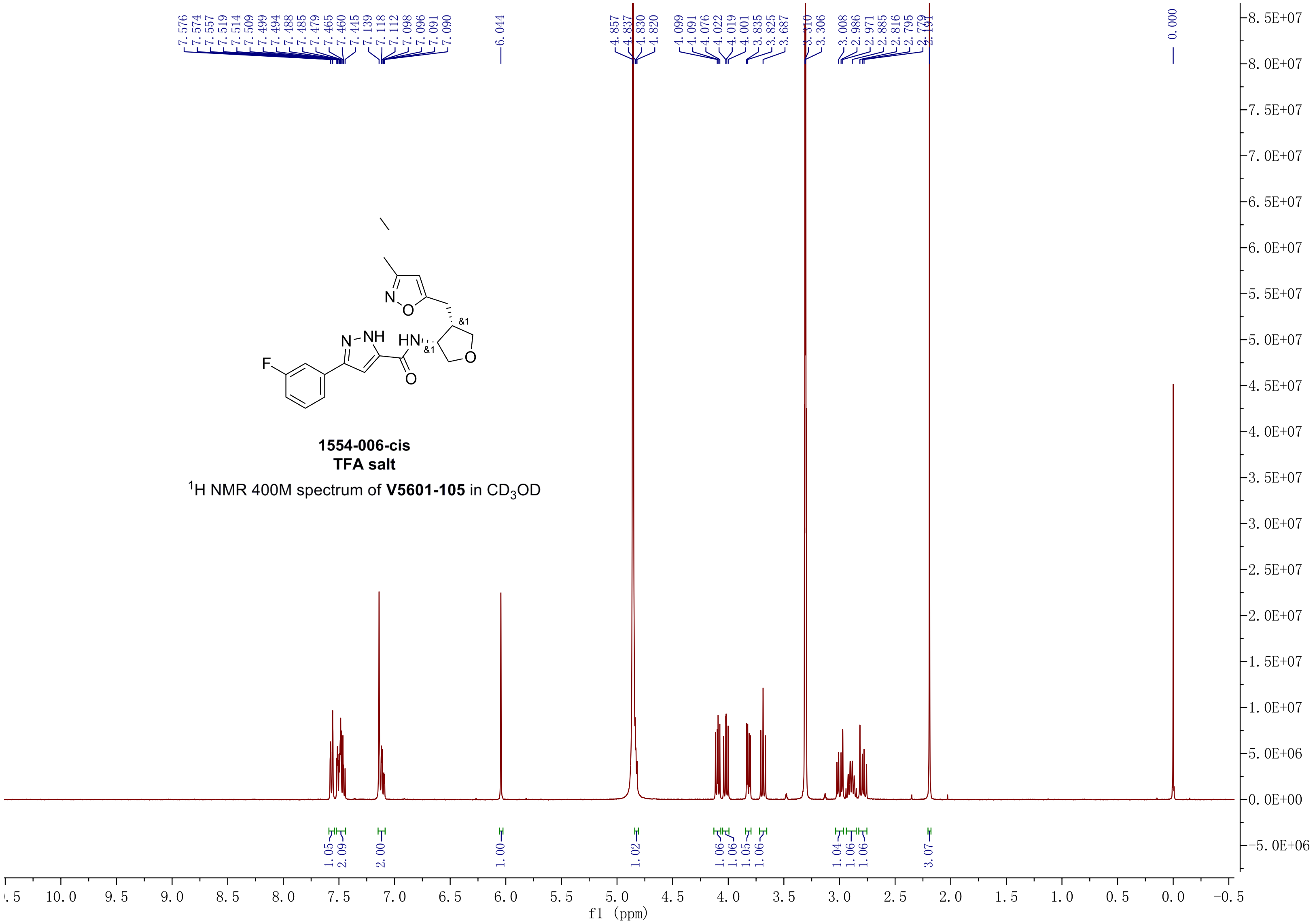

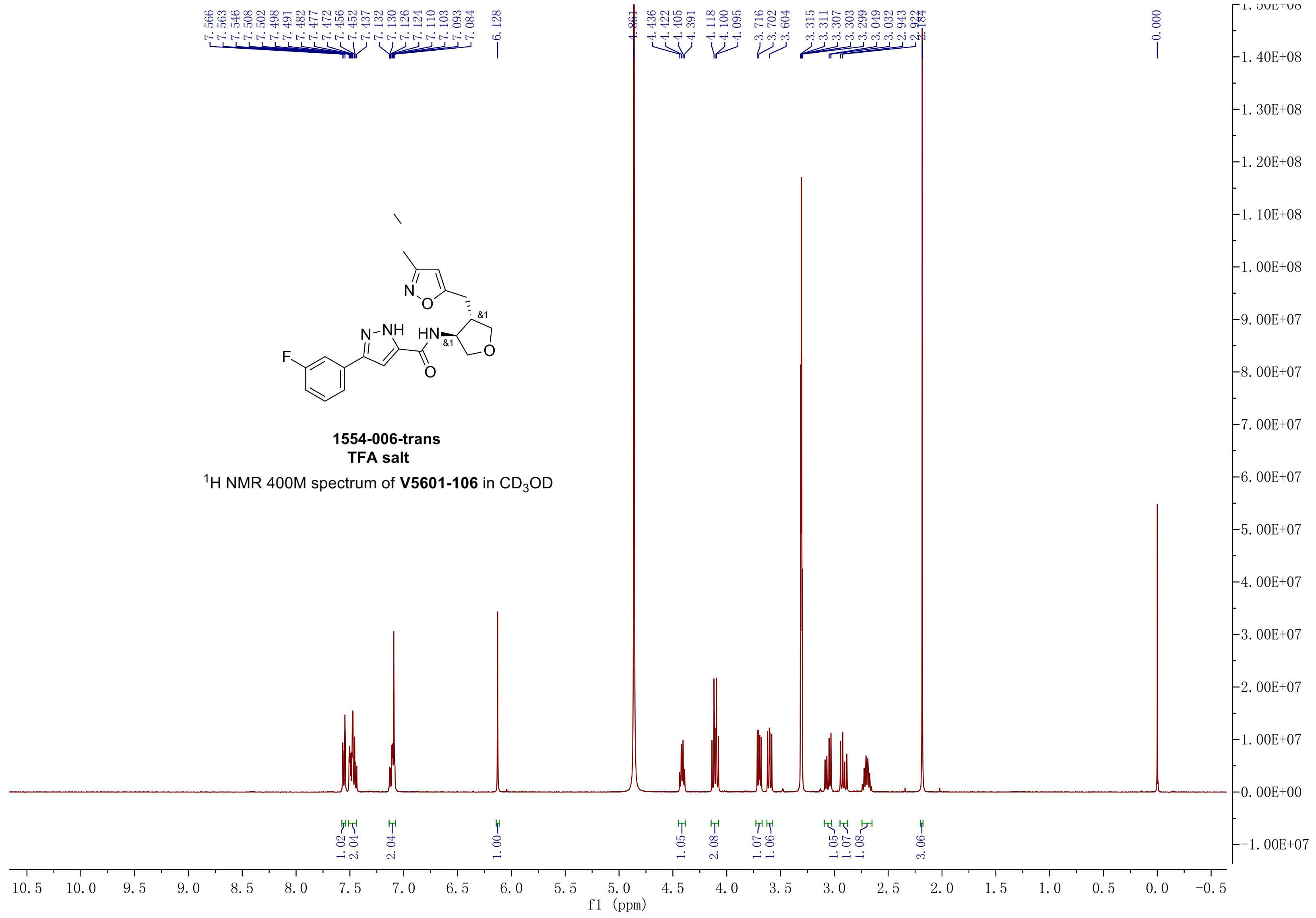

F
N
NH
HN
&1
O
O
N
O
&1
1554-006-trans
TFA salt
$^1$H NMR 400M spectrum of **V5601-106** in $CD_3OD$
7.566 7.563 7.546 7.508 7.502 7.498 7.491 7.482 7.477 7.472 7.456 7.452 7.437 7.132 7.130 7.126 7.124 7.110 7.103 7.093 7.084
6.128
4.861
4.436 4.422 4.405 4.391
4.118 4.100 4.095
3.716 3.702 3.604
3.315 3.311 3.307 3.303 3.299 3.049 3.032 2.943 2.922 2.184
0.000
1.02 2.04 2.04 1.00 1.05 2.08 1.07 1.06 1.05 1.07 1.08 3.06
f1 (ppm)

1554-06-3S,4R
1554-06-cis1

Molecular Weight: 370.38

1H NMR 400M spectrum of V5899-007-P1 in MeOH-$d_4$

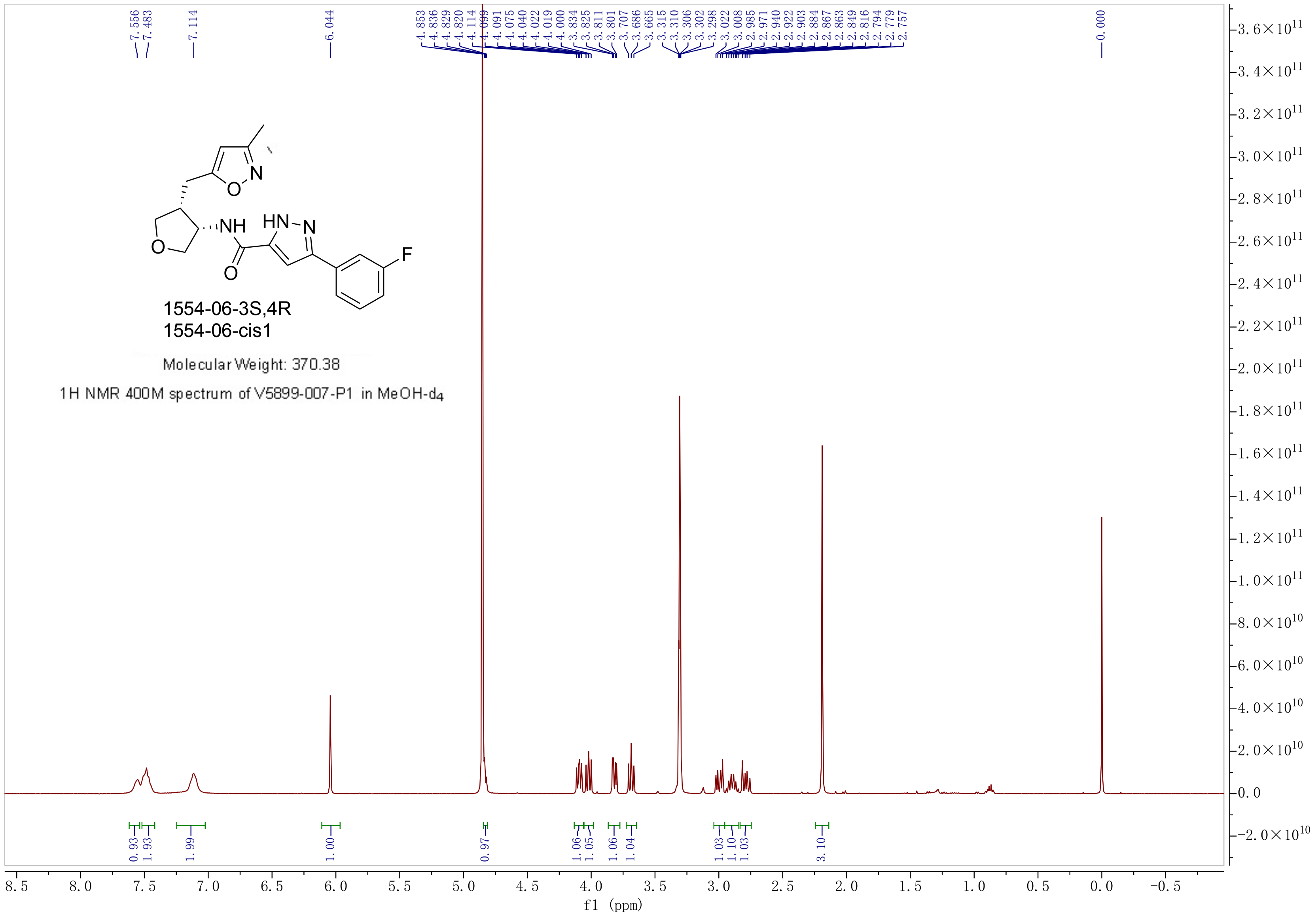

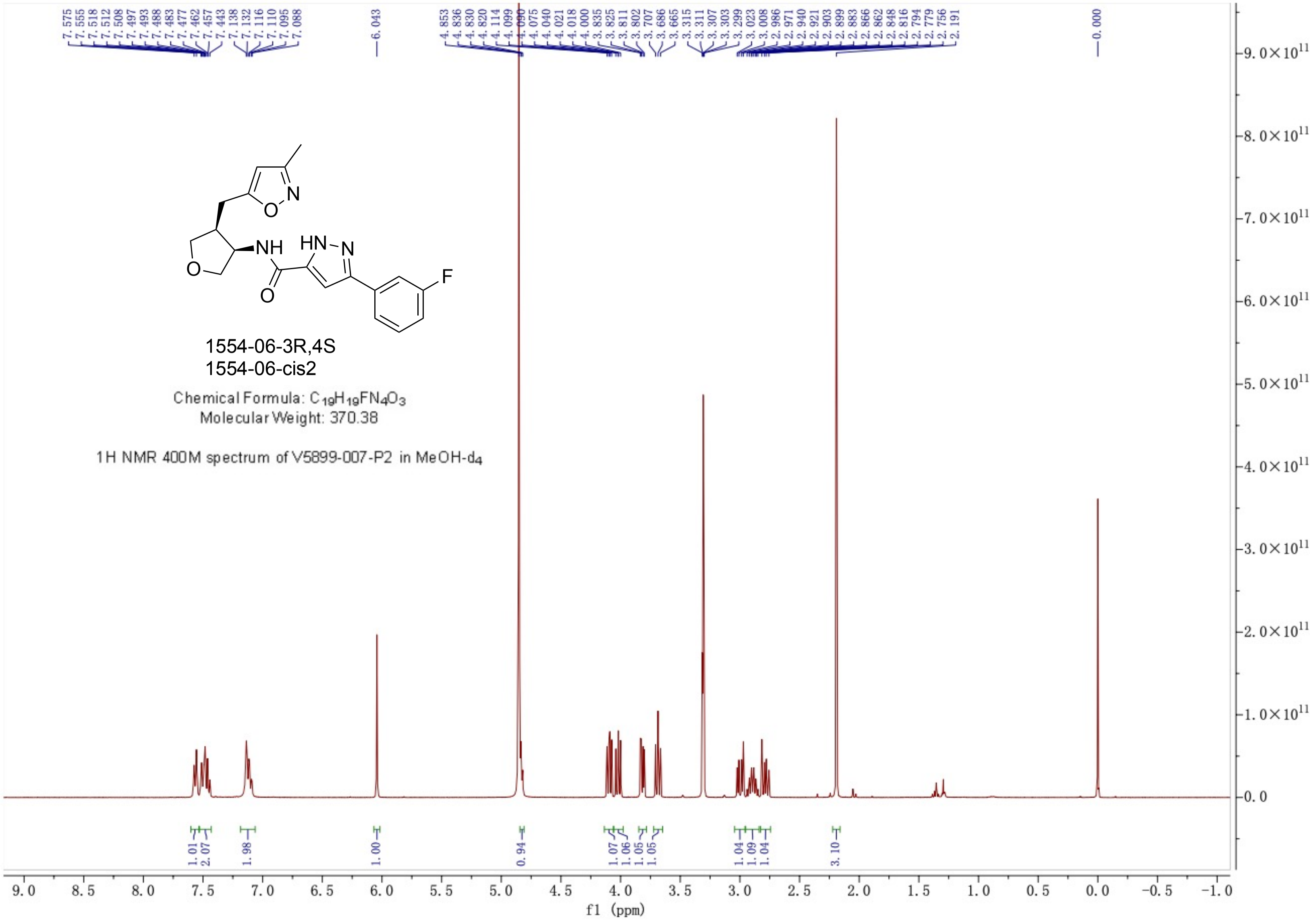
7.575 7.555 7.518 7.512 7.508 7.497 7.493 7.488 7.483 7.477 7.462 7.457 7.443 7.138 7.132 7.116 7.110 7.095 7.088
6.043
4.853 4.836 4.830 4.820 4.114 4.099 4.090 4.075 4.040 4.021 4.018 4.000 3.835 3.825 3.811 3.802 3.707 3.686 3.665 3.315 3.311 3.307 3.303 3.299 3.023 3.008 2.986 2.971 2.940 2.921 2.903 2.899 2.883 2.866 2.862 2.848 2.816 2.794 2.779 2.756 2.191
0.000
1554-06-3R,4S
1554-06-cis2
Chemical Formula: $C_{19}H_{19}FN_4O_3$
Molecular Weight: 370.38
1H NMR 400M spectrum of V5899-007-P2 in MeOH-$d_4$
1.01 2.07 1.98 1.00 0.94 1.07 1.06 1.05 1.05 1.04 1.09 1.04 3.10
f1 (ppm)

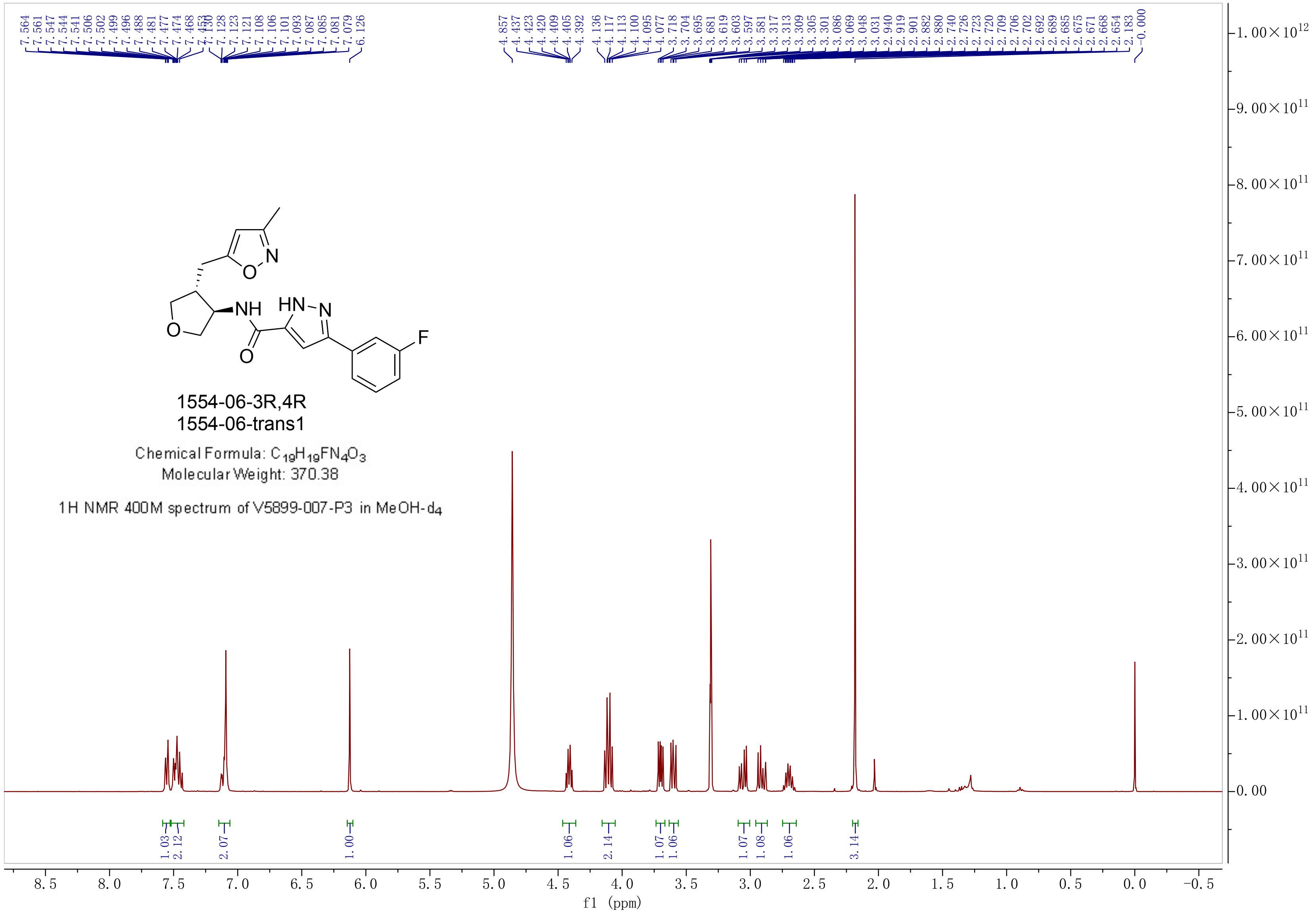
1554-06-3R,4R
1554-06-trans1
Chemical Formula: $C_{19}H_{19}FN_4O_3$
Molecular Weight: 370.38
1H NMR 400M spectrum of V5899-007-P3 in MeOH-$d_4$
f1 (ppm)

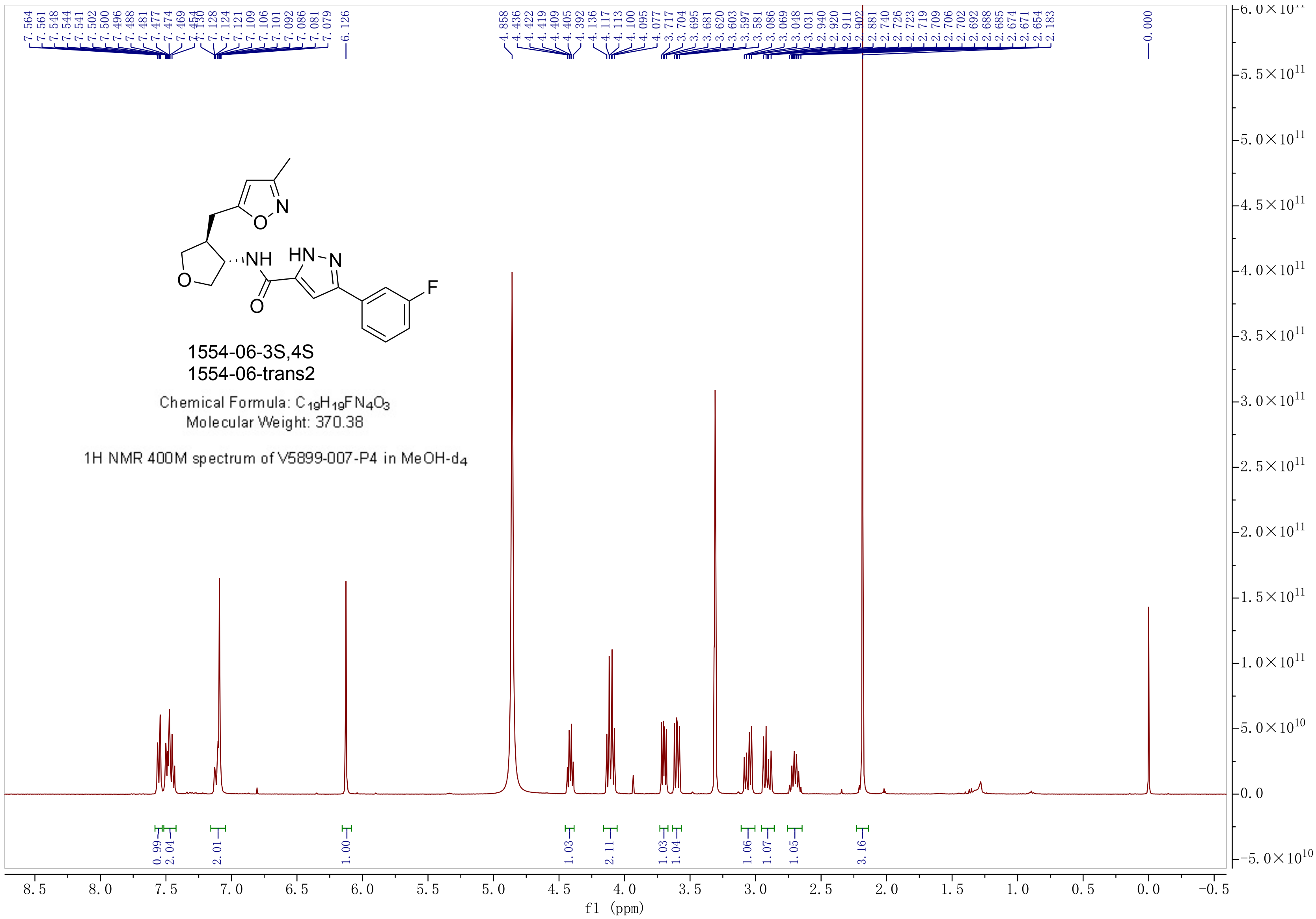


1554-06-3S,4S
1554-06-trans2

Chemical Formula: $C_{19}H_{19}FN_4O_3$
Molecular Weight: 370.38

1H NMR 400M spectrum of V5899-007-P4 in MeOH-$d_4$

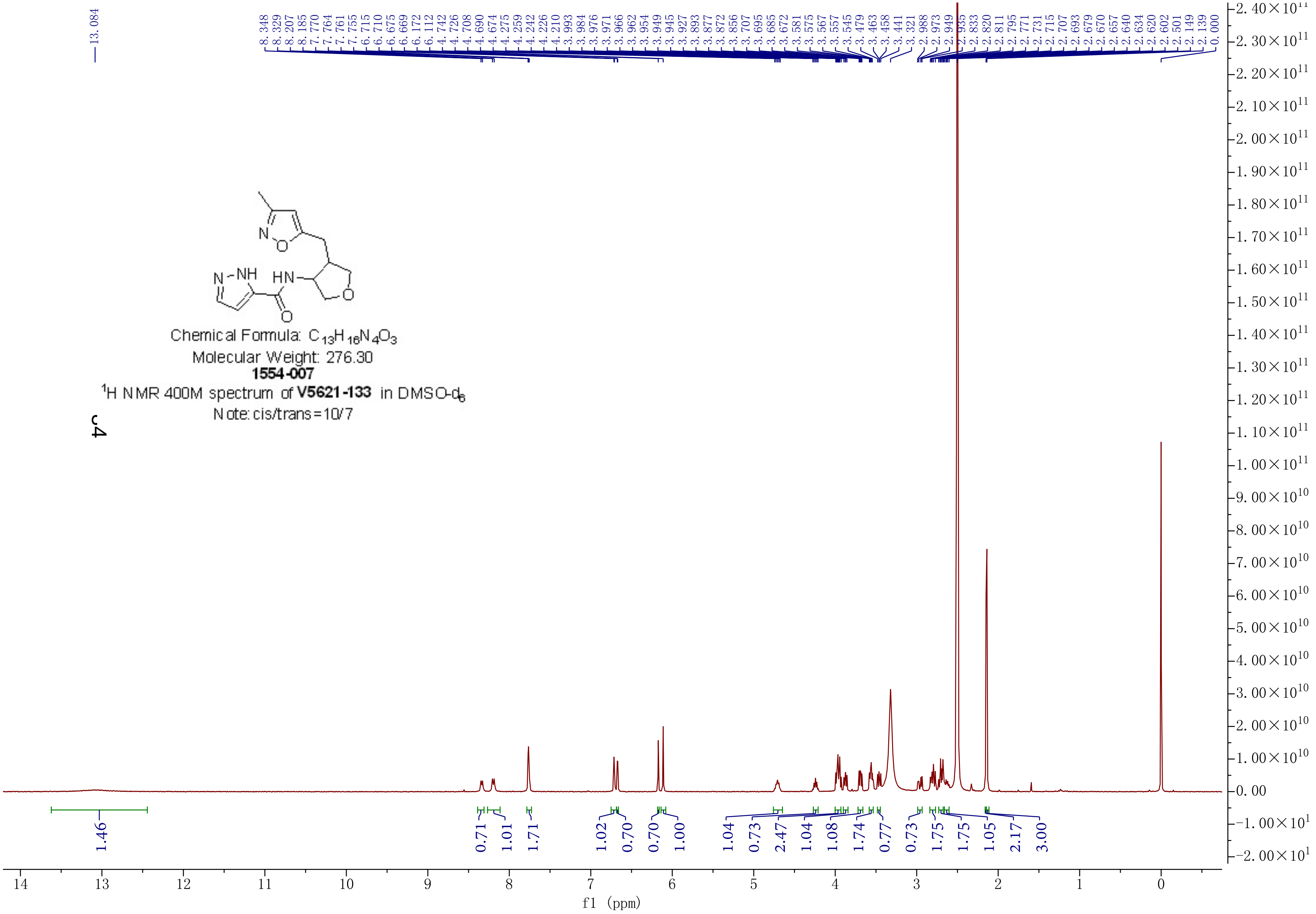


Chemical Formula: $C_{13}H_{16}N_4O_3$

Molecular Weight: 276.30

**1554-007**

$^1$H NMR 400M spectrum of **V5621-133** in DMSO-$d_6$

Note: cis/trans=10/7

[illegible]4

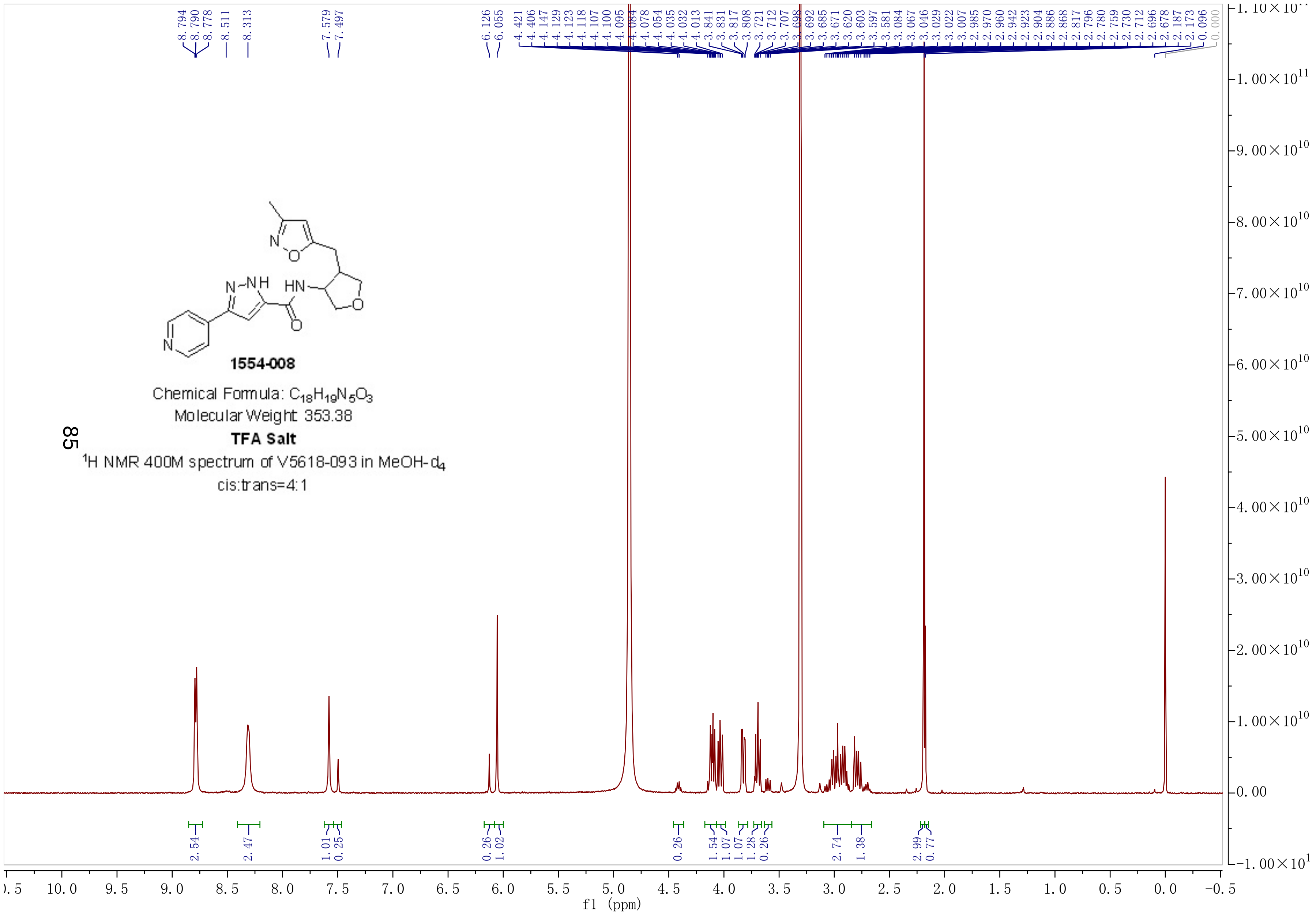


**1554-008**

Chemical Formula: $C_{18}H_{19}N_5O_3$

Molecular Weight: 353.38

**TFA Salt**

$^1H$ NMR 400M spectrum of V5618-093 in MeOH-$d_4$

cis:trans=4:1

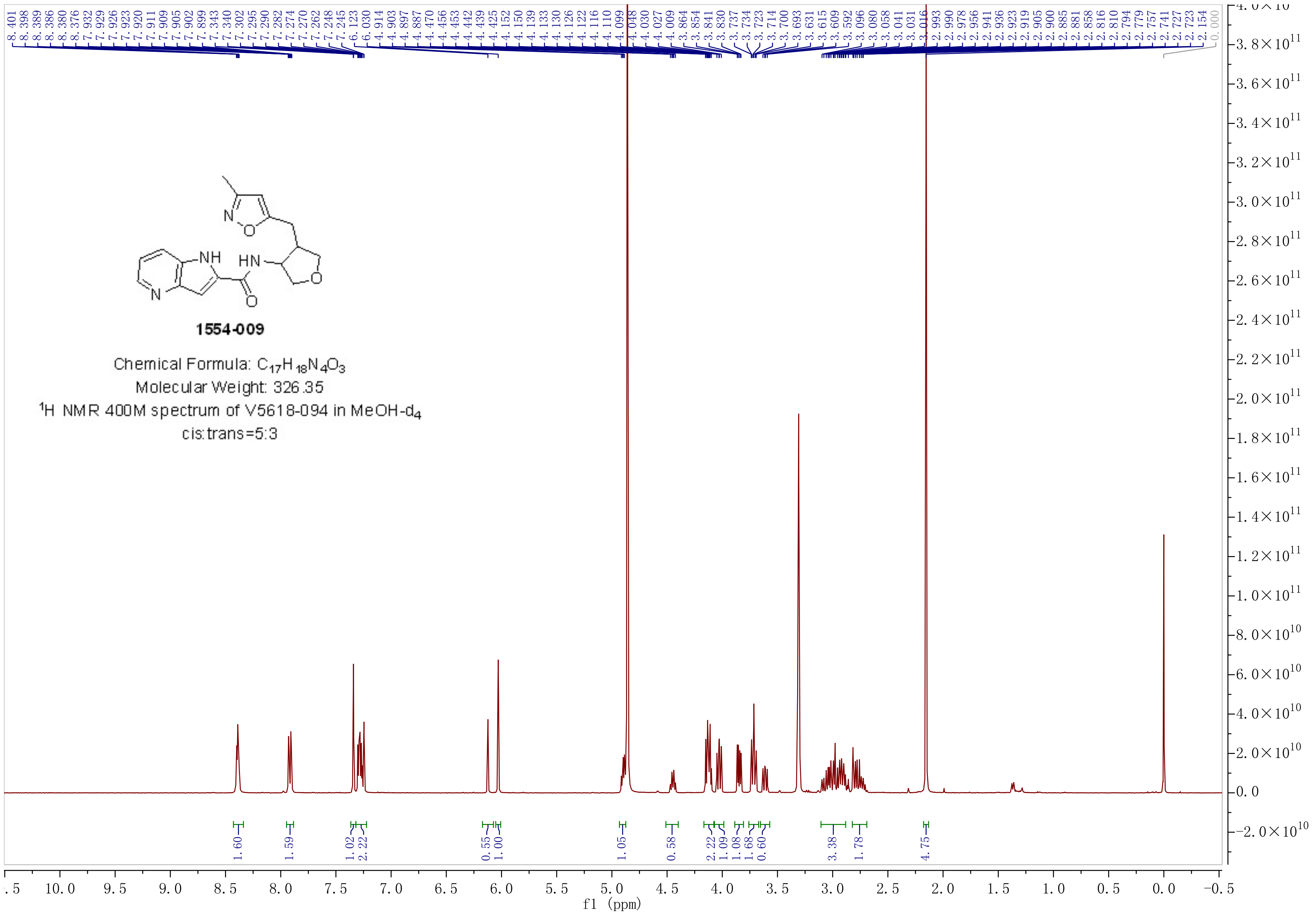

1554-009
Chemical Formula: $C_{17}H_{18}N_4O_3$
Molecular Weight: 326.35
$^1$H NMR 400M spectrum of V5618-094 in MeOH-$d_4$
cis:trans=5:3
8.401 8.398 8.389 8.386 8.380 8.376 7.932 7.929 7.926 7.923 7.920 7.911 7.909 7.905 7.902 7.899 7.343 7.340 7.302 7.295 7.290 7.282 7.274 7.270 7.262 7.248 7.245 6.123 6.030 4.914 4.903 4.897 4.887 4.470 4.456 4.453 4.442 4.439 4.425 4.152 4.150 4.139 4.133 4.130 4.126 4.122 4.116 4.110 4.099 4.048 4.030 4.027 4.009 3.864 3.854 3.841 3.830 3.737 3.734 3.723 3.714 3.700 3.693 3.631 3.615 3.609 3.592 3.096 3.080 3.058 3.041 3.031 3.016 2.993 2.990 2.978 2.956 2.941 2.936 2.923 2.919 2.905 2.900 2.885 2.881 2.858 2.816 2.810 2.794 2.779 2.757 2.741 2.727 2.723 2.154 0.000
1.60 1.59 1.02 2.22 0.55 1.00 1.05 0.58 2.22 1.09 1.08 1.68 0.60 3.38 1.78 4.75
f1 (ppm)

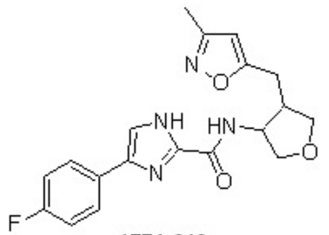


**1554-010**

Chemical Formula: $C_{19}H_{19}FN_4O_3$

Molecular Weight: 370.38

**TFA Salt**

$^1$H NMR 400M spectrum of V5618-095 in MeOH-$d_4$

cis:trans=4:3

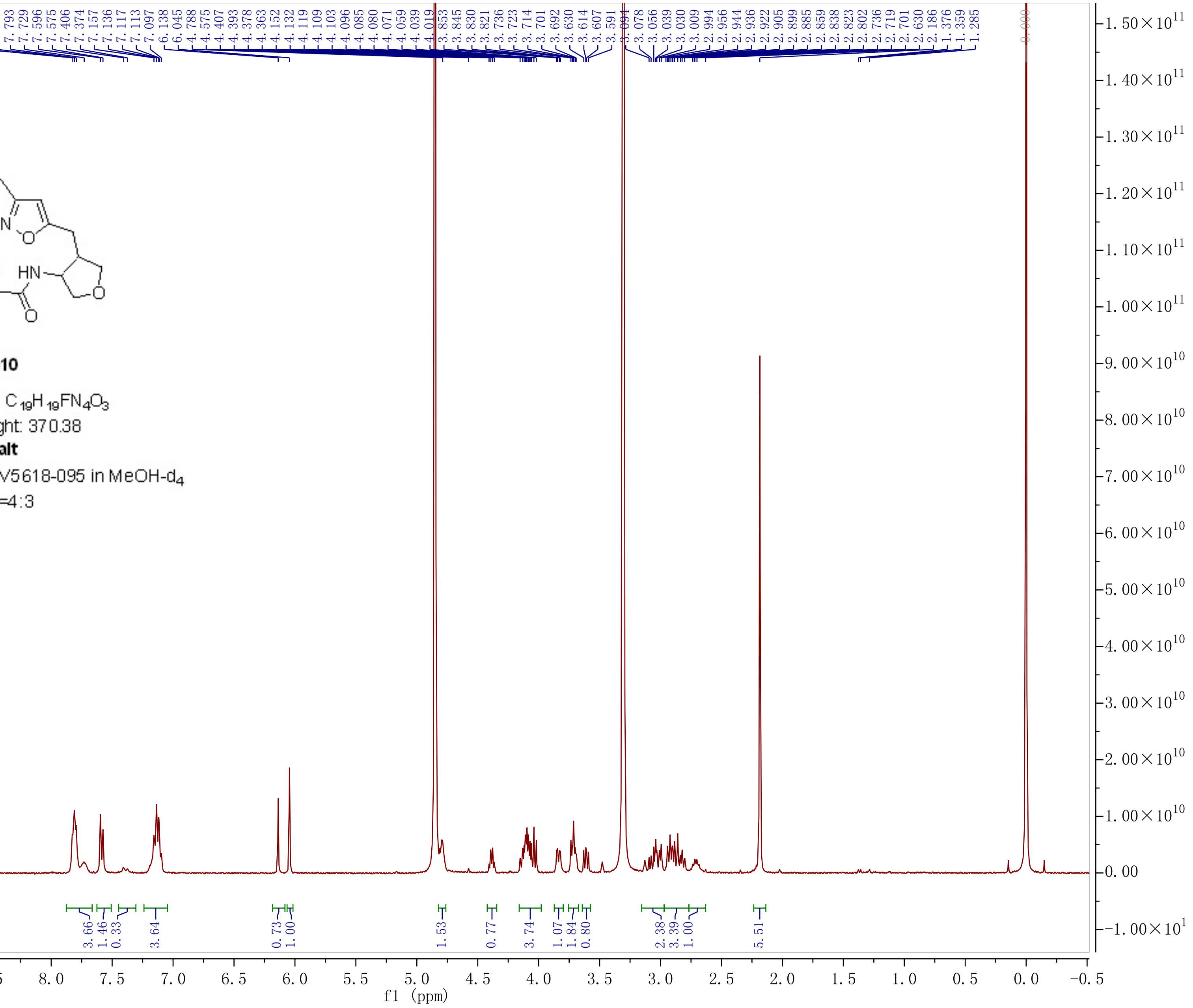

Chemical Formula: $C_{19}H_{19}FN_4O_2S$
Molecular Weight: 386.44

**1554-011**

**TFA salt**

$^1$H NMR 400M spectrum of **V5601-058** in $CD_3OD$

Note: cis/trans = 2/1

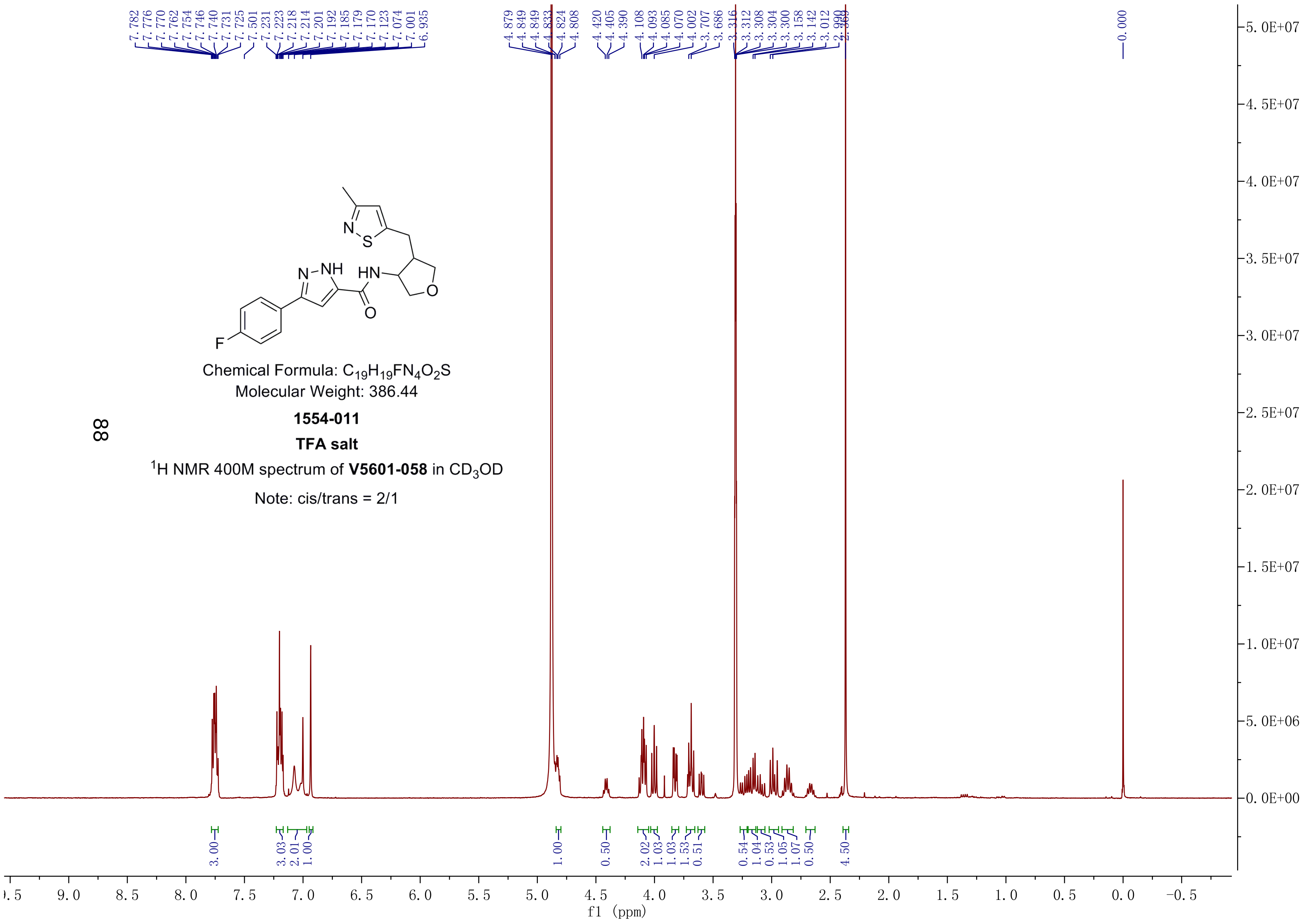

Chemical Formula: $C_{20}H_{21}FN_4O_3$
Molecular Weight: 384.40

**1554-012**

**TFA salt**

$^1$H NMR 400M spectrum of **V5601-089** in $CD_3OD$

Note: cis/trans = 2/1

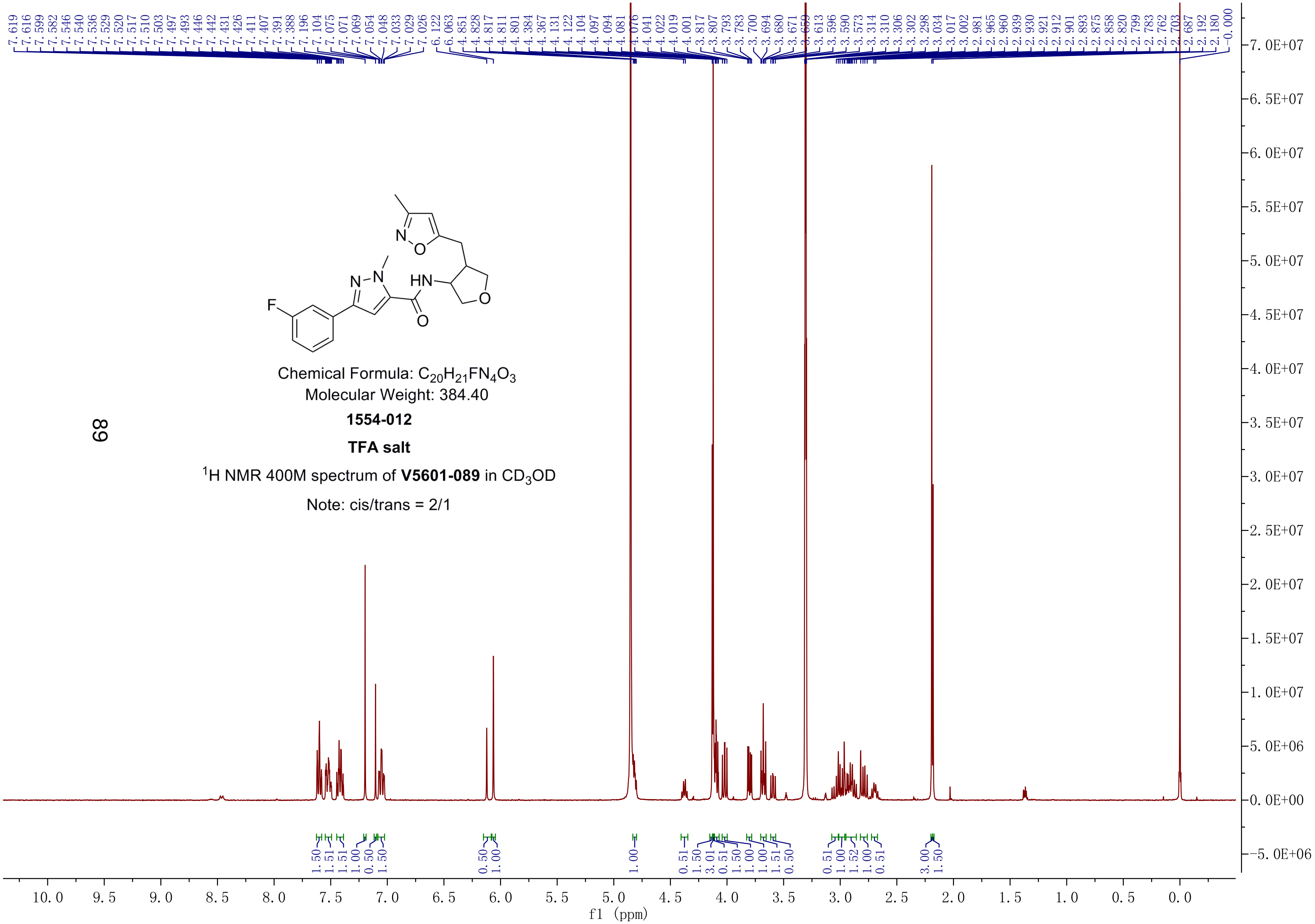

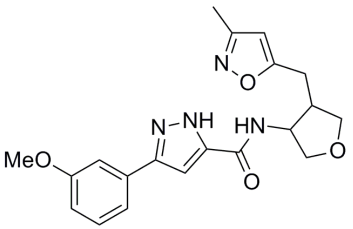

Chemical Formula: $C_{20}H_{22}N_4O_4$
Molecular Weight: 382.41
**1554-013**
**TFA salt**
$^1$H NMR 400M spectrum of **V5601-090** in $CD_3OD$
Note: cis/trans = 5/2

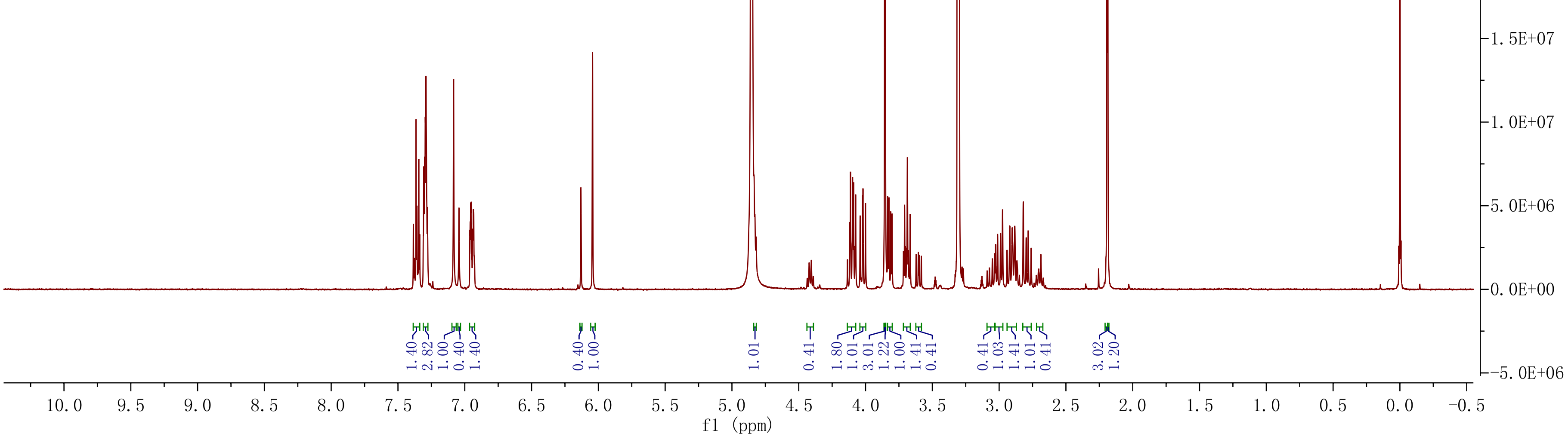

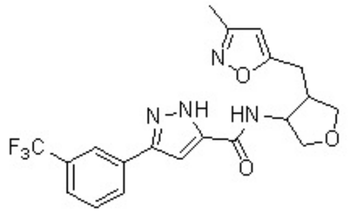

**1554-014**

Chemical Formula: $C_{20}H_{19}F_3N_4O_3$

Molecular Weight: 420.39

**TFA Salt**

$^1H$ NMR 400M spectrum of V5618-130 in $DMSOd_6$

cis:trans=5:3

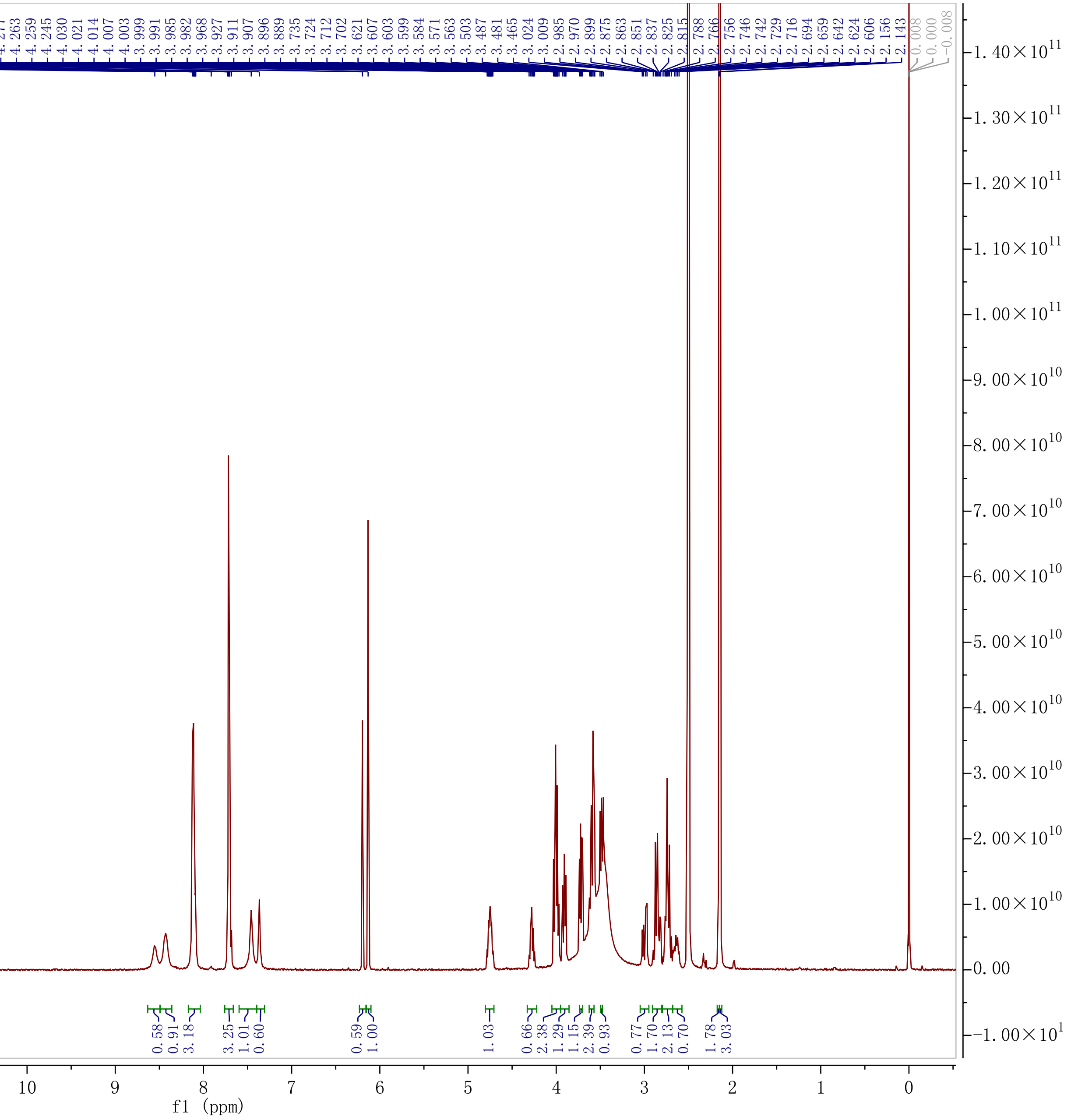

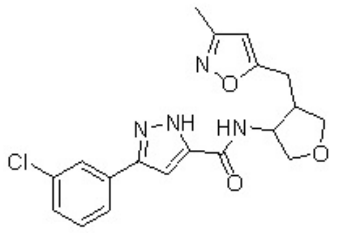


**1554-015**

Chemical Formula: $C_{19}H_{19}ClN_4O_3$

Molecular Weight: 386.83

$^1$H NMR 400M spectrum of V5618-129 in DMSO$d_6$(298 K)

cis:trans=9:4

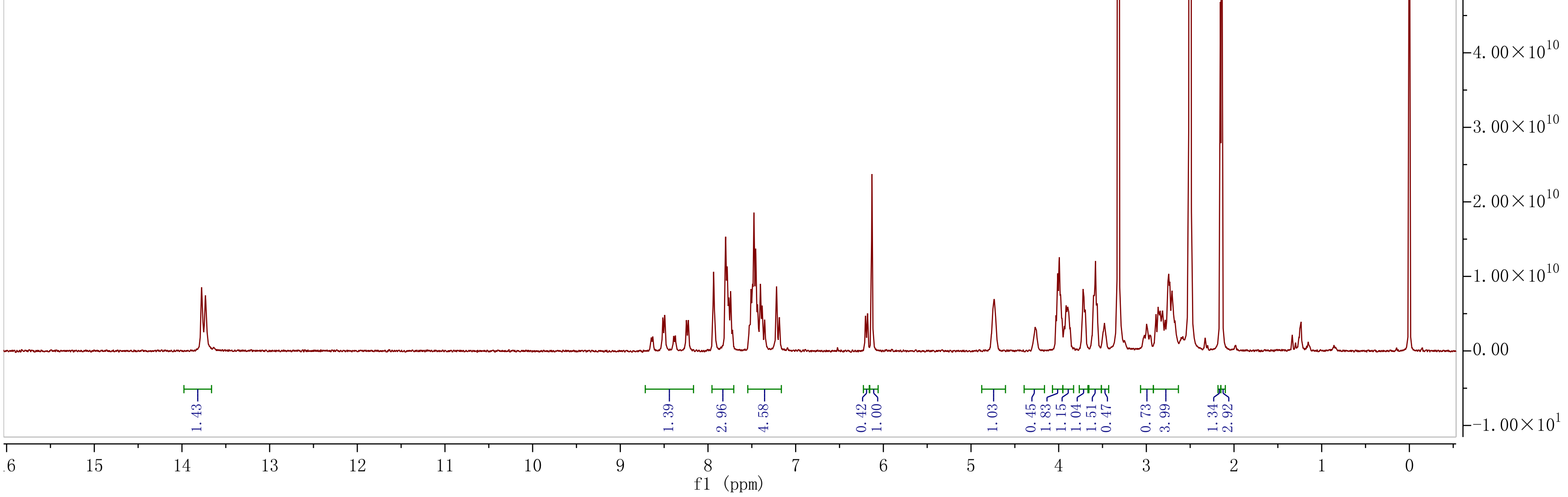

Chemical Formula: $C_{18}H_{19}N_5O_3$
Molecular Weight: 353.38
**1554-017**
TFA salt
$^1$H NMR 400M spectrum of **V4505-110** in $CD_3OD$
Note:cis/trans=10/7

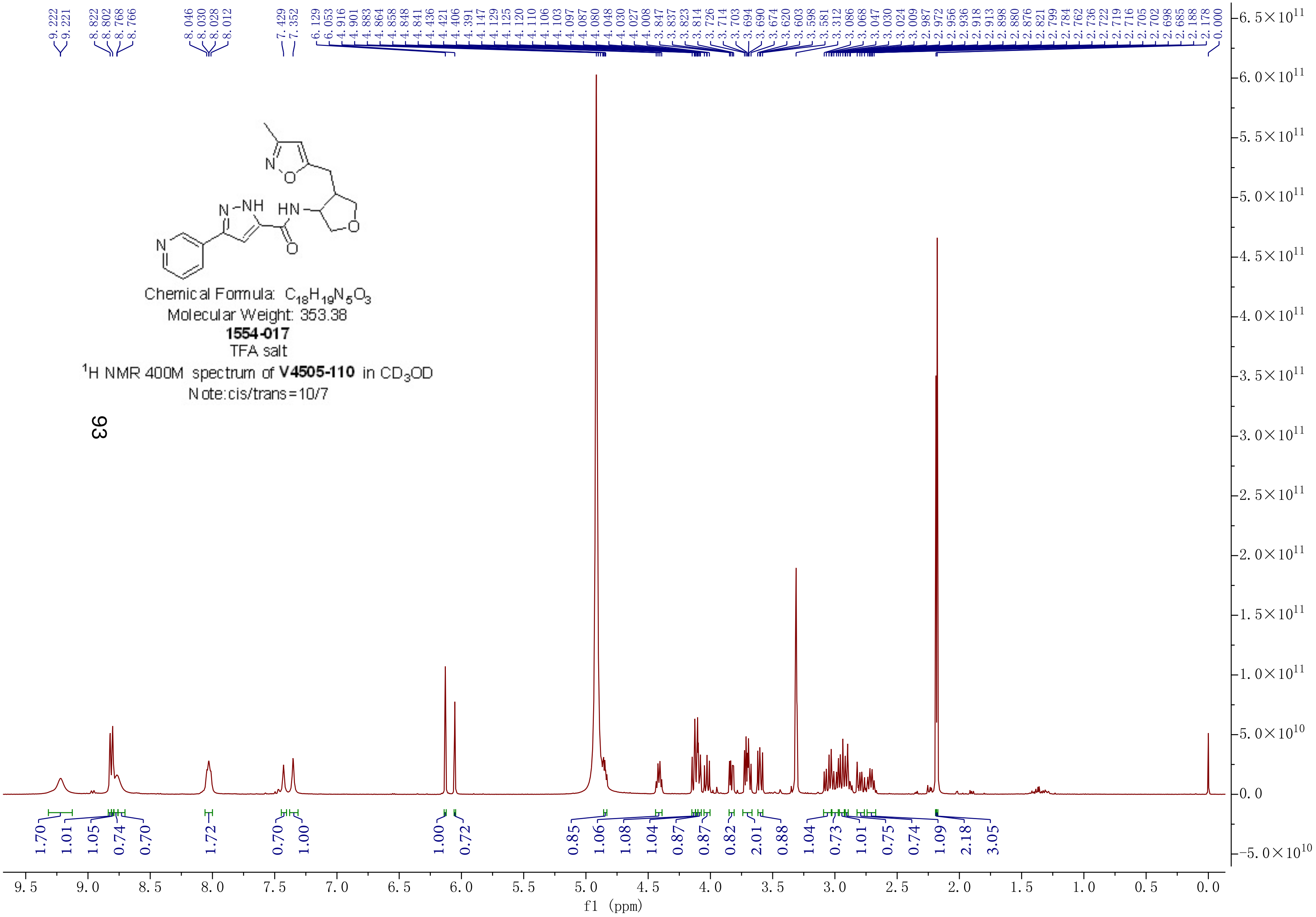

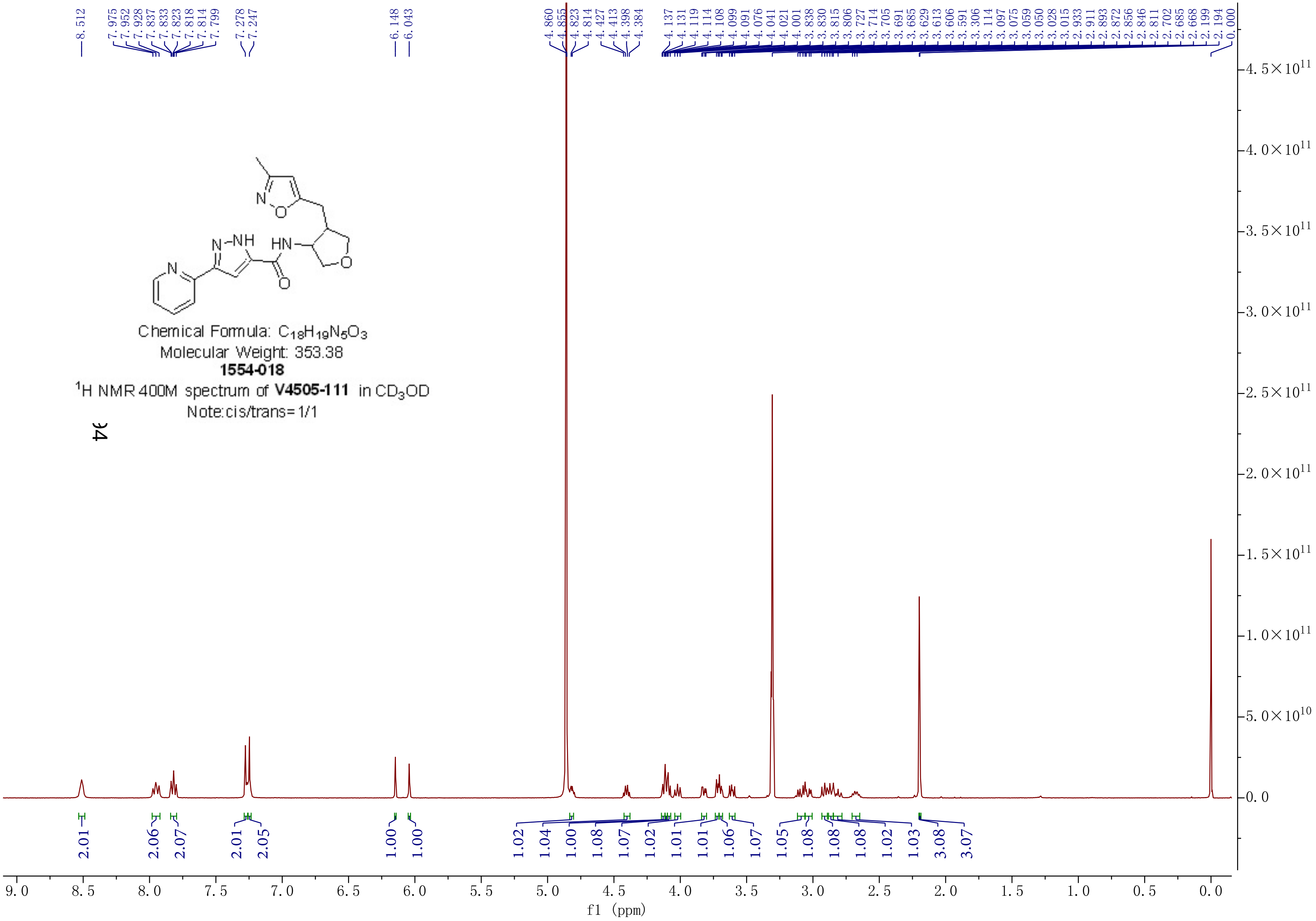
Chemical Formula: $C_{18}H_{19}N_5O_3$
Molecular Weight: 353.38
**1554-018**
$^1$H NMR 400M spectrum of **V4505-111** in $CD_3OD$
Note:cis/trans= 1/1
f1 (ppm)

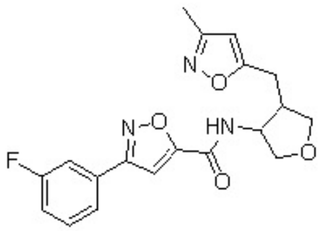

**1554-019**

Chemical Formula: $C_{19}H_{18}FN_3O_4$

Molecular Weight: 371.36

**TFA Salt**

$^1$H NMR 400M spectrum of V5618-146 in MeOH-$d_4$

cis:trans=7:1

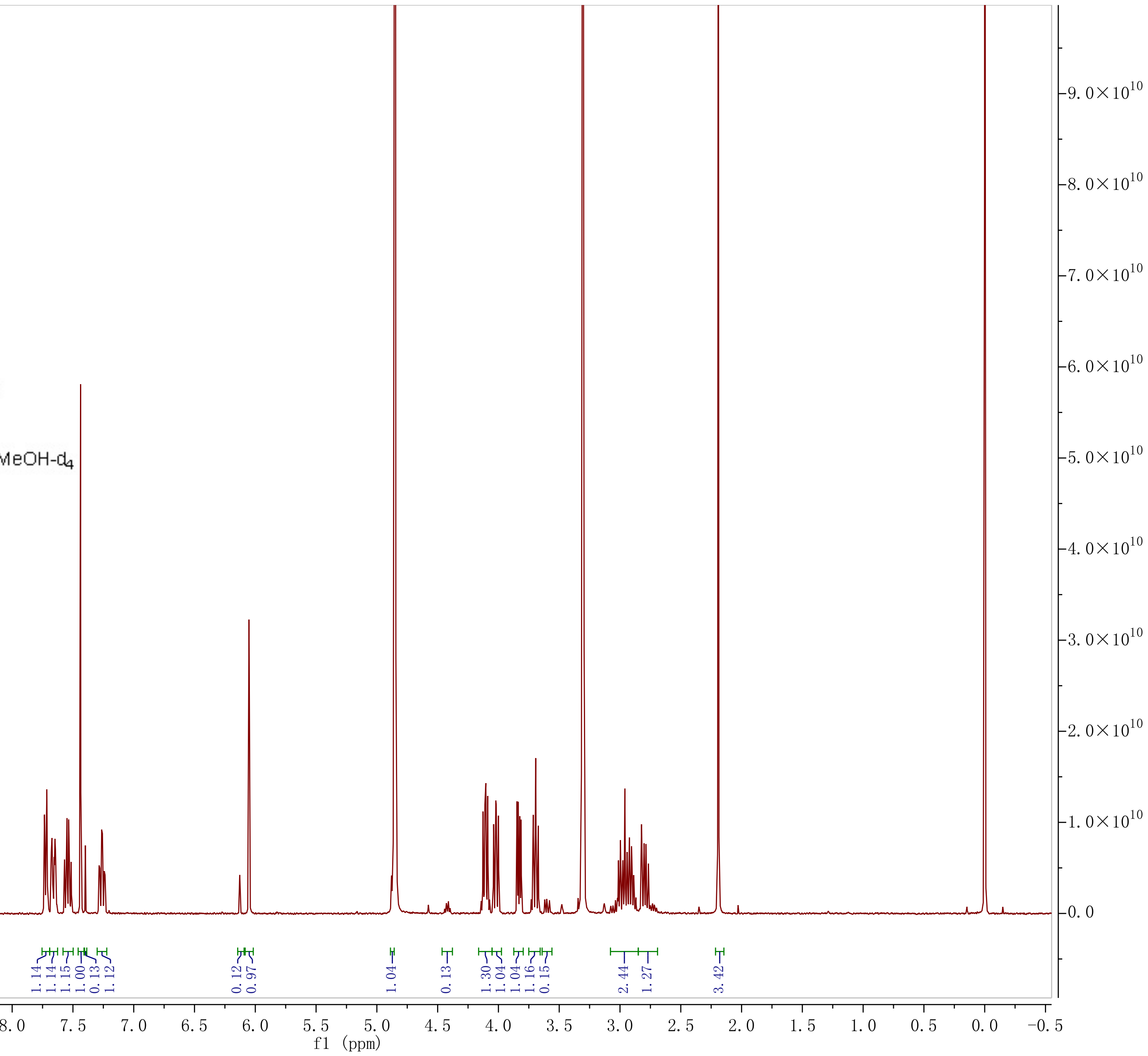

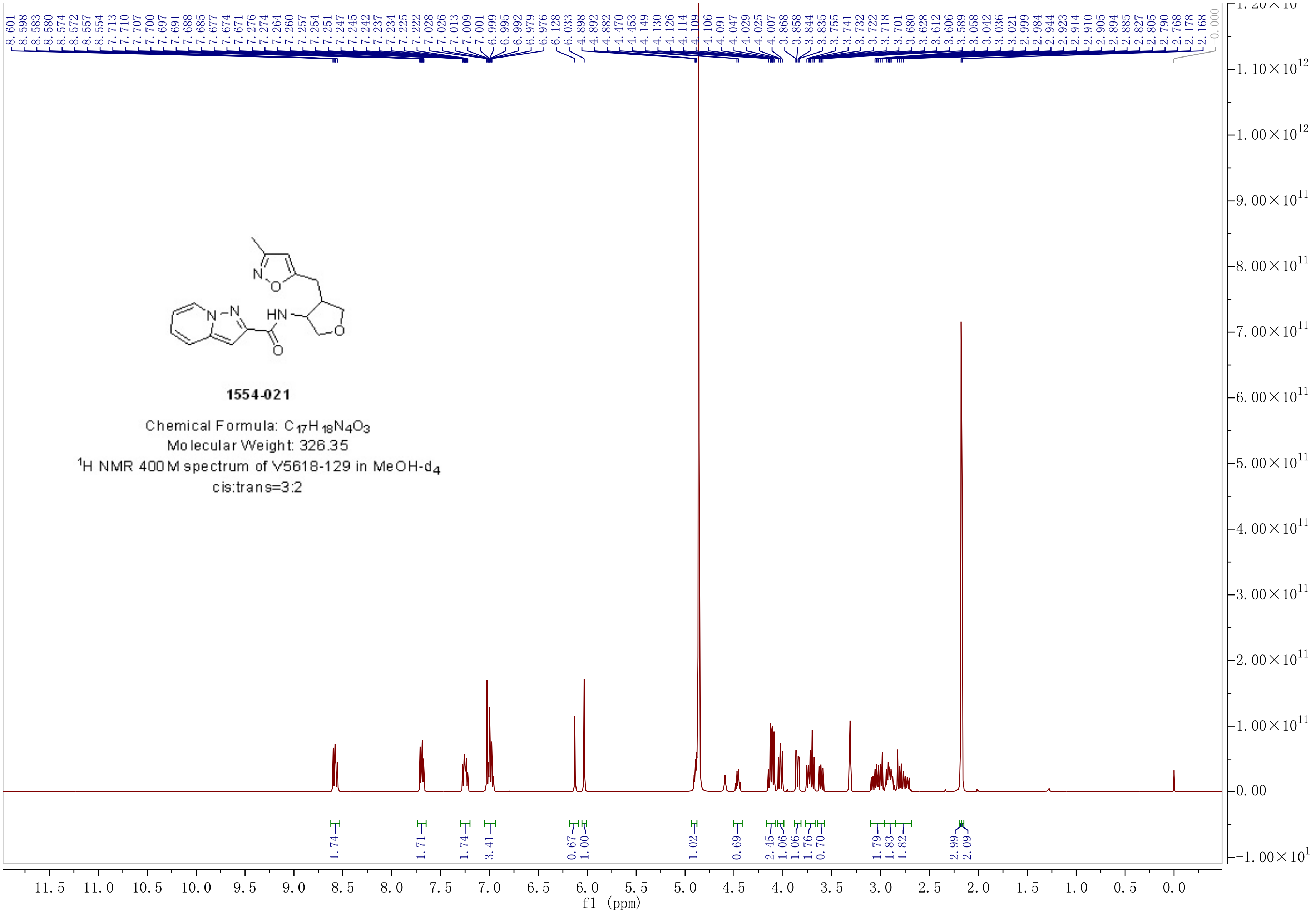

**1554-021**

Chemical Formula: $C_{17}H_{18}N_4O_3$

Molecular Weight: 326.35

$^1$H NMR 400M spectrum of V5618-129 in MeOH-$d_4$

cis:trans=3:2

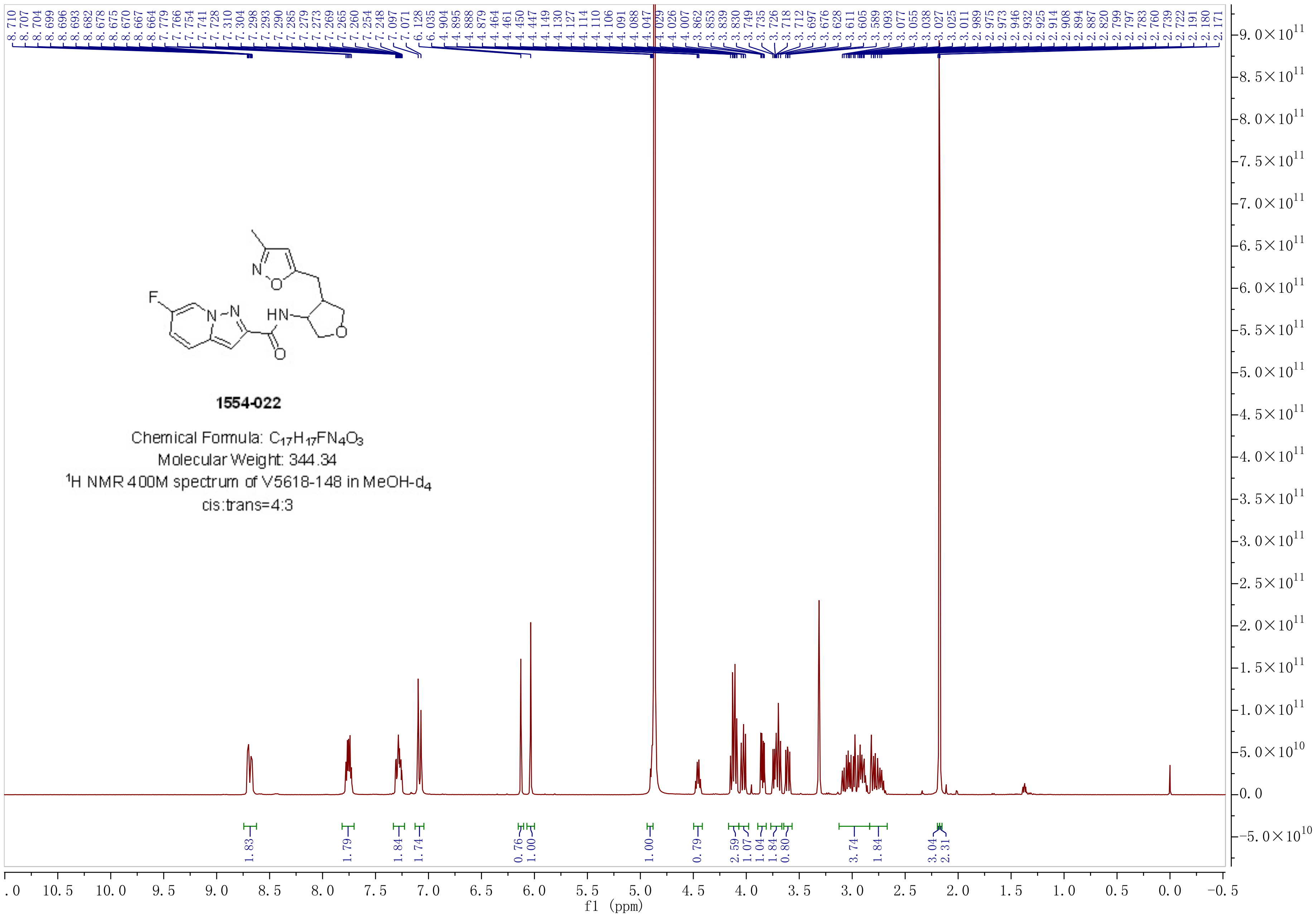
1554-022
Chemical Formula: C17H17FN4O3
Molecular Weight: 344.34
1H NMR 400M spectrum of V5618-148 in MeOH-d4
cis:trans=4:3
8.710 8.707 8.704 8.699 8.696 8.693 8.682 8.678 8.675 8.670 8.667 8.664 7.779 7.766 7.754 7.741 7.728 7.310 7.304 7.298 7.293 7.290 7.285 7.279 7.273 7.269 7.265 7.260 7.254 7.248 7.097 7.071 6.128 6.035 4.904 4.895 4.888 4.879 4.464 4.461 4.450 4.447 4.149 4.130 4.127 4.114 4.110 4.106 4.091 4.088 4.047 4.029 4.026 4.007 3.862 3.853 3.839 3.830 3.749 3.735 3.726 3.718 3.712 3.697 3.676 3.628 3.611 3.605 3.589 3.093 3.077 3.055 3.038 3.027 3.025 3.011 2.989 2.975 2.973 2.946 2.932 2.925 2.914 2.908 2.894 2.887 2.820 2.799 2.797 2.783 2.760 2.739 2.722 2.191 2.180 2.171
1.83 1.79 1.84 1.74 0.76 1.00 1.00 0.79 2.59 1.07 1.04 1.84 0.80 3.74 1.84 3.04 2.31
f1 (ppm)

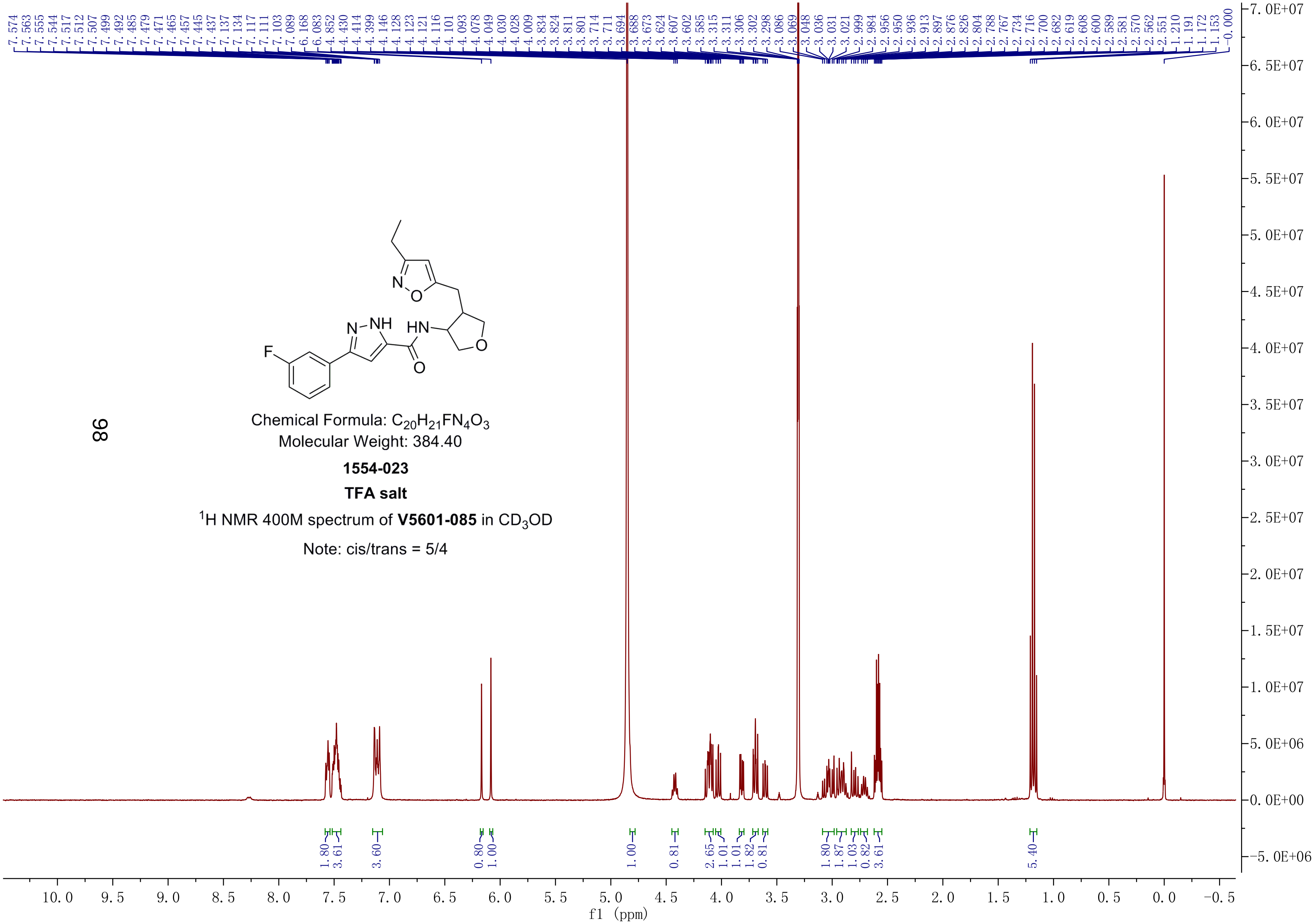


Chemical Formula: $C_{20}H_{21}FN_4O_3$
Molecular Weight: 384.40

**1554-023**

**TFA salt**

$^1$H NMR 400M spectrum of **V5601-085** in $CD_3OD$

Note: cis/trans = 5/4

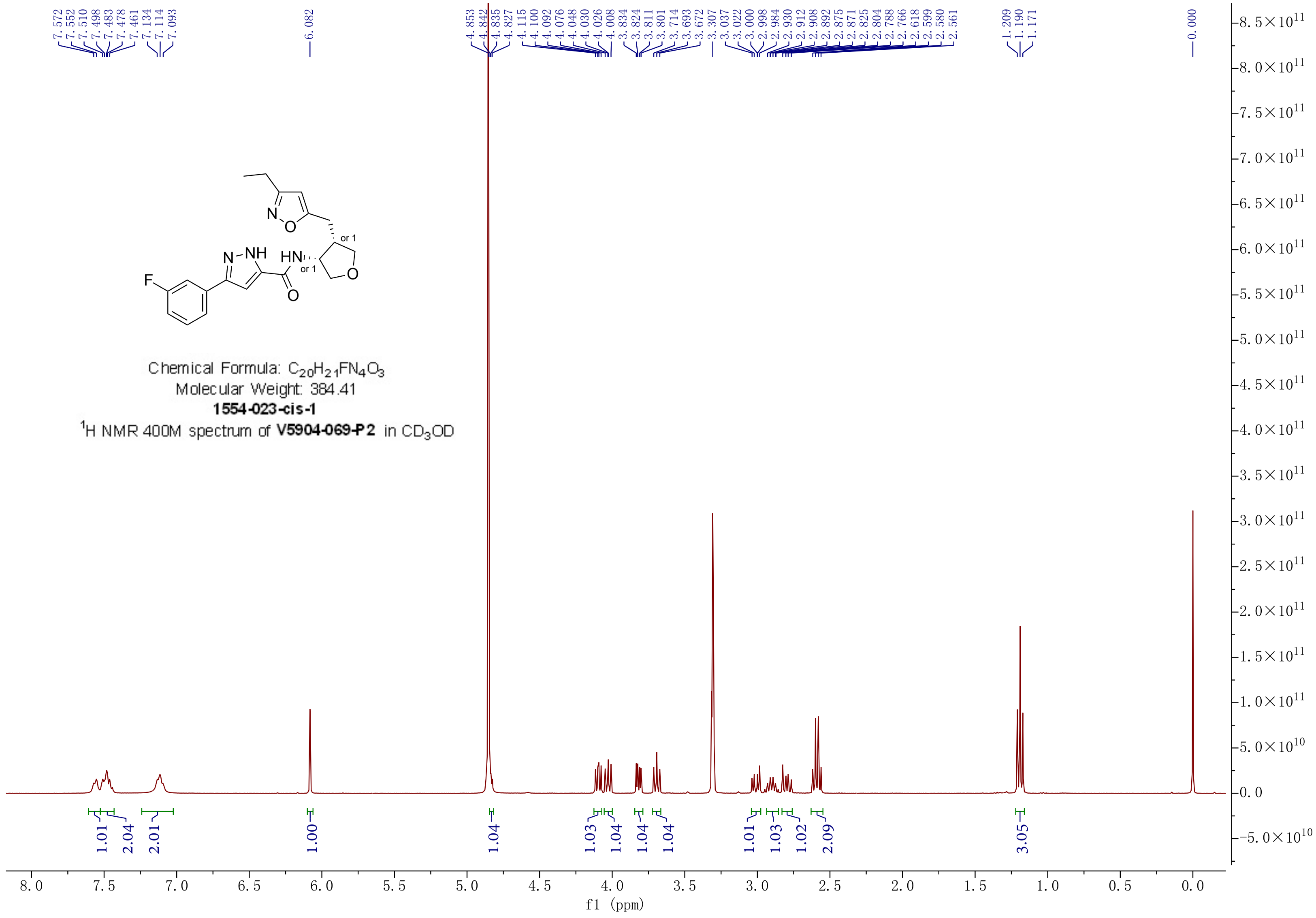
Chemical Formula: $C_{20}H_{21}FN_4O_3$
Molecular Weight: 384.41
**1554-023-cis-1**
$^1$H NMR 400M spectrum of **V5904-069-P2** in $CD_3OD$
7.572 7.552 7.510 7.498 7.483 7.478 7.461 7.134 7.114 7.093
6.082
4.853 4.842 4.835 4.827 4.115 4.100 4.092 4.076 4.048 4.030 4.026 4.008 3.834 3.824 3.811 3.801 3.714 3.693 3.672 3.307 3.037 3.022 3.000 2.998 2.984 2.930 2.912 2.908 2.892 2.875 2.871 2.825 2.804 2.788 2.766 2.618 2.599 2.580 2.561
1.209 1.190 1.171
0.000
1.01 2.04 2.01 1.00 1.04 1.03 1.04 1.04 1.04 1.01 1.03 1.02 2.09 3.05
f1 (ppm)

Chemical Formula: $C_{20}H_{21}FN_4O_3$
Molecular Weight: 384.41
**1554-023-cis-2**
$^1$H NMR 400M spectrum of **V5904-069-P4** in $CD_3OD$

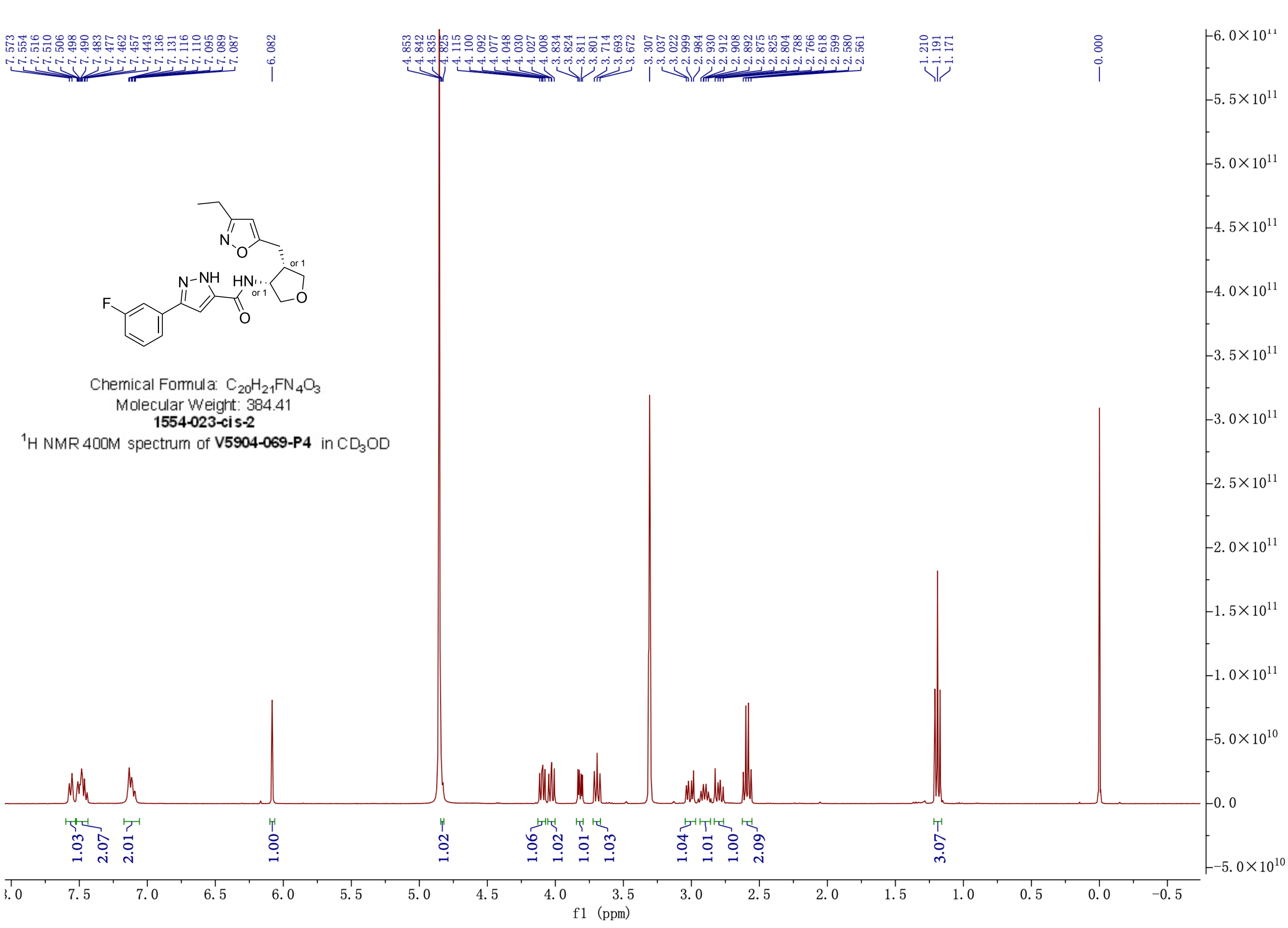

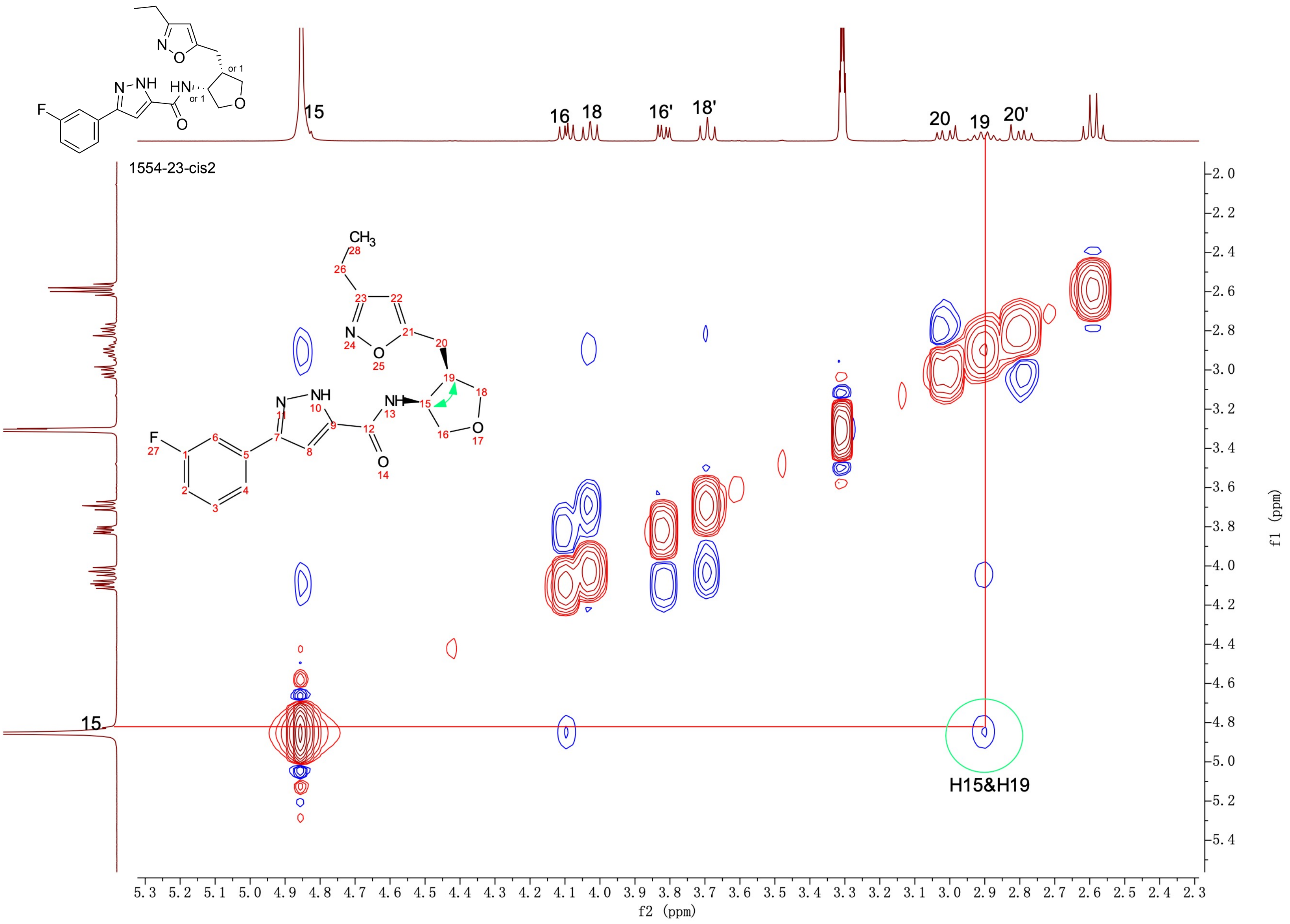

1554-23-cis2
15
16
18
16'
18'
20
19
20'
H15&H19
f2 (ppm)
f1 (ppm)

Chemical Formula: $C_{20}H_{21}FN_4O_3$
Molecular Weight: 384.41
**1554-023-trans-1**
$^1$H NMR 400M spectrum of **V5904-069-P1** in $CD_3OD$

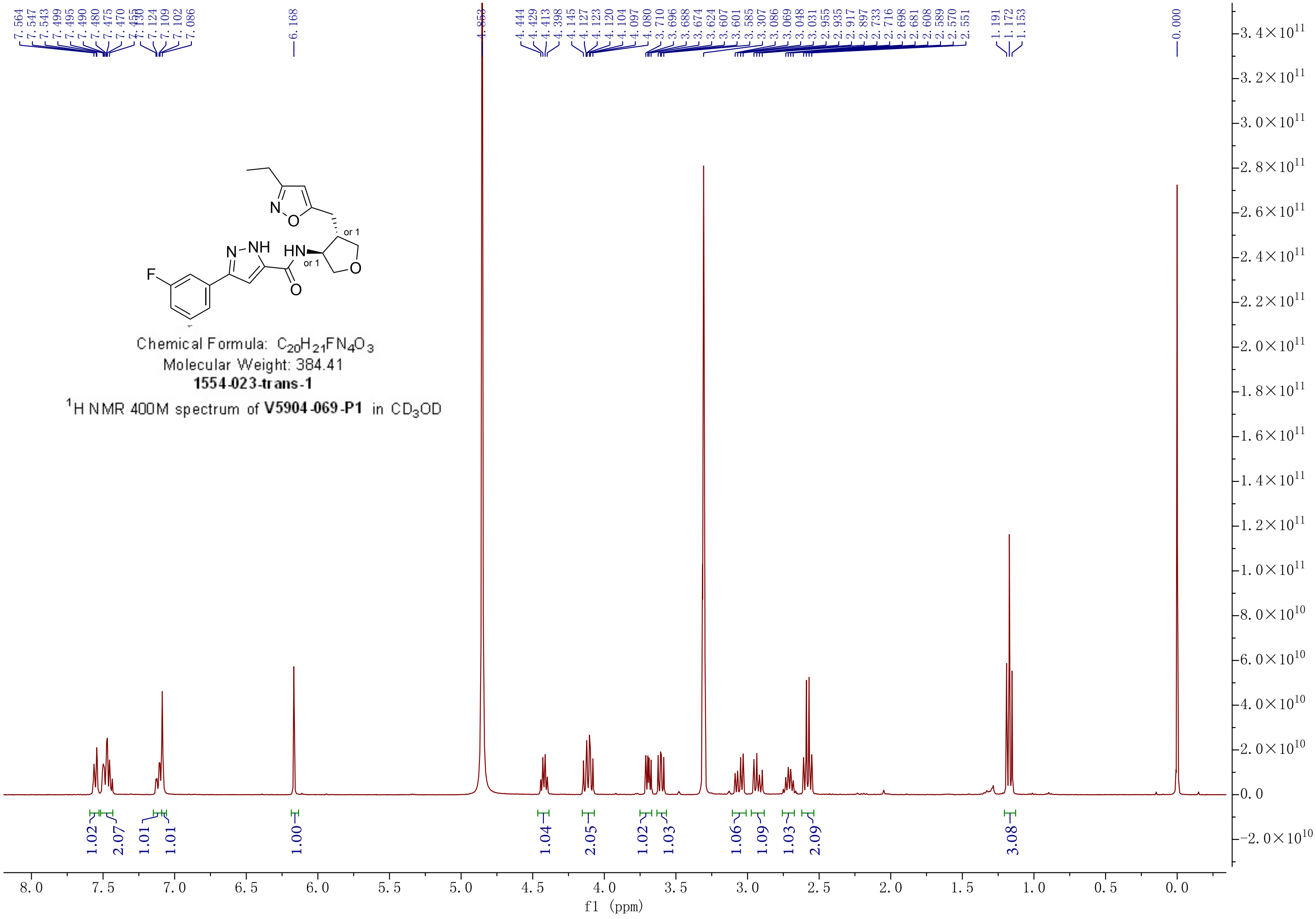

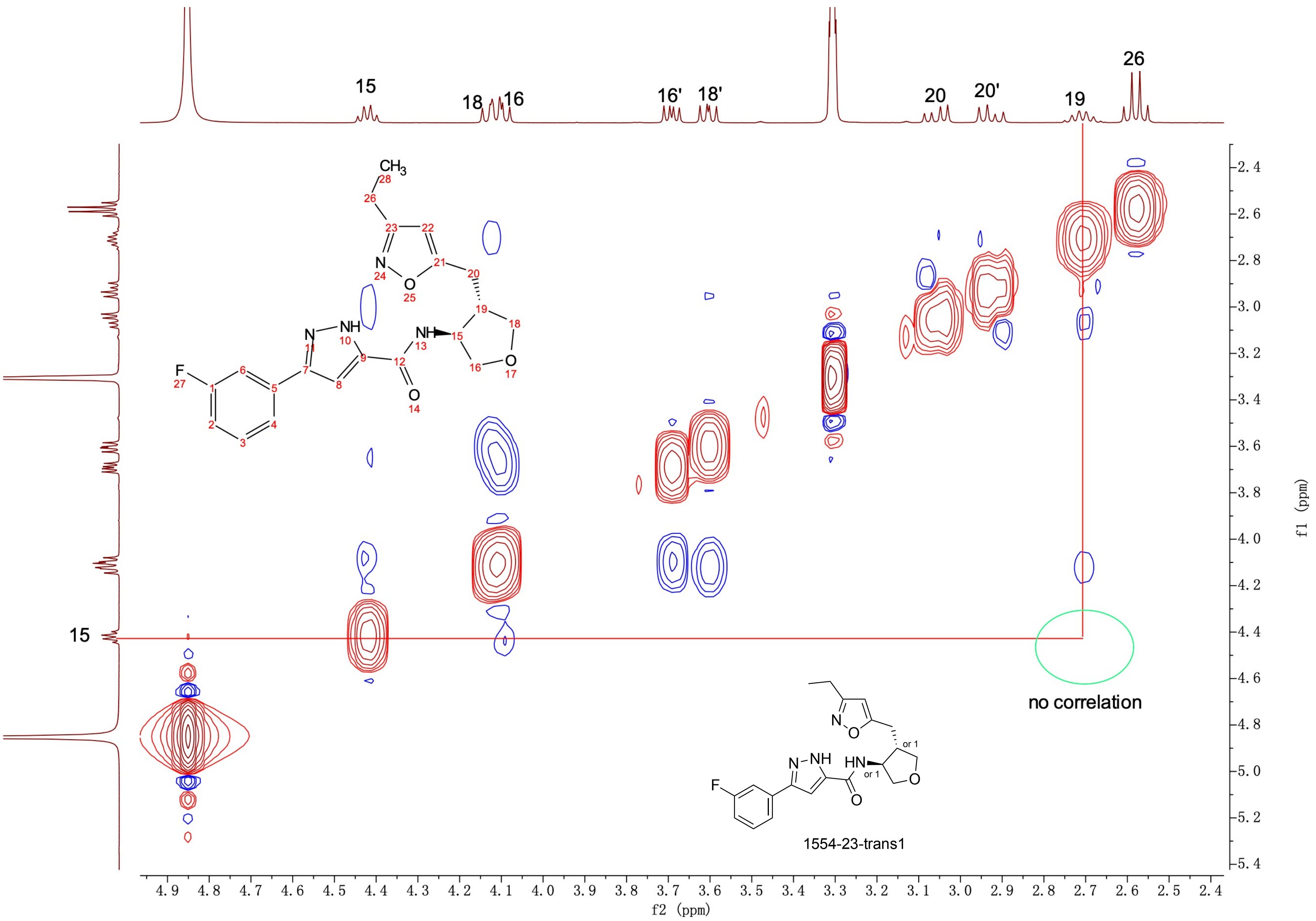

15
18
16
16'
18'
20
20'
19
26
no correlation
1554-23-trans1
or 1
f2 (ppm)
f1 (ppm)

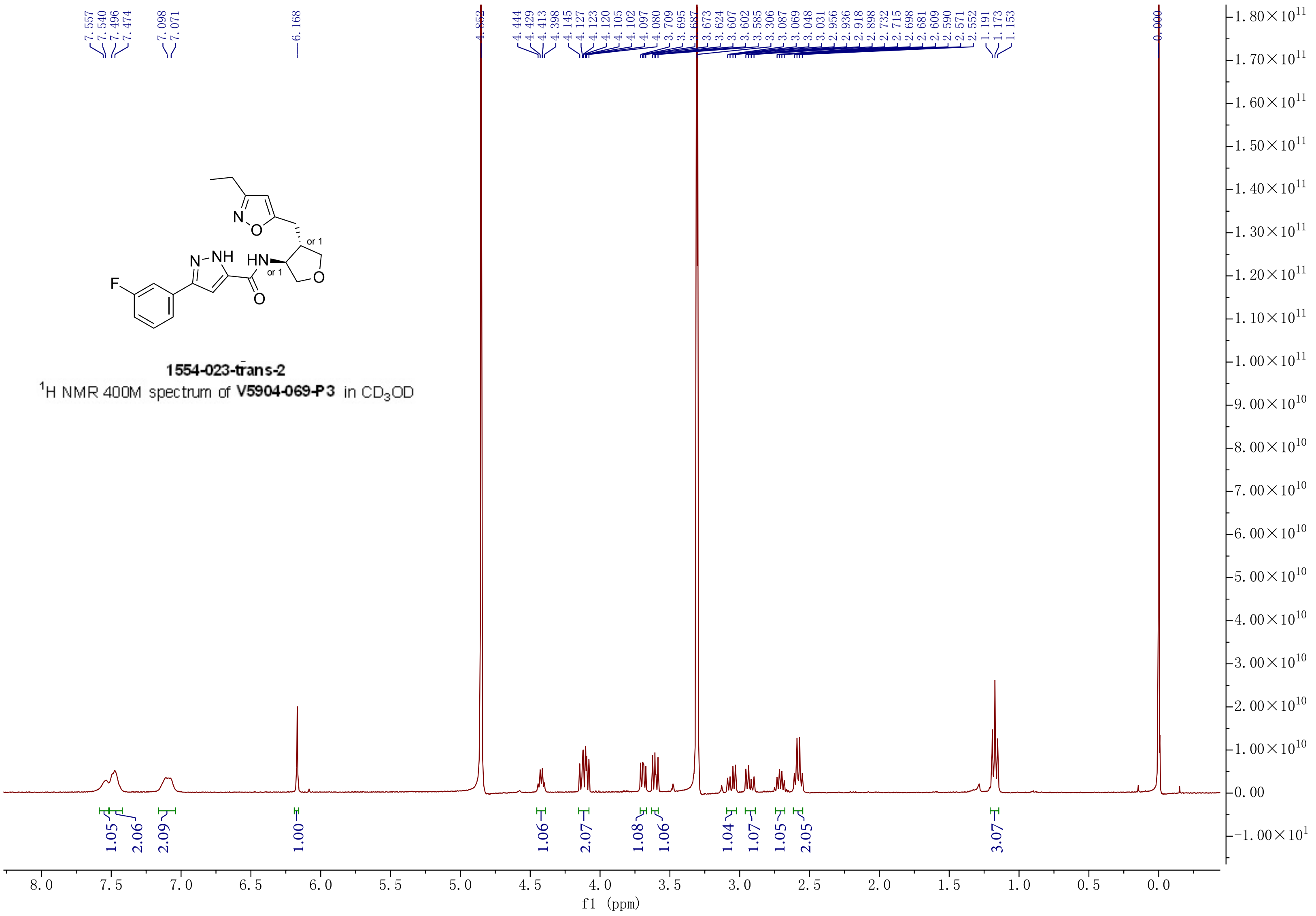


**1554-023-trans-2**

$^1$H NMR 400M spectrum of **V5904-069-P3** in $CD_3OD$

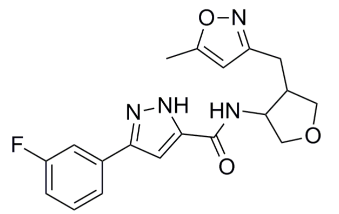


Chemical Formula: $C_{19}H_{19}FN_4O_3$
Molecular Weight: 370.38

**1554-024**

**TFA salt**

$^1$H NMR 400M spectrum of **V5601-087** in $CD_3OD$

Ratio: 10/7

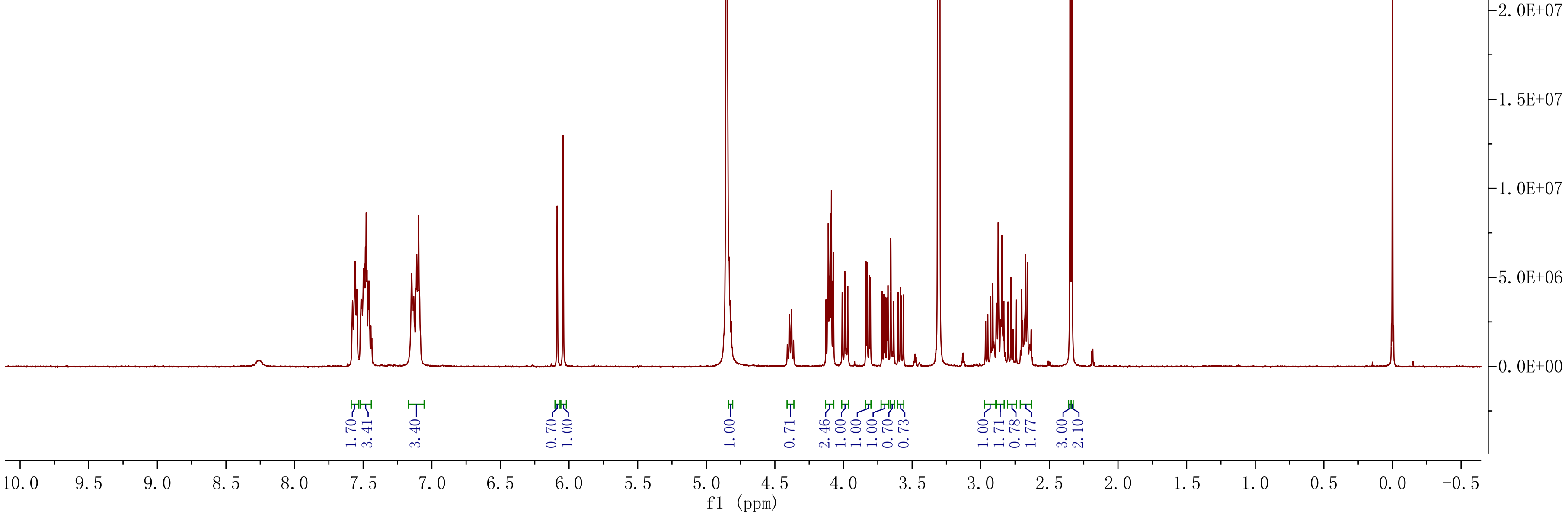

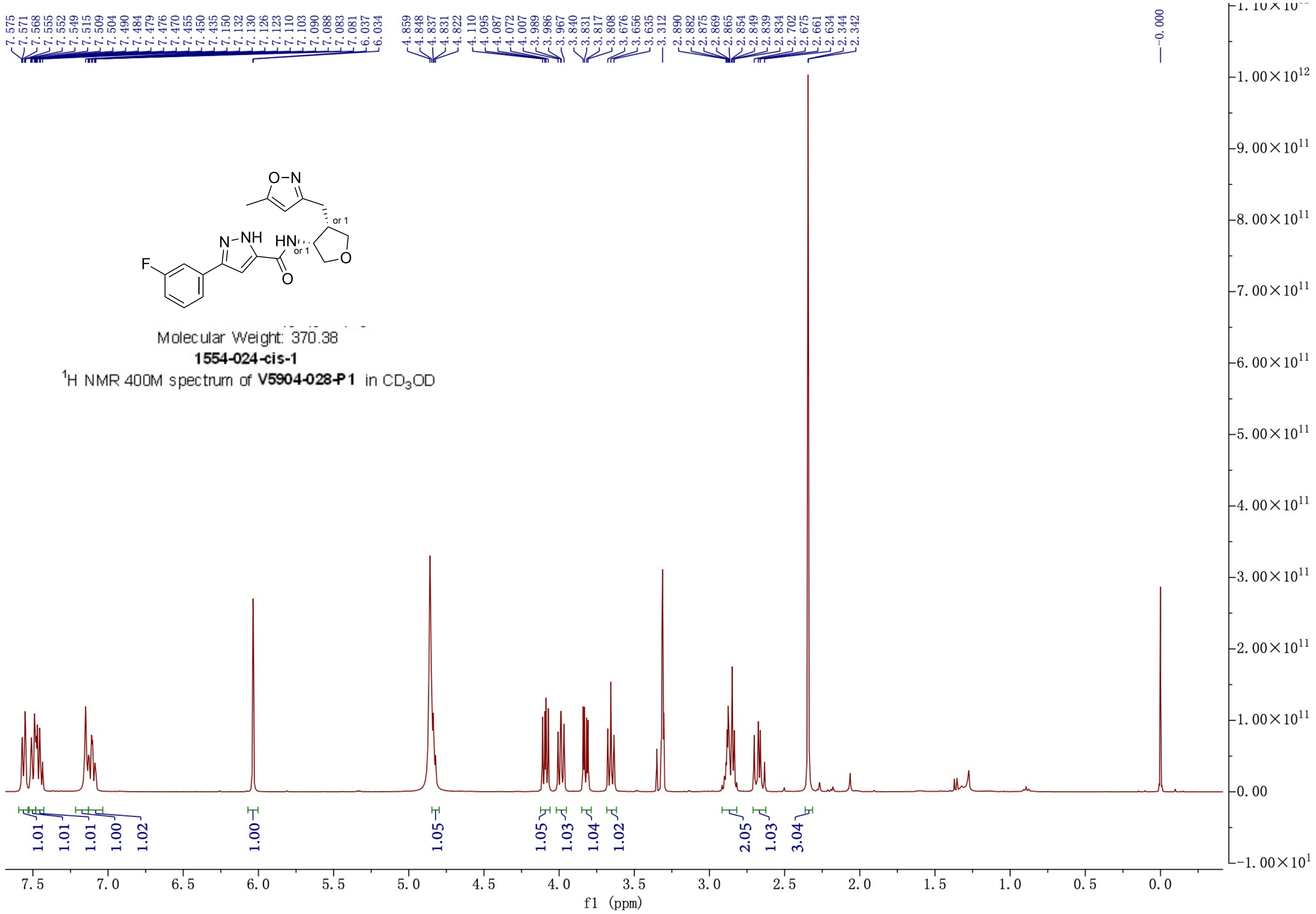

Molecular Weight: 370.38
1554-024-cis-1
$^1H$ NMR 400M spectrum of V5904-028-P1 in $CD_3OD$
f1 (ppm)

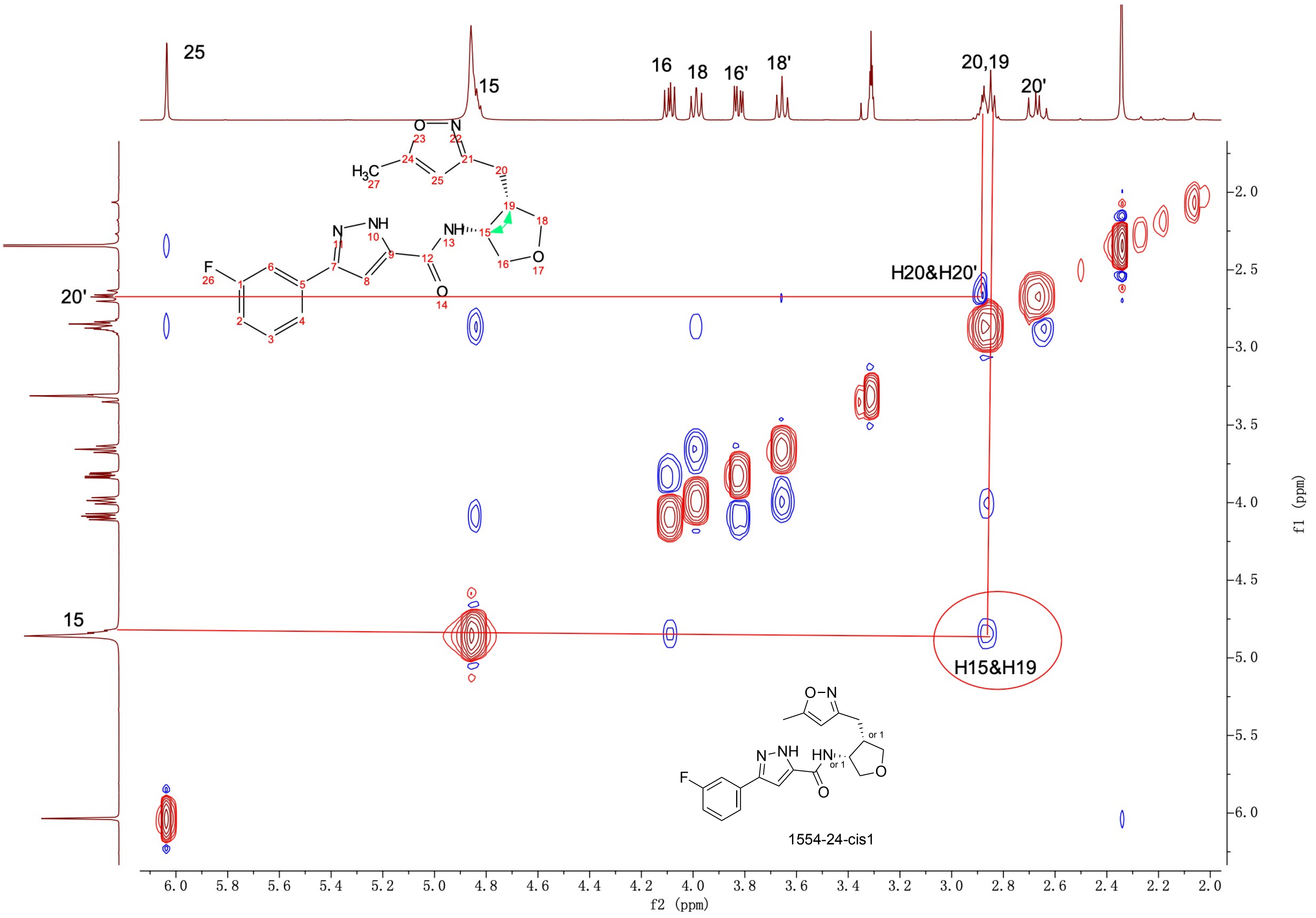
25
15
16
18
16'
18'
20,19
20'
H20&H20'
H15&H19
1554-24-cis1
f2 (ppm)
f1 (ppm)

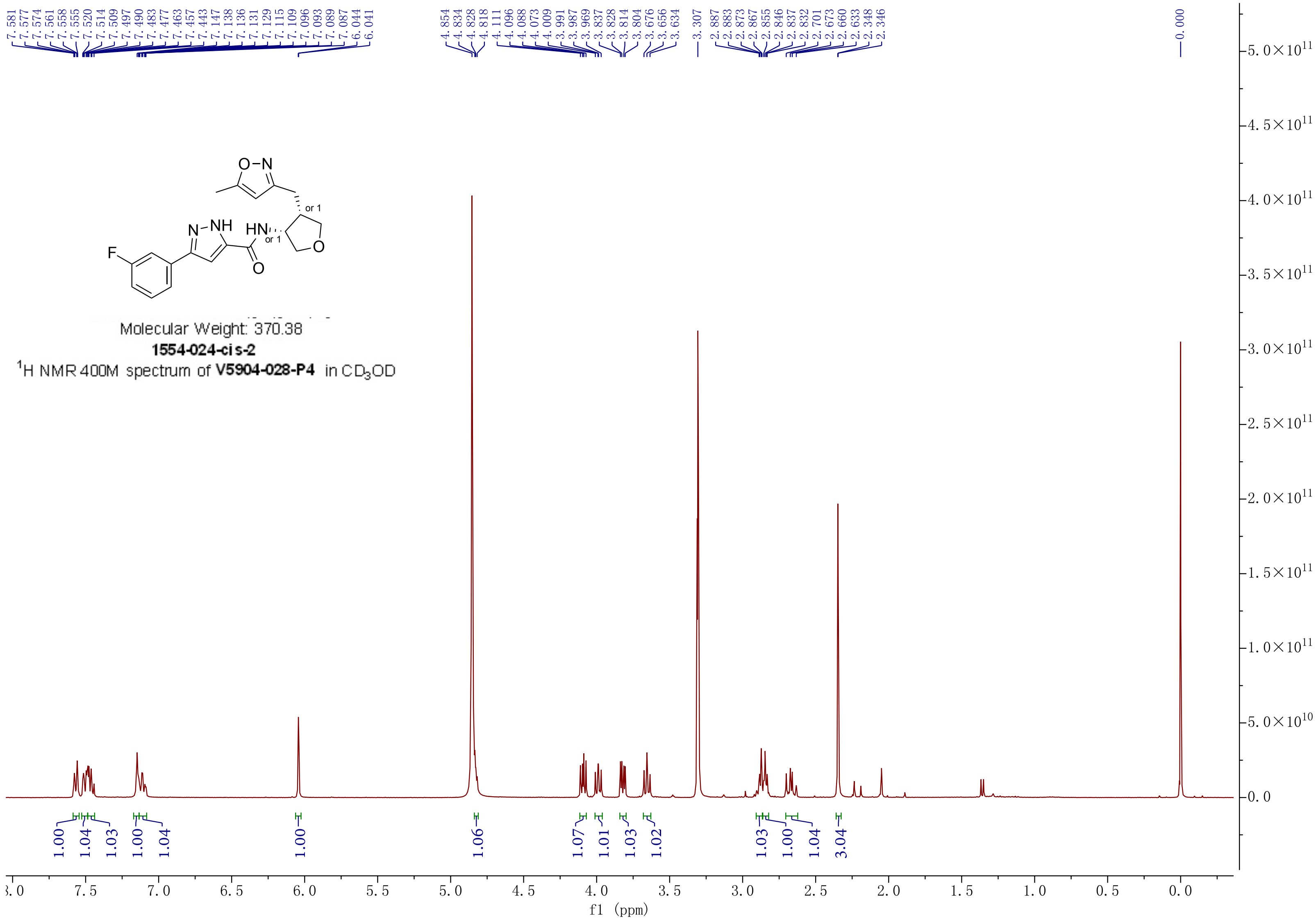

7.581 7.577 7.574 7.561 7.558 7.555 7.520 7.514 7.509 7.497 7.490 7.483 7.477 7.463 7.457 7.443 7.147 7.138 7.136 7.131 7.129 7.115 7.109 7.096 7.093 7.089 7.087 6.044 6.041
4.854 4.834 4.828 4.818 4.111 4.096 4.088 4.073 4.009 3.991 3.987 3.969 3.837 3.828 3.814 3.804 3.676 3.656 3.634
3.307
2.887 2.883 2.873 2.867 2.855 2.846 2.837 2.832 2.701 2.673 2.660 2.633 2.348 2.346
0.000
Molecular Weight: 370.38
1554-024-cis-2
$^1$H NMR 400M spectrum of V5904-028-P4 in $CD_3OD$
1.00 1.04 1.03 1.00 1.04 1.00 1.06 1.07 1.01 1.03 1.02 1.03 1.00 1.04 3.04
f1 (ppm)

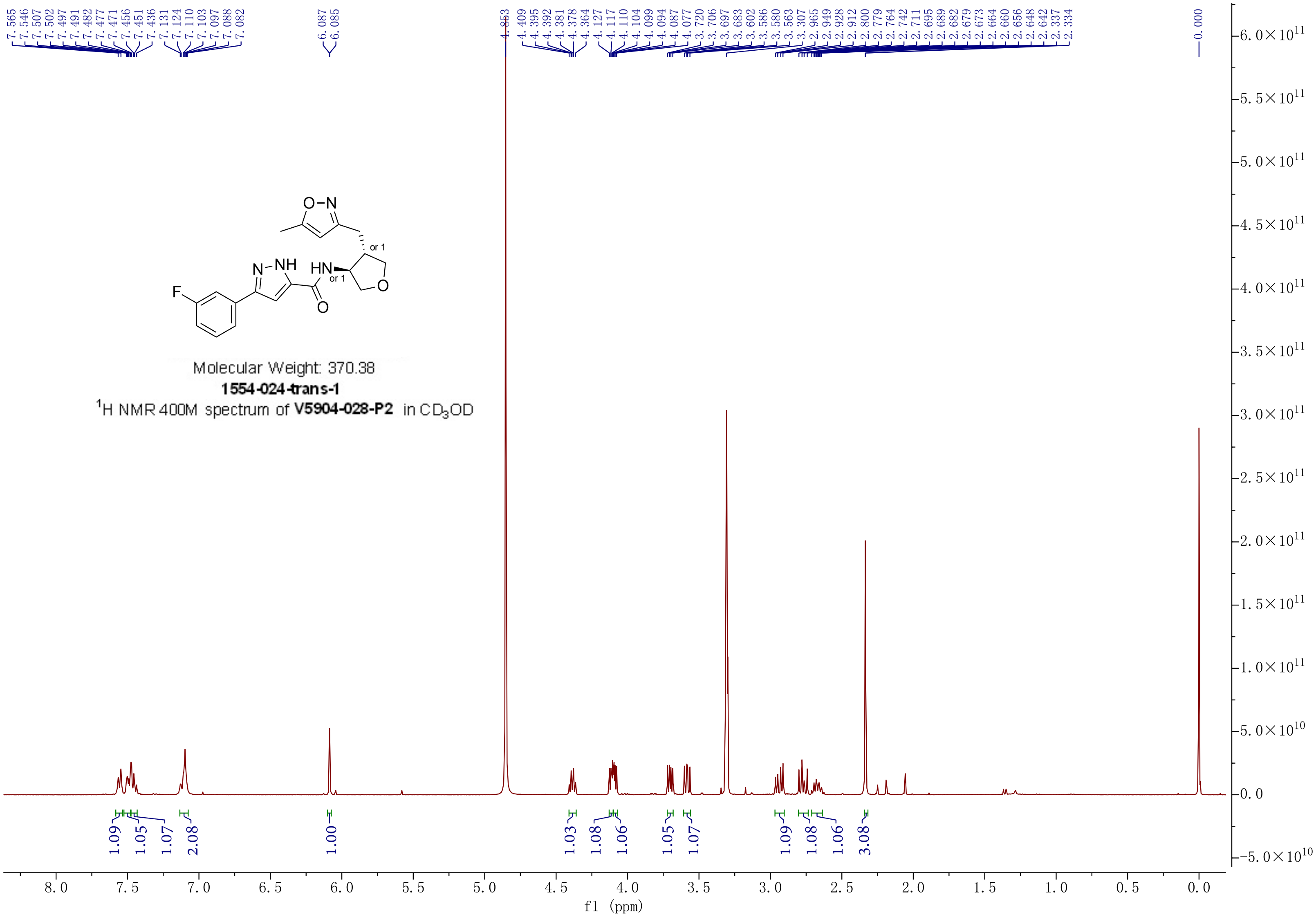


Molecular Weight: 370.38

**1554-024-trans-1**

$^1H$ NMR 400M spectrum of **V5904-028-P2** in $CD_3OD$

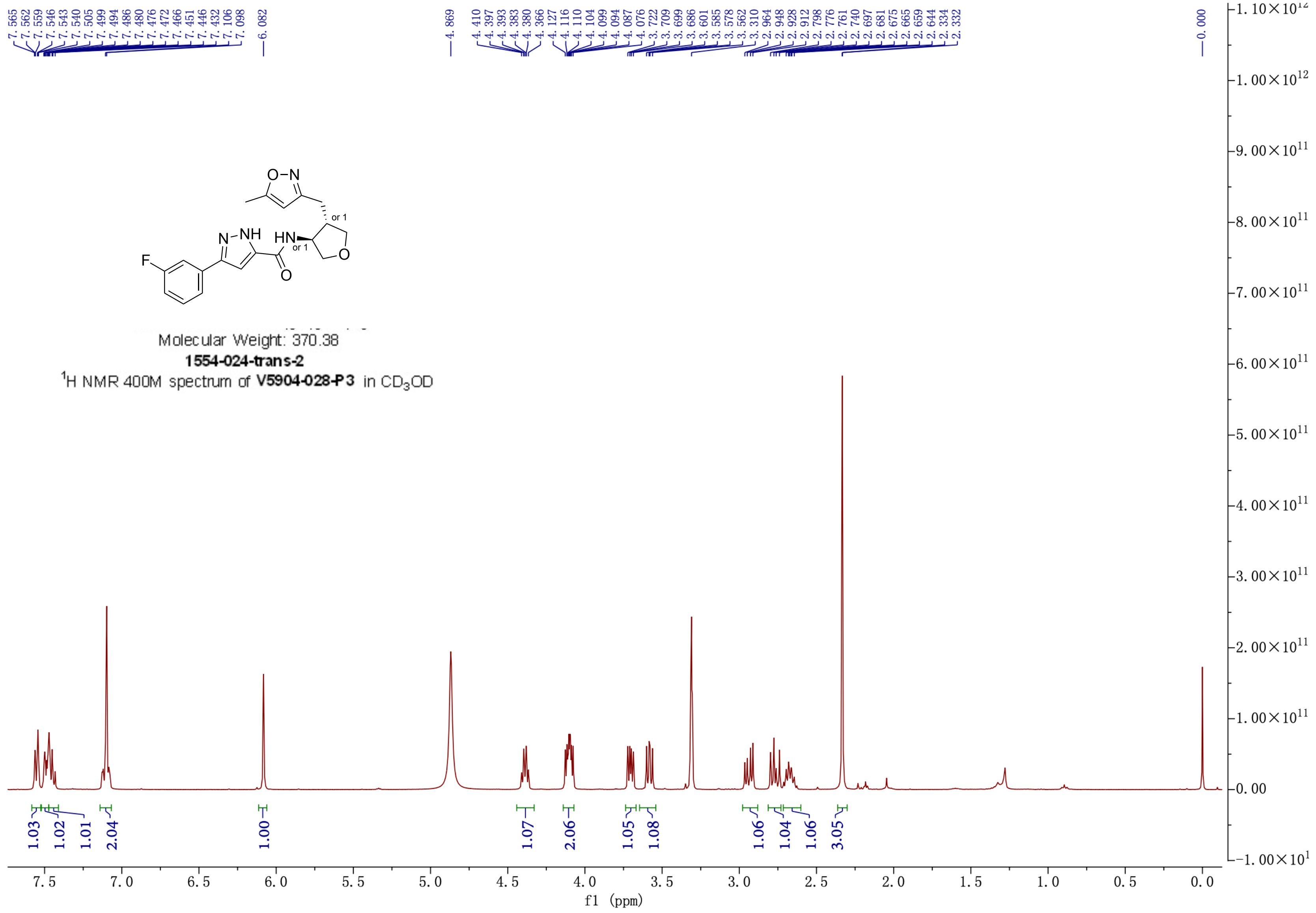


Molecular Weight: 370.38

**1554-024-trans-2**

$^1$H NMR 400M spectrum of **V5904-028-P3** in $CD_3OD$

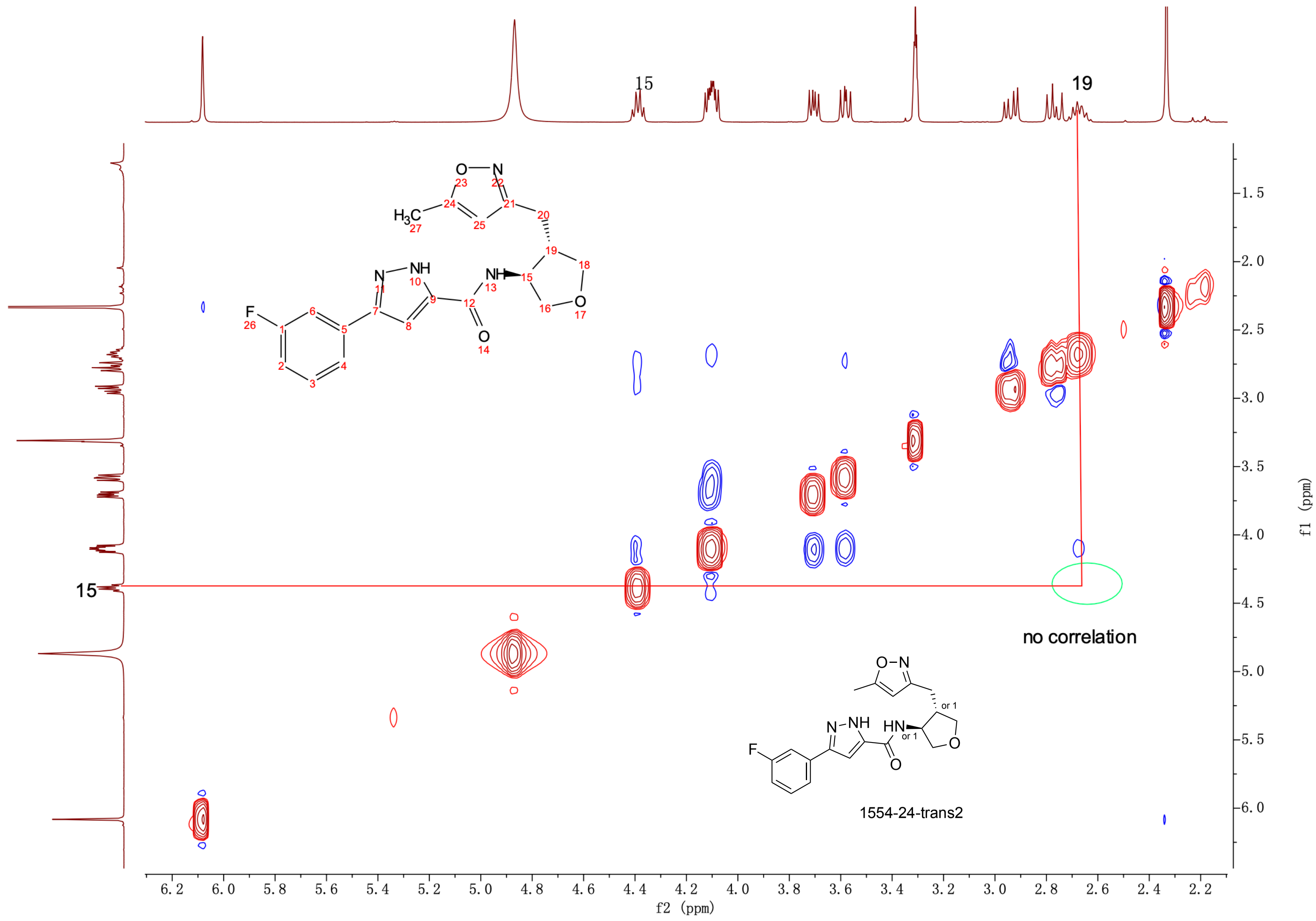
15
19
no correlation
1554-24-trans2
f2 (ppm)
f1 (ppm)
6.2
6.0
5.8
5.6
5.4
5.2
5.0
4.8
4.6
4.4
4.2
4.0
3.8
3.6
3.4
3.2
3.0
2.8
2.6
2.4
2.2
1.5
2.0
2.5
3.0
3.5
4.0
4.5
5.0
5.5
6.0

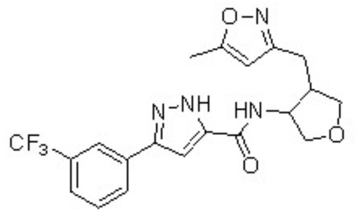

Chemical Formula: $C_{20}H_{19}F_3N_4O_3$
Molecular Weight: 420.39
**1554-025**
TFA salt
$^{1}$H NMR 400M spectrum of **V4505-136** in $CD_3OD$
Note:cis/trans=10/7

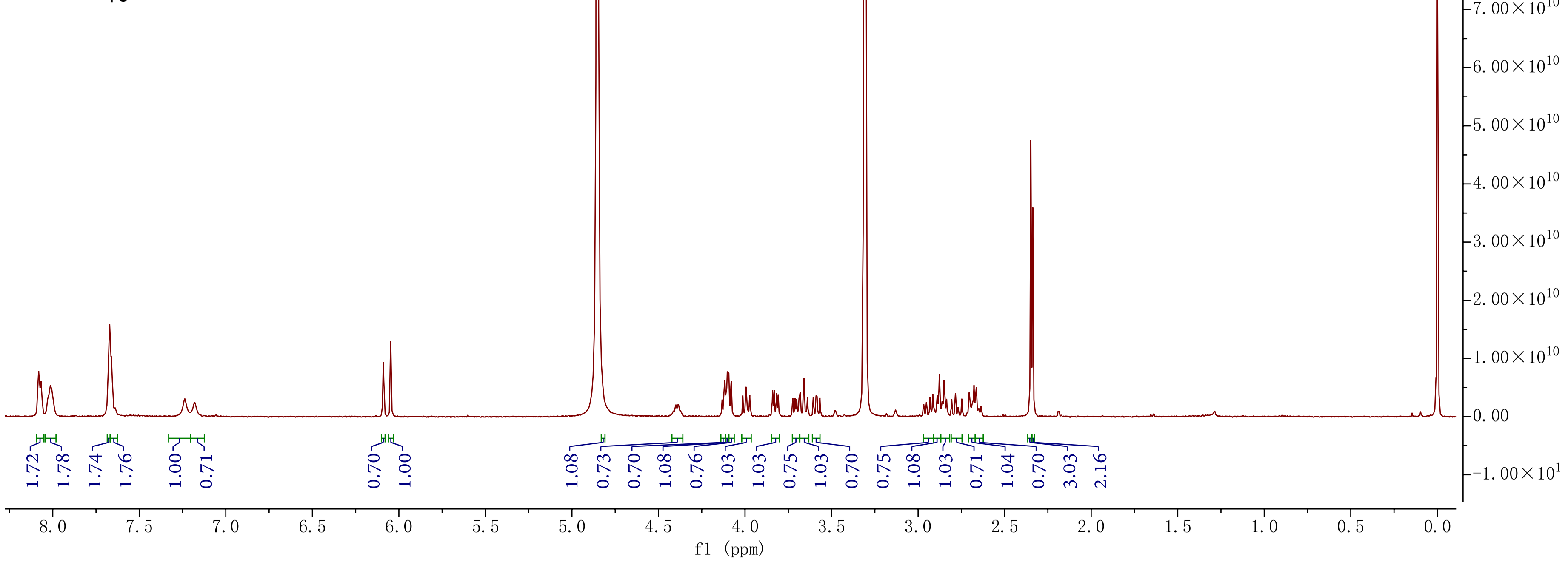

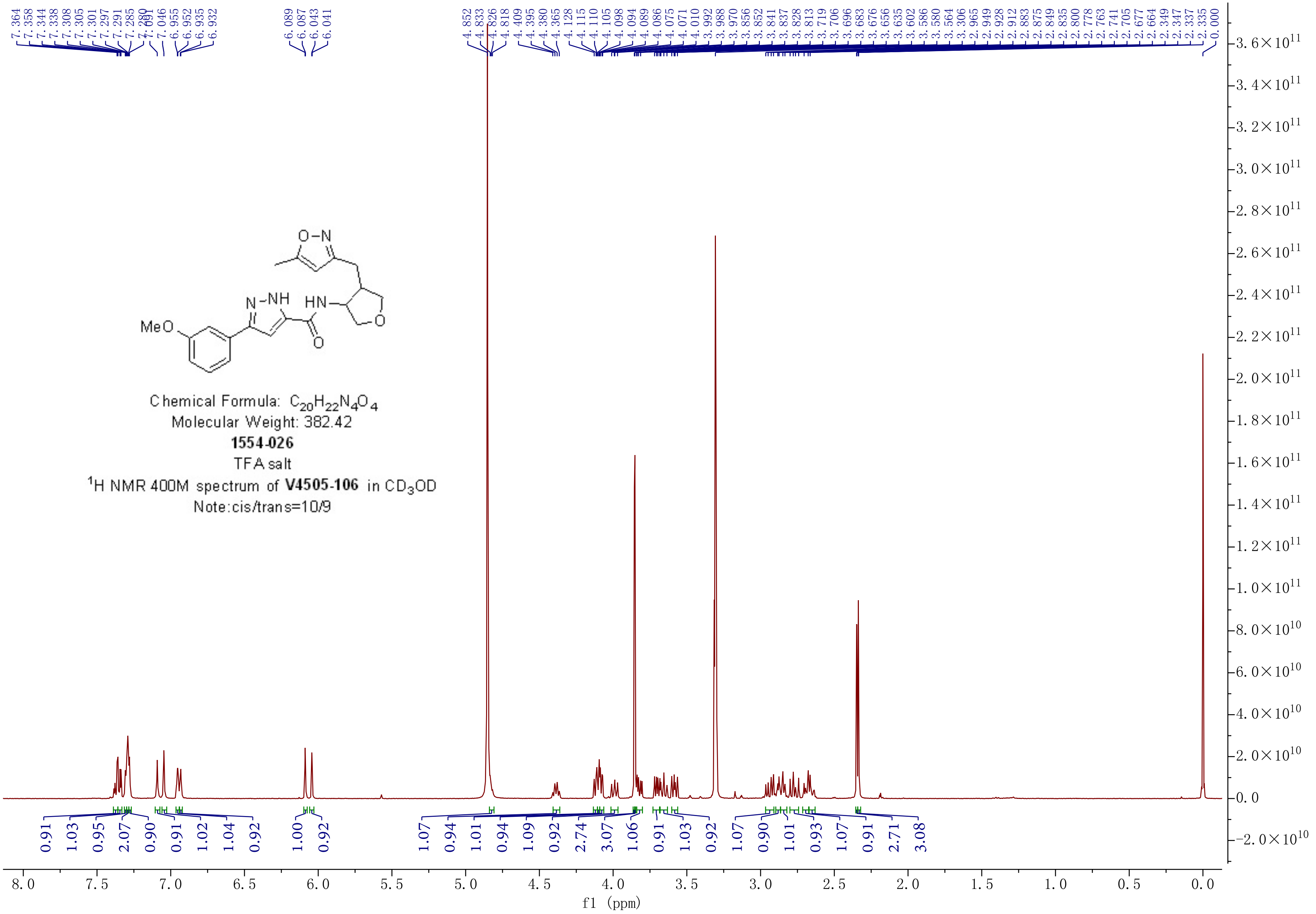
MeO
N-NH
HN
O-N
O
Chemical Formula: $C_{20}H_{22}N_4O_4$
Molecular Weight: 382.42
1554-026
TFA salt
$^1$H NMR 400M spectrum of V4505-106 in $CD_3OD$
Note:cis/trans=10/9
f1 (ppm)

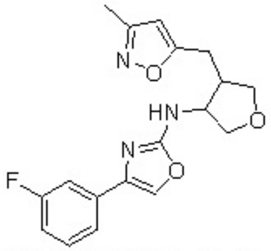


Chemical Formula: $C_{18}H_{18}FN_3O_3$
Molecular Weight: 343.36
**1554-027**
$^1$H NMR 400M spectrum of **V4505-140** in $CD_3OD$
Note:cis/trans=5/2

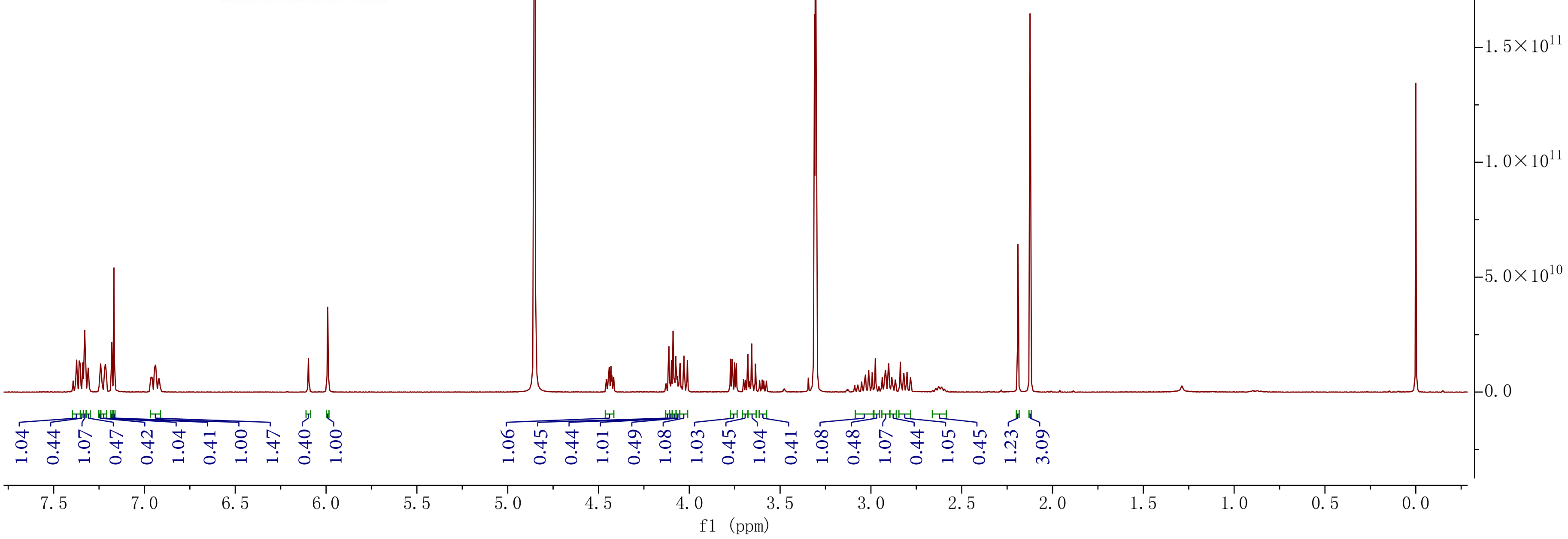

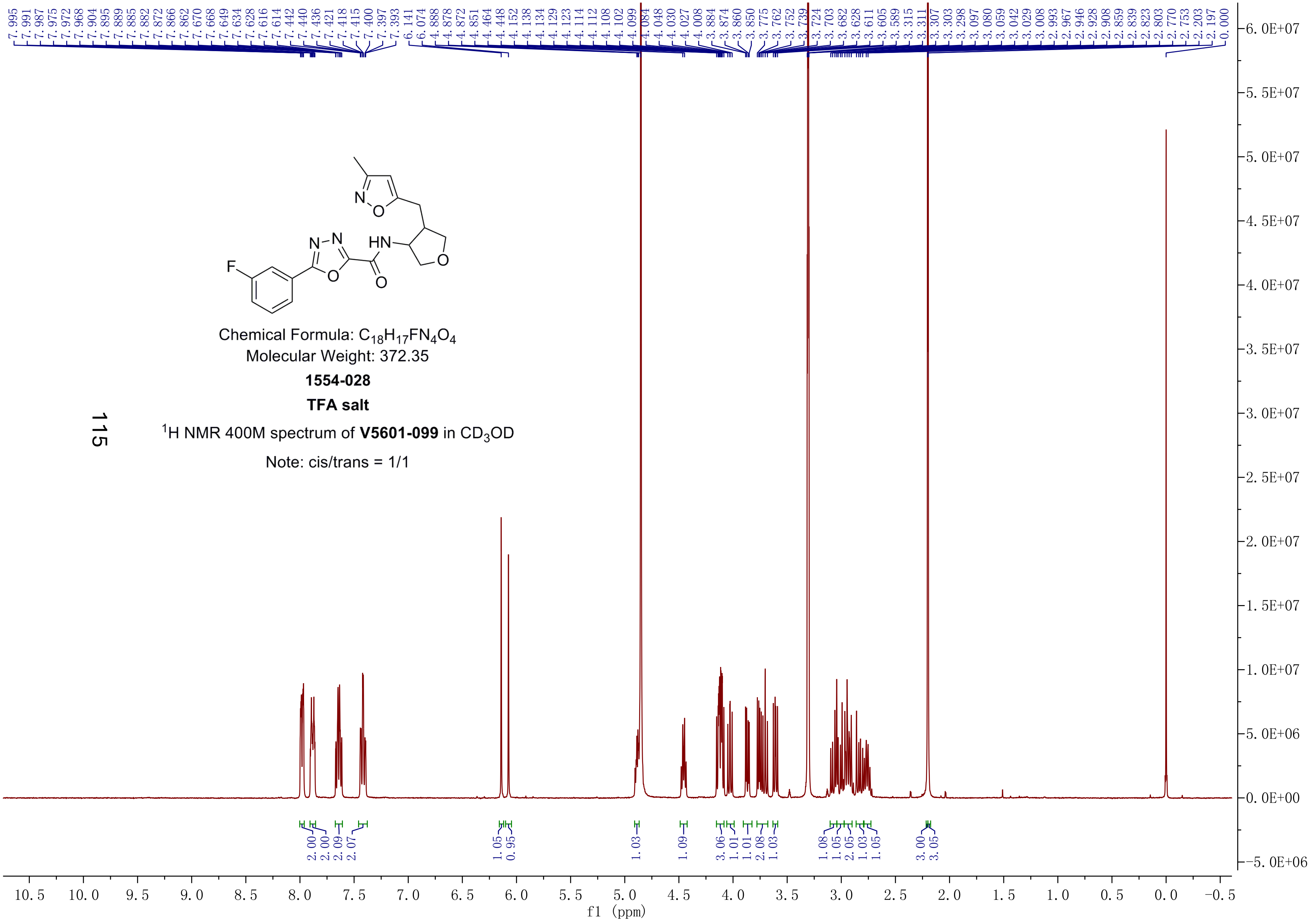

Chemical Formula: $C_{18}H_{17}FN_4O_4$
Molecular Weight: 372.35
**1554-028**
**TFA salt**
$^1$H NMR 400M spectrum of **V5601-099** in $CD_3OD$
Note: cis/trans = 1/1

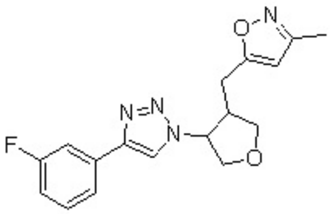

Chemical Formula: $C_{17}H_{17}FN_4O_2$
Molecular Weight: 328.35
**1554-029**
TFA salt
$^1H$ NMR 400M spectrum of **V4505-139** in $CD_3OD$
Note:cis/trans=5/4

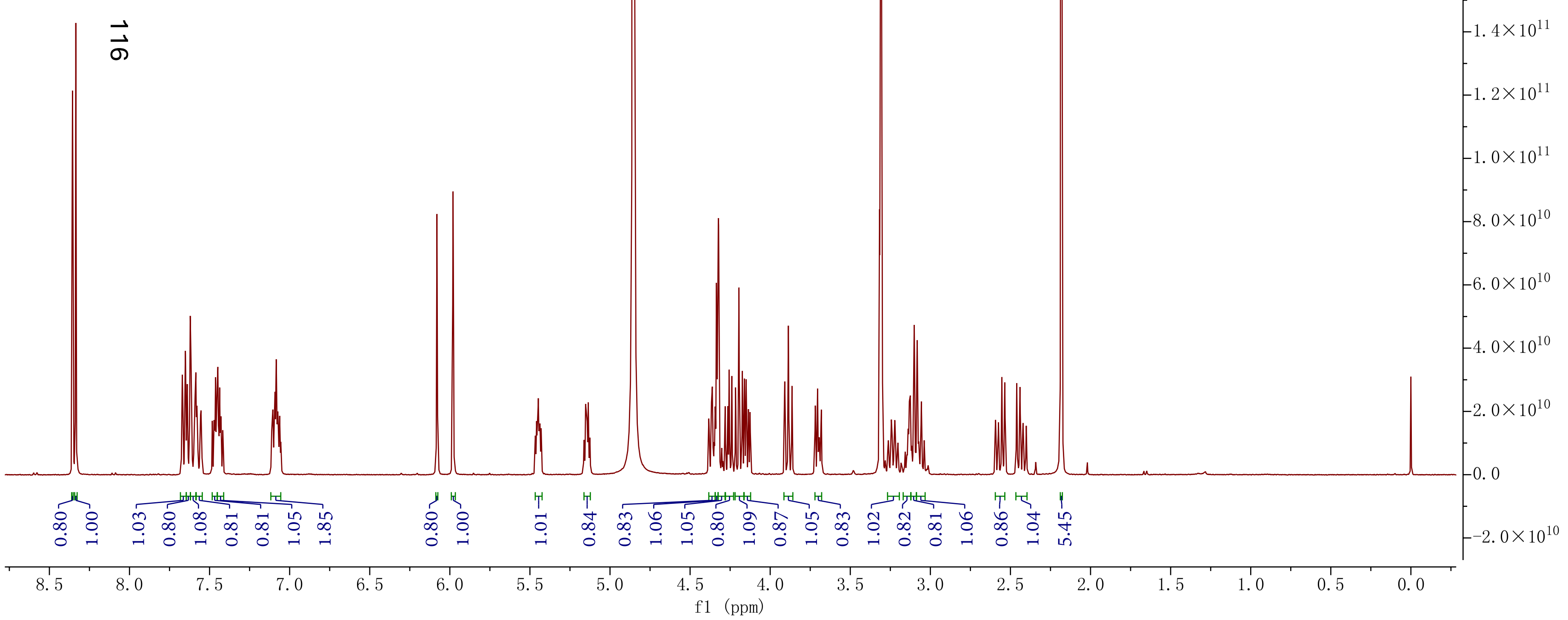

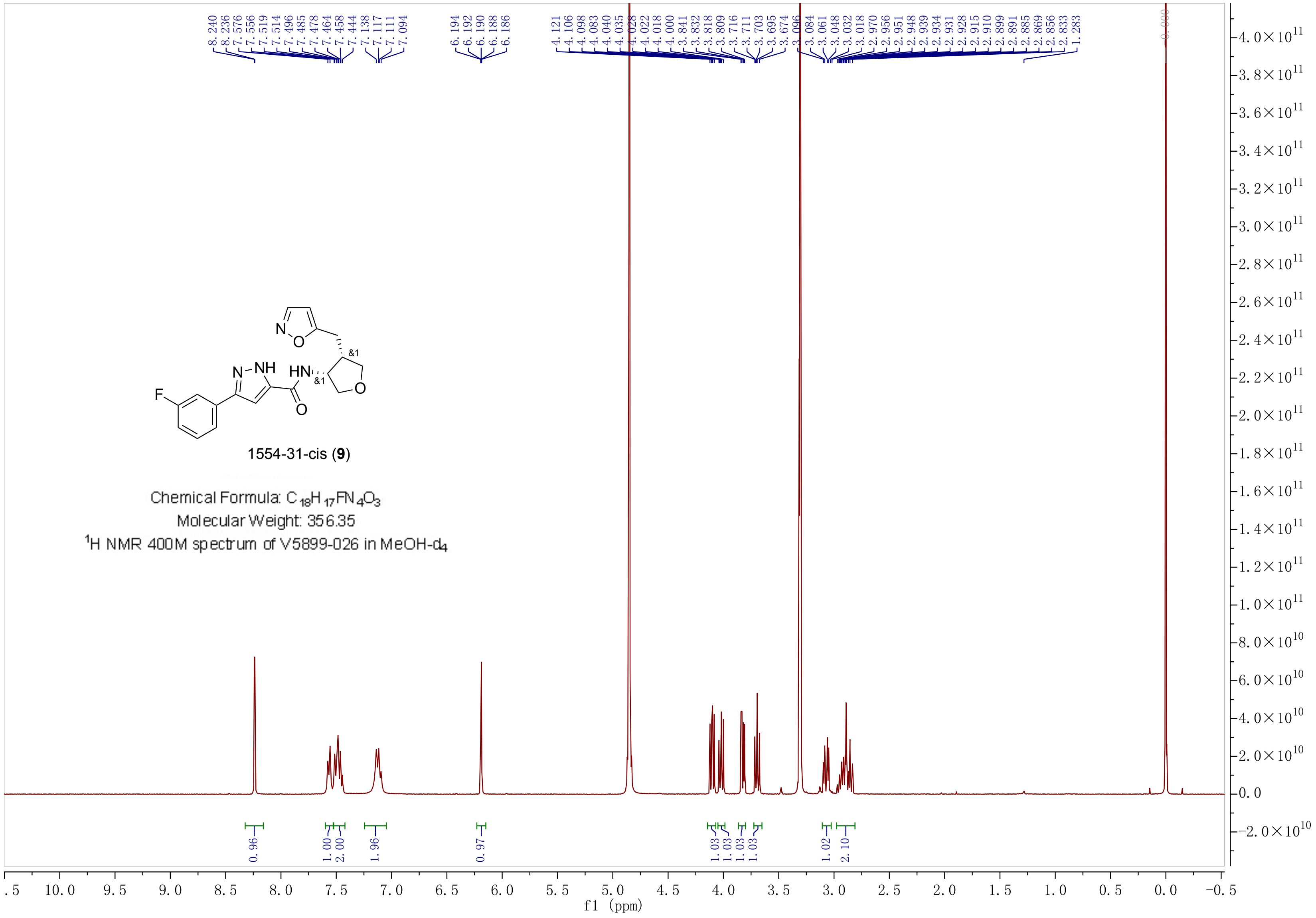


1554-31-cis (**9**)

Chemical Formula: $C_{18}H_{17}FN_4O_3$

Molecular Weight: 356.35

$^1$H NMR 400M spectrum of V5899-026 in MeOH-$d_4$

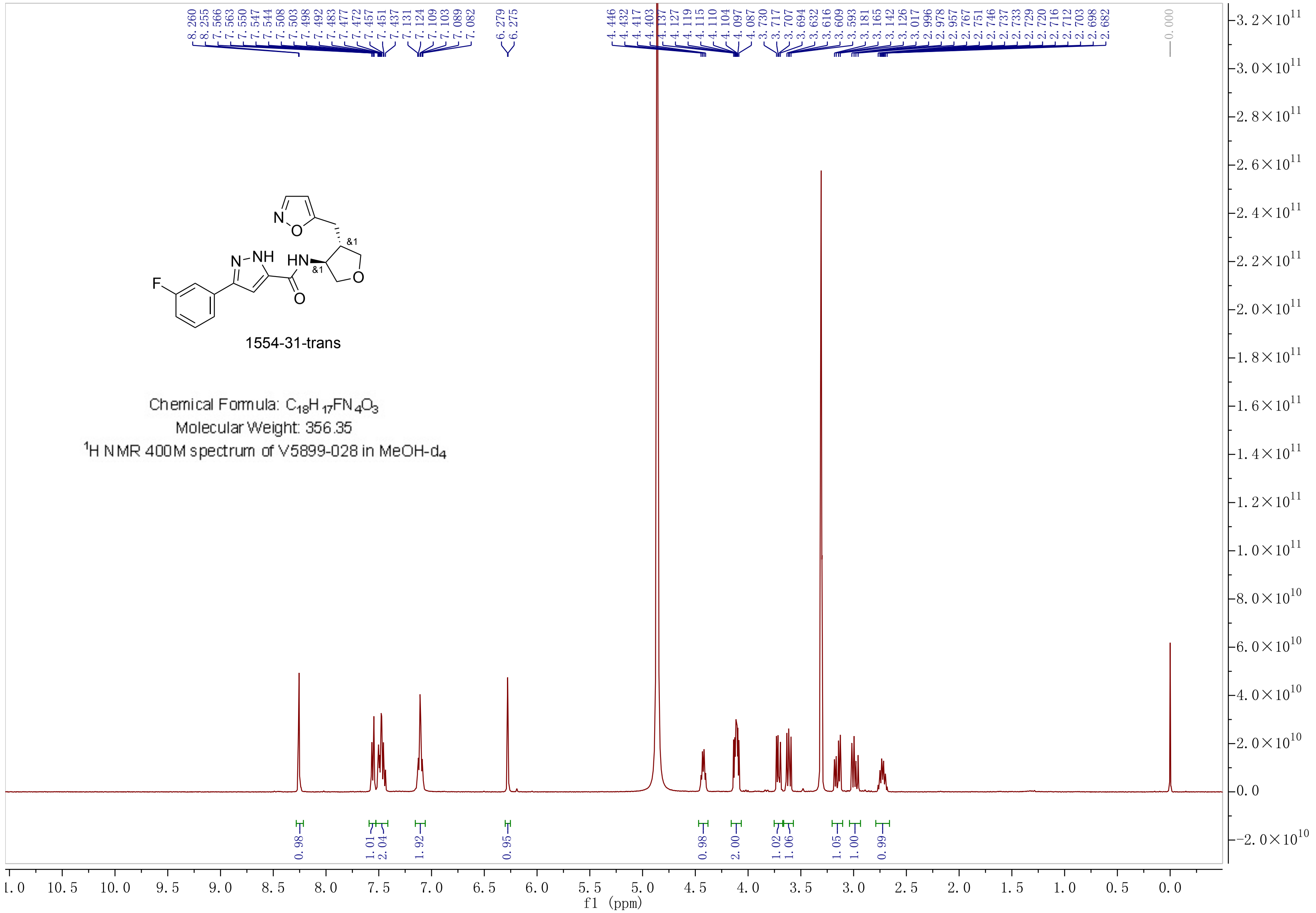
1554-31-trans
Chemical Formula: $C_{18}H_{17}FN_4O_3$
Molecular Weight: 356.35
$^1$H NMR 400M spectrum of V5899-028 in MeOH-d4
8.260 8.255 7.566 7.563 7.550 7.547 7.544 7.508 7.503 7.498 7.492 7.483 7.477 7.472 7.457 7.451 7.437 7.131 7.124 7.109 7.103 7.089 7.082
6.279 6.275
4.446 4.432 4.417 4.403 4.137 4.127 4.119 4.115 4.110 4.104 4.097 4.087 3.730 3.717 3.707 3.694 3.632 3.616 3.609 3.593 3.181 3.165 3.142 3.126 3.017 2.996 2.978 2.957 2.767 2.751 2.746 2.737 2.733 2.729 2.720 2.716 2.712 2.703 2.698 2.682
0.000
0.98 1.01 2.04 1.92 0.95 0.98 2.00 1.02 1.06 1.05 1.00 0.99
f1 (ppm)

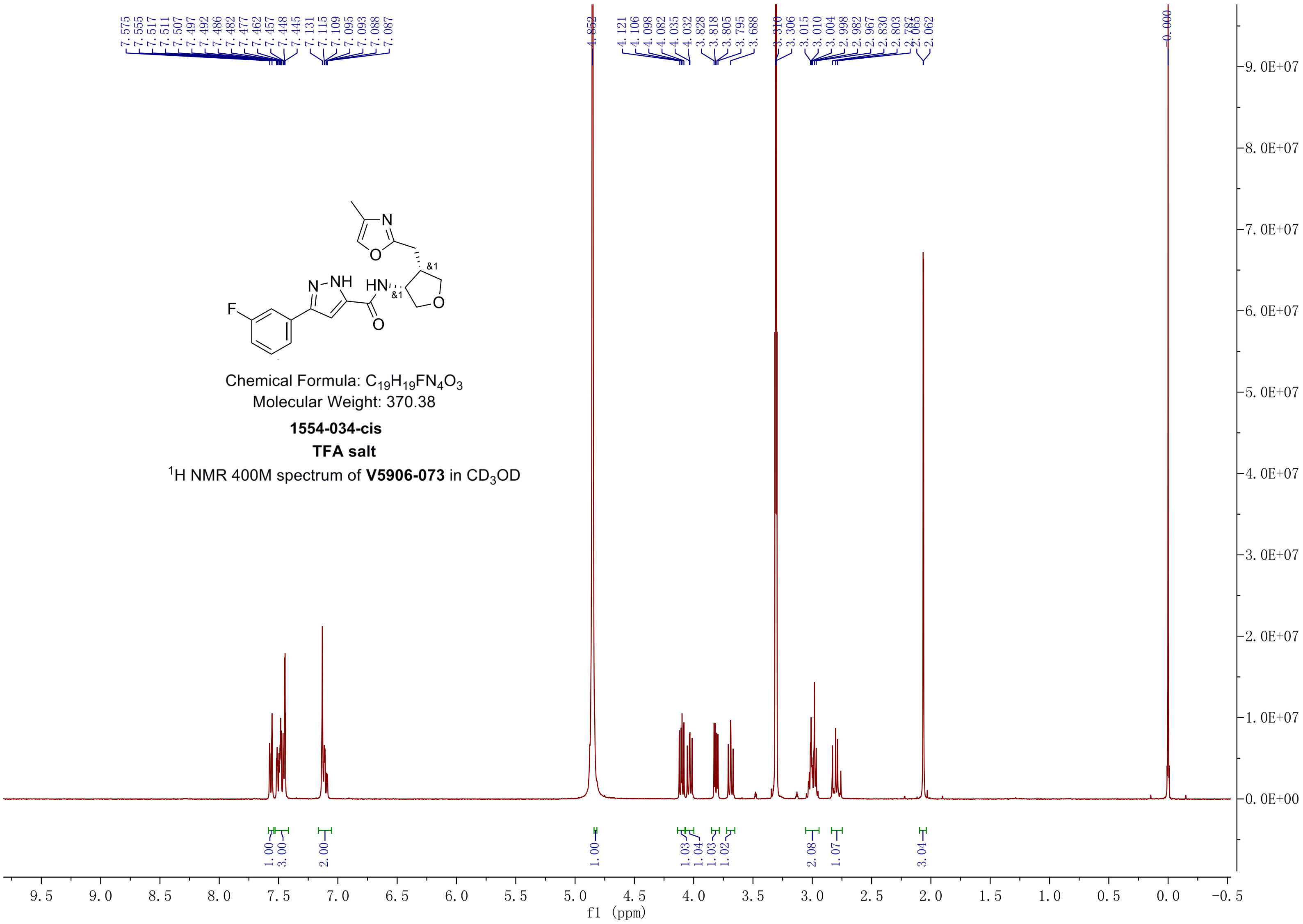


Chemical Formula: $C_{19}H_{19}FN_4O_3$
Molecular Weight: 370.38

**1554-034-cis**

**TFA salt**

$^1$H NMR 400M spectrum of **V5906-073** in $CD_3OD$

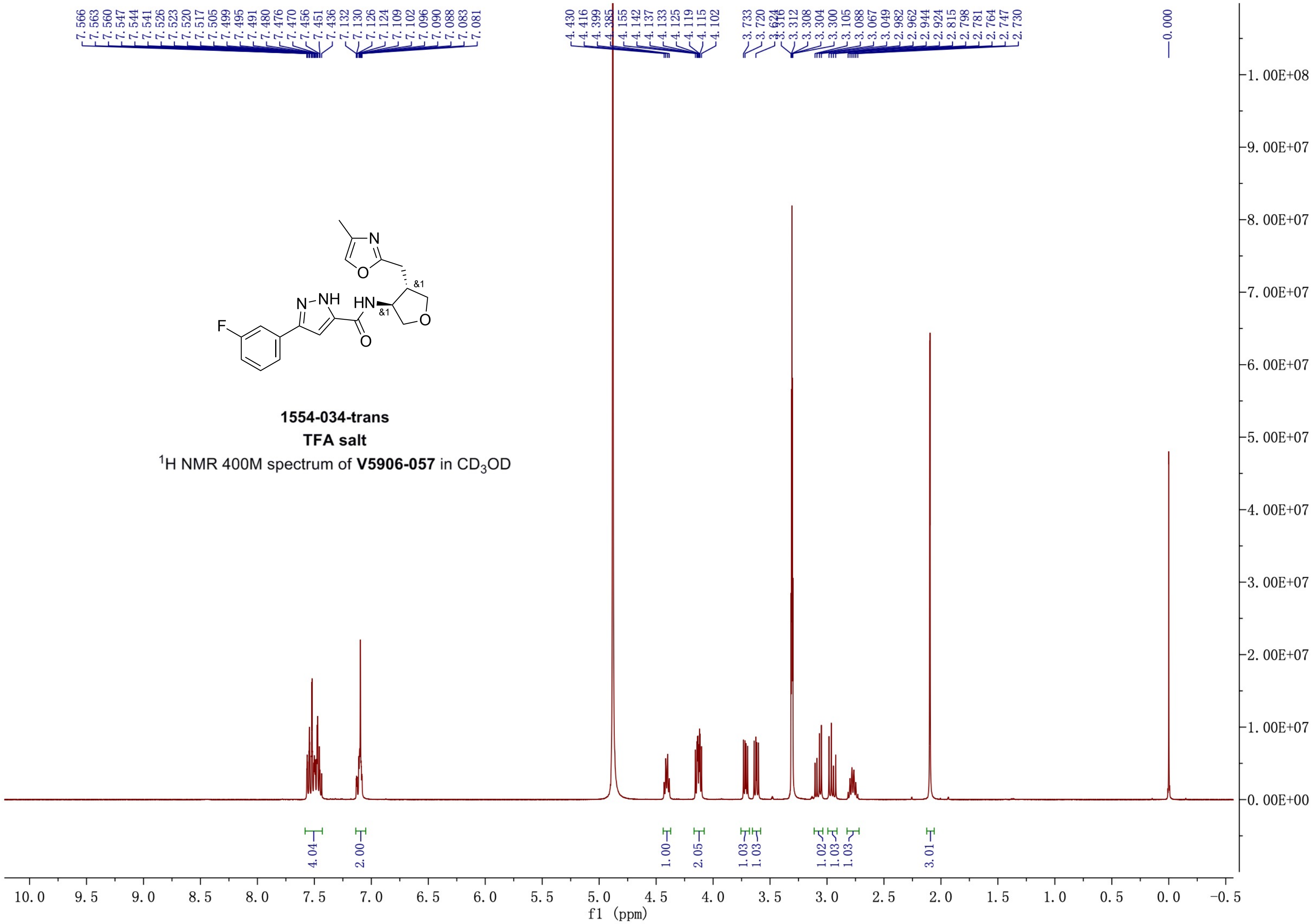


1554-034-trans
TFA salt
$^1$H NMR 400M spectrum of **V5906-057** in $CD_3OD$

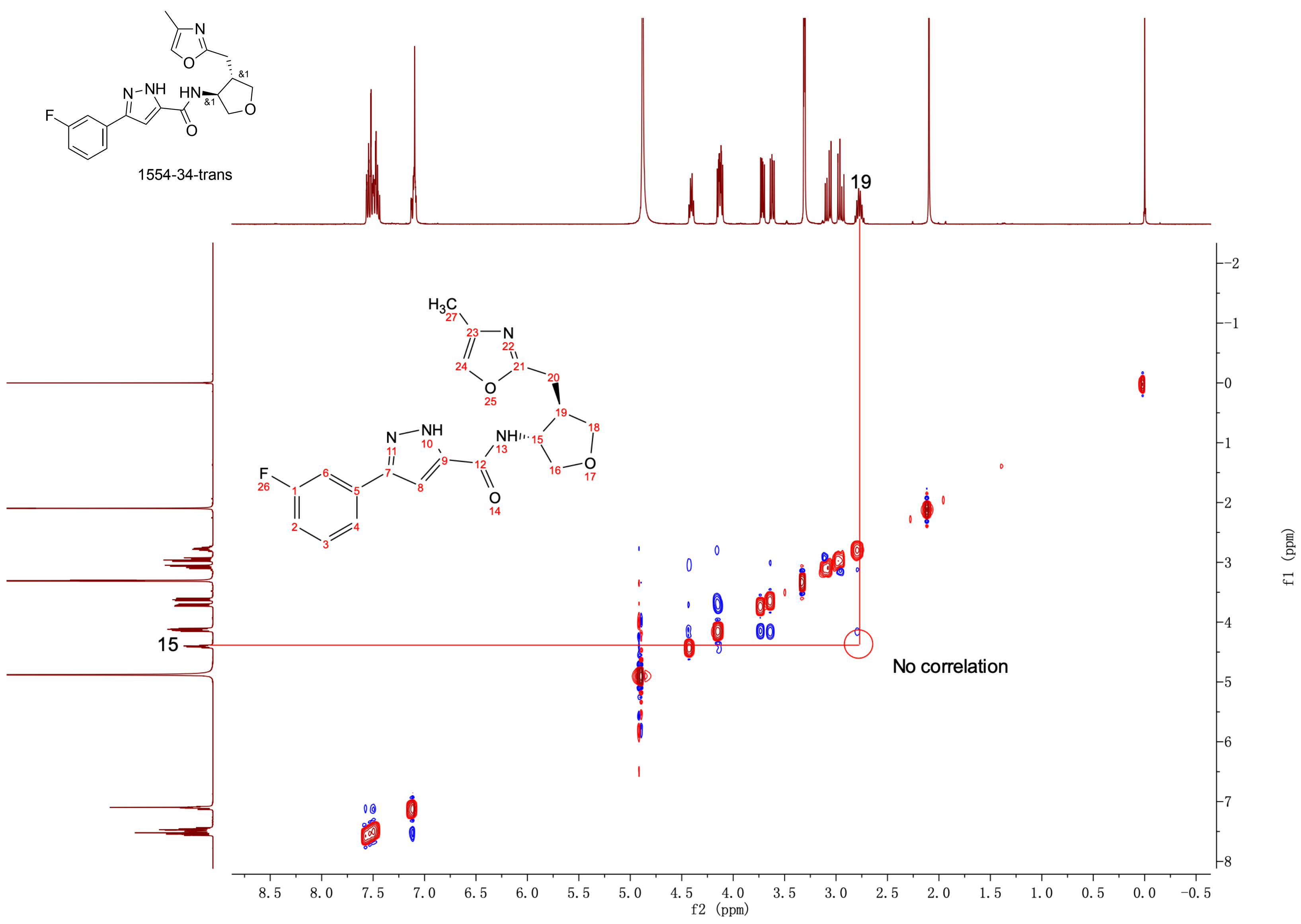
1554-34-trans
19
15
No correlation
f2 (ppm)
f1 (ppm)

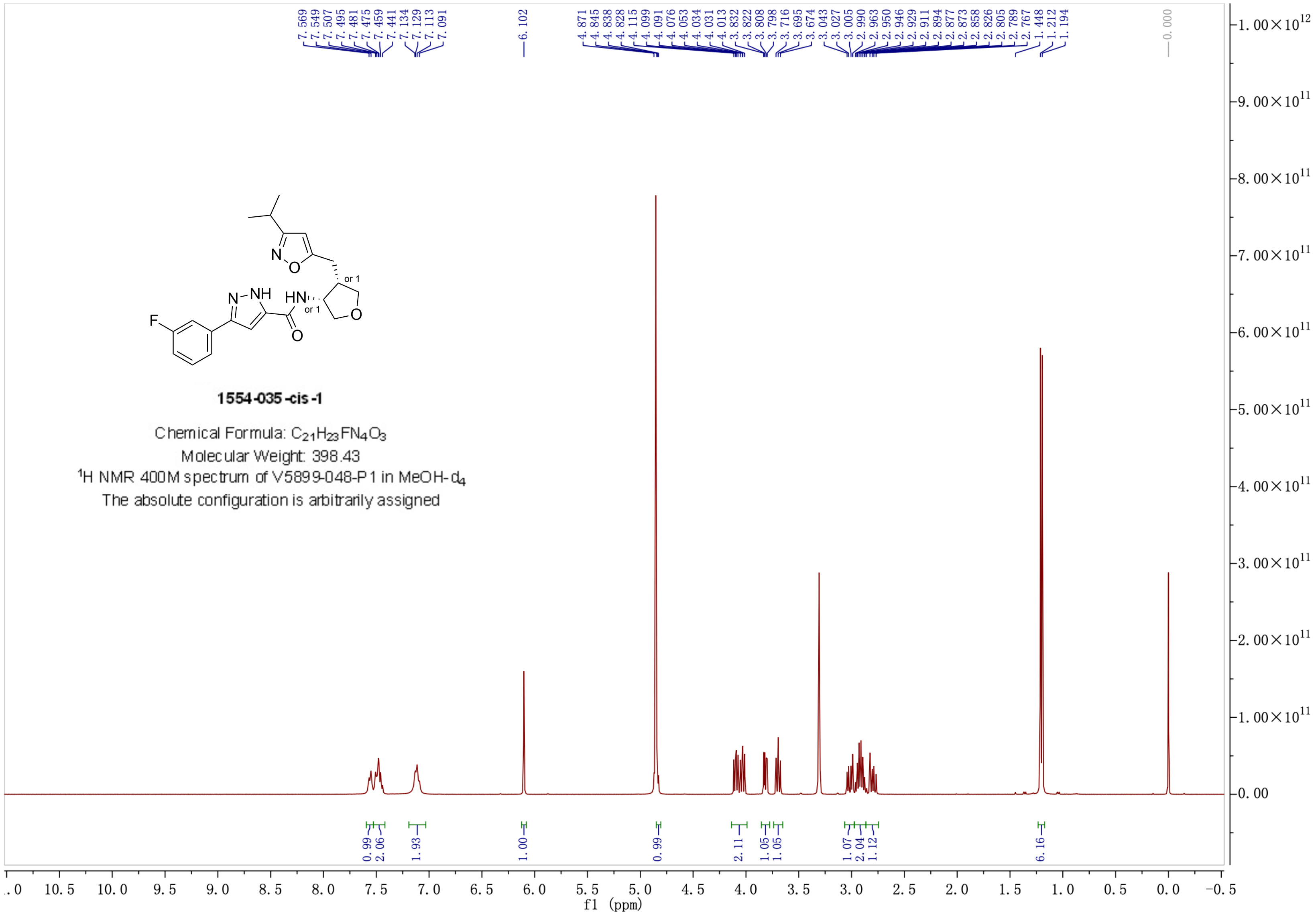


**1554-035-cis-1**

Chemical Formula: $C_{21}H_{23}FN_4O_3$

Molecular Weight: 398.43

$^1H$ NMR 400M spectrum of V5899-048-P1 in MeOH-$d_4$

The absolute configuration is arbitrarily assigned

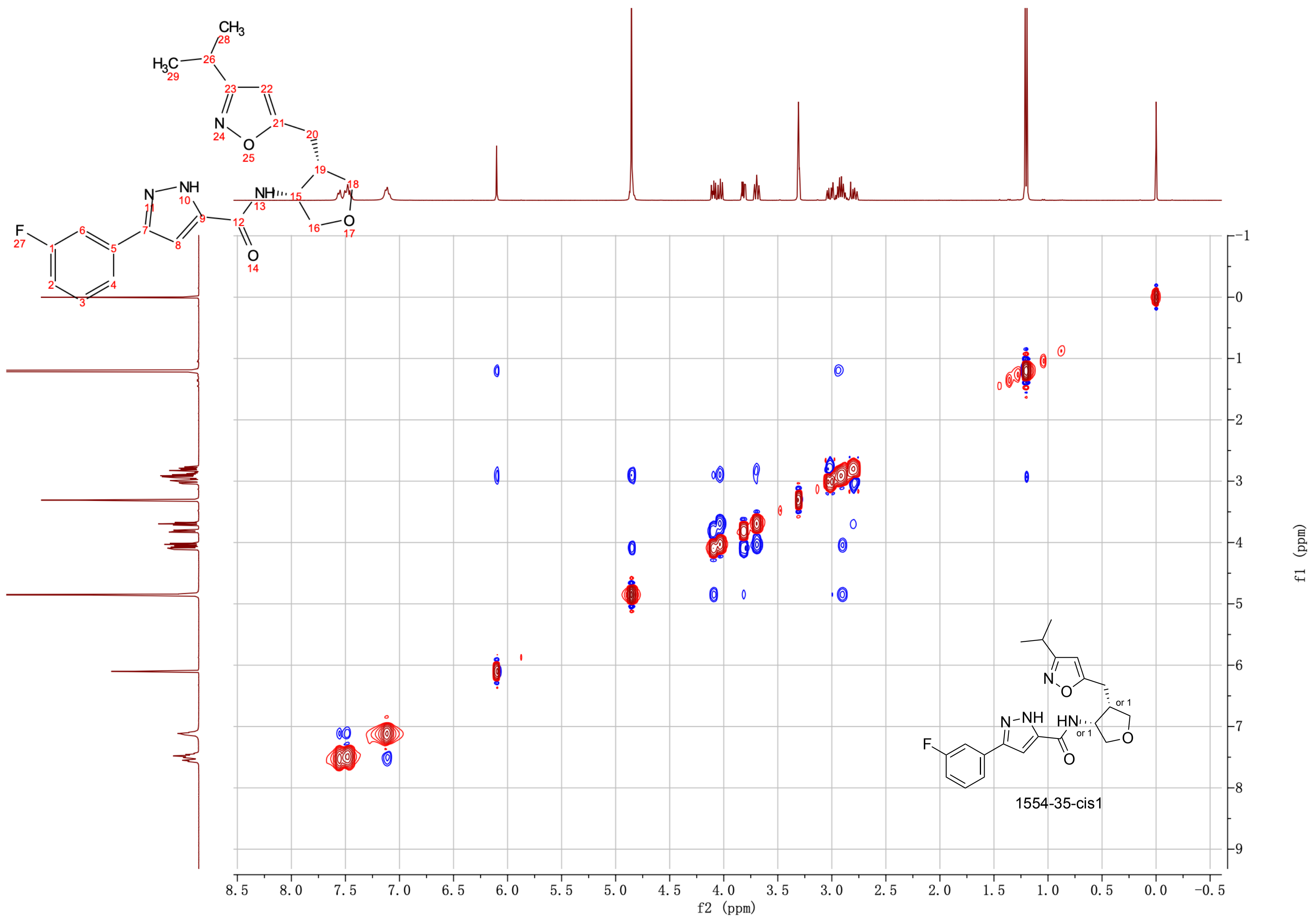
1554-35-cis1
f2 (ppm)
f1 (ppm)

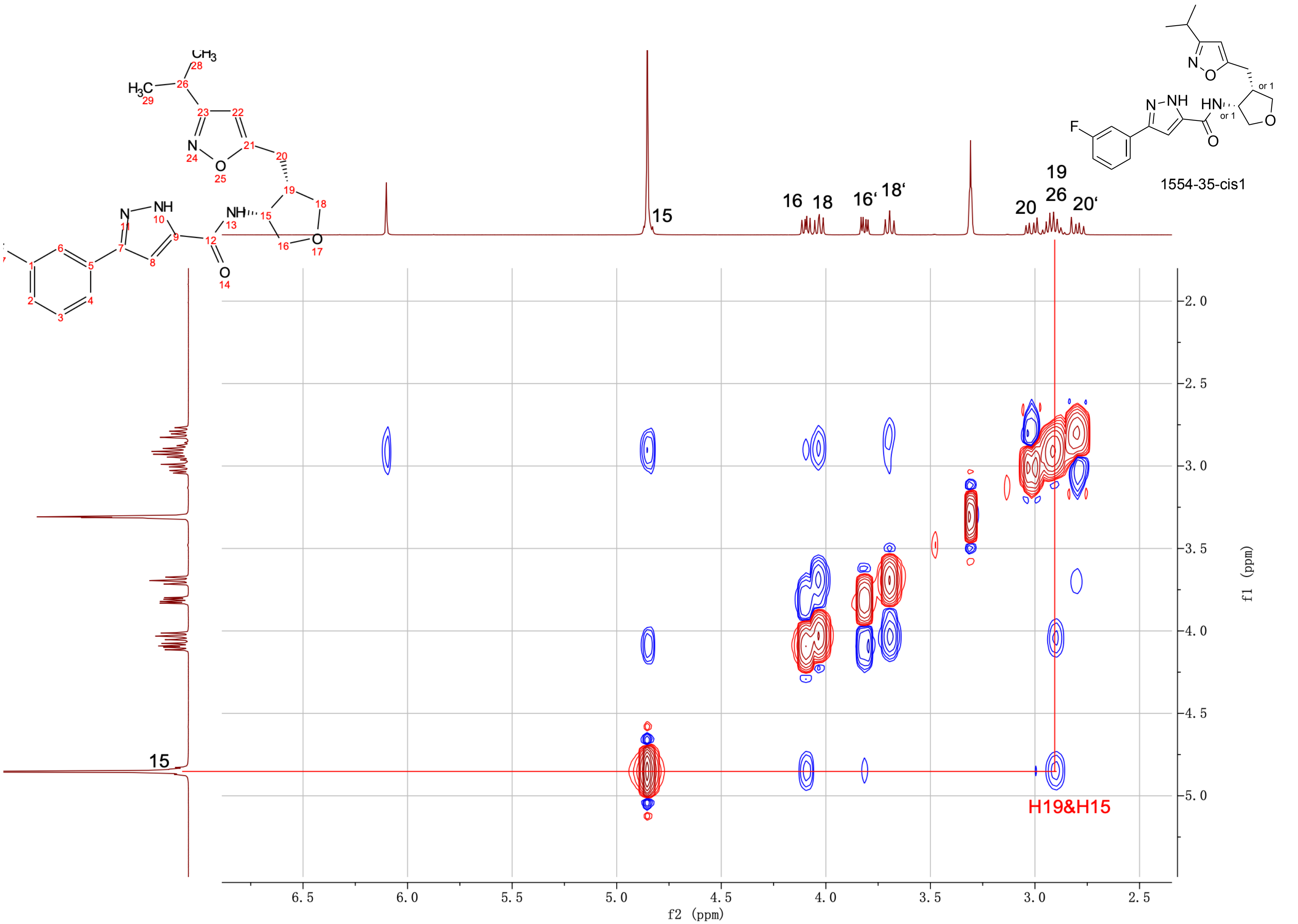

1554-35-cis1
16 18 16' 18'
15
20 19 26 20'
H19&H15
f2 (ppm)
f1 (ppm)

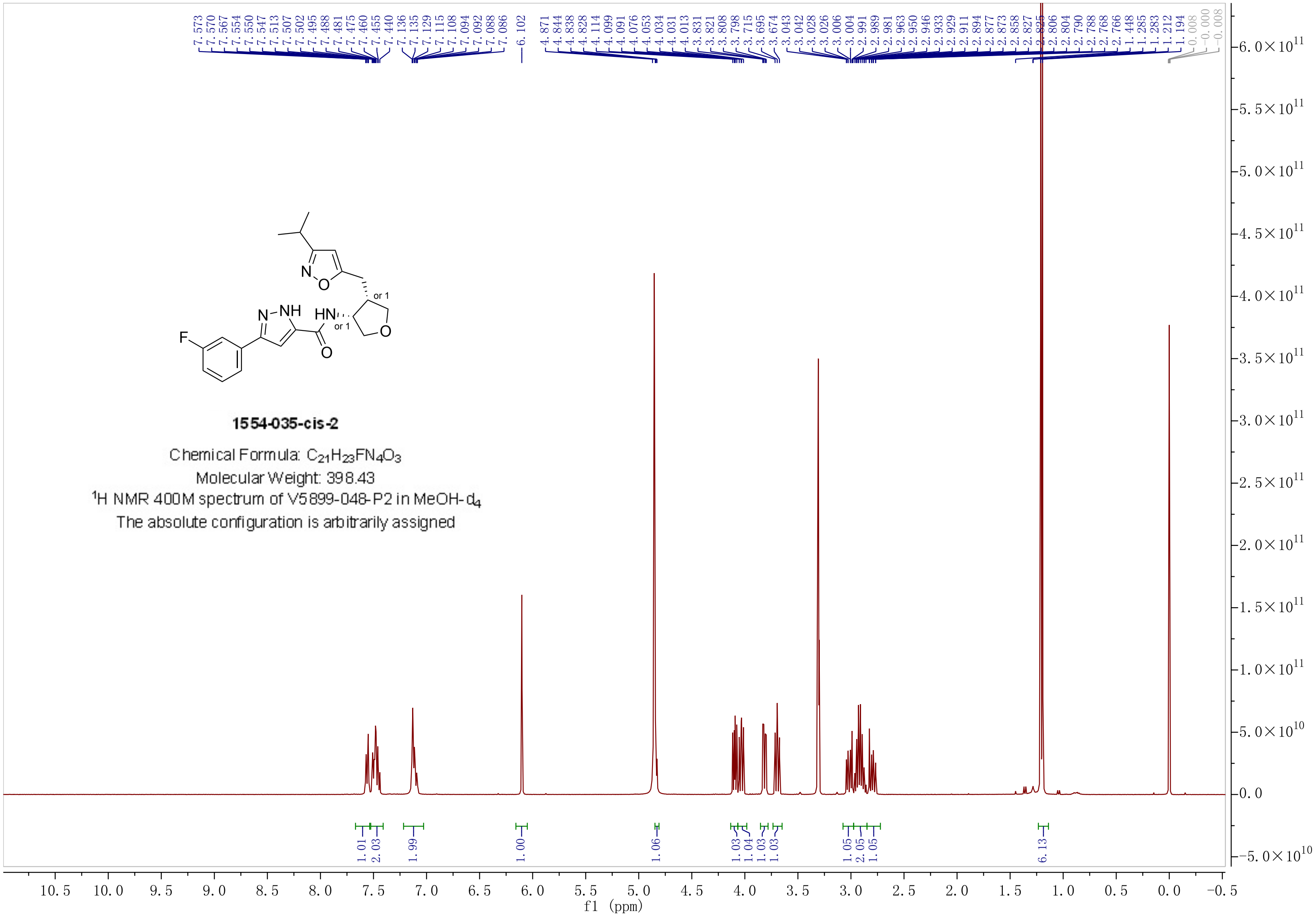


**1554-035-cis-2**

Chemical Formula: $C_{21}H_{23}FN_4O_3$

Molecular Weight: 398.43

$^1$H NMR 400M spectrum of V5899-048-P2 in MeOH-$d_4$

The absolute configuration is arbitrarily assigned

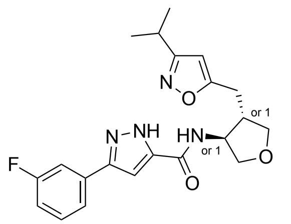


**1554-035-trans-1**

Chemical Formula: $C_{21}H_{23}FN_4O_3$
Molecular Weight: 398.43
$^1$H NMR 400M spectrum of V5899-048-P3 in MeOH-$d_4$
The absolute configuration is arbitrarily assigned

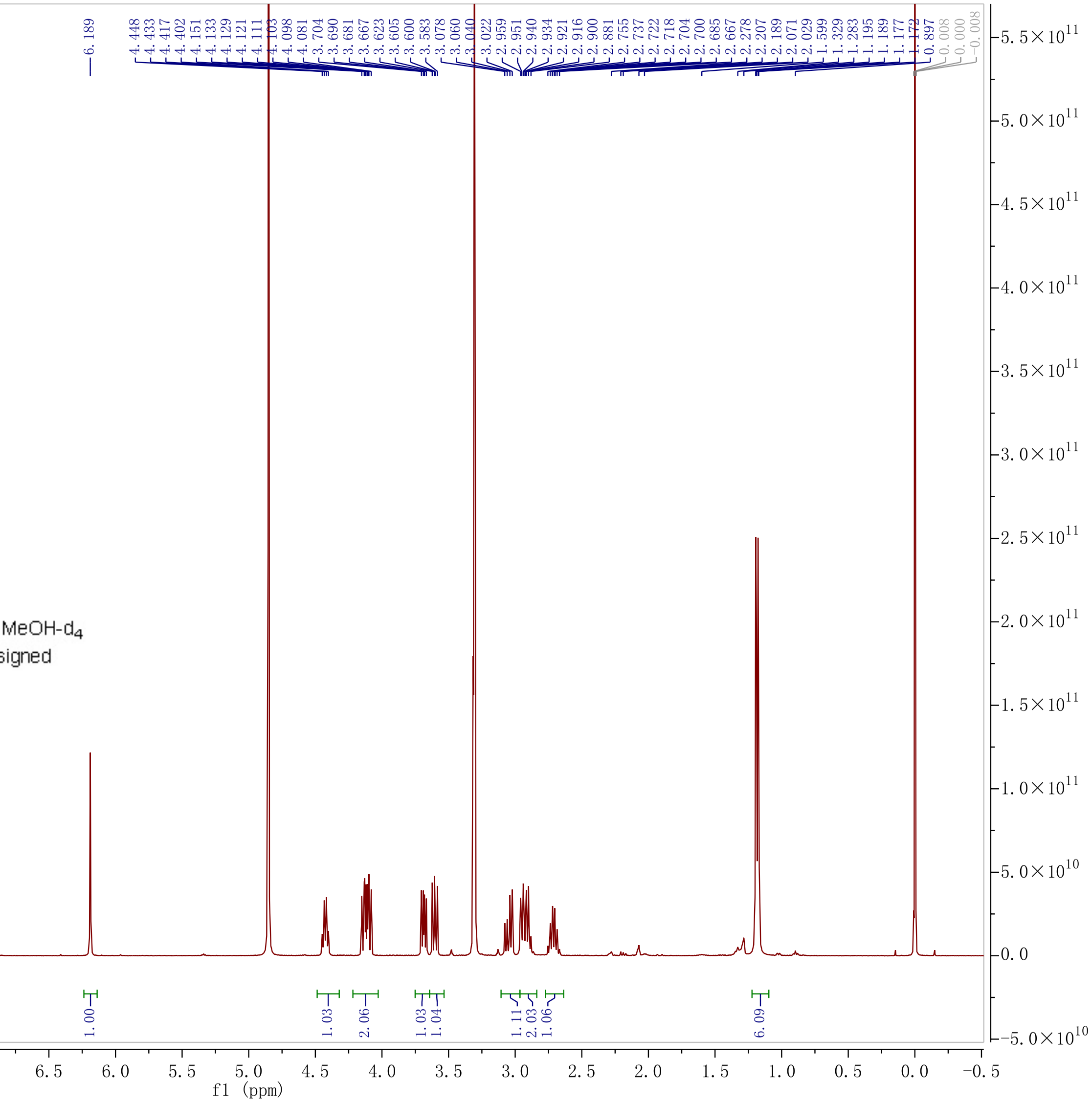

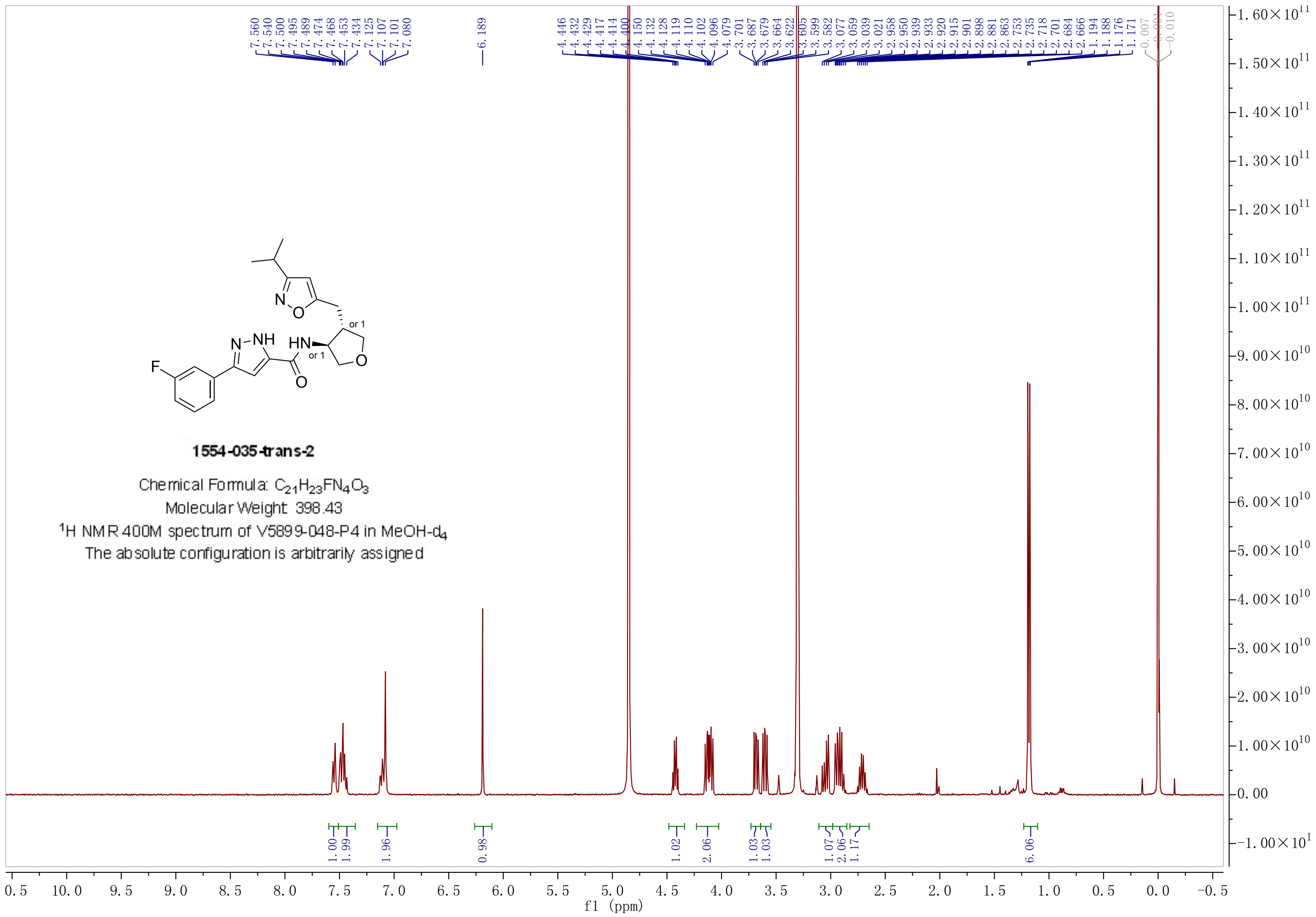

**1554-035-trans-2**

Chemical Formula: $C_{21}H_{23}FN_4O_3$

Molecular Weight: 398.43

$^1$H NMR 400M spectrum of V5899-048-P4 in MeOH-$d_4$

The absolute configuration is arbitrarily assigned

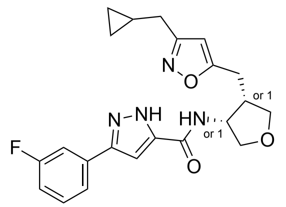

**1554-036-cis-1**

Chemical Formula: $C_{22}H_{23}FN_4O_3$

Molecular Weight: 410.44

$^1$H NMR 400M spectrum of V5899-057-P1 in MeOH-$d_4$

The absolute configuration is arbitrarily assigned

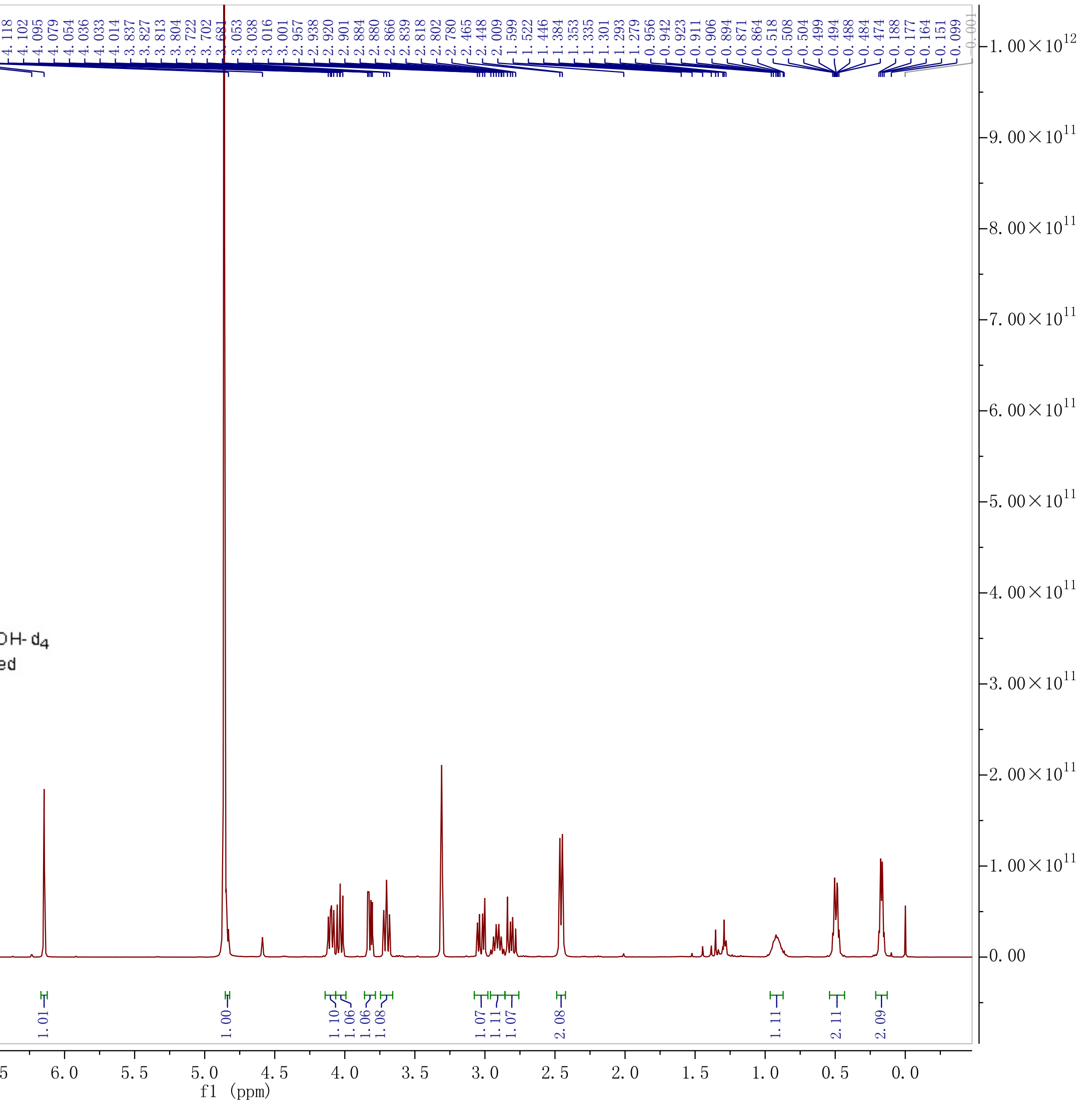

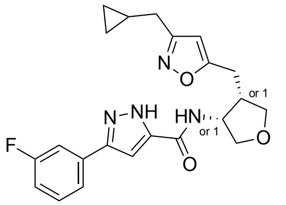

**1554-036-cis-2**

Chemical Formula: $C_{22}H_{23}FN_4O_3$
Molecular Weight: 410.44
$^1$H NMR 400M spectrum of V5899-057-P2 in MeOH-$d_4$
The absolute configuration is arbitrarily assigned

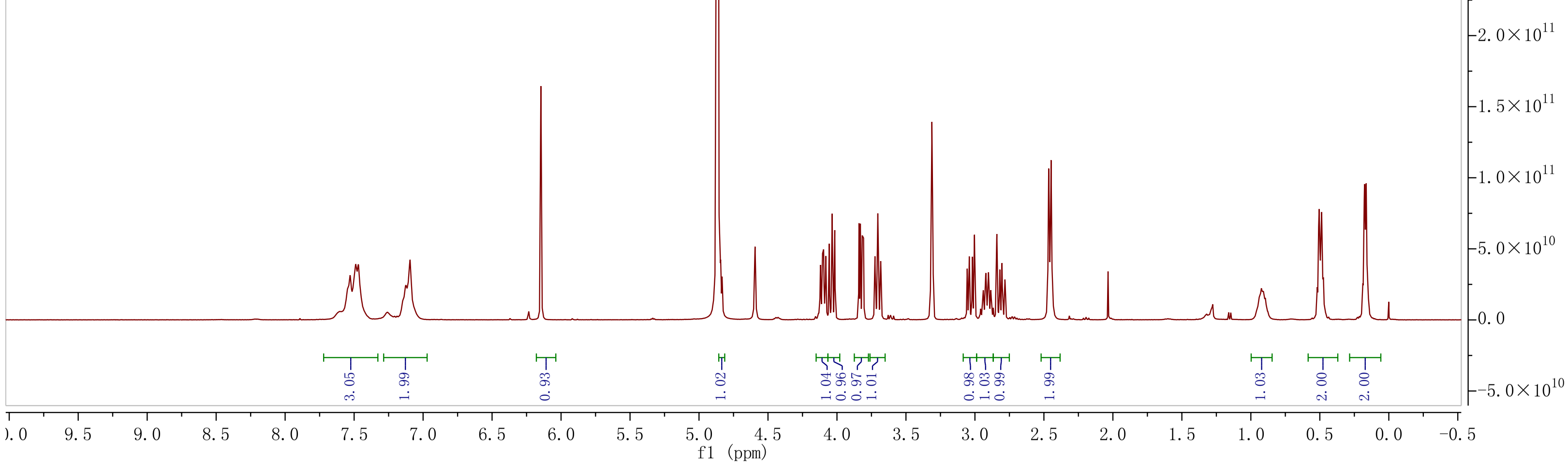

**1554-036-trans-1**

Chemical Formula: $C_{22}H_{23}FN_4O_3$

Molecular Weight: 410.44

**TFA Salt**

$^1$H NMR 400M spectrum of V5899-057-P3 in MeOH-$d_4$

The absolute configuration is arbitrarily assigned

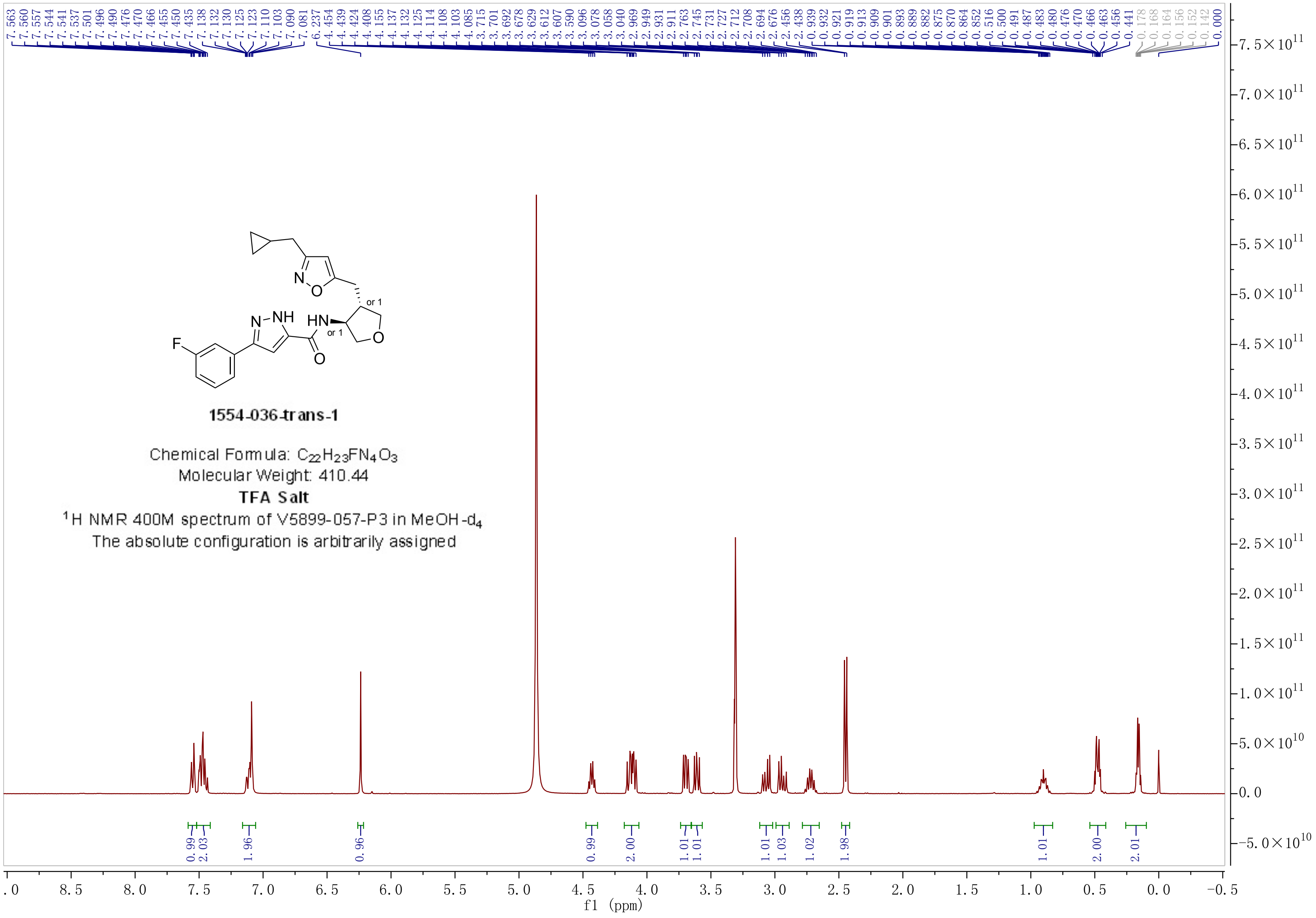

### SAMPLE INFORMATION

| | | | |
|---|---|---|---|
| Sample Name: | V5899-007-P1 | | |
| Acq. Method Set: | AD_55%_B2 | Sample Set Name: | 20240715 |
| Co_Solvent: | EtOH[1%NH3(7M in MeOH)] | Processing Method: | VIVA CRO |
| Column Name: | AD-3 4.6*100mm 3um | Injection #: | 1 |
| Vial: | 1:A,4 | Back Pressure: | 2000 psi |
| Flow Rate: | 3.0 mL/min | Injection Volume: | 10.00 ul |
| Channel Name: | PDA Ch1 MaxPlot(210-400)nm | Run Time: | 6.0 Minutes |
| Proc. Chnl. Descr.: | PDA Ch1 MaxPlot(210-400)nm | Column Temperature: | 40oC |
| Date Acquired: | 7/16/2024 3:01:56 AM CST | Date Processed: | 7/16/2024 10:30:28 AM |

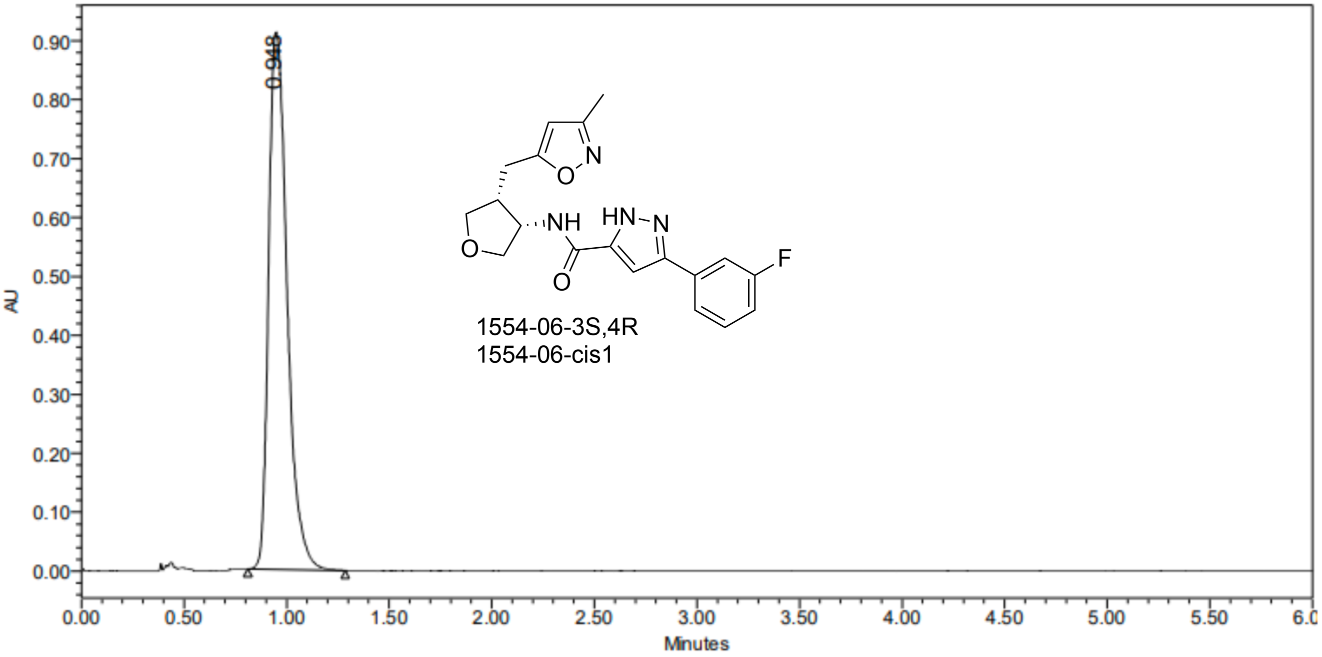


| | RT | Area | % Area | Height |
|---|---|---|---|---|
| 1 | 0.948 | 5701637 | 100.00 | 911728 |

### SAMPLE INFORMATION

| | | | |
|---|---|---|---|
| Sample Name: | V5899-007-P2 | | |
| Acq. Method Set: | AD_55%_B2 | Sample Set Name: | 20240715 |
| Co_Solvent: | EtOH[1%NH3(7M in MeOH)] | Processing Method: | VIVA CRO |
| Column Name: | AD-3 4.6*100mm 3um | Injection #: | 1 |
| Vial: | 1:A,5 | Back Pressure: | 2000 psi |
| Flow Rate: | 3.0 mL/min | Injection Volume: | 10.00 ul |
| Channel Name: | PDA Ch1 MaxPlot(210-400)nm | Run Time: | 6.0 Minutes |
| Proc. Chnl. Descr.: | PDA Ch1 MaxPlot(210-400)nm | Column Temperature: | 40oC |
| Date Acquired: | 7/16/2024 3:09:05 AM CST | Date Processed: | 7/16/2024 10:30:39 AM |

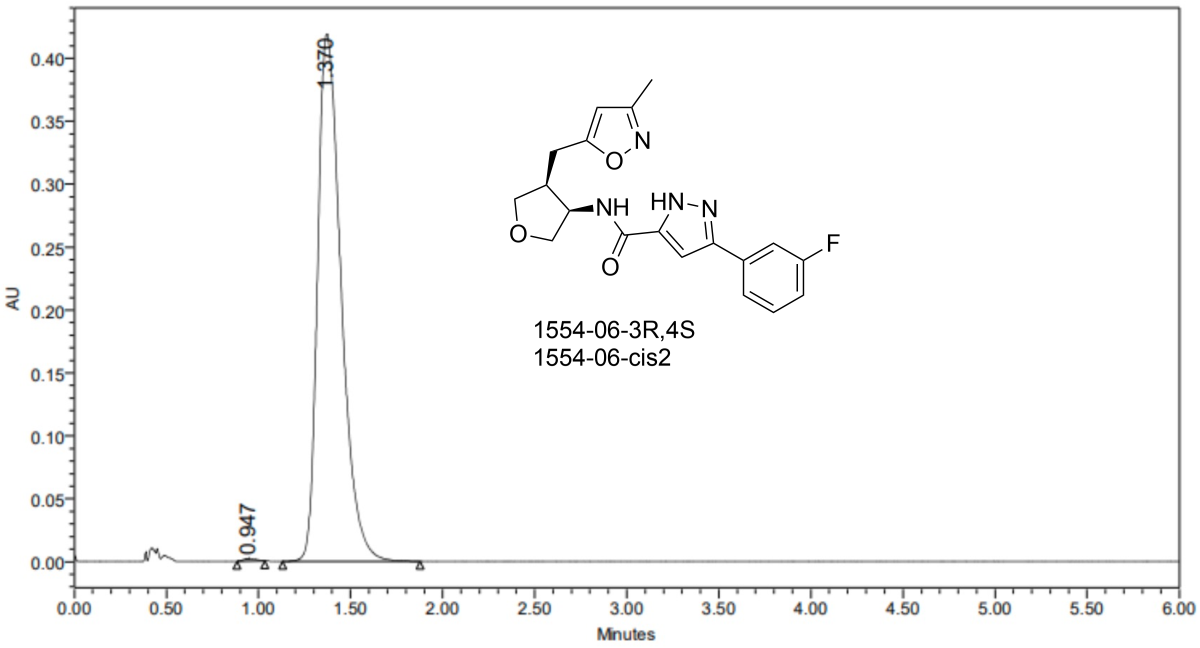


| | RT | Area | % Area | Height |
|---|---|---|---|---|
| 1 | 0.947 | 9642 | 0.26 | 2094 |
| 2 | 1.370 | 3725270 | 99.74 | 418832 |

### SAMPLE INFORMATION

| | | | |
|---|---|---|---|
| Sample Name: | V5899-007-trans-P1 | | |
| Acq. Method Set: | WHELK_35%_B1 | Sample Set Name: | 20240822 |
| Co_Solvent: | MeOH[0.2%NH3(7M in MeOH)] | Processing Method: | VIVA CRO |
| Column Name: | (R,R)Whelk-O1 4.6*100mm 3.5um | Injection #: | 1 |
| Vial: | 2:A,5 | Back Pressure: | 2000 psi |
| Flow Rate: | 3.0 mL/min | Injection Volume: | 10.00 ul |
| Channel Name: | PDA Ch1 MaxPlot(210-400)nm | Run Time: | 2.0 Minutes |
| Proc. Chnl. Descr.: | PDA Ch1 MaxPlot(210-400)nm | Column Temperature: | 40oC |
| Date Acquired: | 8/22/2024 11:16:30 AM CST | Date Processed: | 8/22/2024 12:41:34 PM |

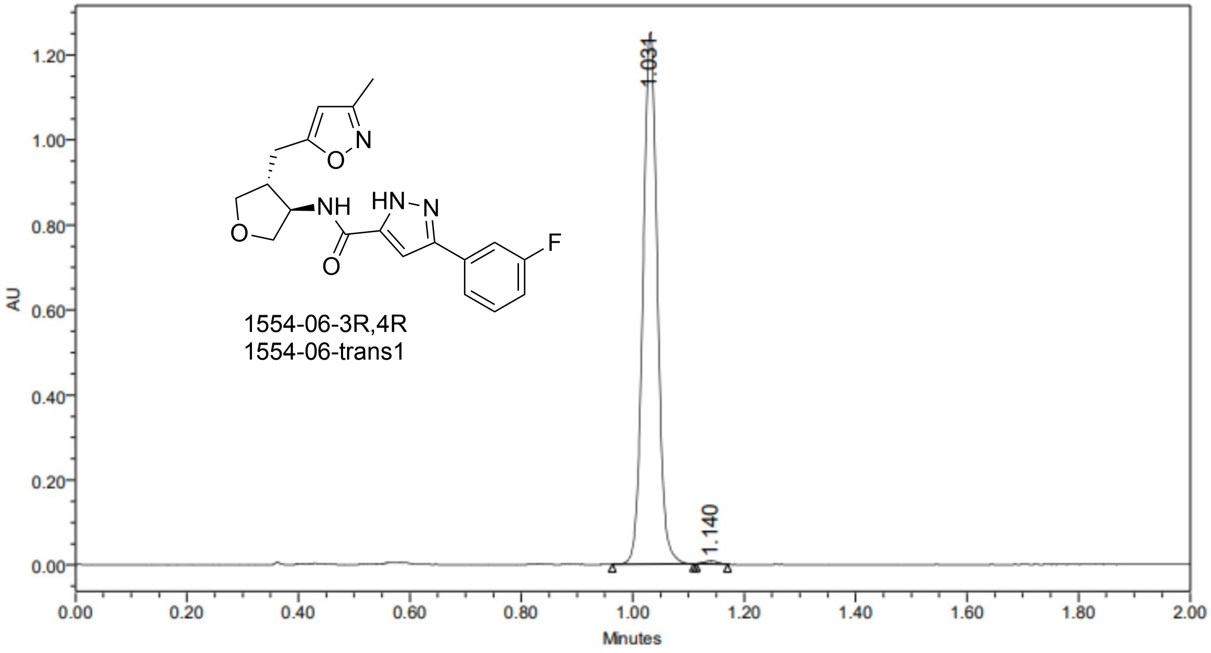


| | RT | Area | % Area | Height |
|---|---|---|---|---|
| 1 | 1.031 | 2241586 | 99.48 | 1251175 |
| 2 | 1.140 | 11811 | 0.52 | 6836 |

### SAMPLE INFORMATION

| | | | |
|---|---|---|---|
| Sample Name: | V5899-007-trans-P2 | | |
| Acq. Method Set: | WHELK_35%_B1 | Sample Set Name: | 20240822 |
| Co_Solvent: | MeOH[0.2%NH3(7M in MeOH)] | Processing Method: | VIVA CRO |
| Column Name: | (R,R)Whelk-O1 4.6*100mm 3.5um | Injection #: | 1 |
| Vial: | 2:A,6 | Back Pressure: | 2000 psi |
| Flow Rate: | 3.0 mL/min | Injection Volume: | 10.00 ul |
| Channel Name: | PDA Ch1 MaxPlot(210-400)nm | Run Time: | 2.0 Minutes |
| Proc. Chnl. Descr.: | PDA Ch1 MaxPlot(210-400)nm | Column Temperature: | 40oC |
| Date Acquired: | 8/22/2024 11:19:41 AM CST | Date Processed: | 8/22/2024 12:41:59 PM |

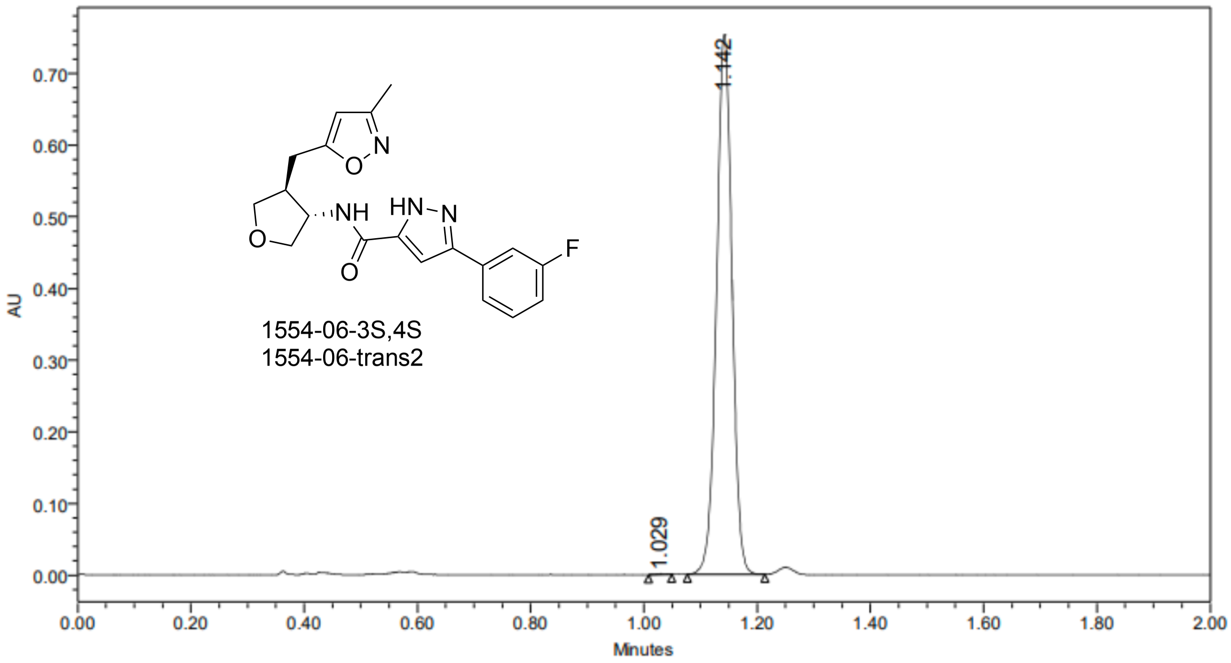


| | RT | Area | % Area | Height |
|---|---|---|---|---|
| 1 | 1.029 | 1853 | 0.13 | 1341 |
| 2 | 1.142 | 1436515 | 99.87 | 752949 |

#### SAMPLE INFORMATION

| | | | |
|---|---|---|---|
| Sample Name: | V5904-069-P2 | | |
| Acq. Method Set: | OJ_20%_B2 | Sample Set Name: | 20240926 |
| Co_Solvent: | EtOH[1%NH3(7M in MeOH)] | Processing Method: | VIVA CRO |
| Column Name: | OJ-3 4.6*100mm 3um | Injection #: | 1 |
| Vial: | 1:E,5 | Back Pressure: | 2000 psi |
| Flow Rate: | 3.0 mL/min | Injection Volume: | 10.00 ul |
| Channel Name: | PDA Ch1 MaxPlot(210-400)nm | Run Time: | 4.0 Minutes |
| Proc. Chnl. Descr.: | PDA Ch1 MaxPlot(210-400)nm | Column Temperature: | 40oC |
| Date Acquired: | 9/26/2024 4:39:53 PM CST | Date Processed: | 9/27/2024 9:29:30 AM CST |

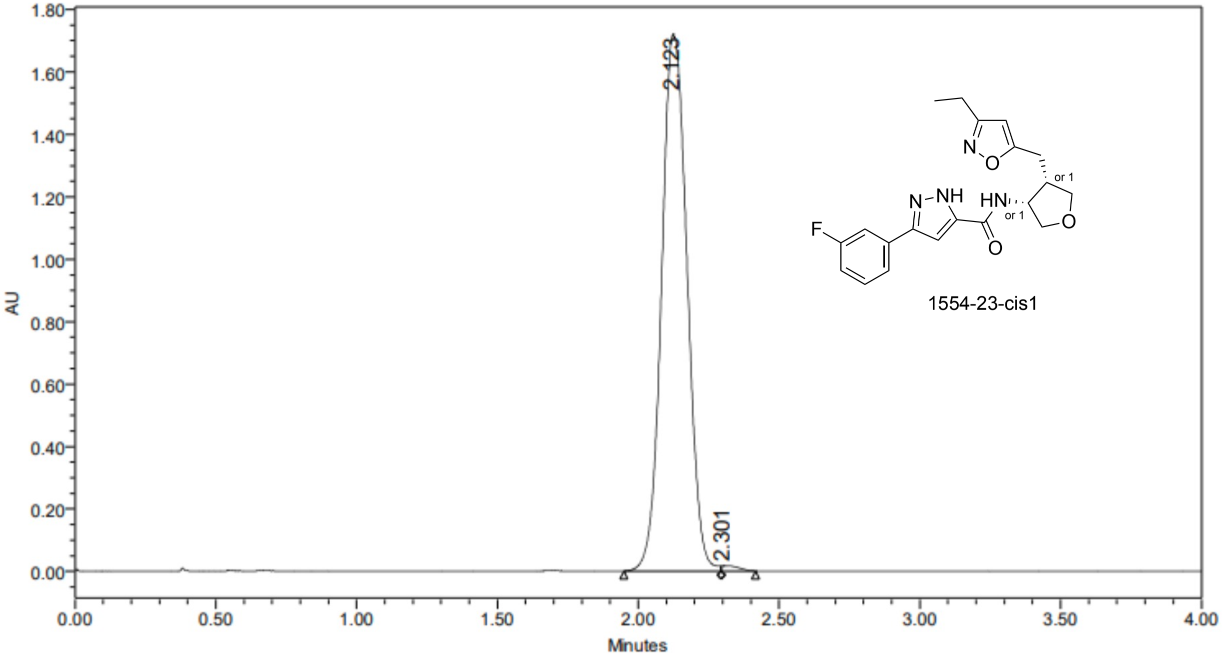


| | RT | Area | % Area | Height |
|---|---|---|---|---|
| 1 | 2.123 | 10813268 | 99.33 | 1720016 |
| 2 | 2.301 | 72570 | 0.67 | 17177 |

### SAMPLE INFORMATION

| | | | |
|---|---|---|---|
| Sample Name: | V5904-069-P4 | | |
| Acq. Method Set: | OJ_20%_B2 | Sample Set Name: | 20240926 |
| Co_Solvent: | EtOH[1%NH3(7M in MeOH)] | Processing Method: | VIVA CRO |
| Column Name: | OJ-3 4.6*100mm 3um | Injection #: | 1 |
| Vial: | 2:D,5 | Back Pressure: | 2000 psi |
| Flow Rate: | 3.0 mL/min | Injection Volume: | 10.00 ul |
| Channel Name: | PDA Ch1 MaxPlot(210-400)nm | Run Time: | 4.0 Minutes |
| Proc. Chnl. Descr.: | PDA Ch1 MaxPlot(210-400)nm | Column Temperature: | 40oC |
| Date Acquired: | 9/26/2024 4:18:27 PM CST | Date Processed: | 9/27/2024 9:28:40 AM CST |

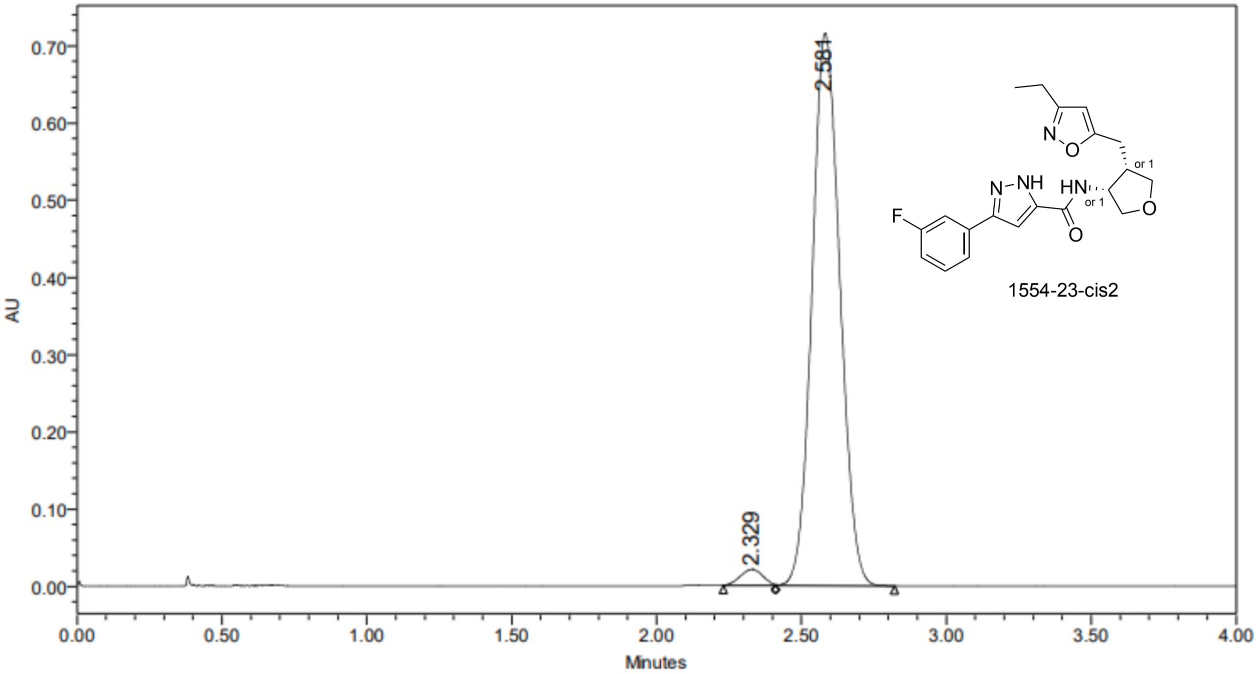


| | RT | Area | % Area | Height |
|---|---|---|---|---|
| 1 | 2.329 | 113814 | 2.33 | 21202 |
| 2 | 2.581 | 4778304 | 97.67 | 715086 |

### SAMPLE INFORMATION

| | | | |
|---|---|---|---|
| Sample Name: | V5904-069-P1 | | |
| Acq. Method Set: | OJ_20%_B2 | Sample Set Name: | 20240926 |
| Co_Solvent: | EtOH[1%NH3(7M in MeOH)] | Processing Method: | VIVA CRO |
| Column Name: | OJ-3 4.6*100mm 3um | Injection #: | 1 |
| Vial: | 1:E,4 | Back Pressure: | 2000 psi |
| Flow Rate: | 3.0 mL/min | Injection Volume: | 10.00 ul |
| Channel Name: | PDA Ch1 MaxPlot(210-400)nm | Run Time: | 4.0 Minutes |
| Proc. Chnl. Descr.: | PDA Ch1 MaxPlot(210-400)nm | Column Temperature: | 40oC |
| Date Acquired: | 9/26/2024 4:34:42 PM CST | Date Processed: | 9/27/2024 9:29:07 AM CST |

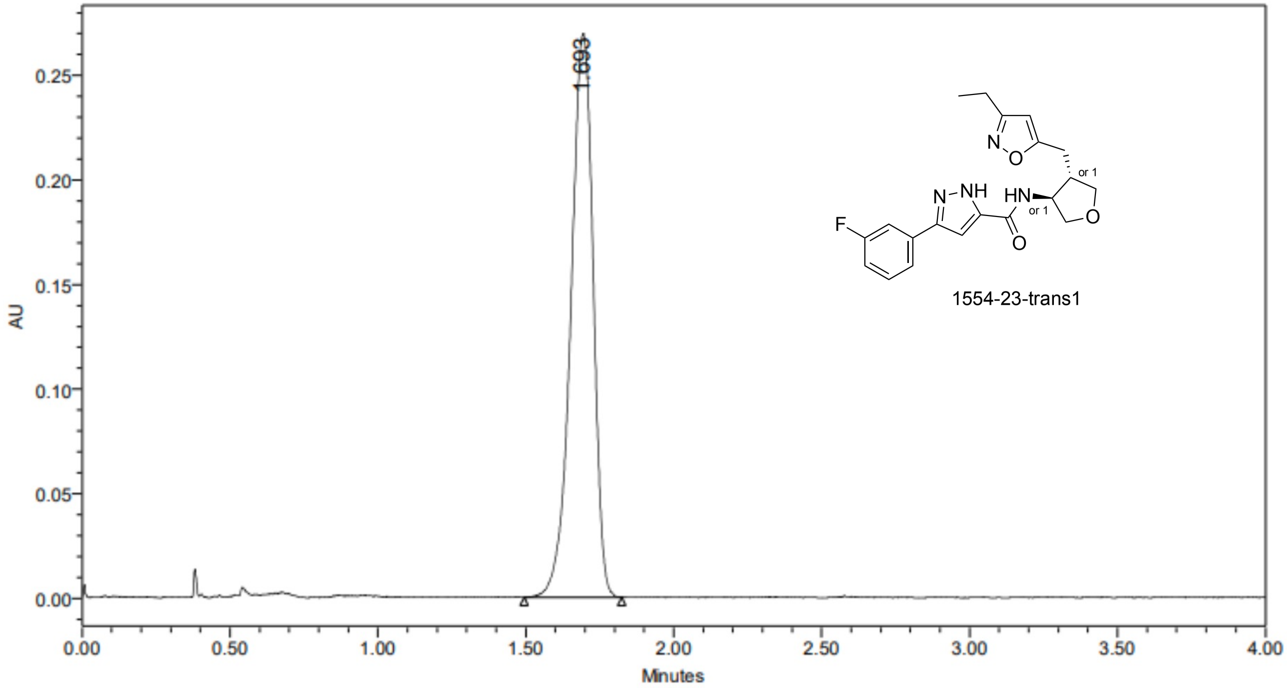


1554-23-trans1

| | RT | Area | % Area | Height |
|---|---|---|---|---|
| 1 | 1.693 | 1405119 | 100.00 | 269489 |

### SAMPLE INFORMATION

| | | | |
|---|---|---|---|
| Sample Name: | V5904-069-P3 | | |
| Acq. Method Set: | OJ_20%_B2 | Sample Set Name: | 20240926 |
| Co_Solvent: | EtOH[1%NH3(7M in MeOH)] | Processing Method: | VIVA CRO |
| Column Name: | OJ-3 4.6*100mm 3um | Injection #: | 1 |
| Vial: | 2:D,6 | Back Pressure: | 2000 psi |
| Flow Rate: | 3.0 mL/min | Injection Volume: | 10.00 ul |
| Channel Name: | PDA Ch1 MaxPlot(210-400)nm | Run Time: | 4.0 Minutes |
| Proc. Chnl. Descr.: | PDA Ch1 MaxPlot(210-400)nm | Column Temperature: | 40oC |
| Date Acquired: | 9/26/2024 4:29:31 PM CST | Date Processed: | 9/27/2024 9:28:07 AM CST |

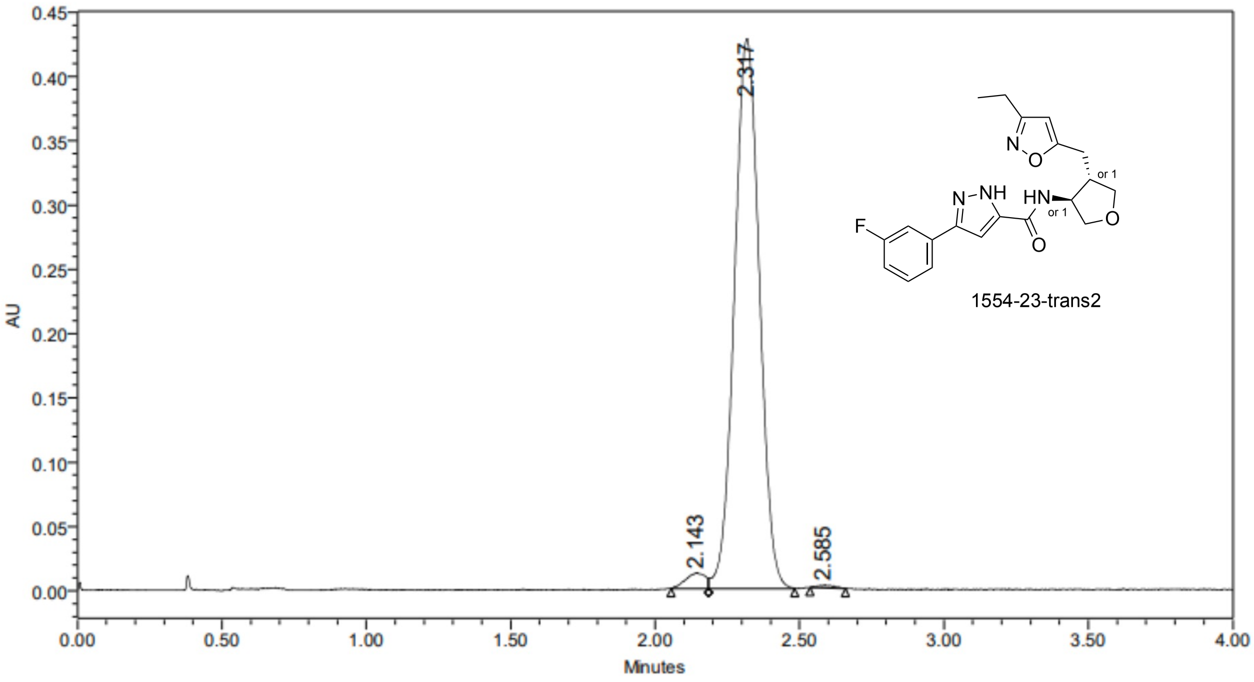


| | RT | Area | % Area | Height |
|---|---|---|---|---|
| 1 | 2.143 | 56989 | 2.16 | 12133 |
| 2 | 2.317 | 2576735 | 97.51 | 427135 |
| 3 | 2.585 | 8861 | 0.34 | 2061 |

### SAMPLE INFORMATION

| | | | |
|---|---|---|---|
| Sample Name: | V5904-069-P2 | | |
| Acq. Method Set: | OJ_20%_B2 | Sample Set Name: | 20240926 |
| Co_Solvent: | EtOH[1%NH3(7M in MeOH)] | Processing Method: | VIVA CRO |
| Column Name: | OJ-3 4.6*100mm 3um | Injection #: | 1 |
| Vial: | 1:E,5 | Back Pressure: | 2000 psi |
| Flow Rate: | 3.0 mL/min | Injection Volume: | 10.00 ul |
| Channel Name: | PDA Ch1 MaxPlot(210-400)nm | Run Time: | 4.0 Minutes |
| Proc. Chnl. Descr.: | PDA Ch1 MaxPlot(210-400)nm | Column Temperature: | 40oC |
| Date Acquired: | 9/26/2024 4:39:53 PM CST | Date Processed: | 9/27/2024 9:29:30 AM CST |

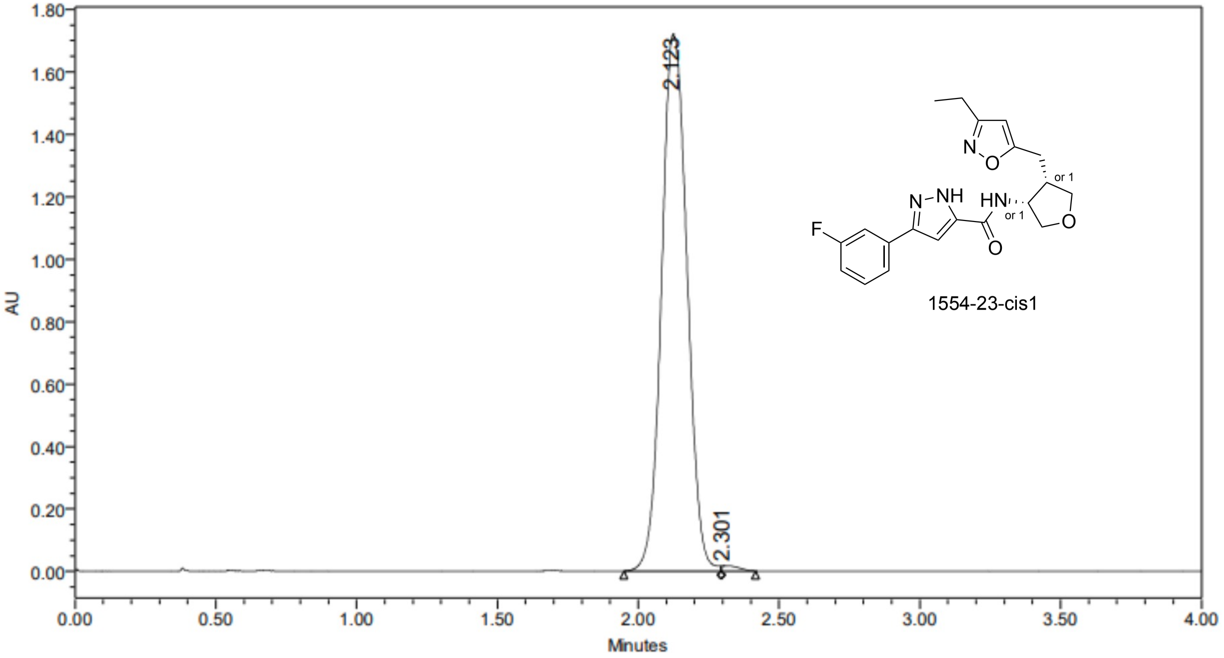


| | RT | Area | % Area | Height |
|---|---|---|---|---|
| 1 | 2.123 | 10813268 | 99.33 | 1720016 |
| 2 | 2.301 | 72570 | 0.67 | 17177 |

### SAMPLE INFORMATION

| | | | |
|---|---|---|---|
| Sample Name: | V5904-069-P4 | | |
| Acq. Method Set: | OJ_20%_B2 | Sample Set Name: | 20240926 |
| Co_Solvent: | EtOH[1%NH3(7M in MeOH)] | Processing Method: | VIVA CRO |
| Column Name: | OJ-3 4.6*100mm 3um | Injection #: | 1 |
| Vial: | 2:D,5 | Back Pressure: | 2000 psi |
| Flow Rate: | 3.0 mL/min | Injection Volume: | 10.00 ul |
| Channel Name: | PDA Ch1 MaxPlot(210-400)nm | Run Time: | 4.0 Minutes |
| Proc. Chnl. Descr.: | PDA Ch1 MaxPlot(210-400)nm | Column Temperature: | 40oC |
| Date Acquired: | 9/26/2024 4:18:27 PM CST | Date Processed: | 9/27/2024 9:28:40 AM CST |

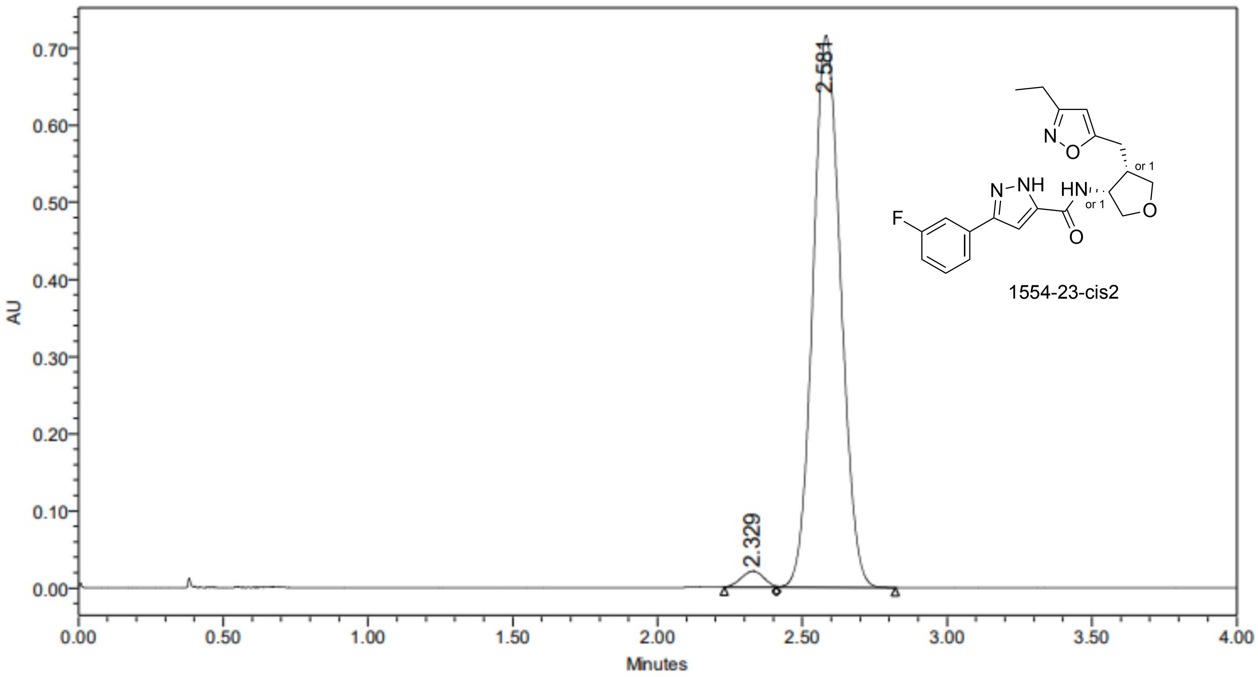


| | RT | Area | % Area | Height |
|---|---|---|---|---|
| 1 | 2.329 | 113814 | 2.33 | 21202 |
| 2 | 2.581 | 4778304 | 97.67 | 715086 |

### SAMPLE INFORMATION

| | | | |
|---|---|---|---|
| Sample Name: | V5904-069-P1 | | |
| Acq. Method Set: | OJ_20%_B2 | Sample Set Name: | 20240926 |
| Co_Solvent: | EtOH[1%NH3(7M in MeOH)] | Processing Method: | VIVA CRO |
| Column Name: | OJ-3 4.6*100mm 3um | Injection #: | 1 |
| Vial: | 1:E,4 | Back Pressure: | 2000 psi |
| Flow Rate: | 3.0 mL/min | Injection Volume: | 10.00 ul |
| Channel Name: | PDA Ch1 MaxPlot(210-400)nm | Run Time: | 4.0 Minutes |
| Proc. Chnl. Descr.: | PDA Ch1 MaxPlot(210-400)nm | Column Temperature: | 40oC |
| Date Acquired: | 9/26/2024 4:34:42 PM CST | Date Processed: | 9/27/2024 9:29:07 AM CST |

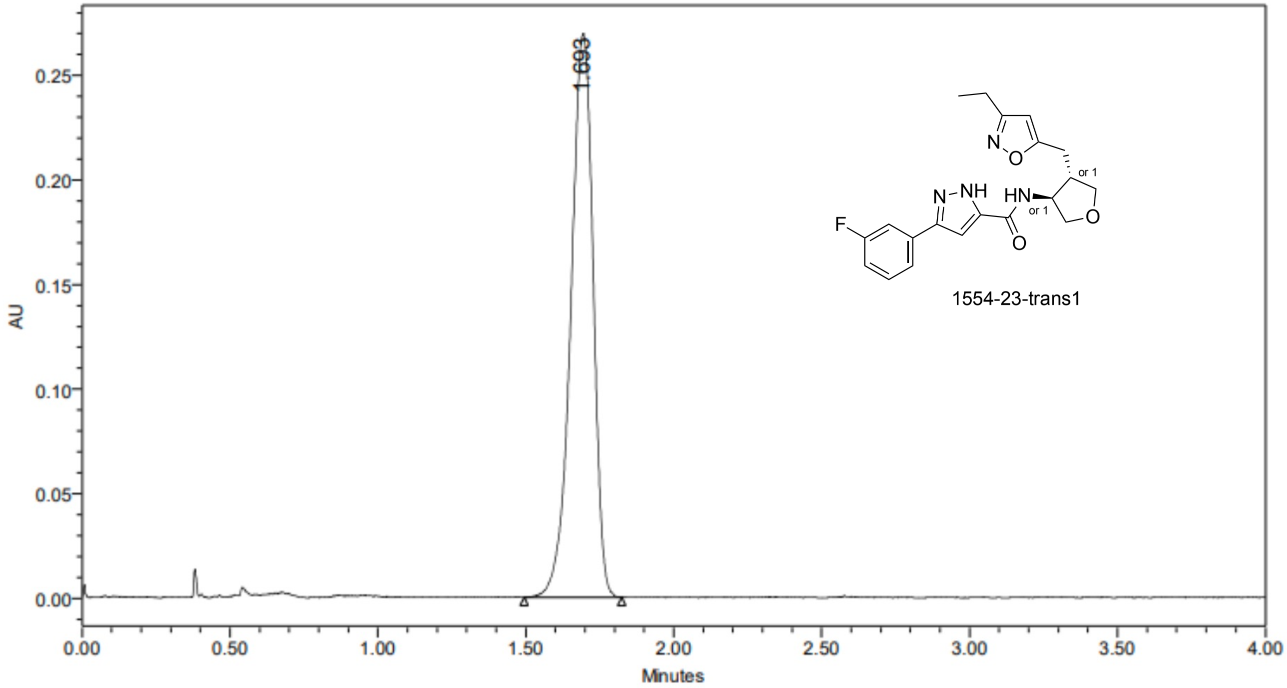


| | RT | Area | % Area | Height |
|---|---|---|---|---|
| 1 | 1.693 | 1405119 | 100.00 | 269489 |

### SAMPLE INFORMATION

| | | | |
|---|---|---|---|
| Sample Name: | V5904-069-P3 | | |
| Acq. Method Set: | OJ_20%_B2 | Sample Set Name: | 20240926 |
| Co_Solvent: | EtOH[1%NH3(7M in MeOH)] | Processing Method: | VIVA CRO |
| Column Name: | OJ-3 4.6*100mm 3um | Injection #: | 1 |
| Vial: | 2:D,6 | Back Pressure: | 2000 psi |
| Flow Rate: | 3.0 mL/min | Injection Volume: | 10.00 ul |
| Channel Name: | PDA Ch1 MaxPlot(210-400)nm | Run Time: | 4.0 Minutes |
| Proc. Chnl. Descr.: | PDA Ch1 MaxPlot(210-400)nm | Column Temperature: | 40oC |
| Date Acquired: | 9/26/2024 4:29:31 PM CST | Date Processed: | 9/27/2024 9:28:07 AM CST |

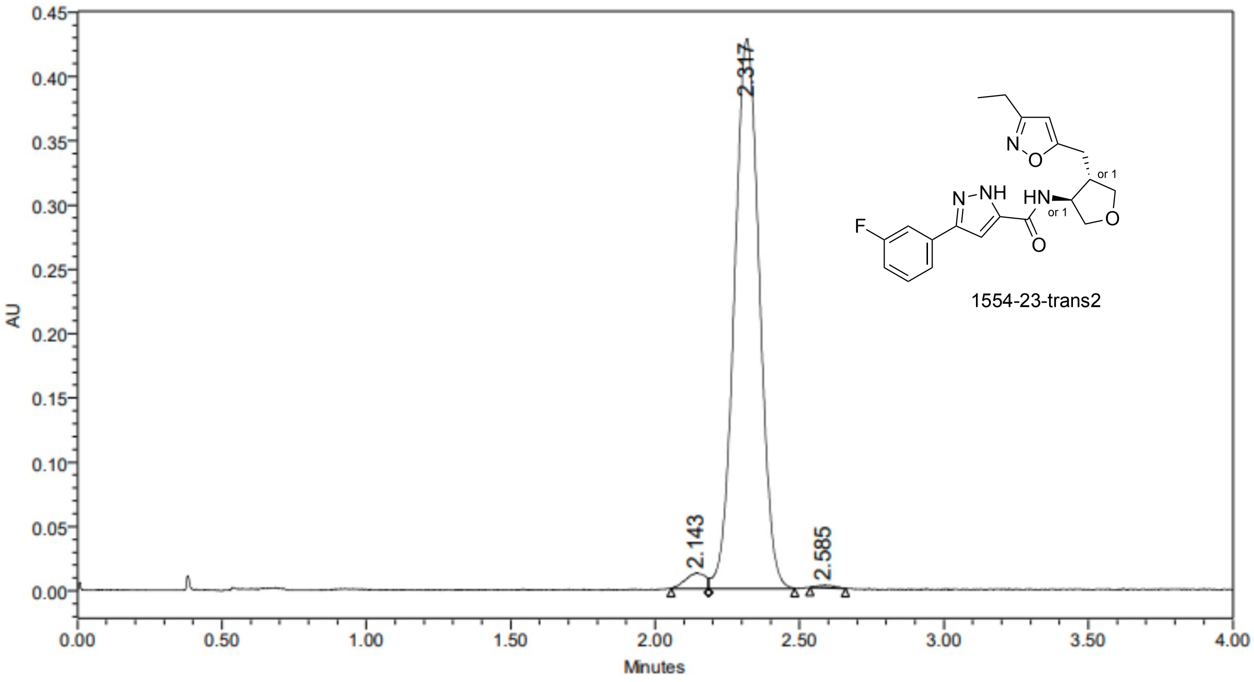


| | RT | Area | % Area | Height |
|---|---|---|---|---|
| 1 | 2.143 | 56989 | 2.16 | 12133 |
| 2 | 2.317 | 2576735 | 97.51 | 427135 |
| 3 | 2.585 | 8861 | 0.34 | 2061 |

### SAMPLE INFORMATION

| | | | |
|---|---|---|---|
| Sample Name: | V5904-028-P1 | | |
| Acq. Method Set: | OJ_15%_B1 | Sample Set Name: | 20240812 |
| Co_Solvent: | MeOH[0.2%NH3(7M in MeOH)] | Processing Method: | VIVA CRO |
| Column Name: | OJ-3 4.6*100mm 3um | Injection #: | 1 |
| Vial: | 2:E,2 | Back Pressure: | 2000 psi |
| Flow Rate: | 3.0 mL/min | Injection Volume: | 10.00 ul |
| Channel Name: | PDA Ch1 MaxPlot(210-400)nm | Run Time: | 6.0 Minutes |
| Proc. Chnl. Descr.: | PDA Ch1 MaxPlot(210-400)nm | Column Temperature: | 40oC |
| Date Acquired: | 8/12/2024 10:54:50 AM CST | Date Processed: | 8/14/2024 1:24:34 PM CST |

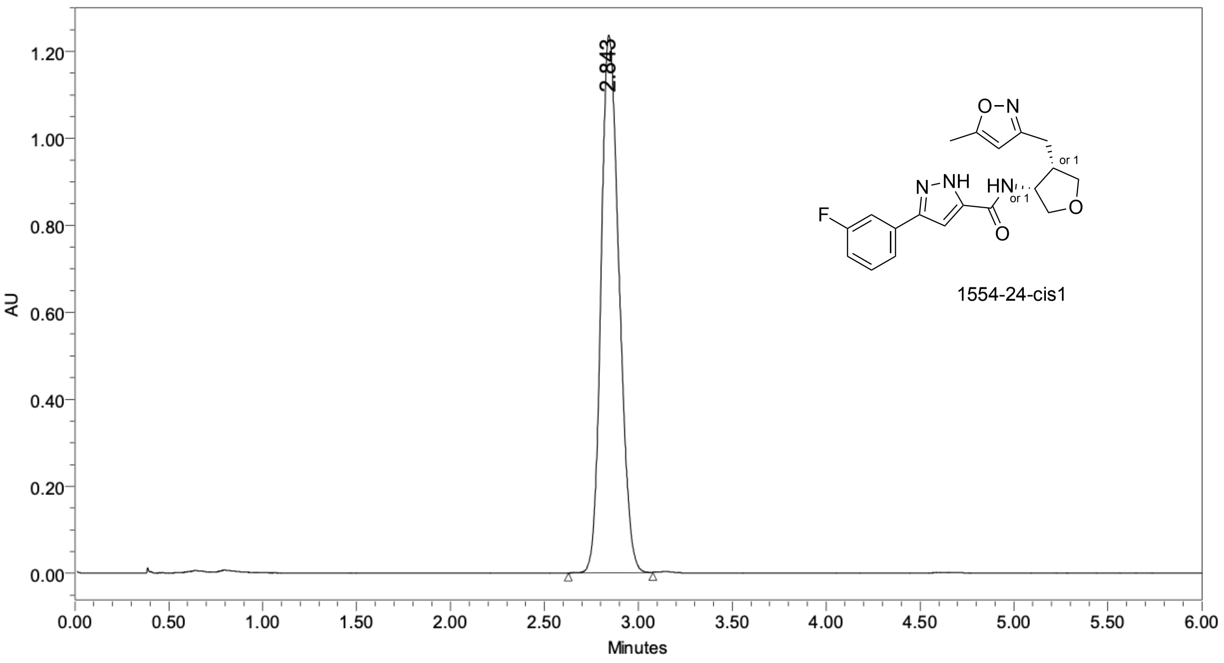


| | RT | Area | % Area | Height |
|---|---|---|---|---|
| 1 | 2.843 | 8253236 | 100.00 | 1236344 |

### SAMPLE INFORMATION

| | | | |
|---|---|---|---|
| Sample Name: | V5904-028-P4 | | |
| Acq. Method Set: | OJ_15%_B1 | Sample Set Name: | 20240812 |
| Co_Solvent: | MeOH[0.2%NH3(7M in MeOH)] | Processing Method: | VIVA CRO |
| Column Name: | OJ-3 4.6*100mm 3um | Injection #: | 1 |
| Vial: | 2:E,5 | Back Pressure: | 2000 psi |
| Flow Rate: | 3.0 mL/min | Injection Volume: | 10.00 ul |
| Channel Name: | PDA Ch1 MaxPlot(210-400)nm | Run Time: | 6.0 Minutes |
| Proc. Chnl. Descr.: | PDA Ch1 MaxPlot(210-400)nm | Column Temperature: | 40oC |
| Date Acquired: | 8/12/2024 11:16:22 AM CST | Date Processed: | 8/14/2024 1:27:27 PM CST |

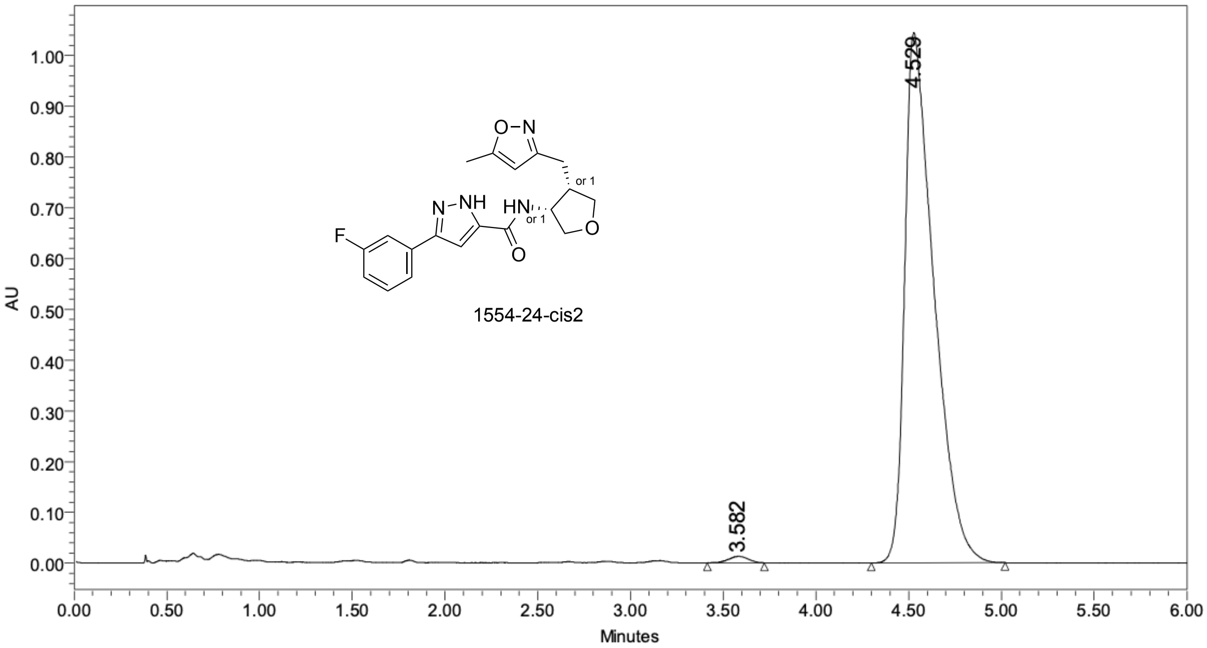


| | RT | Area | % Area | Height |
|---|---|---|---|---|
| 1 | 3.582 | 94228 | 0.82 | 12778 |
| 2 | 4.529 | 11432220 | 99.18 | 1044821 |

### SAMPLE INFORMATION

| | | | |
|---|---|---|---|
| Sample Name: | V5904-028-P2 | | |
| Acq. Method Set: | OJ_15%_B1 | Sample Set Name: | 20240812 |
| Co_Solvent: | MeOH[0.2%NH3(7M in MeOH)] | Processing Method: | VIVA CRO |
| Column Name: | OJ-3 4.6*100mm 3um | Injection #: | 1 |
| Vial: | 2:E,3 | Back Pressure: | 2000 psi |
| Flow Rate: | 3.0 mL/min | Injection Volume: | 10.00 ul |
| Channel Name: | PDA Ch1 MaxPlot(210-400)nm | Run Time: | 6.0 Minutes |
| Proc. Chnl. Descr.: | PDA Ch1 MaxPlot(210-400)nm | Column Temperature: | 40oC |
| Date Acquired: | 8/12/2024 11:02:01 AM CST | Date Processed: | 8/15/2024 10:55:51 AM |

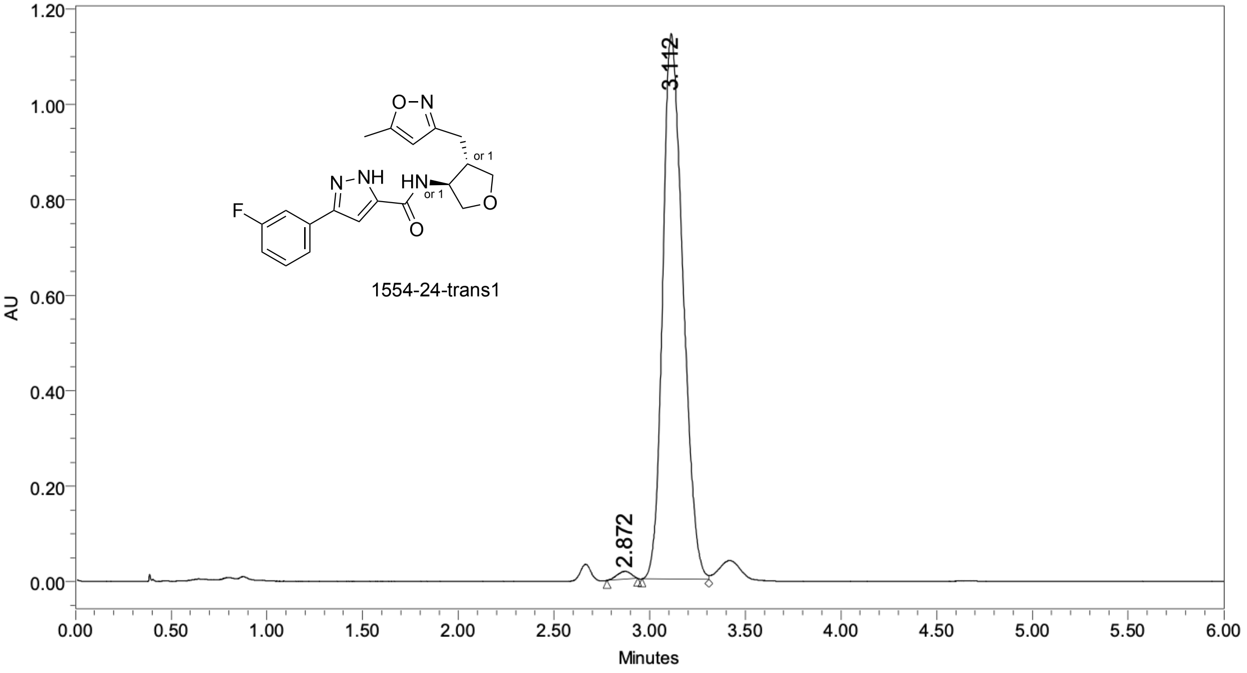


| | RT | Area | % Area | Height |
|---|---|---|---|---|
| 1 | 2.872 | 83795 | 0.99 | 16494 |
| 2 | 3.112 | 8395370 | 99.01 | 1143312 |

### SAMPLE INFORMATION

| | | | |
|---|---|---|---|
| Sample Name: | V5904-028-P3 | | |
| Acq. Method Set: | OJ_15%_B1 | Sample Set Name: | 20240812 |
| Co_Solvent: | MeOH[0.2%NH3(7M in MeOH)] | Processing Method: | VIVA CRO |
| Column Name: | OJ-3 4.6*100mm 3um | Injection #: | 1 |
| Vial: | 2:E,4 | Back Pressure: | 2000 psi |
| Flow Rate: | 3.0 mL/min | Injection Volume: | 10.00 ul |
| Channel Name: | PDA Ch1 MaxPlot(210-400)nm | Run Time: | 6.0 Minutes |
| Proc. Chnl. Descr.: | PDA Ch1 MaxPlot(210-400)nm | Column Temperature: | 40oC |
| Date Acquired: | 8/12/2024 11:09:12 AM CST | Date Processed: | 8/14/2024 1:27:47 PM CST |

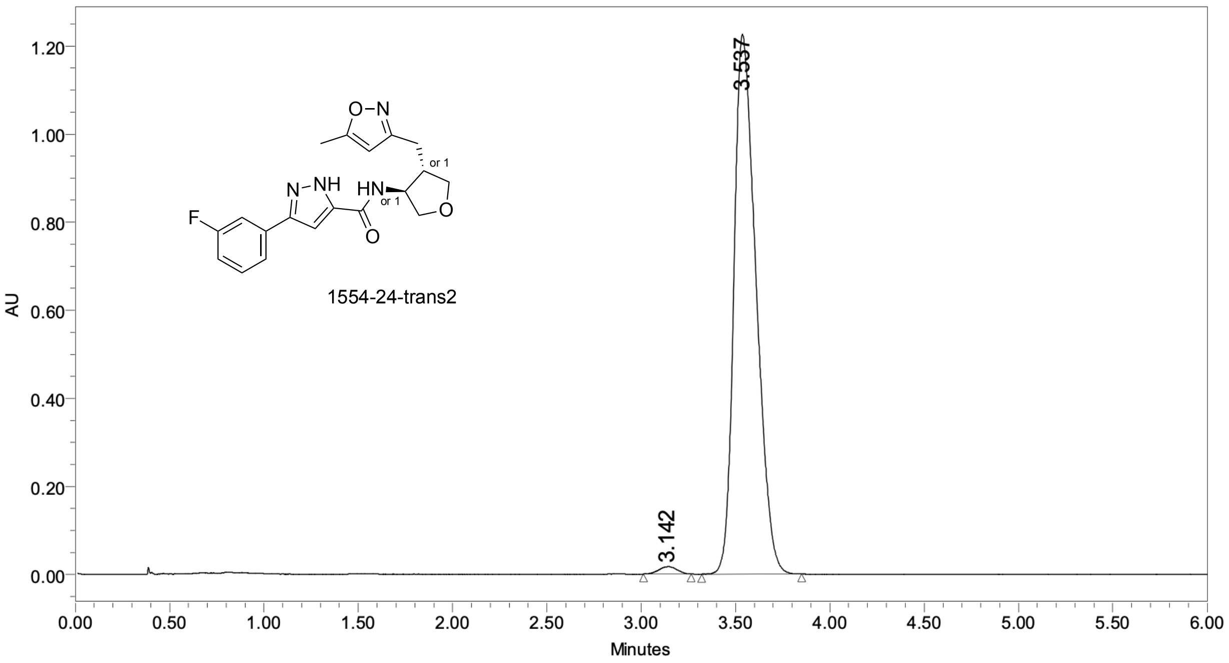


| | RT | Area | % Area | Height |
|---|---|---|---|---|
| 1 | 3.142 | 115349 | 1.15 | 17143 |
| 2 | 3.537 | 9933678 | 98.85 | 1226528 |

### SAMPLE INFORMATION

| | | | |
|---|---|---|---|
| Sample Name: | V5899-048-P1 | | |
| Acq. Method Set: | AD_55%_B2 | Sample Set Name: | 20240929 |
| Co_Solvent: | EtOH[1%NH3(7M in MeOH)] | Processing Method: | VIVA CRO |
| Column Name: | AD-3 4.6*100mm 3um | Injection #: | 1 |
| Vial: | 2:F,5 | Back Pressure: | 2000 psi |
| Flow Rate: | 3.0 mL/min | Injection Volume: | 10.00 ul |
| Channel Name: | PDA Ch1 MaxPlot(210-400)nm | Run Time: | 4.0 Minutes |
| Proc. Chnl. Descr.: | PDA Ch1 MaxPlot(210-400)nm | Column Temperature: | 40oC |
| Date Acquired: | 9/29/2024 2:42:08 PM CST | Date Processed: | 9/29/2024 3:14:39 PM CST |

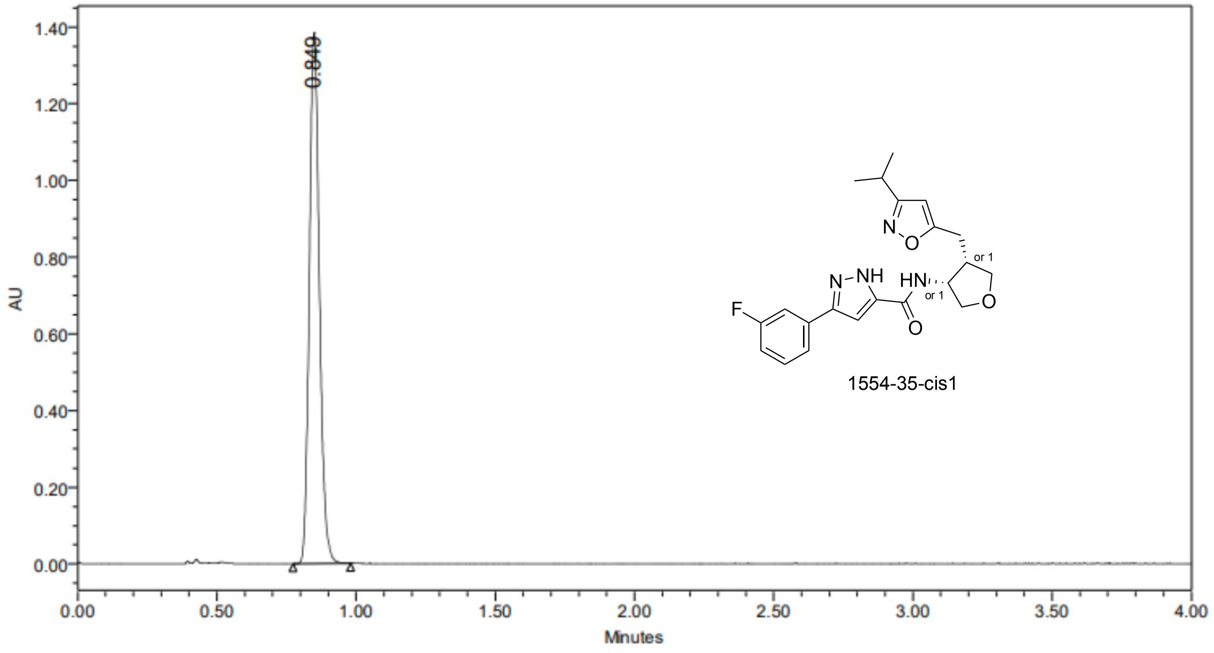


| | RT | Area | % Area | Height |
|---|---|---|---|---|
| 1 | 0.849 | 3471302 | 100.00 | 1383980 |

### SAMPLE INFORMATION

| | | | |
|---|---|---|---|
| Sample Name: | V5899-048-P2 | | |
| Acq. Method Set: | AD_55%_B2 | Sample Set Name: | 20240929 |
| Co_Solvent: | EtOH[1%NH3(7M in MeOH)] | Processing Method: | VIVA CRO |
| Column Name: | AD-3 4.6*100mm 3um | Injection #: | 1 |
| Vial: | 2:F,6 | Back Pressure: | 2000 psi |
| Flow Rate: | 3.0 mL/min | Injection Volume: | 10.00 ul |
| Channel Name: | PDA Ch1 MaxPlot(210-400)nm | Run Time: | 4.0 Minutes |
| Proc. Chnl. Descr.: | PDA Ch1 MaxPlot(210-400)nm | Column Temperature: | 40oC |
| Date Acquired: | 9/29/2024 2:47:20 PM CST | Date Processed: | 9/29/2024 3:14:55 PM CST |

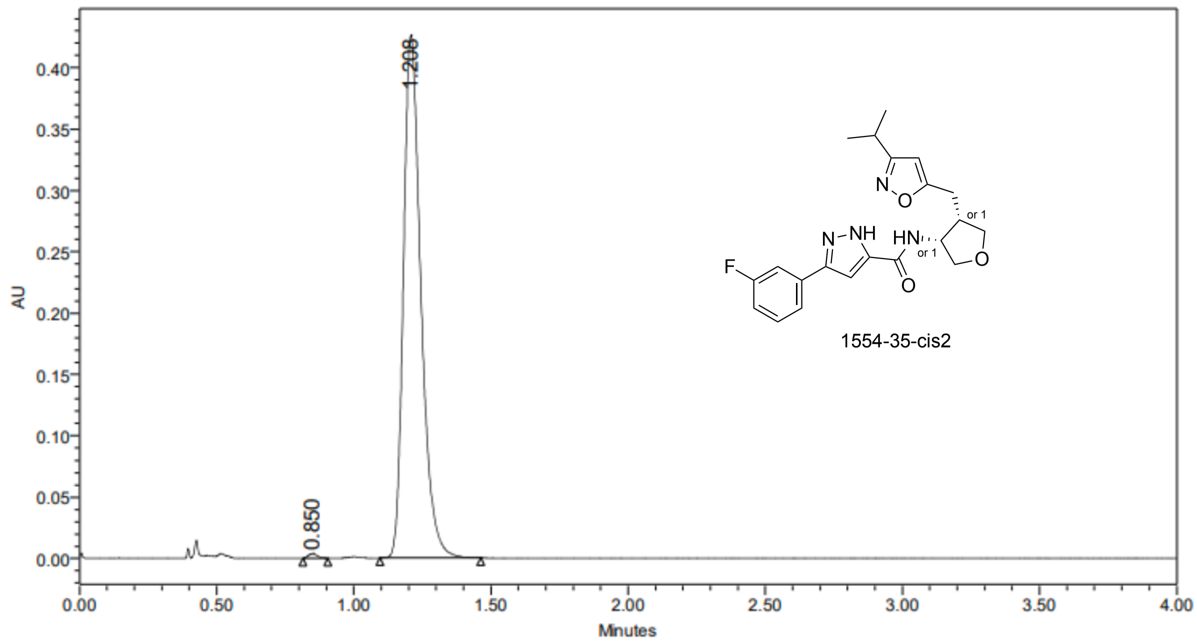


| | RT | Area | % Area | Height |
|---|---|---|---|---|
| 1 | 0.850 | 8293 | 0.44 | 3708 |
| 2 | 1.208 | 1885287 | 99.56 | 426151 |

### SAMPLE INFORMATION

| | | | |
|---|---|---|---|
| Sample Name: | V5899-048-P3 | | |
| Acq. Method Set: | AD_55%_B2 | Sample Set Name: | 20240930 |
| Co_Solvent: | EtOH[1%NH3(7M in MeOH)] | Processing Method: | VIVA CRO |
| Column Name: | AD-3 4.6*100mm 3um | Injection #: | 1 |
| Vial: | 2:A,6 | Back Pressure: | 2000 psi |
| Flow Rate: | 3.0 mL/min | Injection Volume: | 10.00 ul |
| Channel Name: | PDA Ch1 MaxPlot(210-400)nm | Run Time: | 4.0 Minutes |
| Proc. Chnl. Descr.: | PDA Ch1 MaxPlot(210-400)nm | Column Temperature: | 40oC |
| Date Acquired: | 9/30/2024 10:49:19 AM CST | Date Processed: | 9/30/2024 10:57:08 AM |

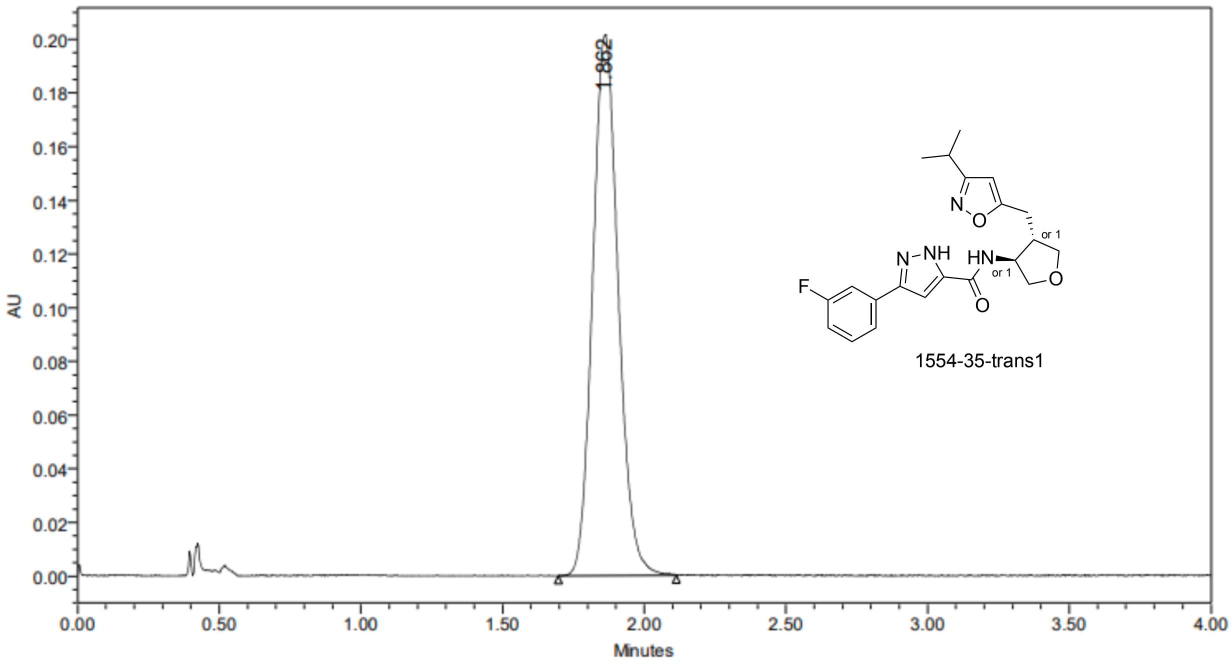


| | RT | Area | % Area | Height |
|---|---|---|---|---|
| 1 | 1.862 | 1263976 | 100.00 | 201346 |

### SAMPLE INFORMATION

| | | | |
|---|---|---|---|
| Sample Name: | V5899-048-P3 | | |
| Acq. Method Set: | AD_55%_B2 | Sample Set Name: | 20240930 |
| Co_Solvent: | EtOH[1%NH3(7M in MeOH)] | Processing Method: | VIVA CRO |
| Column Name: | AD-3 4.6*100mm 3um | Injection #: | 1 |
| Vial: | 2:A,6 | Back Pressure: | 2000 psi |
| Flow Rate: | 3.0 mL/min | Injection Volume: | 10.00 ul |
| Channel Name: | PDA Ch1 MaxPlot(210-400)nm | Run Time: | 4.0 Minutes |
| Proc. Chnl. Descr.: | PDA Ch1 MaxPlot(210-400)nm | Column Temperature: | 40oC |
| Date Acquired: | 9/30/2024 10:49:19 AM CST | Date Processed: | 9/30/2024 10:57:08 AM |

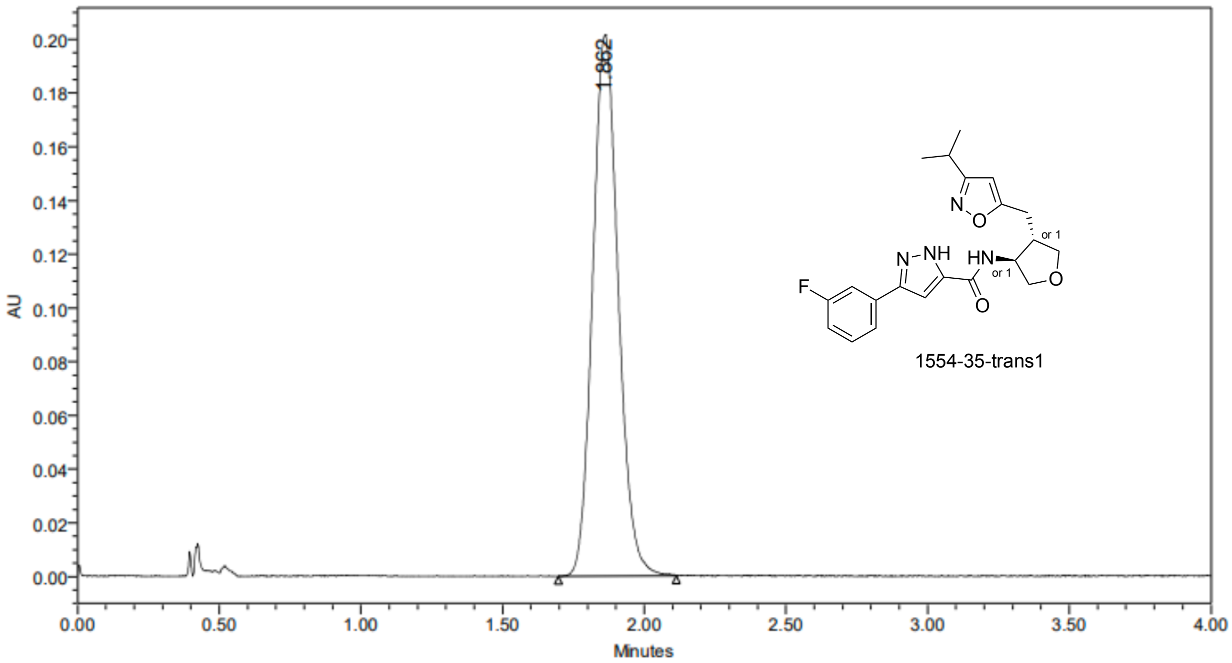


| | RT | Area | % Area | Height |
|---|---|---|---|---|
| 1 | 1.862 | 1263976 | 100.00 | 201346 |

### SAMPLE INFORMATION

| | | | |
|---|---|---|---|
| Sample Name: | V5899-048-P4 | | |
| Acq. Method Set: | AD_55%_B2 | Sample Set Name: | 20240930 |
| Co_Solvent: | EtOH[1%NH3(7M in MeOH)] | Processing Method: | VIVA CRO |
| Column Name: | AD-3 4.6*100mm 3um | Injection #: | 1 |
| Vial: | 2:A,3 | Back Pressure: | 2000 psi |
| Flow Rate: | 3.0 mL/min | Injection Volume: | 10.00 ul |
| Channel Name: | PDA Ch1 MaxPlot(210-400)nm | Run Time: | 4.0 Minutes |
| Proc. Chnl. Descr.: | PDA Ch1 MaxPlot(210-400)nm | Column Temperature: | 40oC |
| Date Acquired: | 9/30/2024 9:11:07 AM CST | Date Processed: | 9/30/2024 10:58:31 AM |

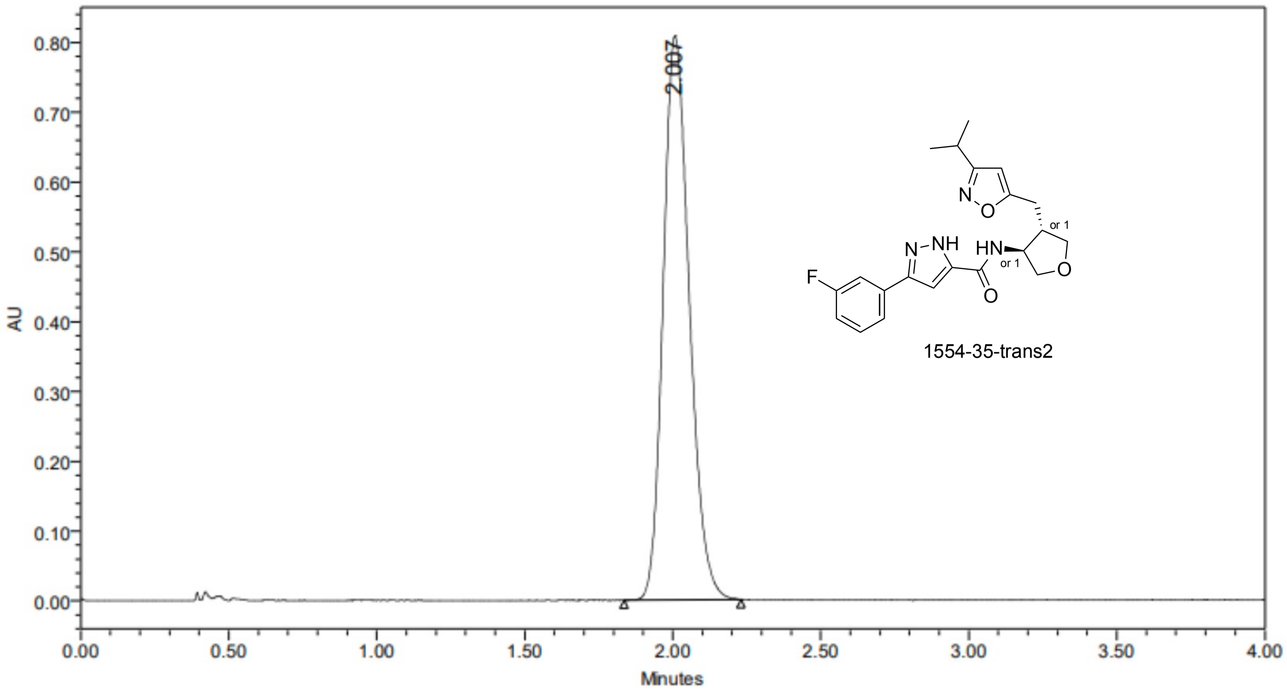


| | RT | Area | % Area | Height |
|---|---|---|---|---|
| 1 | 2.007 | 4959872 | 100.00 | 808274 |

### SAMPLE INFORMATION

| | | | |
|---|---|---|---|
| Sample Name: | V5899-057-P1 | | |
| Acq. Method Set: | AD_55%_B2 | Sample Set Name: | 20241025 |
| Co_Solvent: | EtOH[1%NH3(7M in MeOH)] | Processing Method: | VIVA CRO |
| Column Name: | AD-3 4.6*100mm 3um | Injection #: | 1 |
| Vial: | 2:A,5 | Back Pressure: | 2000 psi |
| Flow Rate: | 3.0 mL/min | Injection Volume: | 8.00 ul |
| Channel Name: | 254.0nm | Run Time: | 7.0 Minutes |
| Proc. Chnl. Descr.: | PDA Spectrum PDA 254.0 nm (PDA | Column Temperature: | 40oC |
| Date Acquired: | 10/25/2024 10:31:38 AM CST | Date Processed: | 10/28/2024 1:51:46 PM |

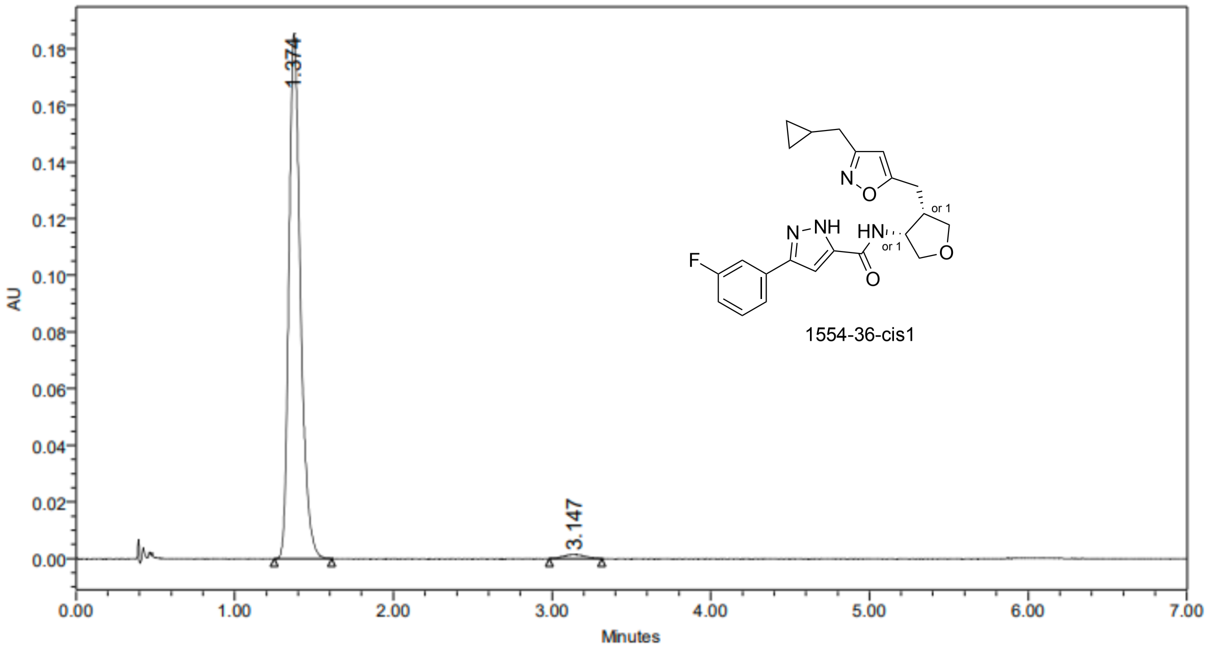


| | RT | Area | % Area | Height |
|---|---|---|---|---|
| 1 | 1.374 | 941006 | 98.57 | 185361 |
| 2 | 3.147 | 13628 | 1.43 | 1430 |

### SAMPLE INFORMATION

| | | | |
|---|---|---|---|
| Sample Name: | V5899-057-P2 | | |
| Acq. Method Set: | AD_55%_B2 | Sample Set Name: | 20241025 |
| Co_Solvent: | EtOH[1%NH3(7M in MeOH)] | Processing Method: | VIVA CRO |
| Column Name: | AD-3 4.6*100mm 3um | Injection #: | 1 |
| Vial: | 2:C,1 | Back Pressure: | 2000 psi |
| Flow Rate: | 3.0 mL/min | Injection Volume: | 10.00 ul |
| Channel Name: | 254.0nm | Run Time: | 8.0 Minutes |
| Proc. Chnl. Descr.: | PDA Spectrum PDA 254.0 nm (PDA | Column Temperature: | 40oC |
| Date Acquired: | 10/26/2024 3:59:51 AM CST | Date Processed: | 10/28/2024 2:00:29 PM |

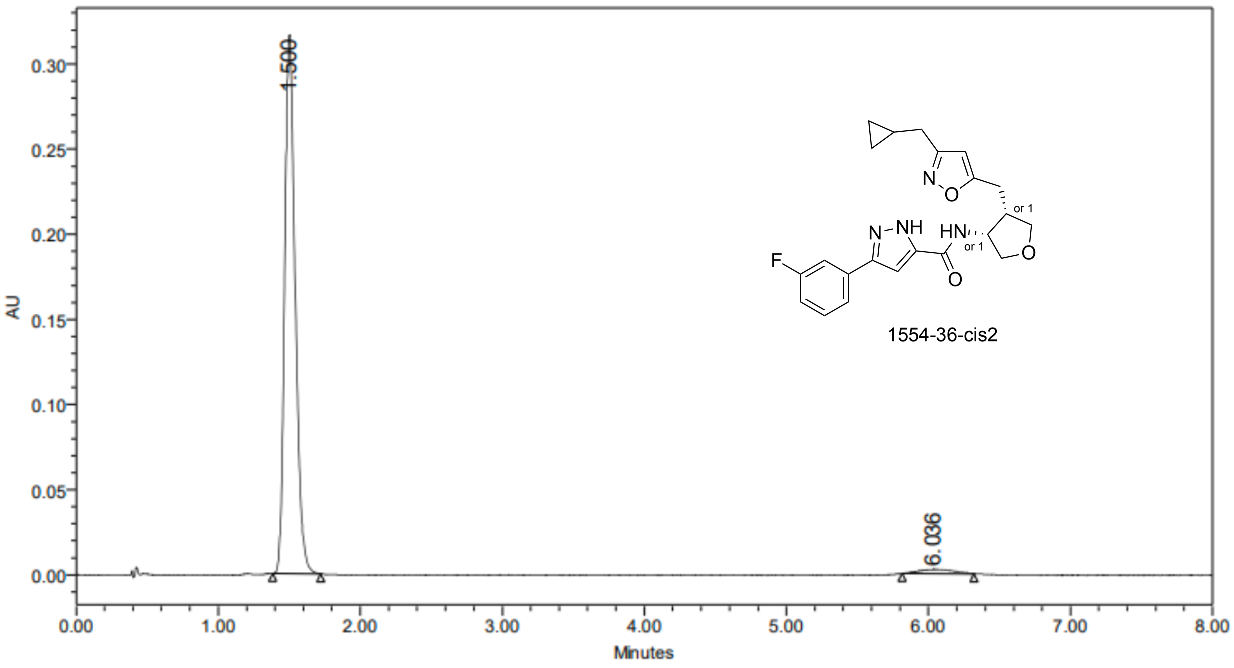


| | RT | Area | % Area | Height |
|---|---|---|---|---|
| 1 | 1.500 | 1634656 | 97.53 | 316110 |
| 2 | 6.036 | 41465 | 2.47 | 2451 |

### SAMPLE INFORMATION

| | | | |
|---|---|---|---|
| Sample Name: | V5899-057-P3 | | |
| Acq. Method Set: | AD_55%_B2 | Sample Set Name: | 20241025 |
| Co_Solvent: | EtOH[1%NH3(7M in MeOH)] | Processing Method: | VIVA CRO |
| Column Name: | AD-3 4.6*100mm 3um | Injection #: | 1 |
| Vial: | 2:A,4 | Back Pressure: | 2000 psi |
| Flow Rate: | 3.0 mL/min | Injection Volume: | 8.00 ul |
| Channel Name: | 254.0nm | Run Time: | 7.0 Minutes |
| Proc. Chnl. Descr.: | PDA Spectrum PDA 254.0 nm (PDA | Column Temperature: | 40oC |
| Date Acquired: | 10/25/2024 10:23:32 AM CST | Date Processed: | 10/28/2024 1:51:11 PM |

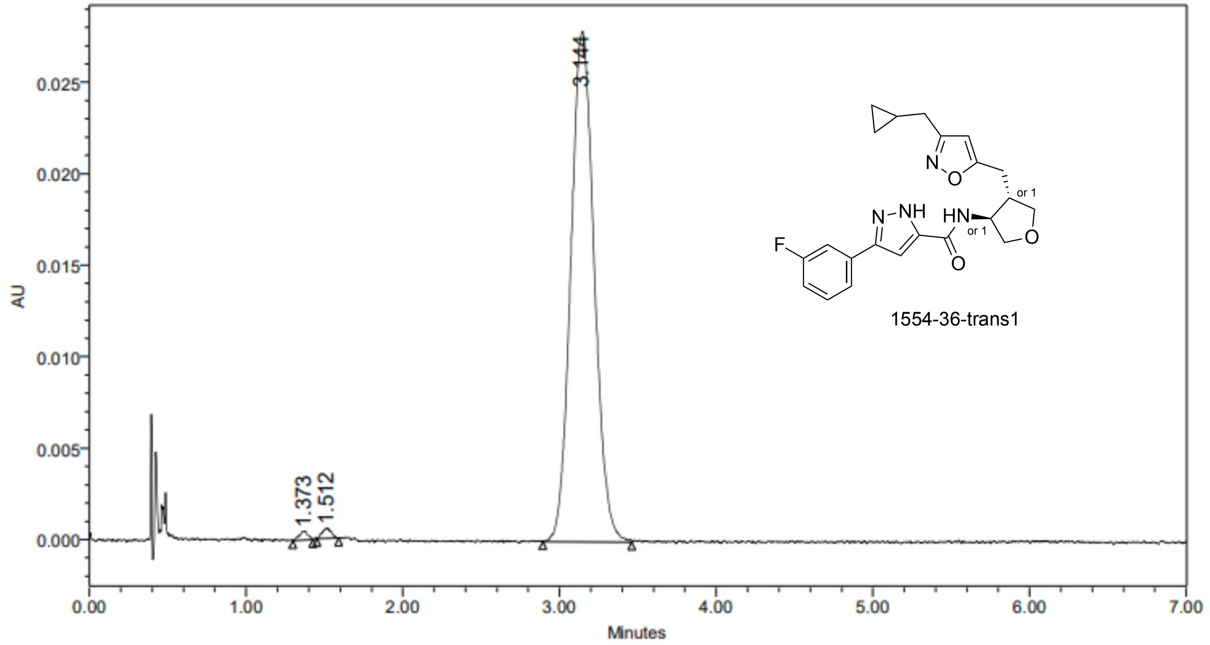


1554-36-trans1

| | RT | Area | % Area | Height |
|---|---|---|---|---|
| 1 | 1.373 | 1838 | 0.63 | 479 |
| 2 | 1.512 | 2313 | 0.79 | 552 |
| 3 | 3.144 | 289024 | 98.58 | 27870 |
